# Multimodal brain imaging study of 36,678 participants reveals adverse effects of moderate drinking

**DOI:** 10.1101/2020.03.27.011791

**Authors:** Remi Daviet, Gökhan Aydogan, Kanchana Jagannathan, Nathaniel Spilka, Philipp D. Koellinger, Henry R. Kranzler, Gideon Nave, Reagan R. Wetherill

**Author notes:** Corresponding Authors: Reagan R. Wetherill, Department of Psychiatry, 3535 Market Street, Suite 500, Philadelphia, PA 19104 USA, Gideon Nave, The Wharton School, University of Pennsylvania, 3730 Walnut St., Philadelphia, PA 19104 USA.

## Abstract

Heavy alcohol consumption can have significant deleterious neural consequences, including brain atrophy, neuronal loss, poorer white matter fiber integrity, and cognitive decline. However, the effects of light-to-moderate alcohol consumption on brain structure remain unclear. Here, we examine the associations between alcohol intake and brain structure using multimodal imaging data from 36,678 generally healthy middle-aged and older adults from the UK Biobank, controlling for numerous potential confounds. We find negative associations between alcohol intake and global gray matter volume (GMV) and white matter volume (WMV), which become stronger as intake increases. An examination of the associations between alcohol intake and 139 regional GMV imaging-derived phenotypes (IDPs) and 375 WM microstructure IDPs yielded 304 (59.1%) significant findings, including 125 GMV IDPs that are spread across the brain and 179 WM microstructure IDPs across multiple tract regions. In general, findings comport with the existing literature. However, a daily alcohol intake of as little as one to two units – 250 to 500 ml of a 4% beer or 76 to 146 ml of a 13% wine – is already associated with GMV deficits and altered WMV microstructure, placing moderate drinkers at risk.

**One Sentence Summary:** Moderate alcohol intake, consuming one or more daily alcohol units, has adverse effects on brain health.

---

Alcohol consumption is one of the leading contributors to the global burden of disease and to high healthcare and economic costs. Alcoholism, now diagnosed as Alcohol Use Disorder (AUD)^1^, is one of the most prevalent mental conditions worldwide^2^, with harmful effects on physical, cognitive, and social functioning^3^. Chronic excessive alcohol use leaves heavy drinkers vulnerable to direct and indirect adverse effects, including (but not limited to) cardiovascular disease^4^, nutritional deficiency^5^, cancer^6^, and accelerated aging^7–9^.

Chronic alcohol use affects both brain structure and connectivity^9–11^. Decades of neuroimaging studies have shown that chronic excessive alcohol consumption is associated with widespread patterns of macrostructural and microstructural changes, mostly affecting frontal, diencephalic, hippocampal, and cerebellar structures^9,10,12^. A recent meta-analysis of individuals with AUD (n = 433) showed lower gray matter volume (GMV) in the corticostriatal-limbic circuits, including regions of the prefrontal cortex, insula, superior temporal gyrus, striatum, thalamus, and hippocampus compared to healthy controls (n = 498)^13^. Notably, lower GMV in striatal, frontal, and thalamic regions was associated with AUD duration or lifetime alcohol consumption. Although alcohol consumption can produce global and regional tissue volume deficits, frontal regions are particularly vulnerable to alcohol toxicity^14–16^ and the interactive effects of alcohol and age^9,17^.

Alcohol-related white matter (WM) microstructural alterations are a hallmark injury of AUD^18–20^. Neuroimaging studies have consistently shown WM degeneration of the corpus callosum in AUD^3,21,22^. However, the effects of chronic alcohol use on WM microstructure, as evidenced by decreased fractional anisotropy (FA) and increased mean diffusivity (MD), are not limited to the corpus callosum, but are also seen in the internal and external capsules, fornix, frontal forceps, superior cingulate, and longitudinal fasciculi^3,21,23^. Further, research indicates that anterior and superior WM systems are more vulnerable to heavy drinking than posterior and inferior systems^24^. However, because conventional diffusion tensor imaging (DTI) measures (FA and MD) are based on a simplistic brain tissue microstructure model, they fail to account for the complexities of neurite geometry^25^. For example, the lower FA observed in individuals with AUD may reflect lower neurite density and/or greater orientation dispersion of neurites, which conventional DTI measures do not differentiate^26,27^.

Despite an extensive literature on brain structure and microstructure in individuals with AUD, research exploring associations between alcohol consumption across the drinking spectrum and brain structure and microstructure measures in the general population is limited. In some studies of middle-aged and older adults, moderate alcohol consumption was associated with lower total cerebral volume^28^, gray matter atrophy^29,30^, and lower density of gray matter in frontal and parietal brain regions^30^. However, other studies have failed to show an association^31^, and one study showed a positive association of light-to-moderate alcohol consumption and GMV in older men^32^. One interpretation of these findings is that an inverse U-shaped, dose-dependent association exists between alcohol use and brain structure, with light-to-moderate drinking being protective against and heavy drinking being a risk factor for reductions in GMV^32^. This interpretation was not supported by a longitudinal cohort study, which showed no difference in structural brain measures between abstinent individuals and light drinkers, and moderate-to-heavy drinkers showed GMV atrophy in the hippocampi and impaired WM microstructure (lower FA, higher MD) in the corpus callosum^33^.

The inconclusive nature of the evidence regarding the association between moderate alcohol intake and brain structure in the general population may reflect the literature’s patchwork nature, consisting as it does of mostly small, unrepresentative studies with limited statistical power^34,35^. Moreover, most studies have not accounted for the effects of many relevant covariates and, therefore, have yielded potentially biased findings. Potential confounds that may be associated with individual differences in both alcohol intake and neuroanatomy include sex (women are more vulnerable than men)^36^, body mass index (BMI) (vulnerability increases as a function of BMI)^37^, age (older adults are more vulnerable than younger adults)^38,39^, and genetic population structure (i.e., biological characteristics that are correlated with environmental causes)^40^. Similar to other research fields, progress in this area may also be limited by publication bias^41^.

The current study examines the associations between alcohol intake and measures of GM structure and WM microstructure in the brain in the largest population sample available to date. We address the existing literature’s limitations through a preregistered analysis of multimodal imaging data from the UK Biobank (UKB)^42–44^ that controls for numerous potential confounds. The UKB, a prospective cohort study representative of the United Kingdom (UK) population aged 40-69 years, is the largest available collection of high-quality MRI brain scans, alcohol-related behavioral phenotypes, and measurements of the socioeconomic environment. The UKB brain imaging data include three structural modalities, resting and task-based fMRI, and diffusion imaging^42–44^. The WM fiber integrity measures available in the UKB include the conventional FA and MD metrics and neurite orientation dispersion and density imaging (NODDI) measures^26^. Such measures offer information on WM microstructure and estimates of neurite density (i.e., intracellular volume fraction; ICVF), extracellular water diffusion (i.e., isotropic volume fraction; ISOVF), and tract complexity/fanning (i.e., orientation dispersion, OD). This allows us to assess the nature of the association between alcohol intake and WM microstructure in greater detail than any previous studies on the topic. Specifically, we assess associations between alcohol intake (i.e., mean daily alcohol units; one unit=10 ml or 8 g of ethanol) and imaging derived phenotypes (IDPs) of brain structure (total GMV, total WMV, and 139 regional GMVs), as well as 375 IDPs of WM microstructure (DTI and NODDI indices), using data from 36,678 UKB participants. Our analyses adjust for numerous covariates (see Methods for an exhaustive list of these).

Our sample size provides us statistical power of 90% to detect effects as small as *f*^2^ > 0.00078 at the 5% significance level, after accounting for multiple hypotheses testing (*p*_uncorrected_ <1.64×10^−4^). Given previous findings, we hypothesized to see a reduction in global GMV and WMV in heavy drinkers. However, the large general population sample provided sufficient sensitivity to qualitatively and quantitatively assess how effects vary across the entire drinking spectrum and test at what threshold effects emerge. Our well-powered design also allowed us to explore whether the effects of alcohol intake on GMV and WM microstructure are localized in specific regions or conversely widespread across the brain and compare the effects across various WM integrity indicators.

## RESULTS

Table 1 summarizes the characteristics of the 36,678 participants (52.8% female), including the distributions of age, daily alcohol units and global GMV and WMV. Participants were healthy, middle-aged and older adults. We normalize all of the IDPs for head size by multiplying the raw IDP by the head size scaling factor.

**Table 1.**
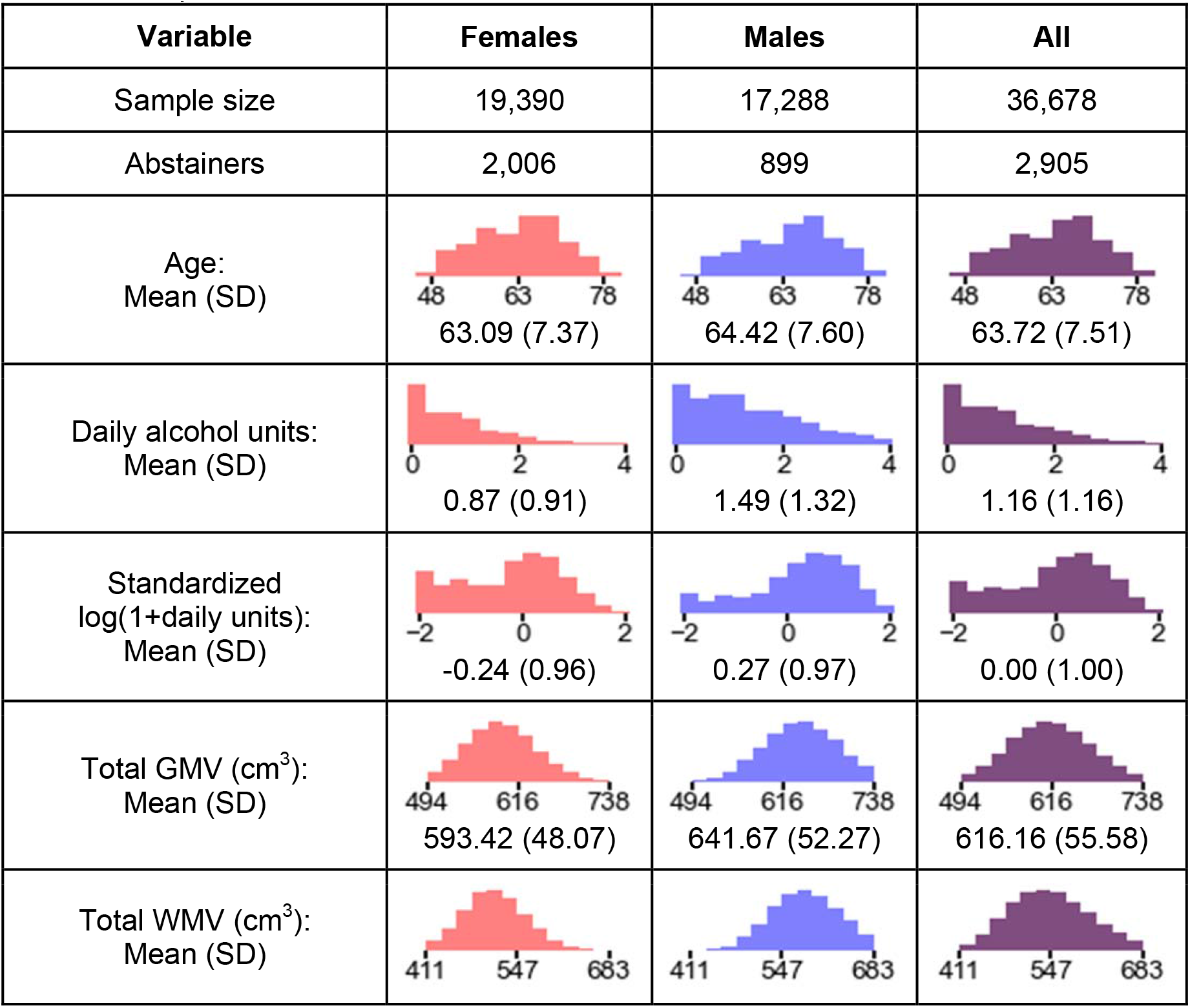
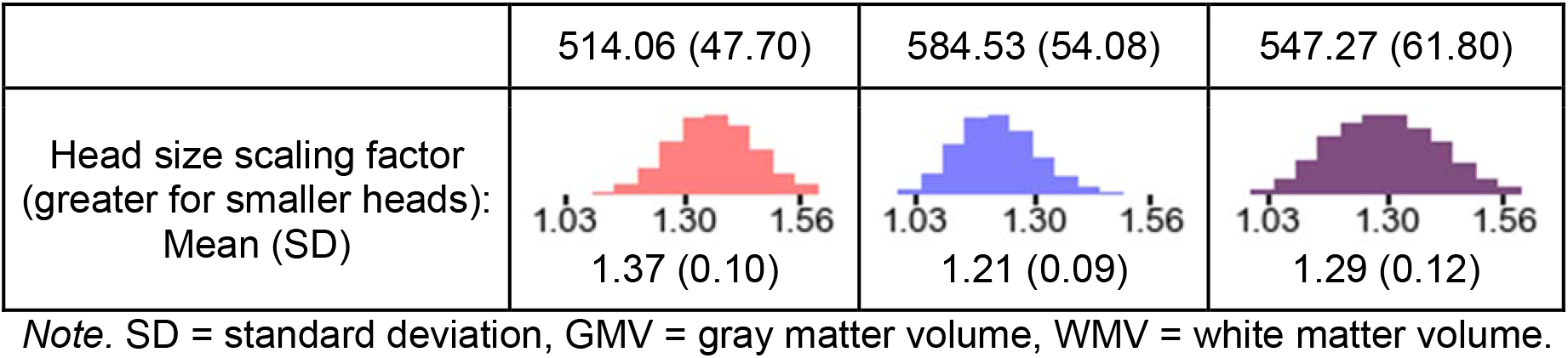
Empirical distributions of variables.

### Relationship between Global GMV and WMV and alcohol intake

The scatter plots in Figure 1 and Figure 2 illustrate the relationships of the global IDPs (GMV and WMV, both normalized for head size) with age and daily alcohol intake (in log scale) within sex. The figures include local polynomial regression lines (LOWESS), which indicate negative trends on every dimension. This preliminary analysis also demonstrates slight non-linearities in both dimensions, with curves appearing concave. Therefore, our subsequent regression models include both linear and quadratic terms for age and logged alcohol intake, test the joint significance of the associations of the two terms with brain structure via an F-test (see Methods), and quantify the effect size via the IDP variance explained by alcohol intake, above the other covariates (Δ*R*^2^).

**Figure 1.**
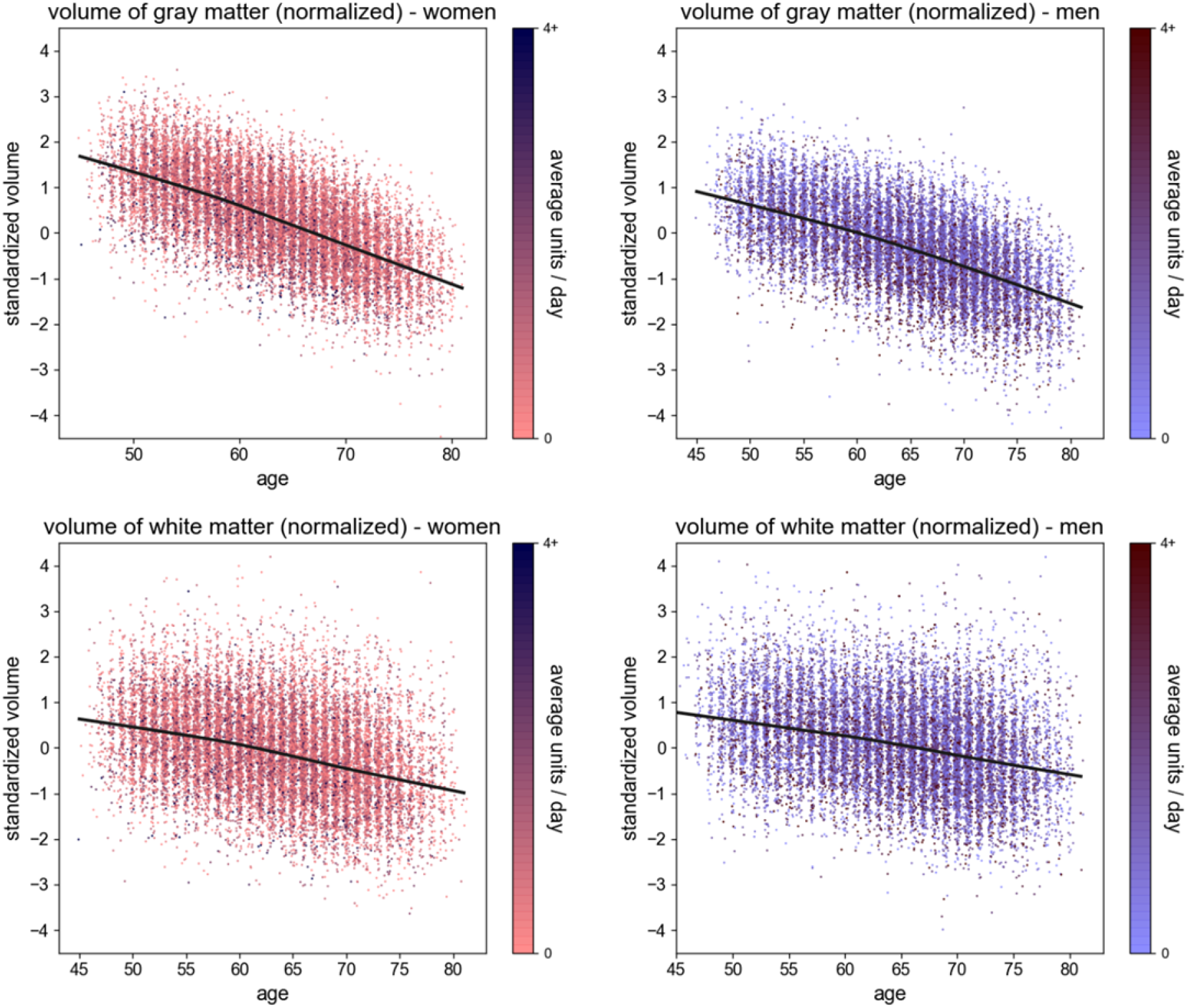
Scatter plots of whole-brain standardized gray matter volume (women, upper left; men, upper right) and standardized white matter volume (women, lower left; men, lower right), all normalized for head size, against the individual’s age (x-axis). The plots also show the LOWESS regression line (smoothness: a=0.2). The 95% confidence interval is indistinguishable from the regression line. The colors are representative of the average daily alcohol consumption.

**Figure 2.**
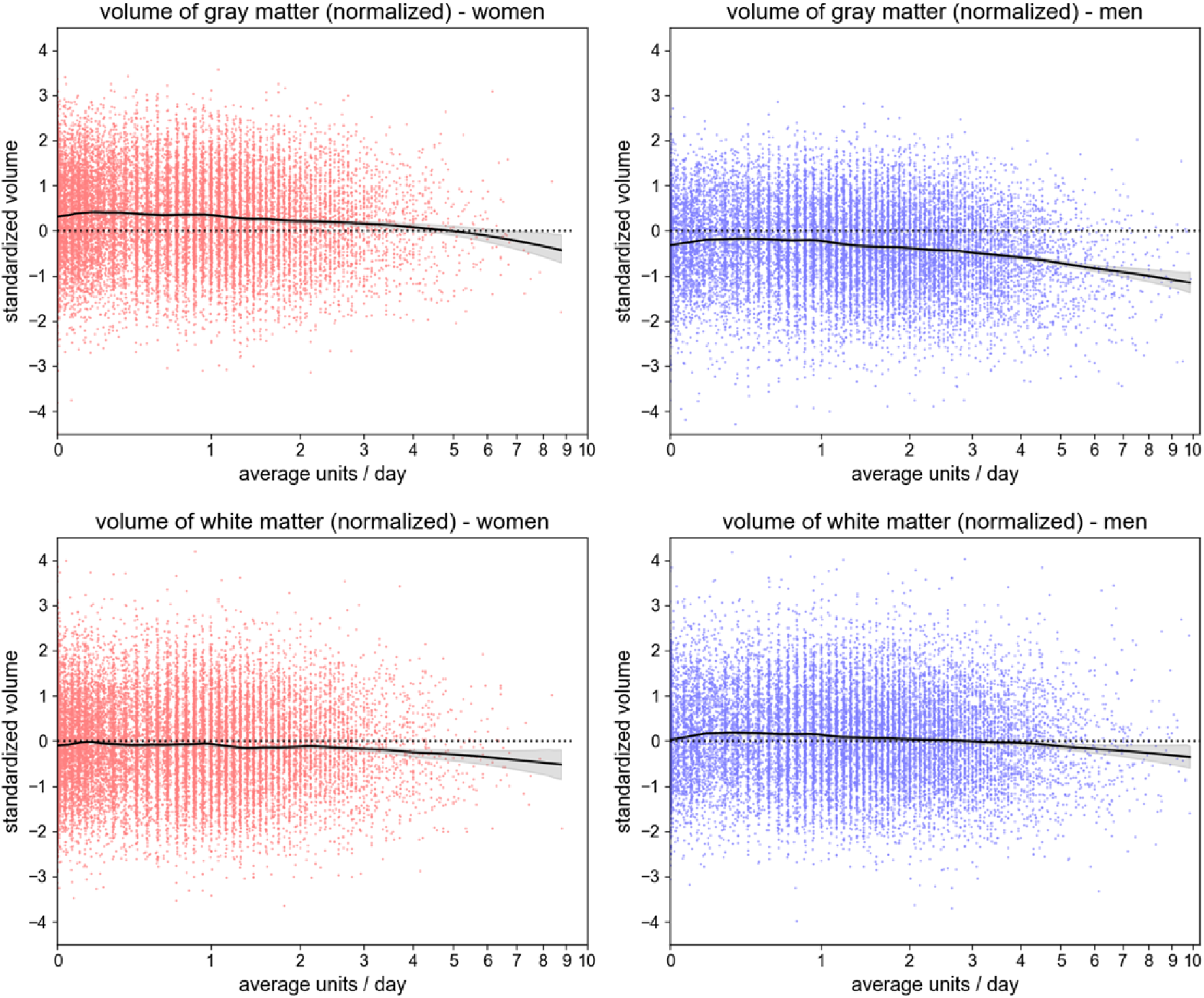
Scatter plots of whole-brain standardized gray matter volume (women, upper left; men, upper right) and standardized white matter volume (women, lower left; men, lower right), all normalized for head size, against the individual’s daily alcohol consumption (x-axis, in log scale). The plots also show the LOWESS regression line (smoothness: a=0.2), with its 95% confidence interval.

We estimate linear regressions to quantify the relationships between daily alcohol intake, as well as its interactions with age and sex, and the global IDPs. Our main analyses (*N* = 36,585) controls for age, height, handedness, sex, smoking status, socioeconomic status, genetic ancestry, and county of residence (see Methods). Table 2 summarizes the results, revealing that both global IDPs decrease as a function of daily alcohol intake. Alcohol intake explains 1% of the variance in global GMV and 0.3% of the variance in global WMV across individuals beyond all other control variables (both *p* < 10^−16^). Additional analyses excluding abstainers (*N* = 33,733) or heavy drinkers (*N* = 34,383), as well as models using an extended set of covariates (addition of BMI, educational attainment, and weight; *N* = 36,678) yield similar findings, though the variance explained by alcohol intake beyond other control variables is reduced to 0.4% for GMV and 0.1% for WMV when heavy drinkers are excluded (Extended Data Tables 1 and 2).

**Table 2.** Regression analysis with global IDPs as outcome variables. All regressions include standard controls. Intake is measured in log(1 + daily units of alcohol).

| Variable | Dependent variable: global GMV |  | Dependent variable: global WMV |  |
| --- | --- | --- | --- | --- |
| | N: 36,678 (d.f.: 36,585), $R^2$: 0.514 | | N: 36,678 (d.f.: 36,585), $R^2$: 0.514 | |
|  | Regression Coefficient (S.Err),<br>95% CI | t-stat<br>(p-value) | Regression Coefficient (S.Err),<br>95% CI | t-stat<br>(p-value) |
| intake | -0.1095 (0.0058),<br>CI: [-0.1209,-0.0982] | -19.0<br>($p < 1.0\text{e-}16$) | -0.0650 (0.0078),<br>CI: [-0.0802,-0.0498] | -8.4<br>($p < 1.0\text{e-}16$) |
| intake<sup>2</sup> | -0.0651 (0.0037),<br>CI: [-0.0723,-0.0579] | -17.7<br>($p < 1.0\text{e-}16$) | -0.0370 (0.0050),<br>CI: [-0.0468,-0.0273] | -7.5<br>($p = 7.8\text{e-}14$) |
| intake x male | 0.0174 (0.0080),<br>CI: [0.0018,0.0330] | 2.2<br>($p = 2.9\text{e-}02$) | 0.0164 (0.0107),<br>CI: [-0.0046,0.0374] | 1.5<br>($p = 1.2\text{e-}01$) |
| intake x std. age | 0.0080 (0.0037),<br>CI: [0.0008,0.0152] | 2.2<br>($p = 3.0\text{e-}02$) | 0.0111 (0.0050),<br>CI: [0.0014,0.0208] | 2.2<br>($p = 2.5\text{e-}02$) |
| std. age | -0.5991 (0.0038),<br>CI: [-0.6066,-0.5916] | -157.0<br>($p < 1.0\text{e-}16$) | -0.3213 (0.0051),<br>CI: [-0.3313,-0.3112] | -62.6<br>($p < 1.0\text{e-}16$) |
| std. age<sup>2</sup> | -0.0378 (0.0034),<br>CI: [-0.0445,-0.0311] | -11.0<br>($p < 1.0\text{e-}16$) | -0.0127 (0.0046),<br>CI: [-0.0217,-0.0037] | -2.8<br>($p = 5.7\text{e-}03$) |
| Against model without intake and interactions<br>Delta $R^2$: 0.0099, F-test: $p < 1.0\text{e-}16$ | | | Against model without intake and interactions<br>Delta $R^2$: 0.0033, F-test: $p < 1.0\text{e-}16$ | |

In the eight regressions we tested, the interaction between alcohol intake and sex is not significant at the 1% level, except weakly for GMV when including the extended control variables (BMI, weight, and educational attainment). Given our large sample size, this indicates that if there is any effect, it is negligible. Similarly, the interaction between intake and age is weakly significant for the regressions excluding abstainers only, indicating that it is also negligible if any effect exists. None of the interaction terms are significant at the 0.1% level. Consequently, we excluded the interaction terms from the analyses of local IDPs.

To evaluate the magnitude of the main effects of alcohol intake on the global IDPs, we use our regression models to calculate the predicted change in global GMV and WMV associated with an increase of daily alcohol intake by one unit (Table 3A). This prediction is similar when using different sets of control variables and when excluding abstainers or heavy drinkers. Given the non-linear relationship between global IDPs and alcohol intake, the effect varies across the drinking range. There is virtually no change (less than .03 standard deviations) in the predicted global GMV and WMV when shifting from abstinence to one daily alcohol unit. However, the effect of intake increases as the number of daily units increases. An increase from one to two daily units is associated with a decrease of 0.127 and 0.074 standard deviations in predicted global GMV and WMV, respectively. A change from two to three daily units is associated with a 75% greater decrease of 0.223 and 1.28 standard deviations in GMV and WMV, respectively. Table 3B benchmarks the predicted effect magnitudes against the effects associated with aging for an average 50-year-old UKB participant, based on our regression models. For illustration, the effect associated with a change from one to two daily alcohol units is equivalent to the effect of aging 2 years, where the increase from two to three daily units is equivalent to aging 3.5 years.

**Table 3A.**
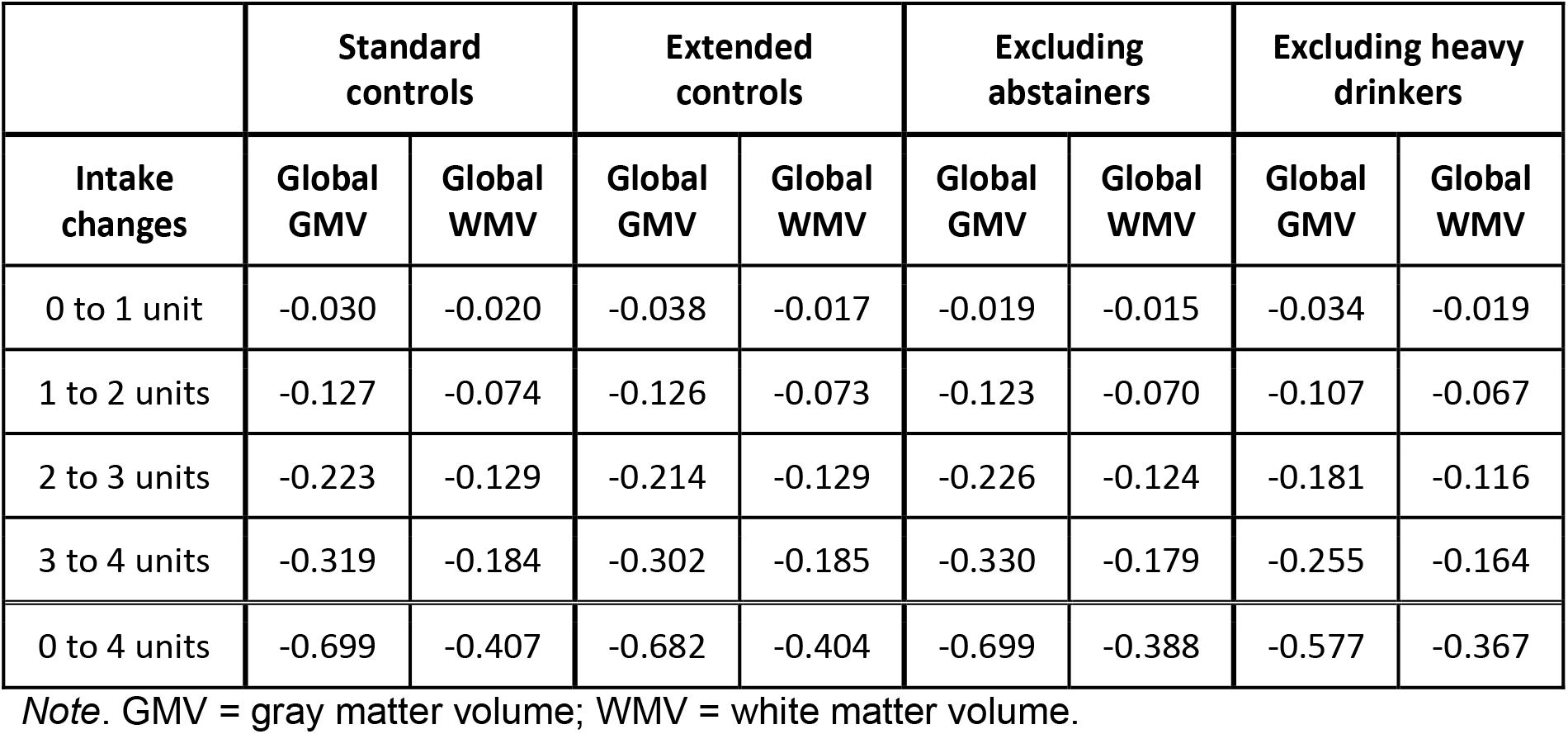
Predicted average additional effect (in standard deviations of IDP) of increasing alcohol intake by one daily unit on whole-brain gray matter volume and white matter volume, for models with different sets of controls (first and second columns), and for standard controls with samples excluding abstainers (third column) and heavy drinkers (last column).

| Intake changes | Standard controls |  | Extended controls |  | Excluding abstainers |  | Excluding heavy drinkers |  |
| --- | --- | --- | --- | --- | --- | --- | --- | --- |
|  | Global GMV | Global WMV | Global GMV | Global WMV | Global GMV | Global WMV | Global GMV | Global WMV |
| 0 to 1 unit | -0.030 | -0.020 | -0.038 | -0.017 | -0.019 | -0.015 | -0.034 | -0.019 |
| 1 to 2 units | -0.127 | -0.074 | -0.126 | -0.073 | -0.123 | -0.070 | -0.107 | -0.067 |
| 2 to 3 units | -0.223 | -0.129 | -0.214 | -0.129 | -0.226 | -0.124 | -0.181 | -0.116 |
| 3 to 4 units | -0.319 | -0.184 | -0.302 | -0.185 | -0.330 | -0.179 | -0.255 | -0.164 |
| 0 to 4 units | -0.699 | -0.407 | -0.682 | -0.404 | -0.699 | -0.388 | -0.577 | -0.367 |

Note. GMV = gray matter volume; WMV = white matter volume.

**Table 3B.** Equivalent effect of ageing in terms of additional years for an average 50-year old individual.

|  | Standard controls |  |  |  |
| --- | --- | --- | --- | --- |
| Intake changes | Global GMV | Equivalent aging at 50 | Global WMV | Equivalent aging at 50 |
| 0 to 1 unit | -0.030 | 0.5 years | -0.020 | 0.5 years |
| 1 to 2 units | -0.127 | 2.0 years | -0.074 | 2.0 years |
| 2 to 3 units | -0.223 | 3.5 years | -0.129 | 3.5 years |
| 3 to 4 units | -0.319 | 4.9 years | -0.184 | 4.9 years |
| 0 to 4 units | -0.699 | 10.2 years | -0.407 | 10.4 years |

*Note.* GMV = gray matter volume.

Figure 3 displays the averages of the two global IDPs in sub-samples binned according to their daily alcohol consumption range, illustrating the non-linear nature of the relationship between daily units and the global measures. The figure includes statistical tests that compare the average of the IDPs in the different sub-samples to their average in participants who consume one daily unit or less. These tests identify statistically significant effects for all bins of participants consuming more than one daily units, including those consuming as little as 1-2 daily units. These effects are observed both in the full sample and within sex.

**Figure 3.**
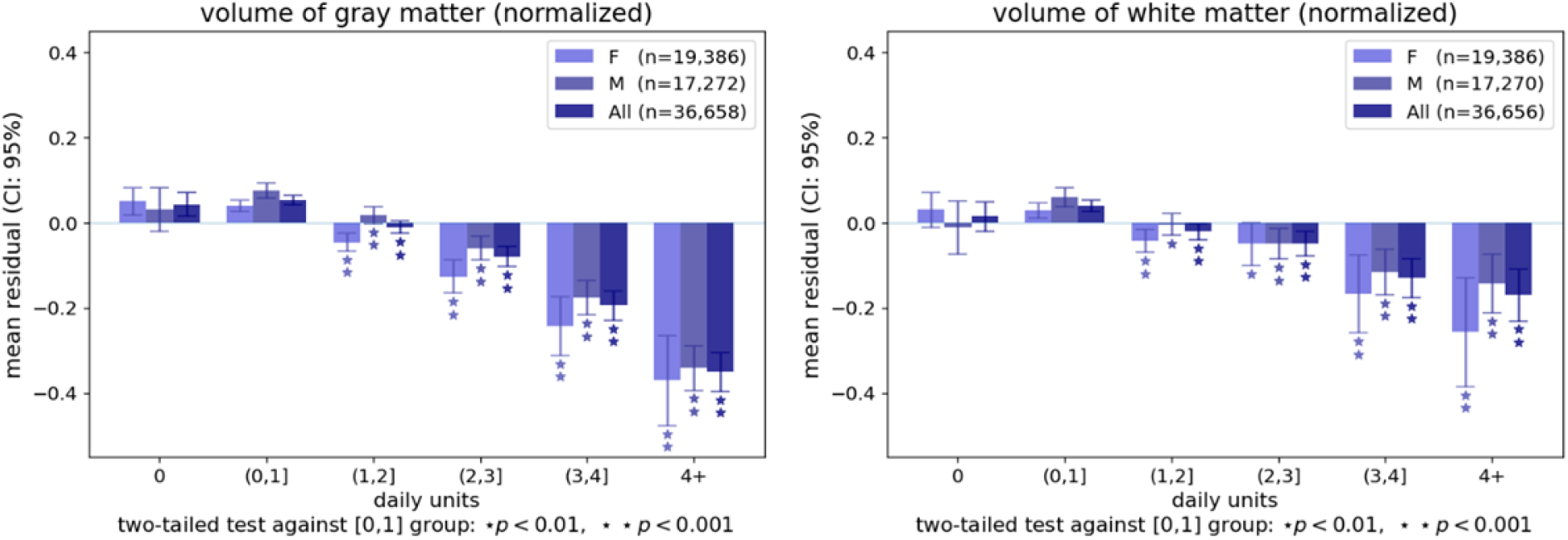
Bar plots representing the average volume of whole-brain gray and white matter volume for individuals grouped by the number of daily alcohol units after controlling for standard control variables (keeping the regression residual). The mean residuals are in terms of standard deviations of the dependent variable. The error bars represent the 95% confidence interval. \**p*<0.01 and \*\**p*<.0001 for groups showing a significant difference against the group consuming up to one alcohol unit daily.

### Relationship between regional GMV and alcohol intake

To investigate whether the reduction in global GMV associated with alcohol intake stems effects of drinking in specific regions, we estimate regression models to quantify the association of alcohol intake with a total of 139 regional GMV IDPs. These IDPs were derived using parcellations from the Harvard-Oxford cortical and subcortical atlases and Diedrichsen cerebellar atlas. Of the 139 GMV IDPs, 125 (88.9%) are significantly associated with log alcohol intake (see Extended Data Table 3). We observe the strongest effects in frontal, parietal, and insular cortices, temporal and cingulate regions, putamen, amygdala and the brain stem. In these regions, alcohol intake explains between 0.3%-0.4% of the variance in local GMV above the other covariates. Extended Data Figure 1 illustrates the marginal effect of increasing daily alcohol units on regional GMV IDPs, grouped by lobe. All of the associations are negative, except the association involving the right pallidum – where the effect size is positive but very small (Δ*R*^2^ = 0.0005). Importantly, the largest regional effect was less than half the size of the association between drinking and global GMV, indicating that the global reduction in GMV associated with alcohol intake is the result of aggregating smaller effects that are widespread across the brain (rather than constrained to specific areas).

In a similar fashion to the analysis using the global IDPs, we calculate the average localized GMV IDP for each daily alcohol unit bin (Extended Data Figure 2) and test their difference against the average of the group drinking up to one unit per day, within sexes and in the overall sample. As expected, the number of regional GMV IDPs showing a significant negative association with alcohol intake, as well as these associations’ magnitudes, increases as the average number of daily alcohol units increases. There are few regions where lower GMV is either not observed as a function of drinking or only apparent among heavy drinkers (e.g., fusiform cortex). However, in most regions, GMV reduction is already visible in the groups that drink moderately (i.e., consuming 1-2 or 2-3 daily units). Thus, the influence of moderate alcohol intake on GMV also appears to be widespread across the brain, and it is detectable in both males and females.

### Relationship between regional WM microstructure and alcohol intake

To evaluate how drinking influences the different indicators of WM integrity at the regional level, we estimate linear regressions to quantify the association of alcohol intake with 375 IDPs, including FA, MD, ICVF, ISOVF, and OD measures extracted via averaging parameters across 74 WM tract regions^45^. Of the 375 WM microstructure IDPs, 179 (47.7%) are significantly associated with alcohol intake (Extended Data Table 4). Generally, alcohol intake is related to lower coherence of water diffusion, lower neurite density, and higher magnitude of water diffusion, indicating less healthy WM microstructure with increasing alcohol intake.

To visualize the magnitude of WM microstructure IDP associations with alcohol intake, Figure 4 displays the statistically significant and non-significant effects, alongside the average change in normalized WM microstructure IDPs associated with a mean daily alcohol intake increasing from 2 to 3 units. Thirteen WM tract regions show consistent significant associations with lower FA and higher ISOVF and MD. The strongest effects of these are in the fornix, where WM integrity was previously found to be affected by drinking in studies of populations with AUD^3,21,23^. In the fornix, alcohol intake accounts for 0.45% of the variance in ISOVF, 0.35% of the variance in MD, and 0.32% of the variance in FA. Other WM tract regions showing a similar pattern yet with effects of weaker magnitude include commissural fibers (genu and body of the corpus callosum, bilateral tapetum), projection fibers (bilateral anterior corona radiata), associative fibers (fornix cres+stria terminalis, left inferior longitudinal fasciculus), and the bilateral anterior thalamic radiations.

**Figure 4.**
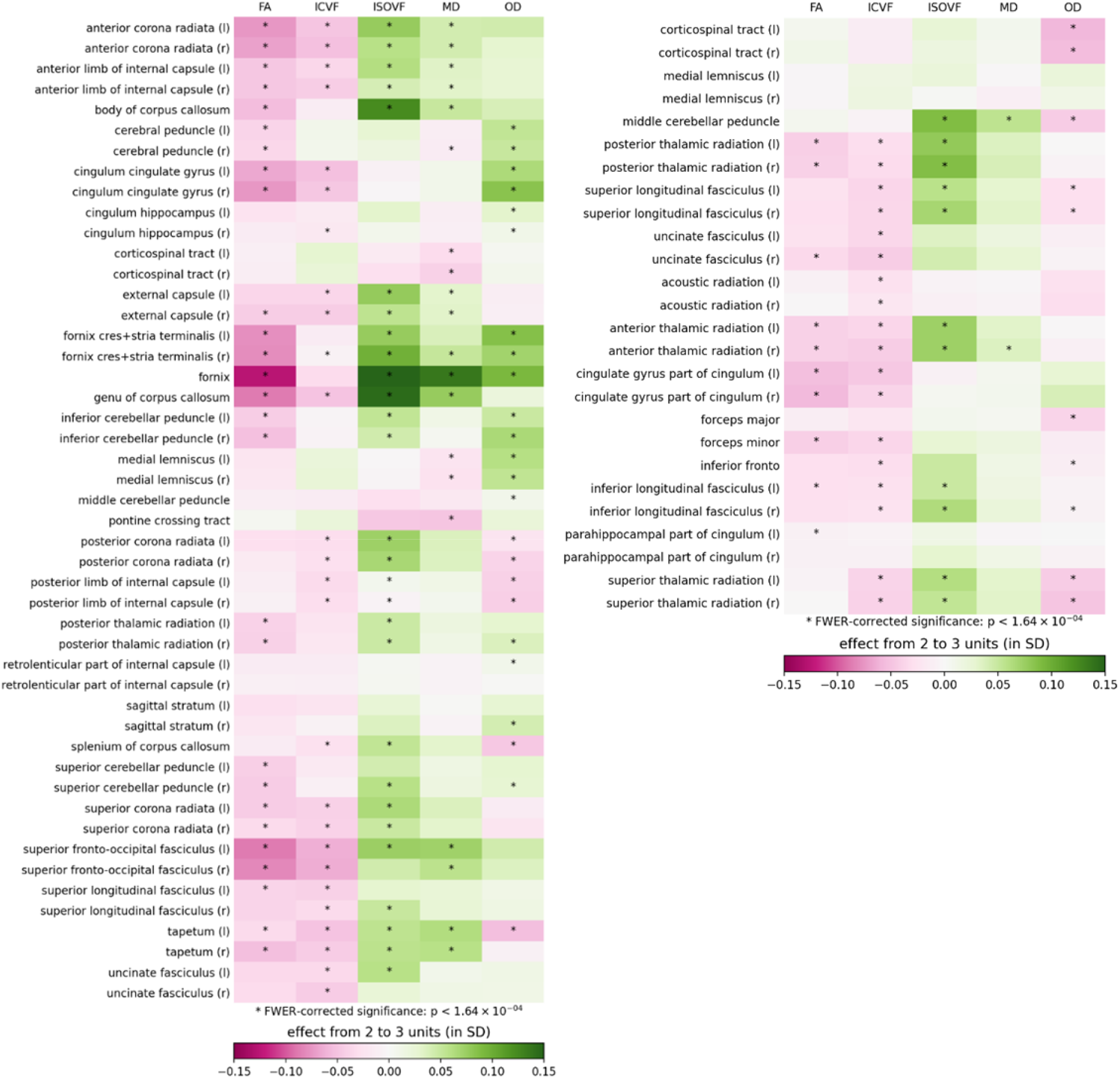
Matrix representing the effect of consuming 2-3 daily alcohol units on water matter microstructure indices of interest across white matter tract regions. r = right, l = left. *\*p* < 1.64×10 ^−4^

Among the NODDI measures, ISOVF showed the strongest effects of alcohol intake all over the brain, most notably in the tract regions discussed above. The associations between drinking and ICVF are also consistently negative yet smaller in size, with daily alcohol intake explaining no more than 0.1% of the variance beyond other control variables in all ICVF IDPs. The associations with OD, which is a measure of tract complexity, are either positive, negative or absent, and while some are statistically significant, they are all very small in size (Δ*R*^2^ < 0.001 for all IDPs).

## DISCUSSION

We report a multimodal brain imaging study of 36,678 middle-aged and older adults of European descent, a population sample whose reported alcohol consumption spanned the spectrum from abstinence to heavy drinking. The scale and granularity of the data provide ample statistical power to identify small effects while accounting for important potential confounds. We observe negative relationships between alcohol intake and global measures of gray and white matter, regional GMVs, and WM microstructure indices. The effects we identify are widespread across the brain, and their magnitude increases with the average absolute number of daily alcohol units consumed.

Notably, the negative associations we observe with global IDPs are already detectable in those who consume between 1 and 2 alcohol units daily. This finding has important implications for recommendations regarding safe drinking levels, both in males and females. In 2016, the UK Chief Medical Officers published new “low-risk” alcohol consumption guidelines that advise limiting alcohol intake to 14 units per week^32^. One alcohol unit is equivalent to 10 ml or 8 g of ethanol, the amount contained in 25 ml of 40% spirits, 250 ml of 4% beer, or 76 ml of 13% wine. Many drinking establishments serve drinks that contain 35-50 ml of 40% spirits (1.4-2 units), 568 ml of 4% beer (2.27 units), and 175 ml of 13% wine (2.30 units)^33^. Thus, in the UK, consuming just one alcoholic drink (or two units of alcohol) daily can have negative effects on brain health. This has important public health implications insofar as 57% of UK adults, or an estimated 29.2 million individuals^28^ endorse drinking during the past week.

The negative associations between alcohol intake and total GMV and WMV are consistent with prior studies of early middle-aged^46^ and older adults^28,47^. Because men consume more alcohol units per day and had larger global GMV and WMV, we further examine the effect of sex in detail. We find negative associations between alcohol intake and the global IDPs for both sexes and weak evidence for interactive effects between alcohol intake and sex on the brain. These findings are similar to a recent study of early middle-aged adult moderate drinkers that showed smaller brain volumes associated with moderate alcohol consumption in both men and women^46^. The weak sex-by-alcohol interactions also comport with the findings of an earlier longitudinal study in individuals with AUD^38^; however, other cross-sectional studies have reported greater volume deficits in women than men^48,49^.

Although nearly 90% of all regional GMVs show significant negative associations with alcohol intake, the most extensively affected regions included the frontal, parietal, and insular cortices, with deficits also in temporal and cingulate regions. Associations are also marked in the brain stem, putamen, and amygdala. The share of variance explained by alcohol intake for these regions is smaller in size than for global GMV, suggesting that the latter is the result of aggregation of many small effects that are widespread, rather than a localized effect that is limited to specific regions. Alcohol intake is further associated with poorer WM microstructure (lower FA and higher ISOVF and MD) in specific classes of WM tract regions. The commissural fibers (genu and body of the corpus callosum, bilateral tapetum), projection fibers (bilateral anterior corona radiata), associative bundles (fornix, fornix cres+stria terminalis, left inferior longitudinal fasciculus), and the bilateral anterior thalamic radiations show the most consistent associations with alcohol intake, with the fornix showing the strongest effects. The fornix is the primary outgoing pathway from the hippocampus^50^, and WM microstructural alterations in the fornix are consistently associated with heavy alcohol use and memory impairments^3,51^. Moreover, recent research indicates that one extreme-drinking episode can cause acute WM damage to the fornix, suggesting that the fornix may be particularly vulnerable to alcohol’s effects.

Our findings are partly consistent with studies of individuals with AUD^18,52^. The pattern of microstructural alterations in our general population sample show that widespread WM alterations are present across multiple WM systems. Like individuals with AUD, alcohol intake in this healthy population sample is associated with microstructural changes in superficial WM systems functionally related to GM networks, including the frontoparietal control and attention networks, and the default mode, sensorimotor, and cerebellar networks. Deeper WM systems (superior longitudinal fasciculus and dorsal frontoparietal systems, inferior longitudinal fasciculus system, and deep frontal WM) thought to be involved in cognitive functioning by regulating reciprocal connectivity^52,53^ are also associated with alcohol intake. Within these WM systems, alcohol intake is most strongly associated with ISOVF, MD, and FA WM microstructure indices; whereas, associations with ICVF are small, and OD associations are inconsistent or nonexistent. Alcohol intake shows positive associations with ISOVF and MD and negative associations with FA. This pattern of alcohol-associated WM microstructural disruption supports previous research showing excessive intracellular and extracellular fluid in individuals with AUD^20^. Given that alcohol increases blood-brain permeability^54^ and activates pro-inflammatory cytokines in the brain^55^, the association between alcohol intake and higher ISOVF (extracellular water diffusion) may be due to inflammatory demyelination. For example, higher ISOVF is evident in WM lesions of multiple sclerosis, characterized histopathologically by inflammatory demyelination associated with blood-brain permeability and axonal injury^56,57^. Additional research is warranted; however, these findings suggest that even low-moderate alcohol intake increases intracellular and extracellular water diffusion in WM, which may be a result of alcohol-induced inflammatory demyelination.

Our study is not without limitations, which provide opportunities for further research. First, we rely on a sample of middle-aged individuals of European ancestry living in the UK. We hope that future work will test the generalizability of our findings to individuals from other populations and in other age groups. It is reasonable to expect that the relationship we observe would differ in younger individuals who have not experienced the chronic effects of alcohol on the brain. An additional limitation stems from the self-reported alcohol intake measures in the UK Biobank, which cover only the past year. Such estimates do not adequately reflect drinking prior to the past year and are susceptible to reporting and recall bias^38,39^. Further, our analyses do not account for individuals with a past diagnosis of AUD. Earlier studies have shown that the brain shows some recovery with prolonged sobriety, but this recovery varies with age and sex, and recovery might be incomplete^58–60^. Thus, a past diagnosis of AUD would likely influence our results. We hope that future studies will shed light on how a history of AUD with prolonged recovery influences brain structure in middle-aged and older adults.

In summary, this large-scale brain imaging sample provides additional evidence of alcohol’s adverse effects on brain macrostructure and microstructure in a general population sample of middle-aged and older adults. Alcohol intake is negatively associated with global brain volume measures, regional GMVs, and WM microstructure. The associations between alcohol intake and regional GMV are evident across the entire brain, with the largest deficits observed in frontal, parietal, and insular cortices, temporal and cingulate regions, the brain stem, putamen, and amygdala. Alcohol intake is related to WM microstructural alterations in several WM tract regions connecting large-scale networks and deeper WM systems. Most of these adverse effects are already apparent with an average consumption of only one to two daily alcohol units. Thus, this multimodal imaging study highlights the risk that even moderate drinking poses on the brain in middle-aged and older adults.

## Methods

### Sample, procedure and exclusion criteria

Our sample comprised 36,678 individuals of European ancestry from the UKB, all study participants whose data were available as of September 1, 2020. All UK Biobank (www.ukbiobank.ac.uk) participants provided written informed consent, and ethical approval was granted by the North West Multi-Centre Ethics committee. Participants provided demographic and health information via touchscreen questionnaires. A nurse conducted a medical history interview, which included self-report of medical diagnoses and other conditions or life events that were used to evaluate eligibility to participate (study details are available at http://www.ukbiobank.ac.uk/key-documents/). Vital signs were obtained, and body mass index was calculated as weight (kg)/height^2^ (m).

The data provided by the UK Biobank and was already subject to quality control^61^. We excluded individuals with IDP values outside a range of four standard deviations (SDs). We chose this lenient threshold as a non-trivial number of observations (97 for GM, 127 for WM) fall between three and four SDs away from the mean, given the large sample size. The IDPs beyond the four SD range are likely the results of processing errors, or the corresponding individuals present severe brain irregularities (5 individuals for GM, 7 for WM). Note that the exclusion of these outliers does not change the statistical significance nor the magnitude of the effects that we report. The exclusion of individuals falling within three SDs of the mean does not change the results either.

### Measures of alcohol consumption

Participants self-reported the number of alcohol units (10 ml of pure ethanol) consumed, in “units per week” (for frequent drinkers) or “units per month” (for less frequent drinkers), across several beverage categories (red wine, white wine/champagne, beer/cider, spirits, fortified wine, and “other”). The UKB assessment defined units of alcohol as follows: a pint or can of beer/lager/cider = two units; a 25-ml single shot of spirits = one unit; and a standard glass of wine (175 ml) = two units. We computed the number of weekly units by summing the weekly units consumed in all categories. When reported monthly, the intake was converted to units per week by dividing by 4.3. The number of weekly units was divided by 7 to determine units per day.

### MRI data acquisition and processing

MRI data were acquired using a Siemens Skyra 3T scanner (Siemens Healthcare, Erlangen, Germany) using a standard 32-channel head coil, according to a freely available protocol (http://www.fmrib.ox.ac.uk/ukbiobank/protocol/V4_23092014.pdf), documentation (http://biobank.ctsu.ox.ac.uk/crystal/docs/brain_mri.pdf), and publication^40^. As part of the scanning protocol, high-resolution T1-weighted images, three-dimensional T2-weighted fluid-attenuated inversion recovery (FLAIR) images, and diffusion data were obtained. High-resolution T1-weighted images were obtained using an MPRAGE sequence with the following parameters: TR=2000ms; TE=2.01ms; 208 sagittal slices; flip angle, 8°; FOV=256 mm; matrix=256×256; slice thickness=1.0mm (voxel size 1×1×1mm); total scan time=4min 54s. 3D FLAIR images were obtained with the following parameters: TR=1800ms; TE=395.0ms; 192 sagittal slices; FOV=256mm; 256×256; slice thickness=1.05mm (voxel size 1.05×1×1mm); total scan time=5min 52s. Diffusion acquisition comprised a spin-echo echo-planar sequence with 10 T2-weighted (b ≈ 0 s mm^−2^) baseline volumes, 50 b = 1000 s mm^−2^ and 50 b = 2000 s mm^−2^ diffusion-weighted volumes, with 100 distinct diffusion-encoding directions and 2 mm isotropic voxels; total scan time=6min 32s.

Structural imaging and diffusion data were processed by the UK Biobank team and made available to approved researchers as imaging-derived phenotypes (IDPs); the full details of the image processing and QC pipeline are available in an open-access article^42,62^. IDPs used in analyses included whole-brain GMV, whole-brain WMV, 139 regional GMV IDPs derived using parcellations from the Harvard-Oxford cortical and subcortical atlases and Diedrichsen cerebellar atlas (UKB fields 25782 to 25920), and 375 tract-averaged measures of fractional anisotropy (FA), mean diffusivity (MD), intra-cellular volume fraction (ICVF), isotropic volume fraction (ISOVF), and orientation diffusion (OD) extracted by averaging parameters over 74 different white-matter tract regions based on subject-specific tractography^63^ and from population-average WM masks^45^. Volumetric IDPs were normalized for head size by multiplying the raw IDP by the T1-based “head size scaling factor”^62^.

### Statistical Analyses

#### Descriptive analysis using global IDPs

We plot global GMV and WMV in males and females separately, normalized for head size, against age (Figure 1) and alcohol intake (i.e., alcohol units/day on a log scale) (Figure 2).

#### Global IDPs, regional GMV, and WM microstructure analyses

Our main analysis estimates a linear regression of several IDPs on alcohol intake in *log(1+daily units)*, including various control variables and interactions. Given the slight concavity of the LOWESS regression lines in the descriptive analysis of the global IDPs, we included both linear and quadratic values for alcohol intake and age in the regression:

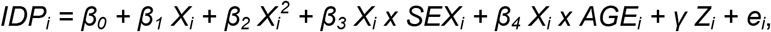

where *IDP_i_* is the IDP normalized for head size, *X_i_* is the standardized alcohol intake in log(1 + daily units), *AGE_i_* is standardized age, *Z_i_* is a vector of control variables, and *e_i_* is an error term.

Our analyses comprise of models that include two different sets of control variables. The standard set includes standardized age, standardized age^2^, standardized height, handedness (right/left/ambidextrous; dummy-coded), sex (female:0, male:1), current smoker status, former light smoker, former heavy smoker, and standardized Townsend index of social deprivation measured at the zip code level^64^. To control for genetic population structure, the models also include the first 40 genetic principal components^65^ and county of residence (dummy-coded)^66^. A second set of extended control variables includes all of the standard controls along with standardized body mass index (BMI), standardized educational attainment^67^, and standardized weight. To determine whether observations at the extreme ends of the drinking distribution bias the estimates of the relationship between alcohol intake and IDPs, we also estimate a model that excludes abstainers and a model that excludes heavy drinkers (i.e., women who reported consuming more than 18 units/week and men who consumed more than 24 units/week), both with standard controls. For each of the IDPs, we test the hypothesis that alcohol had no effect on the outcome measure via an F-test that compares our model against a model with only the control variables (excluding alcohol intake and related interaction terms).

We separate the analysis into two parts: (1) global analysis and (2) regional GMV and WM microstructure analysis, including 514 IDPs in total (139 GMV IDPs, 375 WM microstructure IDPs). The interactions of alcohol intake variables with sex and age are not significant in the global analysis (p > 0.001), so we exclude these them from the regional analyses. To control the family-wise error rate in the regional GMV and WM microstructure analysis, we determine the significance thresholds for all regressions using the Holm method^68^, ensuring a family-wise error rate below 5%. When testing for M hypotheses, this method orders the corresponding p-values from lowest to highest: *p_0_, …, p_M_*, and identifies the minimal index *k* such that *p_k_ > 0.05/(M+1-k)*. All hypotheses with an index *m < k* are then considered to be statistically significant. In our application, the significance threshold was determined to be 1.64 x 10^−4^.

To quantify and visualize associations between alcohol intake and IDPs (i.e., global GMV and WMV, and regional GMV IDPs), we bin participants in the following six categories based on average alcohol intake: (1) abstainers, (2) individuals who drank less than one unit/day, (3) individuals who drank between one (included) and two (excluded) units/day (recommended maximal alcohol consumption based on the UK Chief Medical Officers “low-risk” guidelines^32^), (4) individuals who drank between two (included) and three (excluded) units/day, (5) individuals who drank between three (included) and four (excluded) units/day, and (6) individuals who drank at least four units/day. After regressing the influence of the standard control variables, we then calculate the mean residual values (measured in standard deviations of IDPs) and 95% confidence intervals (CI). By first regressing the dependent variables on the standard control variables, the estimated effect can be interpreted as the part of the change in IDP that is not explained by these other variables, and it is represented in terms of standard deviations from the average. All results are available to the readers in extended data figures and tables. Specifically, Extended Data Tables 3 and 4 include the regression coefficients, *p*-values and incremental variance explained above that of control variables for all of the regional IDPs (both significant and non-significant). Extended Data Figure 2 includes the average GMV of all regions tested (both significant and non-significant), in bins of participants with different daily alcohol intake levels.

#### Pre registration

We registered the analysis plan was preregistered with the Open Science Foundation (https://osf.io/trauf/?view_only=a3795f76c5a54830b2ca443e3e07c0f0).

#### Data Availability

Data and materials are available via UK Biobank at http://www.ukbiobank.ac.uk/.

#### Code Availability

The analysis code used in this study is publicly available with the Open Science Framework (https://osf.io/trauf/?view_only=a3795f76c5a54830b2ca443e3e07c0f0).

## Acknowledgments

This research was carried out under the auspices of the Brain Imaging and Genetics in Behavioral Research Consortium (https://big-bear-research.org/), using UK Biobank resources under application 40830. The study was supported by funding from an ERC Consolidator Grant to PK (647648 EdGe), NSF Early Career Development Program grant (1942917) to GN, the National Institute on Alcohol Abuse and Alcoholism to RRW (K23 AA023894), and the VISN 4 Mental Illness Research, Education and Clinical Center at the Crescenz VA Medical Center. GN thanks Carlos and Rosa de la Cruz for ongoing support.

## Author contributions

RD, PK, HRK, GN, and RRW conceived and designed the study. RD analyzed data. RD, GA, KJ, PK, HRK, GN, and RRW interpreted data. RD, GN and RRW wrote the paper. GA, NS, PK, and HRK, critically edited the work. RD, GN and RRW finalized all edits. All authors approved the final version to be submitted for publication and agree to be accountable for all aspects of this work.

