## Extended Data Table 1 for "Multimodal brain imaging study of 36,678 participants reveals adverse effects of moderate drinking"

**Extended Data Table 1. Regression results: Volume of gray matter (whole brain, normalized).**

log intake is measured in standardized log( 1 + daily units of alcohol ).

| | **Standard Controls** | | **Extended Controls** | |
|---|---|---|---|---|
| | N: 36,678 (d.f.: 36,585), $R^2$: 0.514 | | N: 36,678 (d.f.: 36,578), $R^2$: 0.520 | |
| **Variable** | **Regression Coefficient (S.Err), 95% CI** | **t-stat (p-value)** | **Regression Coefficient (S.Err), 95% CI** | **t-stat (p-value)** |
| log intake | -0.1095 (0.0058), CI: [-0.1209,-0.0982] | -19.0 (p < 1.0e-16) | -0.1125 (0.0057), CI: [-0.1238,-0.1013] | -19.6 (p < 1.0e-16) |
| log intake$^2$ | -0.0651 (0.0037), CI: [-0.0723,-0.0579] | -17.7 (p < 1.0e-16) | -0.0596 (0.0037), CI: [-0.0668,-0.0524] | -16.2 (p < 1.0e-16) |
| log intake x male | 0.0174 (0.0080), CI: [0.0018,0.0330] | 2.2 (p = 2.9e-02) | 0.0224 (0.0079), CI: [0.0068,0.0379] | 2.8 (p = 4.8e-03) |
| log intake x std. age | 0.0080 (0.0037), CI: [0.0008,0.0152] | 2.2 (p = 3.0e-02) | 0.0080 (0.0037), CI: [0.0008,0.0151] | 2.2 (p = 3.0e-02) |
| standardized age | -0.5991 (0.0038), CI: [-0.6066,-0.5916] | -157.0 (p < 1.0e-16) | -0.5995 (0.0038), CI: [-0.6069,-0.5921] | -158.0 (p < 1.0e-16) |
| standardized age$^2$ | -0.0378 (0.0034), CI: [-0.0445,-0.0311] | -11.0 (p < 1.0e-16) | -0.0403 (0.0034), CI: [-0.0469,-0.0336] | -11.8 (p < 1.0e-16) |
| | **Against model without log intake and interactions** Delta $R^2$: 0.0099 F-test: p < 1.0e-16 | | **Against model without log intake and interactions** Delta $R^2$: 0.0099 F-test: p < 1.0e-16 | |

| | **Excluding Abstainers** | | **Excluding Heavy Drinkers** | |
|---|---|---|---|---|
| | N: 33,773 (d.f.: 33,676), R2: 0.517 | | N: 34,383 (d.f.: 34,286), R2: 0.510 | |
| **Variable** | **Regression Coefficient (S.Err), 95% CI** | **t-stat (p-value)** | **Regression Coefficient (S.Err), 95% CI** | **t-stat (p-value)** |
| log intake | -0.1025 (0.0064), CI: [-0.1151,-0.0899] | -15.9 (p < 1.0e-16) | -0.0960 (0.0073), CI: [-0.1104,-0.0817] | -13.1 (p < 1.0e-16) |
| log intake$^2$ | -0.0701 (0.0045), CI: [-0.0790,-0.0611] | -15.4 (p < 1.0e-16) | -0.0499 (0.0049), CI: [-0.0595,-0.0402] | -10.1 (p < 1.0e-16) |
| log intake x male | 0.0114 (0.0095), CI: [-0.0072,0.0299] | 1.2 (p = 2.3e-01) | 0.0214 (0.0088), CI: [0.0041,0.0387] | 2.4 (p = 1.5e-02) |
| log intake x std. age | 0.0127 (0.0043), CI: [0.0042,0.0212] | 2.9 (p = 3.4e-03) | 0.0100 (0.0040), CI: [0.0020,0.0179] | 2.5 (p = 1.4e-02) |
| standardized age | -0.6012 (0.0040), CI: [-0.6091,-0.5933] | -149.9 (p < 1.0e-16) | -0.5989 (0.0040), CI: [-0.6066,-0.5911] | -151.2 (p < 1.0e-16) |
| standardized age$^2$ | -0.0389 (0.0036), CI: [-0.0459,-0.0319] | -11.0 (p < 1.0e-16) | -0.0388 (0.0035), CI: [-0.0457,-0.0319] | -11.0 (p < 1.0e-16) |
| | **Against model without log intake and interactions** Delta $R^2$: 0.0108 F-test: p < 1.0e-16 | | **Against model without log intake and interactions** Delta $R^2$: 0.0038 F-test: p < 1.0e-16 | |
