## Extended Data Table 2 for "Multimodal brain imaging study of 36,678 participants reveals adverse effects of moderate drinking"

**Extended Data Table 2. Regression results: Volume of white matter (whole brain, normalized).**
log intake is measured in standardized log(1 + daily units of alcohol).

| | **Standard Controls** | | **Extended Controls** | |
|---|---|---|---|---|
| | N: 36,678 (d.f.: 36,585), $R^2$: 0.122 | | N: 36,678 (d.f.: 36,578), $R^2$: 0.123 | |
| **Variable** | **Regression Coefficient (S.Err), 95% CI** | **t-stat (p-value)** | **Regression Coefficient (S.Err), 95% CI** | **t-stat (p-value)** |
| log intake | -0.0650 (0.0078), CI: [-0.0802,-0.0498] | -8.4 (p < 1.0e-16) | -0.0632 (0.0078), CI: [-0.0784,-0.0479] | -8.1 (p = 4.3e-16) |
| log intake$^2$ | -0.0370 (0.0050), CI: [-0.0468,-0.0273] | -7.5 (p = 7.8e-14) | -0.0378 (0.0050), CI: [-0.0475,-0.0280] | -7.6 (p = 2.9e-14) |
| log intake x male | 0.0164 (0.0107), CI: [-0.0046,0.0374] | 1.5 (p = 1.2e-01) | 0.0148 (0.0107), CI: [-0.0062,0.0358] | 1.4 (p = 1.7e-01) |
| log intake x std. age | 0.0111 (0.0050), CI: [0.0014,0.0208] | 2.2 (p = 2.5e-02) | 0.0110 (0.0050), CI: [0.0013,0.0207] | 2.2 (p = 2.6e-02) |
| standardized age | -0.3213 (0.0051), CI: [-0.3313,-0.3112] | -62.6 (p < 1.0e-16) | -0.3218 (0.0051), CI: [-0.3319,-0.3118] | -62.7 (p < 1.0e-16) |
| standardized age$^2$ | -0.0127 (0.0046), CI: [-0.0217,-0.0037] | -2.8 (p = 5.7e-03) | -0.0126 (0.0046), CI: [-0.0216,-0.0036] | -2.7 (p = 6.2e-03) |
| | **Against model without log intake and interactions**<br>Delta $R^2$: 0.0033<br>F-test: p < 1.0e-16 | | **Against model without log intake and interactions**<br>Delta $R^2$: 0.0032<br>F-test: p < 1.0e-16 | |

| | **Excluding Abstainers** | | **Excluding Heavy Drinkers** | |
|---|---|---|---|---|
| | N: 33,773 (d.f.: 33,676), R2: 0.124 | | N: 34,383 (d.f.: 34,286), R2: 0.124 | |
| **Variable** | **Regression Coefficient (S.Err), 95% CI** | **t-stat (p-value)** | **Regression Coefficient (S.Err), 95% CI** | **t-stat (p-value)** |
| log intake | -0.0599 (0.0087), CI: [-0.0769,-0.0429] | -6.9 (p = 4.6e-12) | -0.0596 (0.0098), CI: [-0.0789,-0.0403] | -6.1 (p = 1.4e-09) |
| log intake$^2$ | -0.0369 (0.0061), CI: [-0.0489,-0.0249] | -6.0 (p = 1.7e-09) | -0.0327 (0.0066), CI: [-0.0456,-0.0197] | -4.9 (p = 7.9e-07) |
| log intake x male | 0.0073 (0.0127), CI: [-0.0176,0.0323] | 0.6 (p = 5.7e-01) | 0.0148 (0.0119), CI: [-0.0084,0.0381] | 1.3 (p = 2.1e-01) |
| log intake x std. age | 0.0187 (0.0059), CI: [0.0072,0.0301] | 3.2 (p = 1.4e-03) | 0.0088 (0.0054), CI: [-0.0019,0.0194] | 1.6 (p = 1.1e-01) |
| standardized age | -0.3251 (0.0054), CI: [-0.3357,-0.3146] | -60.2 (p < 1.0e-16) | -0.3223 (0.0053), CI: [-0.3327,-0.3119] | -60.6 (p < 1.0e-16) |
| standardized age$^2$ | -0.0133 (0.0048), CI: [-0.0227,-0.0040] | -2.8 (p = 5.3e-03) | -0.0135 (0.0047), CI: [-0.0228,-0.0043] | -2.9 (p = 4.2e-03) |
| | **Against model without log intake and interactions**<br>Delta $R^2$: 0.0036<br>F-test: p < 1.0e-16 | | **Against model without log intake and interactions**<br>Delta $R^2$: 0.0014<br>F-test: p = 5.0e-13 | |
