## Extended Data Table 3 for "Multimodal brain imaging study of 36,678 participants reveals adverse effects of moderate drinking"

**Extended Data Table 3.** Regression results of regional gray matter volume IDPs.

| Name | beta_lin | beta_quad | beta_lin (se) | beta_quad (se) | delta_r2 | p-val | c95_lin (low) | c95_lin (high) | c95_quad (low) | c95_quad (high) | Sig |
|---|---|---|---|---|---|---|---|---|---|---|---|
| GMV in Precentral Gyrus (right) | -0.072 | -0.034 | 0.005 | 0.005 | 0.004 | 1.11E-16 | -0.083 | -0.062 | -0.043 | -0.025 | 1 |
| GMV in Brain-Stem | -0.072 | -0.028 | 0.005 | 0.005 | 0.004 | 1.11E-16 | -0.082 | -0.061 | -0.037 | -0.019 | 1 |
| GMV in Amygdala (right) | -0.062 | -0.047 | 0.005 | 0.005 | 0.004 | 1.11E-16 | -0.073 | -0.052 | -0.056 | -0.038 | 1 |
| GMV in Precentral Gyrus (left) | -0.068 | -0.032 | 0.005 | 0.005 | 0.004 | 1.11E-16 | -0.078 | -0.058 | -0.041 | -0.023 | 1 |
| GMV in Frontal Pole (right) | -0.064 | -0.038 | 0.005 | 0.005 | 0.004 | 1.11E-16 | -0.074 | -0.054 | -0.047 | -0.029 | 1 |
| GMV in Lateral Occipital Cortex, superior division (left) | -0.062 | -0.040 | 0.005 | 0.005 | 0.004 | 1.11E-16 | -0.072 | -0.051 | -0.049 | -0.031 | 1 |
| GMV in Frontal Pole (left) | -0.065 | -0.031 | 0.005 | 0.005 | 0.004 | 1.11E-16 | -0.076 | -0.055 | -0.040 | -0.023 | 1 |
| GMV in Frontal Orbital Cortex (right) | -0.062 | -0.034 | 0.005 | 0.005 | 0.003 | 1.11E-16 | -0.072 | -0.051 | -0.043 | -0.025 | 1 |
| GMV in Precuneous Cortex (right) | -0.061 | -0.033 | 0.005 | 0.005 | 0.003 | 1.11E-16 | -0.072 | -0.050 | -0.042 | -0.023 | 1 |
| GMV in Lateral Occipital Cortex, superior division (right) | -0.058 | -0.029 | 0.005 | 0.005 | 0.003 | 1.11E-16 | -0.069 | -0.048 | -0.038 | -0.020 | 1 |
| GMV in Postcentral Gyrus (right) | -0.056 | -0.031 | 0.005 | 0.004 | 0.003 | 1.11E-16 | -0.066 | -0.046 | -0.040 | -0.023 | 1 |
| GMV in Amygdala (left) | -0.050 | -0.039 | 0.005 | 0.005 | 0.003 | 1.11E-16 | -0.061 | -0.040 | -0.048 | -0.030 | 1 |
| GMV in Planum Polare (left) | -0.054 | -0.033 | 0.005 | 0.005 | 0.003 | 1.11E-16 | -0.065 | -0.044 | -0.042 | -0.024 | 1 |
| GMV in Planum Polare (right) | -0.056 | -0.029 | 0.005 | 0.004 | 0.003 | 1.11E-16 | -0.066 | -0.045 | -0.038 | -0.020 | 1 |
| GMV in Precuneous Cortex (left) | -0.055 | -0.026 | 0.005 | 0.005 | 0.003 | 1.11E-16 | -0.065 | -0.044 | -0.036 | -0.017 | 1 |
| GMV in Ventral Striatum (right) | -0.048 | -0.037 | 0.005 | 0.005 | 0.003 | 1.11E-16 | -0.059 | -0.038 | -0.046 | -0.028 | 1 |
| GMV in Insular Cortex (right) | -0.051 | -0.033 | 0.006 | 0.005 | 0.002 | 1.11E-16 | -0.062 | -0.040 | -0.042 | -0.023 | 1 |
| GMV in Putamen (right) | -0.053 | -0.028 | 0.006 | 0.005 | 0.002 | 1.11E-16 | -0.065 | -0.042 | -0.038 | -0.019 | 1 |
| GMV in VIIIa Cerebellum (left) | -0.037 | -0.044 | 0.005 | 0.005 | 0.002 | 1.11E-16 | -0.048 | -0.027 | -0.053 | -0.035 | 1 |
| GMV in Frontal Orbital Cortex (left) | -0.053 | -0.025 | 0.005 | 0.005 | 0.002 | 1.11E-16 | -0.064 | -0.043 | -0.034 | -0.016 | 1 |
| GMV in Middle Temporal Gyrus, posterior division (left) | -0.050 | -0.030 | 0.006 | 0.005 | 0.002 | 1.11E-16 | -0.061 | -0.040 | -0.040 | -0.021 | 1 |
| GMV in Postcentral Gyrus (left) | -0.052 | -0.026 | 0.005 | 0.004 | 0.002 | 1.11E-16 | -0.062 | -0.042 | -0.035 | -0.017 | 1 |
| GMV in Cuneal Cortex (left) | -0.054 | -0.017 | 0.006 | 0.005 | 0.002 | 1.11E-16 | -0.065 | -0.043 | -0.027 | -0.008 | 1 |
| GMV in Temporal Pole (right) | -0.051 | -0.027 | 0.006 | 0.005 | 0.002 | 1.11E-16 | -0.062 | -0.040 | -0.037 | -0.018 | 1 |
| GMV in Cuneal Cortex (right) | -0.053 | -0.019 | 0.006 | 0.005 | 0.002 | 1.11E-16 | -0.064 | -0.042 | -0.029 | -0.010 | 1 |
| GMV in Central Opercular Cortex | -0.050 | -0.027 | 0.005 | 0.005 | 0.002 | 1.11E-16 | -0.061 | -0.040 | -0.037 | -0.018 | 1 |

| | | | | | | | | | | | |
|---|---|---|---|---|---|---|---|---|---|---|---|
| (right) | | | | | | | | | | | |
| GMV in Central Opercular Cortex (left) | -0.050 | -0.026 | 0.005 | 0.005 | 0.002 | 1.11E-16 | -0.061 | -0.040 | -0.035 | -0.017 | 1 |
| GMV in Occipital Pole (right) | -0.052 | -0.015 | 0.006 | 0.005 | 0.002 | 1.11E-16 | -0.063 | -0.041 | -0.025 | -0.006 | 1 |
| GMV in Heschl's Gyrus (includes H1 and H2) (right) | -0.051 | -0.025 | 0.005 | 0.005 | 0.002 | 1.11E-16 | -0.061 | -0.040 | -0.034 | -0.016 | 1 |
| GMV in VIIIb Cerebellum (right) | -0.031 | -0.043 | 0.005 | 0.005 | 0.002 | 1.11E-16 | -0.042 | -0.020 | -0.052 | -0.034 | 1 |
| GMV in Insular Cortex (left) | -0.049 | -0.029 | 0.005 | 0.005 | 0.002 | 1.11E-16 | -0.059 | -0.038 | -0.038 | -0.019 | 1 |
| GMV in VIIIa Cerebellum (right) | -0.033 | -0.042 | 0.005 | 0.005 | 0.002 | 1.11E-16 | -0.043 | -0.022 | -0.051 | -0.033 | 1 |
| GMV in Middle Temporal Gyrus, posterior division (right) | -0.050 | -0.024 | 0.005 | 0.005 | 0.002 | 1.11E-16 | -0.061 | -0.039 | -0.033 | -0.015 | 1 |
| GMV in Putamen (left) | -0.051 | -0.014 | 0.006 | 0.005 | 0.002 | 1.11E-16 | -0.063 | -0.040 | -0.023 | -0.004 | 1 |
| GMV in Occipital Pole (left) | -0.050 | -0.022 | 0.006 | 0.005 | 0.002 | 1.11E-16 | -0.061 | -0.039 | -0.032 | -0.013 | 1 |
| GMV in Lateral Occipital Cortex, inferior division (right) | -0.046 | -0.030 | 0.006 | 0.005 | 0.002 | 1.11E-16 | -0.058 | -0.035 | -0.040 | -0.020 | 1 |
| GMV in Temporal Pole (left) | -0.048 | -0.027 | 0.006 | 0.005 | 0.002 | 1.11E-16 | -0.059 | -0.037 | -0.036 | -0.017 | 1 |
| GMV in VIIIb Cerebellum (left) | -0.032 | -0.040 | 0.006 | 0.005 | 0.002 | 1.11E-16 | -0.043 | -0.022 | -0.050 | -0.031 | 1 |
| GMV in Crus I Cerebellum (left) | -0.032 | -0.040 | 0.005 | 0.005 | 0.002 | 1.11E-16 | -0.043 | -0.022 | -0.049 | -0.031 | 1 |
| GMV in VIIb Cerebellum (left) | -0.037 | -0.037 | 0.006 | 0.005 | 0.002 | 1.11E-16 | -0.048 | -0.026 | -0.046 | -0.028 | 1 |
| GMV in Subcallosal Cortex (left) | -0.049 | -0.018 | 0.006 | 0.005 | 0.002 | 1.11E-16 | -0.060 | -0.038 | -0.028 | -0.009 | 1 |
| GMV in Ventral Striatum (left) | -0.045 | -0.028 | 0.005 | 0.005 | 0.002 | 1.11E-16 | -0.055 | -0.034 | -0.037 | -0.019 | 1 |
| GMV in Superior Temporal Gyrus, posterior division (right) | -0.046 | -0.022 | 0.006 | 0.005 | 0.002 | 2.22E-16 | -0.057 | -0.035 | -0.032 | -0.013 | 1 |
| GMV in VI Cerebellum (left) | -0.028 | -0.039 | 0.005 | 0.005 | 0.002 | 1.11E-16 | -0.039 | -0.018 | -0.048 | -0.030 | 1 |
| GMV in Crus I Cerebellum (right) | -0.026 | -0.037 | 0.005 | 0.005 | 0.002 | 5.55E-16 | -0.037 | -0.016 | -0.047 | -0.028 | 1 |
| GMV in VI Cerebellum (right) | -0.031 | -0.035 | 0.005 | 0.005 | 0.002 | 5.55E-16 | -0.042 | -0.021 | -0.045 | -0.026 | 1 |
| GMV in Subcallosal Cortex (right) | -0.045 | -0.014 | 0.006 | 0.005 | 0.002 | 1.22E-13 | -0.056 | -0.033 | -0.024 | -0.004 | 1 |
| GMV in Superior Frontal Gyrus (right) | -0.043 | -0.021 | 0.006 | 0.005 | 0.002 | 9.25E-14 | -0.054 | -0.032 | -0.031 | -0.011 | 1 |
| GMV in Inferior Frontal Gyrus, pars opercularis (left) | -0.043 | -0.019 | 0.006 | 0.005 | 0.002 | 4.94E-14 | -0.054 | -0.032 | -0.028 | -0.009 | 1 |
| GMV in Middle Frontal Gyrus (left) | -0.042 | -0.021 | 0.006 | 0.005 | 0.002 | 1.16E-13 | -0.053 | -0.031 | -0.031 | -0.012 | 1 |
| GMV in Paracingulate Gyrus (right) | -0.037 | -0.030 | 0.005 | 0.005 | 0.002 | 1.22E-15 | -0.047 | -0.027 | -0.038 | -0.021 | 1 |
| GMV in Superior Frontal Gyrus (left) | -0.043 | -0.017 | 0.006 | 0.005 | 0.002 | 7.45E-14 | -0.054 | -0.032 | -0.026 | -0.007 | 1 |

| | | | | | | | | | | | |
|---|---|---|---|---|---|---|---|---|---|---|---|
| GMV in Occipital Fusiform Gyrus (left) | -0.041 | -0.023 | 0.006 | 0.005 | 0.002 | 3.81E-14 | -0.052 | -0.031 | -0.032 | -0.014 | 1 |
| GMV in Frontal Operculum Cortex (left) | -0.040 | -0.026 | 0.006 | 0.005 | 0.002 | 2.08E-14 | -0.050 | -0.029 | -0.036 | -0.017 | 1 |
| GMV in Juxtapositional Lobule Cortex (right) | -0.040 | -0.025 | 0.006 | 0.005 | 0.001 | 3.46E-13 | -0.051 | -0.028 | -0.034 | -0.015 | 1 |
| GMV in Intracalcarine Cortex (right) | -0.042 | -0.018 | 0.006 | 0.005 | 0.001 | 8.65E-13 | -0.053 | -0.031 | -0.028 | -0.008 | 1 |
| GMV in VIIb Cerebellum (right) | -0.031 | -0.033 | 0.006 | 0.005 | 0.001 | 1.14E-13 | -0.042 | -0.020 | -0.042 | -0.023 | 1 |
| GMV in Superior Temporal Gyrus, posterior division (left) | -0.040 | -0.024 | 0.006 | 0.005 | 0.001 | 4.15E-13 | -0.051 | -0.029 | -0.033 | -0.014 | 1 |
| GMV in Paracingulate Gyrus (left) | -0.039 | -0.023 | 0.005 | 0.005 | 0.001 | 5.38E-14 | -0.050 | -0.029 | -0.032 | -0.014 | 1 |
| GMV in Vermis VIIIb Cerebellum | -0.030 | -0.032 | 0.006 | 0.005 | 0.001 | 5.92E-13 | -0.041 | -0.019 | -0.042 | -0.023 | 1 |
| GMV in IX Cerebellum (right) | -0.028 | -0.033 | 0.005 | 0.005 | 0.001 | 2.51E-13 | -0.038 | -0.017 | -0.043 | -0.024 | 1 |
| GMV in Superior Temporal Gyrus, anterior division (left) | -0.041 | -0.015 | 0.006 | 0.005 | 0.001 | 5.60E-12 | -0.052 | -0.030 | -0.025 | -0.006 | 1 |
| GMV in Vermis VIIIa Cerebellum | -0.031 | -0.030 | 0.005 | 0.005 | 0.001 | 6.48E-13 | -0.042 | -0.020 | -0.040 | -0.021 | 1 |
| GMV in Supramarginal Gyrus, posterior division (right) | -0.038 | -0.020 | 0.006 | 0.005 | 0.001 | 1.34E-11 | -0.049 | -0.027 | -0.030 | -0.011 | 1 |
| GMV in Supracalcarine Cortex (right) | -0.039 | -0.016 | 0.006 | 0.005 | 0.001 | 4.76E-11 | -0.051 | -0.028 | -0.026 | -0.007 | 1 |
| GMV in X Cerebellum (right) | -0.024 | -0.032 | 0.006 | 0.005 | 0.001 | 8.48E-12 | -0.035 | -0.013 | -0.042 | -0.023 | 1 |
| GMV in Heschl's Gyrus (includes H1 and H2) (left) | -0.036 | -0.024 | 0.005 | 0.005 | 0.001 | 5.33E-12 | -0.046 | -0.025 | -0.033 | -0.014 | 1 |
| GMV in Angular Gyrus (left) | -0.033 | -0.026 | 0.006 | 0.005 | 0.001 | 8.74E-11 | -0.045 | -0.022 | -0.036 | -0.016 | 1 |
| GMV in Cingulate Gyrus, posterior division (left) | -0.035 | -0.024 | 0.006 | 0.005 | 0.001 | 1.01E-10 | -0.046 | -0.024 | -0.034 | -0.014 | 1 |
| GMV in Frontal Operculum Cortex (right) | -0.037 | -0.021 | 0.006 | 0.005 | 0.001 | 2.46E-11 | -0.048 | -0.026 | -0.030 | -0.011 | 1 |
| GMV in Cingulate Gyrus, posterior division (right) | -0.035 | -0.022 | 0.006 | 0.005 | 0.001 | 2.94E-10 | -0.047 | -0.024 | -0.032 | -0.012 | 1 |
| GMV in Crus II Cerebellum (left) | -0.027 | -0.029 | 0.006 | 0.005 | 0.001 | 3.96E-11 | -0.038 | -0.017 | -0.039 | -0.020 | 1 |
| GMV in Hippocampus (left) | -0.033 | -0.025 | 0.005 | 0.005 | 0.001 | 1.14E-11 | -0.043 | -0.022 | -0.034 | -0.016 | 1 |
| GMV in Middle Frontal Gyrus (right) | -0.038 | -0.013 | 0.006 | 0.005 | 0.001 | 1.96E-10 | -0.049 | -0.027 | -0.023 | -0.003 | 1 |
| GMV in Occipital Fusiform Gyrus (right) | -0.034 | -0.023 | 0.006 | 0.005 | 0.001 | 6.30E-11 | -0.045 | -0.024 | -0.032 | -0.013 | 1 |
| GMV in IX Cerebellum (left) | -0.028 | -0.028 | 0.005 | 0.005 | 0.001 | 4.88E-11 | -0.039 | -0.018 | -0.038 | -0.019 | 1 |

| | | | | | | | | | | | |
|---|---|---|---|---|---|---|---|---|---|---|---|
| GMV in Superior Temporal Gyrus, anterior division (right) | -0.036 | -0.019 | 0.006 | 0.005 | 0.001 | 3.30E-10 | -0.047 | -0.025 | -0.029 | -0.010 | 1 |
| GMV in Frontal Medial Cortex (left) | -0.037 | -0.013 | 0.006 | 0.005 | 0.001 | 1.89E-10 | -0.048 | -0.026 | -0.023 | -0.004 | 1 |
| GMV in Supramarginal Gyrus, posterior division (left) | -0.035 | -0.021 | 0.006 | 0.005 | 0.001 | 6.55E-10 | -0.046 | -0.024 | -0.031 | -0.011 | 1 |
| GMV in Vermis VIIb Cerebellum | -0.029 | -0.027 | 0.005 | 0.005 | 0.001 | 1.17E-10 | -0.039 | -0.018 | -0.037 | -0.018 | 1 |
| GMV in Frontal Medial Cortex (right) | -0.037 | -0.012 | 0.006 | 0.005 | 0.001 | 4.35E-10 | -0.048 | -0.026 | -0.021 | -0.002 | 1 |
| GMV in Temporal Fusiform Cortex, anterior division (left) | -0.035 | -0.018 | 0.006 | 0.005 | 0.001 | 7.79E-10 | -0.046 | -0.024 | -0.028 | -0.009 | 1 |
| GMV in Superior Parietal Lobule (left) | -0.036 | -0.015 | 0.006 | 0.005 | 0.001 | 8.39E-10 | -0.047 | -0.025 | -0.025 | -0.005 | 1 |
| GMV in Lateral Occipital Cortex, inferior division (left) | -0.035 | -0.019 | 0.006 | 0.005 | 0.001 | 1.62E-09 | -0.046 | -0.023 | -0.028 | -0.009 | 1 |
| GMV in X Cerebellum (left) | -0.023 | -0.029 | 0.006 | 0.005 | 0.001 | 8.04E-10 | -0.034 | -0.012 | -0.039 | -0.020 | 1 |
| GMV in Middle Temporal Gyrus, anterior division (left) | -0.027 | -0.027 | 0.006 | 0.005 | 0.001 | 2.46E-09 | -0.038 | -0.016 | -0.037 | -0.018 | 1 |
| GMV in Inferior Frontal Gyrus, pars opercularis (right) | -0.036 | -0.015 | 0.006 | 0.005 | 0.001 | 1.55E-09 | -0.047 | -0.024 | -0.025 | -0.005 | 1 |
| GMV in Crus II Cerebellum (right) | -0.026 | -0.028 | 0.005 | 0.005 | 0.001 | 4.06E-10 | -0.037 | -0.015 | -0.037 | -0.018 | 1 |
| GMV in Intracalcarine Cortex (left) | -0.035 | -0.014 | 0.006 | 0.005 | 0.001 | 1.04E-08 | -0.046 | -0.023 | -0.024 | -0.005 | 1 |
| GMV in Hippocampus (right) | -0.030 | -0.024 | 0.005 | 0.005 | 0.001 | 6.05E-10 | -0.040 | -0.019 | -0.033 | -0.015 | 1 |
| GMV in Temporal Fusiform Cortex, anterior division (right) | -0.029 | -0.024 | 0.006 | 0.005 | 0.001 | 9.11E-09 | -0.040 | -0.018 | -0.033 | -0.014 | 1 |
| GMV in Juxtapositional Lobule Cortex (left) | -0.033 | -0.015 | 0.006 | 0.005 | 0.001 | 1.65E-08 | -0.045 | -0.022 | -0.025 | -0.005 | 1 |
| GMV in Thalamus (left) | -0.034 | -0.006 | 0.006 | 0.005 | 0.001 | 7.43E-09 | -0.045 | -0.023 | -0.016 | 0.003 | 1 |
| GMV in Supracalcarine Cortex (left) | -0.034 | -0.008 | 0.006 | 0.005 | 0.001 | 3.97E-08 | -0.045 | -0.023 | -0.018 | 0.001 | 1 |
| GMV in Temporal Fusiform Cortex, posterior division (right) | -0.026 | -0.025 | 0.006 | 0.005 | 0.001 | 1.13E-08 | -0.036 | -0.015 | -0.035 | -0.016 | 1 |
| GMV in Angular Gyrus (right) | -0.031 | -0.019 | 0.006 | 0.005 | 0.001 | 3.62E-08 | -0.042 | -0.020 | -0.029 | -0.010 | 1 |
| GMV in Temporal Occipital Fusiform Cortex (left) | -0.026 | -0.024 | 0.006 | 0.005 | 0.001 | 6.35E-08 | -0.037 | -0.015 | -0.034 | -0.014 | 1 |
| GMV in Inferior Frontal Gyrus, pars triangularis (left) | -0.032 | -0.015 | 0.006 | 0.005 | 0.001 | 8.86E-08 | -0.043 | -0.021 | -0.024 | -0.005 | 1 |
| GMV in Inferior Temporal Gyrus, anterior division (right) | -0.026 | -0.023 | 0.006 | 0.005 | 0.001 | 2.42E-07 | -0.037 | -0.015 | -0.033 | -0.013 | 1 |
| GMV in Superior Parietal Lobule (right) | -0.032 | -0.006 | 0.006 | 0.005 | 0.001 | 1.32E-07 | -0.043 | -0.021 | -0.016 | 0.003 | 1 |

| | | | | | | | | | | |
|---|---|---|---|---|---|---|---|---|---|---|
| GMV in V Cerebellum (left) | -0.014 | -0.027 | 0.006 | 0.005 | 0.001 | 1.94E-07 | -0.025 | -0.003 | -0.036 | -0.017 | 1 |
| GMV in Middle Temporal Gyrus, temporooccipital part (right) | -0.027 | -0.020 | 0.006 | 0.005 | 0.001 | 6.05E-07 | -0.038 | -0.015 | -0.030 | -0.010 | 1 |
| GMV in Inferior Temporal Gyrus, posterior division (right) | -0.028 | -0.018 | 0.006 | 0.005 | 0.001 | 4.92E-07 | -0.040 | -0.017 | -0.028 | -0.008 | 1 |
| GMV in Temporal Fusiform Cortex, posterior division (left) | -0.023 | -0.023 | 0.006 | 0.005 | 0.001 | 1.99E-07 | -0.034 | -0.012 | -0.033 | -0.014 | 1 |
| GMV in Vermis VI Cerebellum | -0.023 | -0.023 | 0.005 | 0.005 | 0.001 | 1.25E-07 | -0.033 | -0.012 | -0.033 | -0.014 | 1 |
| GMV in V Cerebellum (right) | -0.014 | -0.026 | 0.006 | 0.005 | 0.001 | 3.04E-07 | -0.025 | -0.003 | -0.036 | -0.017 | 1 |
| GMV in Middle Temporal Gyrus, anterior division (right) | -0.025 | -0.021 | 0.006 | 0.005 | 0.001 | 9.37E-07 | -0.036 | -0.013 | -0.031 | -0.012 | 1 |
| GMV in Supramarginal Gyrus, anterior division (right) | -0.027 | -0.018 | 0.006 | 0.005 | 0.001 | 1.57E-06 | -0.038 | -0.015 | -0.028 | -0.008 | 1 |
| GMV in Lingual Gyrus (right) | -0.023 | -0.022 | 0.006 | 0.005 | 0.001 | 1.39E-06 | -0.034 | -0.012 | -0.031 | -0.012 | 1 |
| GMV in Planum Temporale (right) | -0.028 | -0.016 | 0.006 | 0.005 | 0.001 | 1.56E-06 | -0.039 | -0.016 | -0.025 | -0.006 | 1 |
| GMV in Vermis IX Cerebellum | -0.016 | -0.024 | 0.006 | 0.005 | 0.001 | 1.82E-06 | -0.027 | -0.004 | -0.034 | -0.015 | 1 |
| GMV in Supramarginal Gyrus, anterior division (left) | -0.025 | -0.018 | 0.006 | 0.005 | 0.001 | 3.88E-06 | -0.037 | -0.014 | -0.028 | -0.009 | 1 |
| GMV in Thalamus (right) | -0.025 | 0.002 | 0.006 | 0.005 | 0.001 | 4.83E-06 | -0.036 | -0.014 | -0.007 | 0.012 | 1 |
| GMV in Parietal Operculum Cortex (left) | -0.026 | -0.016 | 0.006 | 0.005 | 0.001 | 8.21E-06 | -0.037 | -0.015 | -0.025 | -0.006 | 1 |
| GMV in Inferior Temporal Gyrus, posterior division (left) | -0.025 | -0.017 | 0.006 | 0.005 | 0.001 | 1.09E-05 | -0.036 | -0.013 | -0.027 | -0.007 | 1 |
| GMV in Inferior Temporal Gyrus, anterior division (left) | -0.027 | -0.013 | 0.006 | 0.005 | 0.001 | 1.25E-05 | -0.038 | -0.015 | -0.023 | -0.004 | 1 |
| GMV in Middle Temporal Gyrus, temporooccipital part (left) | -0.025 | -0.015 | 0.006 | 0.005 | 0.001 | 1.48E-05 | -0.037 | -0.014 | -0.025 | -0.006 | 1 |
| GMV in Temporal Occipital Fusiform Cortex (right) | -0.016 | -0.022 | 0.006 | 0.005 | 0.001 | 1.63E-05 | -0.027 | -0.004 | -0.032 | -0.013 | 1 |
| GMV in I-IV Cerebellum (right) | -0.017 | -0.021 | 0.006 | 0.005 | 0.001 | 1.29E-05 | -0.028 | -0.006 | -0.031 | -0.012 | 1 |
| GMV in Pallidum (right) | 0.025 | 0.000 | 0.006 | 0.005 | 0.001 | 2.54E-05 | 0.014 | 0.036 | -0.009 | 0.010 | 1 |
| GMV in Vermis Crus II Cerebellum | -0.020 | -0.020 | 0.006 | 0.005 | 0.001 | 2.12E-05 | -0.031 | -0.009 | -0.029 | -0.010 | 1 |
| GMV in Parahippocampal Gyrus, anterior division (right) | -0.021 | -0.018 | 0.006 | 0.005 | 0.001 | 4.25E-05 | -0.032 | -0.010 | -0.028 | -0.008 | 1 |
| GMV in Vermis X Cerebellum | -0.019 | -0.019 | 0.005 | 0.005 | 0.001 | 1.35E-05 | -0.030 | -0.009 | -0.028 | -0.010 | 1 |
| GMV in Parietal Operculum Cortex (right) | -0.022 | -0.016 | 0.006 | 0.005 | 0.000 | 7.79E-05 | -0.033 | -0.011 | -0.026 | -0.006 | 1 |
| GMV in Inferior Frontal Gyrus, pars | -0.024 | -0.010 | 0.006 | 0.005 | 0.000 | 1.19E-04 | -0.035 | -0.013 | -0.020 | -0.001 | 1 |

| | | | | | | | | | | | |
|---|---|---|---|---|---|---|---|---|---|---|---|
| triangularis (right) | | | | | | | | | | | |
| GMV in Pallidum (left) | 0.023 | 0.003 | 0.006 | 0.005 | 0.000 | 1.88E-04 | 0.012 | 0.035 | -0.007 | 0.012 | 0 |
| GMV in Planum Temporale (left) | -0.022 | -0.013 | 0.006 | 0.005 | 0.000 | 2.09E-04 | -0.033 | -0.011 | -0.023 | -0.004 | 0 |
| GMV in Inferior Temporal Gyrus, temporooccipital part (left) | -0.020 | -0.016 | 0.006 | 0.005 | 0.000 | 3.73E-04 | -0.031 | -0.008 | -0.026 | -0.006 | 0 |
| GMV in Lingual Gyrus (left) | -0.010 | -0.020 | 0.006 | 0.005 | 0.000 | 3.48E-04 | -0.021 | 0.001 | -0.029 | -0.010 | 0 |
| GMV in Parahippocampal Gyrus, posterior division (left) | -0.021 | -0.013 | 0.006 | 0.005 | 0.000 | 2.50E-04 | -0.032 | -0.010 | -0.022 | -0.003 | 0 |
| GMV in Parahippocampal Gyrus, posterior division (right) | -0.021 | -0.011 | 0.006 | 0.005 | 0.000 | 3.73E-04 | -0.032 | -0.010 | -0.021 | -0.002 | 0 |
| GMV in Parahippocampal Gyrus, anterior division (left) | -0.019 | -0.014 | 0.006 | 0.005 | 0.000 | 7.46E-04 | -0.030 | -0.008 | -0.023 | -0.004 | 0 |
| GMV in I-IV Cerebellum (left) | -0.013 | -0.018 | 0.006 | 0.005 | 0.000 | 6.37E-04 | -0.024 | -0.002 | -0.027 | -0.008 | 0 |
| GMV in Inferior Temporal Gyrus, temporooccipital part (right) | 0.005 | -0.013 | 0.006 | 0.005 | 0.000 | 7.28E-04 | -0.006 | 0.017 | -0.023 | -0.003 | 0 |
| GMV in Vermis Crus I Cerebellum | -0.012 | -0.004 | 0.006 | 0.005 | 0.000 | 0.102 | -0.024 | -0.001 | -0.013 | 0.006 | 0 |
| GMV in Caudate (right) | -0.011 | 0.001 | 0.006 | 0.005 | 0.000 | 0.112 | -0.022 | 0.001 | -0.008 | 0.011 | 0 |
| GMV in Caudate (left) | -0.008 | 0.004 | 0.006 | 0.005 | 0.000 | 0.154 | -0.019 | 0.003 | -0.006 | 0.014 | 0 |
| GMV in Cingulate Gyrus, anterior division (right) | -0.007 | 0.003 | 0.006 | 0.005 | 0.000 | 0.302 | -0.018 | 0.005 | -0.007 | 0.013 | 0 |
| GMV in Cingulate Gyrus, anterior division (left) | -0.006 | 0.000 | 0.006 | 0.005 | 0.000 | 0.580 | -0.017 | 0.006 | -0.010 | 0.009 | 0 |

*Note*. Model: IDP: gray matter volume in region of interest, normalized for head size, x: standardized log(1 + daily units)

IDP = b0 + b1 x + b2 x^2+ bc controls + error term

List of control variables: standardized age, squared standardized age, genetic principal components 1 to 40, standardized height, handedness, sex (female:0, male:1), standardized townsend, county of residence, current smoker, former light smoker, former heavy smoker

GMV = gray matter volume, beta_lin = regression coefficient b1, for standardized log(1 + daily units), beta_quad = regression coefficient b2, for squared standardized log(1 + daily units), beta_lin (se) = standard error for b1, beta_quad (se)= standard error for b2, delta_r2 = difference in r-squared values for the regressions with and without intake and squared intake, p-val = p-value of the F-test of joint significance for b1 and b2, c95_lin (low) = 95% confidence interval for b1 (lower bound), c95_lin (high) = 95% confidence interval for b1 (upper bound), c95_quad (low) = 95% confidence interval for b2 (lower bound), c95_quad (high) = 95% confidence interval for b2 (upper bound), Sig = family-wise error rate (FWER) corrected significance (Holm method).
