## Extended Data Table 4 for "Multimodal brain imaging study of 36,678 participants reveals adverse effects of moderate drinking"

**Extended Data Table 4.** Regression results for white matter microstructure IDPs.

| Name | beta_lin | beta_quad | beta_lin (se) | beta_quad (se) | delta_r2 | p-val | c95_lin (low) | c95_lin (high) | c95_quad (low) | c95_quad (high) | Sig |
|---|---|---|---|---|---|---|---|---|---|---|---|
| Mean FA in anterior corona radiata (left) | -0.041 | -0.020 | 0.005 | 0.004 | 0.001 | 1.11E-16 | -0.050 | -0.032 | -0.028 | -0.012 | 1 |
| Mean FA in anterior corona radiata (right) | -0.038 | -0.018 | 0.005 | 0.004 | 0.001 | 3.00E-15 | -0.048 | -0.029 | -0.027 | -0.010 | 1 |
| Mean FA in anterior limb of internal capsule (left) | -0.027 | -0.012 | 0.005 | 0.004 | 0.001 | 9.91E-08 | -0.037 | -0.018 | -0.021 | -0.004 | 1 |
| Mean FA in anterior limb of internal capsule (right) | -0.026 | -0.012 | 0.005 | 0.004 | 0.001 | 1.84E-07 | -0.036 | -0.017 | -0.021 | -0.004 | 1 |
| Mean FA in body of corpus callosum | -0.031 | -0.014 | 0.005 | 0.004 | 0.001 | 5.87E-11 | -0.040 | -0.022 | -0.022 | -0.006 | 1 |
| Mean FA in cerebral peduncle (left) | -0.022 | -0.010 | 0.005 | 0.004 | 4.11E-04 | 6.15E-06 | -0.032 | -0.013 | -0.018 | -0.002 | 1 |
| Mean FA in cerebral peduncle (right) | -0.021 | -0.010 | 0.005 | 0.004 | 3.62E-04 | 3.40E-05 | -0.030 | -0.012 | -0.017 | -0.001 | 1 |
| Mean FA in cingulum cingulate gyrus (left) | -0.041 | -0.016 | 0.005 | 0.004 | 0.001 | 1.67E-15 | -0.050 | -0.031 | -0.024 | -0.007 | 1 |
| Mean FA in cingulum cingulate gyrus (right) | -0.044 | -0.019 | 0.005 | 0.004 | 0.002 | 1.11E-16 | -0.054 | -0.034 | -0.027 | -0.010 | 1 |
| Mean FA in cingulum hippocampus (left) | -0.014 | -0.010 | 0.005 | 0.004 | 0.002 | 0.020 | -0.024 | -0.004 | -0.015 | 0.002 | 0 |
| Mean FA in cingulum hippocampus (right) | -0.012 | -0.002 | 0.005 | 0.005 | 1.10E-04 | 0.081 | -0.022 | -0.001 | -0.011 | 0.007 | 0 |
| Mean FA in corticospinal tract (left) | -0.002 | -0.005 | 0.005 | 0.004 | 2.49E-05 | 0.558 | -0.012 | 0.008 | -0.014 | 0.004 | 0 |
| Mean FA in corticospinal tract (right) | -0.004 | -0.004 | 0.005 | 0.005 | 2.40E-05 | 0.595 | -0.015 | 0.006 | -0.013 | 0.005 | 0 |
| Mean FA in external capsule (left) | -0.018 | -0.011 | 0.005 | 0.004 | 2.90E-04 | 2.50E-04 | -0.027 | -0.008 | -0.019 | -0.003 | 0 |
| Mean FA in external capsule (right) | -0.022 | -0.012 | 0.005 | 0.004 | 4.14E-04 | 5.40E-06 | -0.031 | -0.013 | -0.020 | -0.004 | 1 |
| Mean FA in fornix cres+stria terminalis (left) | -0.041 | -0.022 | 0.005 | 0.004 | 0.001 | 1.11E-16 | -0.050 | -0.032 | -0.029 | -0.014 | 1 |
| Mean FA in fornix cres+stria terminalis (right) | -0.042 | -0.022 | 0.005 | 0.004 | 0.002 | 1.11E-16 | -0.051 | -0.033 | -0.030 | -0.014 | 1 |
| Mean FA in fornix | -0.058 | -0.039 | 0.005 | 0.004 | 0.003 | 1.11E-16 | -0.067 | -0.048 | -0.047 | -0.031 | 1 |
| Mean FA in genu of corpus callosum | -0.045 | -0.025 | 0.005 | 0.004 | 0.002 | 1.11E-16 | -0.054 | -0.035 | -0.033 | -0.017 | 1 |

| | | | | | | | | | | | |
|---|---|---|---|---|---|---|---|---|---|---|---|
| Mean FA in inferior cerebellar peduncle (left) | -0.019 | -0.014 | 0.005 | 0.004 | 3.69E-04 | 6.53E-05 | -0.029 | -0.010 | -0.022 | -0.005 | 1 |
| Mean FA in inferior cerebellar peduncle (right) | -0.023 | -0.016 | 0.005 | 0.004 | 5.37E-04 | 6.82E-07 | -0.033 | -0.014 | -0.025 | -0.008 | 1 |
| Mean FA in medial lemniscus (left) | -0.015 | -0.005 | 0.005 | 0.004 | 2.67E-04 | 0.015 | -0.024 | -0.005 | -0.014 | 0.003 | 0 |
| Mean FA in medial lemniscus (right) | -0.011 | -0.005 | 0.005 | 0.004 | 1.08E-04 | 0.067 | -0.021 | -0.002 | -0.014 | 0.003 | 0 |
| Mean FA in middle cerebellar peduncle | -0.010 | -0.002 | 0.005 | 0.004 | 7.16E-05 | 0.139 | -0.019 | 0.000 | -0.010 | 0.006 | 0 |
| Mean FA in pontine crossing tract | -0.010 | 0.006 | 0.005 | 0.004 | 1.19E-04 | 0.054 | -0.017 | 0.003 | -0.003 | 0.015 | 0 |
| Mean FA in posterior corona radiata (left) | -0.020 | -0.006 | 0.005 | 0.004 | 3.10E-04 | 2.25E-04 | -0.029 | -0.010 | -0.014 | 0.002 | 0 |
| Mean FA in posterior corona radiata (right) | -0.014 | -0.002 | 0.005 | 0.004 | 1.62E-04 | 0.013 | -0.024 | -0.005 | -0.010 | 0.006 | 0 |
| Mean FA in posterior limb of internal capsule (left) | -0.013 | -0.001 | 0.005 | 0.004 | 1.38E-04 | 0.019 | -0.022 | -0.004 | -0.009 | 0.007 | 0 |
| Mean FA in posterior limb of internal capsule (right) | -0.011 | 0.001 | 0.005 | 0.004 | 1.19E-04 | 0.036 | -0.021 | -0.002 | -0.007 | 0.009 | 0 |
| Mean FA in posterior thalamic radiation (left) | -0.020 | -0.011 | 0.005 | 0.004 | 3.52E-04 | 4.29E-05 | -0.029 | -0.011 | -0.019 | -0.003 | 1 |
| Mean FA in posterior thalamic radiation (right) | -0.020 | -0.012 | 0.005 | 0.004 | 3.56E-04 | 3.84E-05 | -0.029 | -0.011 | -0.020 | -0.004 | 1 |
| Mean FA in retrolenticular part of internal capsule (left) | -0.012 | -0.002 | 0.005 | 0.004 | 1.09E-04 | 0.042 | -0.021 | -0.003 | -0.010 | 0.005 | 0 |
| Mean FA in retrolenticular part of internal capsule (right) | -0.009 | -0.001 | 0.005 | 0.004 | 6.79E-05 | 0.162 | -0.019 | 0.000 | -0.009 | 0.007 | 0 |
| Mean FA in sagittal stratum (left) | -0.016 | -0.007 | 0.005 | 0.004 | 2.07E-04 | 0.003 | -0.025 | -0.006 | -0.016 | 0.001 | 0 |
| Mean FA in sagittal stratum (right) | -0.015 | -0.004 | 0.005 | 0.004 | 1.70E-04 | 0.010 | -0.024 | -0.005 | -0.013 | 0.004 | 0 |
| Mean FA in splenium of corpus callosum | -0.013 | -0.002 | 0.005 | 0.004 | 1.32E-04 | 0.019 | -0.022 | -0.004 | -0.010 | 0.006 | 0 |
| Mean FA in superior cerebellar peduncle (left) | -0.018 | -0.013 | 0.005 | 0.004 | 3.28E-04 | 1.04E-04 | -0.027 | -0.008 | -0.021 | -0.005 | 1 |
| Mean FA in superior cerebellar peduncle (right) | -0.019 | -0.014 | 0.005 | 0.004 | 3.72E-04 | 2.91E-05 | -0.028 | -0.009 | -0.022 | -0.006 | 1 |
| Mean FA in superior corona radiata (left) | -0.026 | -0.011 | 0.005 | 0.004 | 5.50E-04 | 1.76E-07 | -0.035 | -0.017 | -0.019 | -0.003 | 1 |
| Mean FA in superior corona radiata (right) | -0.022 | -0.007 | 0.005 | 0.004 | 3.92E-04 | 1.42E-05 | -0.032 | -0.013 | -0.015 | 0.001 | 1 |

| | | | | | | | | | | | |
|---|---|---|---|---|---|---|---|---|---|---|---|
| Mean FA in superior fronto-occipital fasciculus (left) | -0.038 | -0.028 | 0.005 | 0.004 | 0.001 | 2.22E-16 | -0.048 | -0.028 | -0.036 | -0.019 | 1 |
| Mean FA in superior fronto-occipital fasciculus (right) | -0.035 | -0.026 | 0.005 | 0.004 | 0.001 | 5.15E-14 | -0.045 | -0.025 | -0.035 | -0.017 | 1 |
| Mean FA in superior longitudinal fasciculus (left) | -0.020 | -0.010 | 0.005 | 0.004 | 3.49E-04 | 7.05E-05 | -0.030 | -0.011 | -0.019 | -0.002 | 1 |
| Mean FA in superior longitudinal fasciculus (right) | -0.018 | -0.012 | 0.005 | 0.004 | 3.16E-04 | 1.92E-04 | -0.028 | -0.009 | -0.020 | -0.003 | 0 |
| Mean FA in tapetum (left) | -0.022 | -0.007 | 0.005 | 0.005 | 3.86E-04 | 1.41E-04 | -0.032 | -0.012 | -0.016 | 0.002 | 1 |
| Mean FA in tapetum (right) | -0.028 | -0.015 | 0.005 | 0.004 | 6.86E-04 | 6.60E-08 | -0.038 | -0.018 | -0.024 | -0.006 | 1 |
| Mean FA in uncinate fasciculus (left) | -0.016 | -0.011 | 0.005 | 0.004 | 2.44E-04 | 0.003 | -0.026 | -0.006 | -0.019 | -0.002 | 0 |
| Mean FA in uncinate fasciculus (right) | -0.015 | -0.010 | 0.005 | 0.005 | 2.18E-04 | 0.006 | -0.025 | -0.005 | -0.019 | -0.001 | 0 |
| Mean ICVF in anterior corona radiata (left) | -0.033 | -0.010 | 0.005 | 0.004 | 8.48E-04 | 4.16E-10 | -0.043 | -0.023 | -0.019 | -0.002 | 1 |
| Mean ICVF in anterior corona radiata (right) | -0.034 | -0.011 | 0.005 | 0.004 | 8.95E-04 | 1.07E-10 | -0.043 | -0.024 | -0.020 | -0.004 | 1 |
| Mean ICVF in anterior limb of internal capsule (left) | -0.024 | -0.008 | 0.005 | 0.004 | 4.59E-04 | 2.64E-06 | -0.033 | -0.015 | -0.016 | 0.000 | 1 |
| Mean ICVF in anterior limb of internal capsule (right) | -0.027 | -0.010 | 0.005 | 0.004 | 5.66E-04 | 9.58E-08 | -0.036 | -0.017 | -0.018 | -0.002 | 1 |
| Mean ICVF in body of corpus callosum | -0.013 | -0.001 | 0.005 | 0.004 | 1.48E-04 | 0.018 | -0.023 | -0.004 | -0.009 | 0.007 | 0 |
| Mean ICVF in cerebral peduncle (left) | -0.005 | 0.007 | 0.005 | 0.004 | 9.94E-05 | 0.071 | -0.014 | 0.005 | -0.001 | 0.015 | 0 |
| Mean ICVF in cerebral peduncle (right) | -0.007 | 0.008 | 0.005 | 0.004 | 1.52E-04 | 0.0196 | -0.017 | 0.002 | -0.001 | 0.016 | 0 |
| Mean ICVF in cingulum cingulate gyrus (left) | -0.033 | -0.012 | 0.005 | 0.004 | 8.71E-04 | 2.91E-10 | -0.043 | -0.023 | -0.020 | -0.003 | 1 |
| Mean ICVF in cingulum cingulate gyrus (right) | -0.030 | -0.011 | 0.005 | 0.004 | 7.08E-04 | 1.95E-08 | -0.040 | -0.020 | -0.020 | -0.003 | 1 |
| Mean ICVF in cingulum hippocampus (left) | -0.019 | 1.50E-04 | 0.005 | 0.004 | 3.28E-04 | 3.66E-04 | -0.029 | -0.009 | -0.009 | 0.009 | 0 |
| Mean ICVF in cingulum hippocampus (right) | -0.023 | 1.61E-04 | 0.005 | 0.005 | 4.69E-04 | 1.75E-05 | -0.033 | -0.013 | -0.009 | 0.009 | 1 |
| Mean ICVF in corticospinal tract (left) | 0.002 | 0.014 | 0.005 | 0.004 | 2.03E-04 | 0.008 | -0.008 | 0.013 | 0.005 | 0.022 | 0 |
| Mean ICVF in corticospinal tract (right) | -0.003 | 0.013 | 0.005 | 0.005 | 2.48E-04 | 0.0031 | -0.013 | 0.007 | 0.005 | 0.022 | 0 |
| Mean ICVF in external capsule | -0.022 | -0.009 | 0.005 | 0.004 | 3.86E-04 | 2.94E-05 | -0.031 | -0.012 | -0.018 | -0.001 | 1 |

| | | | | | | | | | | | |
|---|---|---|---|---|---|---|---|---|---|---|---|
| (left) | | | | | | | | | | | |
| Mean ICVF in external capsule (right) | -0.025 | -0.010 | 0.005 | 0.004 | 5.08E-04 | 1.06E-06 | -0.034 | -0.015 | -0.019 | -0.003 | 1 |
| Mean ICVF in fornix cres+stria terminalis (left) | -0.017 | -2.00E-04 | 0.005 | 0.004 | 2.58E-04 | 0.001 | -0.027 | -0.007 | -0.008 | 0.008 | 0 |
| Mean ICVF in fornix cres+stria terminalis (right) | -0.018 | 0.004 | 0.005 | 0.004 | 3.69E-04 | 1.10E-04 | -0.028 | -0.009 | -0.005 | 0.013 | 1 |
| Mean ICVF in fornix | -0.018 | -0.009 | 0.005 | 0.005 | 2.71E-04 | 0.003 | -0.029 | -0.007 | -0.018 | 0.000 | 0 |
| Mean ICVF in genu of corpus callosum | -0.032 | -0.011 | 0.005 | 0.004 | 8.00E-04 | 8.38E-10 | -0.041 | -0.022 | -0.020 | -0.003 | 1 |
| Mean ICVF in inferior cerebellar peduncle (left) | -0.015 | 0.004 | 0.005 | 0.004 | 2.69E-04 | 0.001 | -0.025 | -0.005 | -0.004 | 0.013 | 0 |
| Mean ICVF in inferior cerebellar peduncle (right) | -0.014 | 0.003 | 0.005 | 0.004 | 2.13E-04 | 0.005 | -0.024 | -0.004 | -0.006 | 0.011 | 0 |
| Mean ICVF in medial lemniscus (left) | -0.004 | 0.013 | 0.005 | 0.004 | 2.50E-04 | 0.002 | -0.014 | 0.006 | 0.004 | 0.021 | 0 |
| Mean ICVF in medial lemniscus (right) | -0.004 | 0.014 | 0.005 | 0.004 | 2.65E-04 | 0.001 | -0.014 | 0.006 | 0.005 | 0.022 | 0 |
| Mean ICVF in middle cerebellar peduncle | -0.016 | -2.71E-04 | 0.005 | 0.004 | 2.26E-04 | 0.002 | -0.025 | -0.007 | -0.008 | 0.008 | 0 |
| Mean ICVF in pontine crossing tract | 0.002 | 0.010 | 0.005 | 0.004 | 1.13E-04 | 0.064 | -0.008 | 0.012 | 0.002 | 0.019 | 0 |
| Mean ICVF in posterior corona radiata (left) | -0.022 | -0.006 | 0.005 | 0.004 | 3.79E-04 | 6.44E-05 | -0.032 | -0.012 | -0.015 | 0.002 | 1 |
| Mean ICVF in posterior corona radiata (right) | -0.021 | -0.006 | 0.005 | 0.004 | 3.53E-04 | 1.31E-04 | -0.031 | -0.011 | -0.014 | 0.003 | 1 |
| Mean ICVF in posterior limb of internal capsule (left) | -0.026 | -0.006 | 0.005 | 0.004 | 5.24E-04 | 2.28E-07 | -0.035 | -0.017 | -0.013 | 0.002 | 1 |
| Mean ICVF in posterior limb of internal capsule (right) | -0.025 | -0.005 | 0.005 | 0.004 | 5.07E-04 | 4.92E-07 | -0.034 | -0.016 | -0.013 | 0.003 | 1 |
| Mean ICVF in posterior thalamic radiation (left) | -0.016 | -0.005 | 0.005 | 0.004 | 2.11E-04 | 0.004 | -0.026 | -0.007 | -0.014 | 0.003 | 0 |
| Mean ICVF in posterior thalamic radiation (right) | -0.015 | -0.004 | 0.005 | 0.004 | 1.88E-04 | 0.008 | -0.025 | -0.006 | -0.012 | 0.005 | 0 |
| Mean ICVF in retrolenticular part of internal capsule (left) | -0.018 | 2.71E-04 | 0.005 | 0.004 | 2.76E-04 | 7.19E-04 | -0.027 | -0.008 | -0.008 | 0.009 | 0 |
| Mean ICVF in retrolenticular part of internal capsule (right) | -0.016 | 0.002 | 0.005 | 0.004 | 2.48E-04 | 0.002 | -0.026 | -0.006 | -0.007 | 0.010 | 0 |
| Mean ICVF in sagittal stratum (left) | -0.016 | -0.005 | 0.005 | 0.004 | 2.04E-04 | 0.005 | -0.026 | -0.006 | -0.013 | 0.003 | 0 |
| Mean ICVF in sagittal stratum | -0.012 | 0.002 | 0.005 | 0.004 | 1.44E-04 | 0.027 | -0.022 | -0.002 | -0.007 | 0.010 | 0 |

| | | | | | | | | | | | |
|---|---|---|---|---|---|---|---|---|---|---|---|
| (right) | | | | | | | | | | | |
| Mean ICVF in splenium of corpus callosum | -0.021 | -0.005 | 0.005 | 0.004 | 3.47E-04 | 4.46E-05 | -0.030 | -0.012 | -0.013 | 0.003 | 1 |
| Mean ICVF in superior cerebellar peduncle (left) | -0.011 | -0.004 | 0.005 | 0.004 | 9.79E-05 | 0.065 | -0.020 | -0.002 | -0.012 | 0.004 | 0 |
| Mean ICVF in superior cerebellar peduncle (right) | -0.011 | -3.98E-05 | 0.005 | 0.004 | 1.10E-04 | 0.051 | -0.021 | -0.002 | -0.008 | 0.008 | 0 |
| Mean ICVF in superior corona radiata (left) | -0.026 | -0.009 | 0.005 | 0.004 | 5.32E-04 | 3.71E-07 | -0.035 | -0.017 | -0.017 | -0.001 | 1 |
| Mean ICVF in superior corona radiata (right) | -0.023 | -0.009 | 0.005 | 0.004 | 4.44E-04 | 4.43E-06 | -0.033 | -0.014 | -0.018 | -0.001 | 1 |
| Mean ICVF in superior fronto-occipital fasciculus (left) | -0.031 | -0.018 | 0.005 | 0.004 | 8.59E-04 | 1.42E-10 | -0.040 | -0.021 | -0.027 | -0.010 | 1 |
| Mean ICVF in superior fronto-occipital fasciculus (right) | -0.031 | -0.019 | 0.005 | 0.004 | 8.74E-04 | 8.84E-11 | -0.040 | -0.021 | -0.027 | -0.011 | 1 |
| Mean ICVF in superior longitudinal fasciculus (left) | -0.025 | -0.010 | 0.005 | 0.004 | 5.00E-04 | 1.30E-06 | -0.034 | -0.015 | -0.018 | -0.002 | 1 |
| Mean ICVF in superior longitudinal fasciculus (right) | -0.022 | -0.008 | 0.005 | 0.004 | 3.75E-04 | 5.14E-05 | -0.031 | -0.012 | -0.016 | 0.001 | 1 |
| Mean ICVF in tapetum (left) | -0.031 | -0.012 | 0.005 | 0.004 | 7.65E-04 | 8.63E-09 | -0.041 | -0.021 | -0.021 | -0.003 | 1 |
| Mean ICVF in tapetum (right) | -0.025 | -0.012 | 0.005 | 0.004 | 5.27E-04 | 2.88E-06 | -0.035 | -0.015 | -0.020 | -0.003 | 1 |
| Mean ICVF in uncinate fasciculus (left) | -0.021 | -0.008 | 0.005 | 0.004 | 3.64E-04 | 5.73E-05 | -0.031 | -0.012 | -0.017 | 0.000 | 1 |
| Mean ICVF in uncinate fasciculus (right) | -0.024 | -0.012 | 0.005 | 0.004 | 4.86E-04 | 2.63E-06 | -0.034 | -0.015 | -0.020 | -0.003 | 1 |
| Mean ISOVF in anterior corona radiata (left) | 0.0120 | 0.034 | 0.006 | 0.005 | 0.001 | 8.17E-11 | 0.001 | 0.023 | 0.024 | 0.044 | 1 |
| Mean ISOVF in anterior corona radiata (right) | 3.10E-04 | 0.030 | 0.006 | 0.005 | 0.001 | 3.51E-09 | -0.011 | 0.012 | 0.020 | 0.040 | 1 |
| Mean ISOVF in anterior limb of internal capsule (left) | 0.004 | 0.029 | 0.006 | 0.005 | 0.001 | 3.08E-09 | -0.007 | 0.015 | 0.020 | 0.039 | 1 |
| Mean ISOVF in anterior limb of internal capsule (right) | -0.002 | 0.021 | 0.006 | 0.005 | 0.001 | 1.21E-05 | -0.013 | 0.009 | 0.012 | 0.031 | 1 |
| Mean ISOVF in body of corpus callosum | 0.048 | 0.041 | 0.005 | 0.005 | 0.003 | 1.11E-16 | 0.038 | 0.059 | 0.032 | 0.050 | 1 |
| Mean ISOVF in cerebral peduncle (left) | 0.007 | 0.011 | 0.006 | 0.005 | 1.35E-04 | 0.080 | -0.004 | 0.018 | 0.001 | 0.021 | 0 |
| Mean ISOVF in cerebral peduncle (right) | -0.004 | 0.010 | 0.006 | 0.005 | 1.60E-04 | 0.049 | -0.015 | 0.008 | 0.000 | 0.020 | 0 |
| Mean ISOVF in cingulum cingulate gyrus (left) | -0.020 | 0.004 | 0.006 | 0.005 | 4.13E-04 | 5.86E-04 | -0.031 | -0.008 | -0.006 | 0.014 | 0 |

| | | | | | | | | | | | |
|---|---|---|---|---|---|---|---|---|---|---|---|
| Mean ISOVF in cingulum cingulate gyrus (right) | -0.011 | 0.003 | 0.006 | 0.005 | 1.50E-04 | 0.068 | -0.023 | 0.000 | -0.007 | 0.013 | 0 |
| Mean ISOVF in cingulum hippocampus (left) | 0.010 | 0.012 | 0.006 | 0.005 | 1.70E-04 | 0.044 | -0.001 | 0.022 | 0.002 | 0.021 | 0 |
| Mean ISOVF in cingulum hippocampus (right) | -0.007 | 0.007 | 0.006 | 0.005 | 1.45E-04 | 0.075 | -0.019 | 0.005 | -0.003 | 0.018 | 0 |
| Mean ISOVF in corticospinal tract (left) | -0.017 | -4.29E-04 | 0.006 | 0.005 | 2.59E-04 | 0.010 | -0.029 | -0.006 | -0.010 | 0.010 | 0 |
| Mean ISOVF in corticospinal tract (right) | -0.023 | -0.005 | 0.006 | 0.005 | 4.15E-04 | 5.23E-04 | -0.034 | -0.011 | -0.015 | 0.005 | 0 |
| Mean ISOVF in external capsule (left) | 0.020 | 0.031 | 0.006 | 0.005 | 0.001 | 2.18E-09 | 0.008 | 0.031 | 0.021 | 0.041 | 1 |
| Mean ISOVF in external capsule (right) | 0.009 | 0.024 | 0.006 | 0.005 | 6.26E-04 | 6.94E-06 | -0.002 | 0.020 | 0.015 | 0.034 | 1 |
| Mean ISOVF in fornix cres+stria terminalis (left) | 0.036 | 0.023 | 0.006 | 0.005 | 0.001 | 1.59E-11 | 0.025 | 0.047 | 0.014 | 0.032 | 1 |
| Mean ISOVF in fornix cres+stria terminalis (right) | 0.039 | 0.032 | 0.006 | 0.005 | 0.002 | 1.11E-16 | 0.028 | 0.050 | 0.023 | 0.042 | 1 |
| Mean ISOVF in fornix | 0.067 | 0.047 | 0.005 | 0.004 | 0.004 | 1.11E-16 | 0.057 | 0.077 | 0.039 | 0.056 | 1 |
| Mean ISOVF in genu of corpus callosum | 0.054 | 0.050 | 0.005 | 0.005 | 0.004 | 1.11E-16 | 0.043 | 0.064 | 0.041 | 0.059 | 1 |
| Mean ISOVF in inferior cerebellar peduncle (left) | 0.006 | 0.025 | 0.006 | 0.005 | 6.61E-04 | 4.73E-06 | -0.006 | 0.017 | 0.015 | 0.035 | 1 |
| Mean ISOVF in inferior cerebellar peduncle (right) | 0.002 | 0.021 | 0.006 | 0.005 | 0.0004992 | 8.88E-05 | -0.009 | 0.014 | 0.011 | 0.031 | 1 |
| Mean ISOVF in medial lemniscus (left) | -0.009 | 0.003 | 0.006 | 0.005 | 8.69E-05 | 0.201 | -0.020 | 0.003 | -0.007 | 0.013 | 0 |
| Mean ISOVF in medial lemniscus (right) | -0.011 | 0.003 | 0.006 | 0.005 | 0.0001458 | 0.068 | -0.023 | 0.000 | -0.007 | 0.013 | 0 |
| Mean ISOVF in middle cerebellar peduncle | -0.019 | -0.007 | 0.006 | 0.005 | 0.0002981 | 0.002 | -0.030 | -0.008 | -0.016 | 0.003 | 0 |
| Mean ISOVF in pontine crossing tract | -0.020 | -0.014 | 0.006 | 0.005 | 0.0004111 | 4.70E-04 | -0.032 | -0.009 | -0.024 | -0.005 | 0 |
| Mean ISOVF in posterior corona radiata (left) | 0.030 | 0.024 | 0.006 | 0.005 | 0.0010019 | 2.20E-09 | 0.019 | 0.041 | 0.015 | 0.034 | 1 |
| Mean ISOVF in posterior corona radiata (right) | 0.031 | 0.021 | 0.006 | 0.005 | 0.0009290 | 6.87E-09 | 0.020 | 0.042 | 0.011 | 0.030 | 1 |
| Mean ISOVF in posterior limb of internal capsule (left) | -0.019 | 0.013 | 0.006 | 0.005 | 0.0006807 | 1.09E-06 | -0.030 | -0.008 | 0.003 | 0.022 | 1 |
| Mean ISOVF in posterior limb of internal capsule (right) | -0.019 | 0.006 | 0.006 | 0.005 | 4.42E-04 | 1.33E-04 | -0.030 | -0.008 | -0.004 | 0.015 | 1 |

| | | | | | | | | | | | |
|---|---|---|---|---|---|---|---|---|---|---|---|
| Mean ISOVF in posterior thalamic radiation (left) | 0.026 | 0.013 | 0.006 | 0.005 | 5.55E-04 | 1.23E-05 | 0.015 | 0.036 | 0.004 | 0.022 | 1 |
| Mean ISOVF in posterior thalamic radiation (right) | 0.032 | 0.010 | 0.006 | 0.005 | 8.06E-04 | 7.09E-08 | 0.021 | 0.043 | 0.000 | 0.019 | 1 |
| Mean ISOVF in retrolenticular part of internal capsule (left) | -0.006 | 0.004 | 0.006 | 0.005 | 6.46E-05 | 0.307 | -0.017 | 0.006 | -0.006 | 0.014 | 0 |
| Mean ISOVF in retrolenticular part of internal capsule (right) | -0.010 | 0.007 | 0.006 | 0.005 | 1.99E-04 | 0.017 | -0.021 | 0.001 | -0.002 | 0.017 | 0 |
| Mean ISOVF in sagittal stratum (left) | 0.012 | 0.007 | 0.006 | 0.005 | 1.40E-04 | 0.072 | 0.001 | 0.024 | -0.002 | 0.017 | 0 |
| Mean ISOVF in sagittal stratum (right) | 0.016 | 0.011 | 0.006 | 0.005 | 2.50E-04 | 0.006 | 0.005 | 0.027 | 0.002 | 0.021 | 0 |
| Mean ISOVF in splenium of corpus callosum | 0.017 | 0.020 | 0.006 | 0.005 | 4.90E-04 | 7.14E-05 | 0.006 | 0.028 | 0.010 | 0.029 | 1 |
| Mean ISOVF in superior cerebellar peduncle (left) | 0.010 | 0.017 | 0.006 | 0.005 | 3.28E-04 | 0.002 | -0.001 | 0.021 | 0.008 | 0.027 | 0 |
| Mean ISOVF in superior cerebellar peduncle (right) | 0.014 | 0.024 | 0.006 | 0.005 | 6.09E-04 | 7.46E-06 | 0.003 | 0.026 | 0.014 | 0.033 | 1 |
| Mean ISOVF in superior corona radiata (left) | 0.015 | 0.026 | 0.006 | 0.005 | 7.50E-04 | 7.72E-08 | 0.005 | 0.026 | 0.017 | 0.035 | 1 |
| Mean ISOVF in superior corona radiata (right) | 0.001 | 0.020 | 0.005 | 0.005 | 4.47E-04 | 5.48E-05 | -0.001 | 0.021 | 0.011 | 0.030 | 1 |
| Mean ISOVF in superior fronto-occipital fasciculus (left) | 0.021 | 0.029 | 0.006 | 0.005 | 9.59E-04 | 9.47E-09 | 0.010 | 0.032 | 0.019 | 0.038 | 1 |
| Mean ISOVF in superior fronto-occipital fasciculus (right) | 0.012 | 0.017 | 0.006 | 0.005 | 3.35E-04 | 0.002 | 0.001 | 0.024 | 0.007 | 0.027 | 0 |
| Mean ISOVF in superior longitudinal fasciculus (left) | 0.003 | 0.014 | 0.005 | 0.005 | 1.97E-04 | 0.016 | -0.008 | 0.014 | 0.004 | 0.023 | 0 |
| Mean ISOVF in superior longitudinal fasciculus (right) | 0.011 | 0.020 | 0.005 | 0.005 | 4.45E-04 | 5.21E-05 | 0.001 | 0.022 | 0.011 | 0.029 | 1 |
| Mean ISOVF in tapetum (left) | 0.027 | 0.016 | 0.006 | 0.005 | 6.66E-04 | 3.47E-06 | 0.016 | 0.038 | 0.007 | 0.026 | 1 |
| Mean ISOVF in tapetum (right) | 0.029 | 0.016 | 0.006 | 0.005 | 7.51E-04 | 4.49E-07 | 0.018 | 0.040 | 0.007 | 0.026 | 1 |
| Mean ISOVF in uncinate fasciculus (left) | 0.018 | 0.022 | 0.006 | 0.005 | 5.95E-04 | 2.81E-05 | 0.006 | 0.030 | 0.012 | 0.032 | 1 |
| Mean ISOVF in uncinate fasciculus (right) | -0.002 | 0.013 | 0.006 | 0.005 | 2.20E-04 | 0.021 | -0.014 | 0.010 | 0.003 | 0.023 | 0 |
| Mean MD in anterior corona radiata (left) | 0.008 | 0.018 | 0.005 | 0.004 | 3.49E-04 | 4.07E-05 | -0.001 | 0.017 | 0.010 | 0.026 | 1 |
| Mean MD in anterior corona radiata (right) | 0.006 | 0.019 | 0.005 | 0.004 | 3.67E-04 | 2.56E-05 | -0.003 | 0.016 | 0.011 | 0.027 | 1 |
| Mean MD in anterior limb of | -0.004 | 0.019 | 0.005 | 0.004 | 4.75E-04 | 9.87E-07 | -0.013 | 0.005 | 0.011 | 0.026 | 1 |

| | | | | | | | | | | | |
|---|---|---|---|---|---|---|---|---|---|---|---|
| internal capsule (left) | | | | | | | | | | | |
| Mean MD in anterior limb of internal capsule (right) | -0.004 | 0.018 | 0.005 | 0.004 | 4.50E-04 | 3.11E-06 | -0.013 | 0.005 | 0.010 | 0.026 | 1 |
| Mean MD in body of corpus callosum | 0.011 | 0.020 | 0.005 | 0.004 | 4.43E-04 | 6.55E-06 | 0.001 | 0.020 | 0.012 | 0.029 | 1 |
| Mean MD in cerebral peduncle (left) | -0.017 | 0.001 | 0.005 | 0.004 | 2.68E-04 | 6.37E-04 | -0.026 | -0.007 | -0.007 | 0.009 | 0 |
| Mean MD in cerebral peduncle (right) | -0.019 | 0.001 | 0.005 | 0.004 | 3.53E-04 | 5.75E-05 | -0.029 | -0.010 | -0.007 | 0.009 | 1 |
| Mean MD in cingulum cingulate gyrus (left) | -0.009 | 0.009 | 0.005 | 0.004 | 2.16E-04 | 0.001 | -0.018 | 0.000 | 0.001 | 0.017 | 0 |
| Mean MD in cingulum cingulate gyrus (right) | -0.008 | 0.008 | 0.005 | 0.004 | 1.74E-04 | 0.005 | -0.017 | 0.001 | 0.000 | 0.016 | 0 |
| Mean MD in cingulum hippocampus (left) | -0.014 | -2.72E-04 | 0.005 | 0.004 | 1.79E-04 | 0.007 | -0.024 | -0.005 | -0.008 | 0.008 | 0 |
| Mean MD in cingulum hippocampus (right) | -0.012 | -0.0005 | 0.005 | 0.004 | 1.21E-04 | 0.0303 | -0.021 | -0.003 | -0.008 | 0.007 | 0 |
| Mean MD in corticospinal tract (left) | -0.029 | -0.003 | 0.005 | 0.005 | 7.00E-04 | 1.08E-07 | -0.040 | -0.019 | -0.012 | 0.006 | 1 |
| Mean MD in corticospinal tract (right) | -0.031 | -0.007 | 0.005 | 0.005 | 7.50E-04 | 2.71E-08 | -0.041 | -0.021 | -0.016 | 0.002 | 1 |
| Mean MD in external capsule (left) | -0.002 | 0.017 | 0.005 | 0.004 | 3.65E-04 | 2.89E-05 | -0.011 | 0.007 | 0.009 | 0.025 | 1 |
| Mean MD in external capsule (right) | -0.002 | 0.017 | 0.005 | 0.004 | 3.84E-04 | 1.57E-05 | -0.011 | 0.007 | 0.009 | 0.025 | 1 |
| Mean MD in fornix cres+stria terminalis (left) | 0.012 | 0.014 | 0.005 | 0.004 | 2.44E-04 | 0.002 | 0.002 | 0.022 | 0.006 | 0.022 | 0 |
| Mean MD in fornix cres+stria terminalis (right) | 0.015 | 0.019 | 0.005 | 0.004 | 4.52E-04 | 6.99E-06 | 0.006 | 0.025 | 0.011 | 0.028 | 1 |
| Mean MD in fornix | 0.058 | 0.044 | 0.005 | 0.005 | 3.54E-04 | 1.11E-16 | 0.048 | 0.068 | 0.035 | 0.053 | 1 |
| Mean MD in genu of corpus callosum | 0.022 | 0.029 | 0.005 | 0.004 | 0.001 | 1.75E-12 | 0.013 | 0.032 | 0.021 | 0.038 | 1 |
| Mean MD in inferior cerebellar peduncle (left) | -0.008 | 0.010 | 0.005 | 0.004 | 2.30E-04 | 0.002 | -0.018 | 0.001 | 0.002 | 0.018 | 0 |
| Mean MD in inferior cerebellar peduncle (right) | -0.012 | 0.008 | 0.005 | 0.004 | 2.72E-04 | 3.55E-04 | -0.021 | -0.003 | 0.000 | 0.016 | 0 |
| Mean MD in medial lemniscus (left) | -0.021 | -0.003 | 0.005 | 0.004 | 3.61E-04 | 8.36E-05 | -0.031 | -0.011 | -0.011 | 0.006 | 1 |
| Mean MD in medial lemniscus (right) | -0.023 | -0.003 | 0.005 | 0.004 | 4.21E-04 | 1.36E-05 | -0.032 | -0.013 | -0.011 | 0.005 | 1 |
| Mean MD in middle cerebellar | -0.019 | -0.001 | 0.005 | 0.004 | 2.98E-04 | 2.00E-04 | -0.028 | -0.010 | -0.009 | 0.007 | 0 |

| | | | | | | | | | | | |
|---|---|---|---|---|---|---|---|---|---|---|---|
| peduncle | | | | | | | | | | | |
| Mean MD in pontine crossing tract | -0.029 | -0.012 | 0.005 | 0.004 | 6.80E-04 | 1.01E-07 | -0.039 | -0.019 | -0.021 | -0.003 | 1 |
| Mean MD in posterior corona radiata (left) | 0.008 | 0.015 | 0.005 | 0.004 | 2.51E-04 | 0.001 | -0.002 | 0.018 | 0.007 | 0.024 | 0 |
| Mean MD in posterior corona radiata (right) | 0.009 | 0.014 | 0.005 | 0.004 | 2.31E-04 | 0.003 | -0.001 | 0.019 | 0.006 | 0.023 | 0 |
| Mean MD in posterior limb of internal capsule (left) | -0.010 | 0.011 | 0.005 | 0.004 | 2.98E-04 | 1.67E-04 | -0.019 | -0.001 | 0.003 | 0.019 | 0 |
| Mean MD in posterior limb of internal capsule (right) | -0.010 | 0.008 | 0.005 | 0.004 | 2.27E-04 | 0.001 | -0.019 | -0.001 | 0.000 | 0.016 | 0 |
| Mean MD in posterior thalamic radiation (left) | -0.001 | 0.010 | 0.005 | 0.004 | 1.16E-04 | 0.049 | -0.011 | 0.008 | 0.001 | 0.018 | 0 |
| Mean MD in posterior thalamic radiation (right) | 2.78E-04 | 0.007 | 0.005 | 0.004 | 5.39E-05 | 0.240 | -0.009 | 0.010 | -0.001 | 0.015 | 0 |
| Mean MD in retrolenticular part of internal capsule (left) | -0.011 | 0.002 | 0.005 | 0.004 | 1.43E-04 | 0.017 | -0.021 | -0.002 | -0.006 | 0.010 | 0 |
| Mean MD in retrolenticular part of internal capsule (right) | -0.014 | 0.003 | 0.005 | 0.004 | 1.98E-04 | 0.003 | -0.023 | -0.004 | -0.005 | 0.011 | 0 |
| Mean MD in sagittal stratum (left) | -0.010 | 0.005 | 0.005 | 0.004 | 1.38E-04 | 0.025 | -0.019 | 0.000 | -0.003 | 0.013 | 0 |
| Mean MD in sagittal stratum (right) | -0.010 | 0.002 | 0.005 | 0.004 | 1.06E-04 | 0.057 | -0.020 | -0.001 | -0.006 | 0.010 | 0 |
| Mean MD in splenium of corpus callosum | 0.003 | 0.014 | 0.005 | 0.004 | 2.212E-04 | 0.003 | -0.006 | 0.013 | 0.006 | 0.023 | 0 |
| Mean MD in superior cerebellar peduncle (left) | -0.007 | 0.010 | 0.005 | 0.004 | 2.19E-04 | 0.003 | -0.017 | 0.002 | 0.002 | 0.019 | 0 |
| Mean MD in superior cerebellar peduncle (right) | -0.005 | 0.011 | 0.005 | 0.004 | 2.067E-04 | 0.004 | -0.015 | 0.004 | 0.003 | 0.019 | 0 |
| Mean MD in superior corona radiata (left) | 0.004 | 0.017 | 0.005 | 0.004 | 3.01E-04 | 1.85E-04 | -0.005 | 0.013 | 0.009 | 0.025 | 0 |
| Mean MD in superior corona radiata (right) | 3.53E-04 | 0.015 | 0.005 | 0.004 | 2.77E-04 | 3.31E-04 | -0.009 | 0.010 | 0.007 | 0.023 | 0 |
| Mean MD in superior fronto-occipital fasciculus (left) | 0.018 | 0.029 | 0.005 | 0.004 | 9.05E-04 | 2.97E-10 | 0.008 | 0.028 | 0.020 | 0.037 | 1 |
| Mean MD in superior fronto-occipital fasciculus (right) | 0.012 | 0.024 | 0.005 | 0.004 | 5.99E-04 | 2.57E-07 | 0.002 | 0.022 | 0.015 | 0.032 | 1 |
| Mean MD in superior longitudinal fasciculus (left) | -9.78E-04 | 0.013 | 0.005 | 0.004 | 1.95E-04 | 0.004 | -0.010 | 0.008 | 0.005 | 0.020 | 0 |
| Mean MD in superior longitudinal fasciculus (right) | -0.001 | 0.013 | 0.005 | 0.004 | 2.23E-04 | 0.002 | -0.010 | 0.008 | 0.005 | 0.021 | 0 |

| | | | | | | | | | | |
|---|---|---|---|---|---|---|---|---|---|---|
| Mean MD in tapetum (left) | 0.026 | 0.021 | 0.005 | 0.005 | 7.28E-04 | 1.50E-07 | 0.015 | 0.036 | 0.011 | 0.030 | 1 |
| Mean MD in tapetum (right) | 0.024 | 0.020 | 0.005 | 0.005 | 6.39E-04 | 7.47E-07 | 0.013 | 0.034 | 0.010 | 0.029 | 1 |
| Mean MD in uncinate fasciculus (left) | -0.011 | 0.010 | 0.005 | 0.004 | 2.81E-04 | 0.0006 | -0.020 | -0.001 | 0.002 | 0.018 | 0 |
| Mean MD in uncinate fasciculus (right) | -0.010 | 0.011 | 0.005 | 0.004 | 3.04E-04 | 2.62E-04 | -0.020 | -0.001 | 0.003 | 0.019 | 0 |
| Mean OD in anterior corona radiata (left) | 0.011 | 0.018 | 0.005 | 0.005 | 3.37E-04 | 6.14E-04 | 0.000 | 0.021 | 0.008 | 0.027 | 0 |
| Mean OD in anterior corona radiata (right) | 0.005 | 0.013 | 0.005 | 0.005 | 1.79E-04 | 0.019 | -0.006 | 0.015 | 0.004 | 0.022 | 0 |
| Mean OD in anterior limb of internal capsule (left) | 0.010 | 0.009 | 0.005 | 0.005 | 1.23E04 | 0.066 | 0.000 | 0.021 | 0.000 | 0.018 | 0 |
| Mean OD in anterior limb of internal capsule (right) | 0.010 | 0.009 | 0.005 | 0.005 | 1.24E-04 | 0.060 | -0.001 | 0.020 | 0.000 | 0.018 | 0 |
| Mean OD in body of corpus callosum | 0.012 | 0.015 | 0.006 | 0.005 | 2.69E-04 | 0.005 | 0.001 | 0.023 | 0.005 | 0.025 | 0 |
| Mean OD in cerebral peduncle (left) | 0.008 | 0.023 | 0.005 | 0.005 | 5.61E-04 | 4.05E-06 | -0.002 | 0.019 | 0.014 | 0.032 | 1 |
| Mean OD in cerebral peduncle (right) | 0.008 | 0.022 | 0.005 | 0.005 | 5.06E-04 | 1.50E-05 | -0.002 | 0.019 | 0.013 | 0.031 | 1 |
| Mean OD in cingulum cingulate gyrus (left) | 0.022 | 0.023 | 0.006 | 0.005 | 7.31E04 | 3.53E-07 | 0.011 | 0.033 | 0.014 | 0.033 | 1 |
| Mean OD in cingulum cingulate gyrus (right) | 0.030 | 0.029 | 0.006 | 0.005 | 0.001 | 3.65E-11 | 0.019 | 0.041 | 0.020 | 0.039 | 1 |
| Mean OD in cingulum hippocampus (left) | -0.009 | 0.020 | 0.006 | 0.005 | 6.63E-04 | 2.28E-06 | -0.020 | 0.002 | 0.010 | 0.030 | 1 |
| Mean OD in cingulum hippocampus (right) | -0.020 | 0.015 | 0.006 | 0.005 | 8.12E-04 | 1.13E-07 | -0.031 | -0.009 | 0.005 | 0.024 | 1 |
| Mean OD in corticospinal tract (left) | -0.004 | 0.005 | 0.005 | 0.005 | 5.97E-05 | 0.274 | -0.015 | 0.006 | -0.004 | 0.014 | 0 |
| Mean OD in corticospinal tract (right) | -0.007 | 0.007 | 0.005 | 0.005 | 1.44E-04 | 0.041 | -0.017 | 0.003 | -0.002 | 0.017 | 0 |
| Mean OD in external capsule (left) | -0.018 | 0.002 | 0.005 | 0.005 | 2.94E-04 | 0.002 | -0.028 | -0.007 | -0.008 | 0.011 | 0 |
| Mean OD in external capsule (right) | -0.015 | 0.001 | 0.005 | 0.005 | 2.06E-04 | 0.012 | -0.026 | -0.004 | -0.008 | 0.010 | 0 |
| Mean OD in fornix cres+stria terminalis (left) | 0.019 | 0.035 | 0.006 | 0.005 | 0.001 | 2.09E-12 | 0.008 | 0.030 | 0.025 | 0.044 | 1 |
| Mean OD in fornix cres+stria terminalis (right) | 0.014 | 0.030 | 0.006 | 0.005 | 9.65E-04 | 3.07E-09 | 0.003 | 0.025 | 0.021 | 0.040 | 1 |
| Mean OD in fornix | 0.042 | 0.028 | 0.006 | 0.005 | 0.002 | 7.77E-15 | 0.030 | 0.053 | 0.018 | 0.038 | 1 |

| | | | | | | | | | | | |
|---|---|---|---|---|---|---|---|---|---|---|---|
| Mean OD in genu of corpus callosum | -3.72E-04 | 0.010 | 0.005 | 0.005 | 1.29E-04 | 0.069 | -0.011 | 0.010 | 0.001 | 0.020 | 0 |
| Mean OD in inferior cerebellar peduncle (left) | 0.004 | 0.023 | 0.006 | 0.005 | 5.54E-04 | 1.23E-05 | -0.007 | 0.015 | 0.013 | 0.032 | 1 |
| Mean OD in inferior cerebellar peduncle (right) | 0.008 | 0.030 | 0.006 | 0.005 | 9.68E-04 | 2.87E-09 | -0.003 | 0.019 | 0.021 | 0.040 | 1 |
| Mean OD in medial lemniscus (left) | 0.010 | 0.026 | 0.006 | 0.005 | 7.11E-04 | 3.31E-07 | 0.000 | 0.021 | 0.017 | 0.035 | 1 |
| Mean OD in medial lemniscus (right) | 0.004 | 0.026 | 0.006 | 0.005 | 7.27E-04 | 2.81E-07 | -0.007 | 0.015 | 0.016 | 0.035 | 1 |
| Mean OD in middle cerebellar peduncle | -0.014 | 0.011 | 0.005 | 0.004 | 4.35E-04 | 2.19E-05 | -0.024 | -0.004 | 0.003 | 0.020 | 1 |
| Mean OD in pontine crossing tract | 0.002 | 0.011 | 0.006 | 0.005 | 1.29E-04 | 0.096 | -0.010 | 0.013 | 0.001 | 0.021 | 0 |
| Mean OD in posterior corona radiata (left) | -0.024 | -0.002 | 0.005 | 0.005 | 4.68E-04 | 1.81E-05 | -0.034 | -0.013 | -0.010 | 0.007 | 1 |
| Mean OD in posterior corona radiata (right) | -0.030 | -0.006 | 0.005 | 0.005 | 7.03E-04 | 8.38E-08 | -0.040 | -0.020 | -0.015 | 0.002 | 1 |
| Mean OD in posterior limb of internal capsule (left) | -0.024 | -0.010 | 0.005 | 0.005 | 4.73E-04 | 3.48E-05 | -0.035 | -0.014 | -0.019 | -0.001 | 1 |
| Mean OD in posterior limb of internal capsule (right) | -0.026 | -0.009 | 0.005 | 0.005 | 5.42E-04 | 8.52E-06 | -0.037 | -0.015 | -0.019 | 0.000 | 1 |
| Mean OD in posterior thalamic radiation (left) | -0.002 | 0.016 | 0.006 | 0.005 | 3.32E-04 | 0.001 | -0.013 | 0.009 | 0.007 | 0.026 | 0 |
| Mean OD in posterior thalamic radiation (right) | -0.003 | 0.020 | 0.006 | 0.005 | 5.24E-04 | 2.17E-05 | -0.014 | 0.008 | 0.011 | 0.030 | 1 |
| Mean OD in retrolenticular part of internal capsule (left) | -0.015 | 0.011 | 0.006 | 0.005 | 4.46E-04 | 1.09E-04 | -0.026 | -0.004 | 0.002 | 0.021 | 1 |
| Mean OD in retrolenticular part of internal capsule (right) | -0.014 | 0.007 | 0.006 | 0.005 | 3.00E-04 | 0.002 | -0.025 | -0.003 | -0.002 | 0.016 | 0 |
| Mean OD in sagittal stratum (left) | 3.82E-04 | 0.014 | 0.006 | 0.005 | 2.15E-04 | 0.012 | -0.011 | 0.011 | 0.004 | 0.023 | 0 |
| Mean OD in sagittal stratum (right) | 0.002 | 0.020 | 0.006 | 0.005 | 4.63E-04 | 9.70E-05 | -0.009 | 0.013 | 0.011 | 0.030 | 1 |
| Mean OD in splenium of corpus callosum | -0.037 | -0.007 | 0.005 | 0.005 | 0.001 | 4.41E-11 | -0.048 | -0.026 | -0.016 | 0.002 | 1 |
| Mean OD in superior cerebellar peduncle (left) | -0.003 | 0.016 | 0.005 | 0.004 | 3.55E-04 | 2.36E-04 | -0.014 | 0.007 | 0.007 | 0.025 | 0 |
| Mean OD in superior cerebellar peduncle (right) | -0.006 | 0.017 | 0.005 | 0.004 | 4.30E-04 | 3.43E-05 | -0.016 | 0.004 | 0.008 | 0.025 | 1 |
| Mean OD in superior corona radiata (left) | -0.014 | -0.001 | 0.005 | 0.005 | 1.60E-04 | 0.025 | -0.024 | -0.004 | -0.010 | 0.008 | 0 |

| | | | | | | | | | | | |
|---|---|---|---|---|---|---|---|---|---|---|---|
| Mean OD in superior corona radiata (right) | -0.016 | -0.005 | 0.005 | 0.005 | 2.08E-04 | 0.009 | -0.027 | -0.006 | -0.014 | 0.004 | 0 |
| Mean OD in superior fronto-occipital fasciculus (left) | 0.007 | 0.020 | 0.006 | 0.005 | 4.03E-04 | 2.86E-04 | -0.004 | 0.018 | 0.010 | 0.029 | 0 |
| Mean OD in superior fronto-occipital fasciculus (right) | 0.003 | 0.017 | 0.006 | 0.005 | 3.26E-04 | 0.001 | -0.008 | 0.014 | 0.008 | 0.027 | 0 |
| Mean OD in superior longitudinal fasciculus (left) | -0.010 | 0.010 | 0.005 | 0.005 | 2.85E-04 | 0.002 | -0.021 | 0.001 | 0.001 | 0.019 | 0 |
| Mean OD in superior longitudinal fasciculus (right) | -0.011 | 0.012 | 0.005 | 0.005 | 3.77E-04 | 2.27E-04 | -0.022 | -0.001 | 0.003 | 0.021 | 0 |
| Mean OD in tapetum (left) | -0.030 | -0.014 | 0.006 | 0.005 | 7.49E-04 | 7.75E-07 | -0.041 | -0.019 | -0.024 | -0.004 | 1 |
| Mean OD in tapetum (right) | -0.012 | 0.001 | 0.006 | 0.005 | 1.43E-04 | 0.073 | -0.024 | -0.001 | -0.009 | 0.011 | 0 |
| Mean OD in uncinate fasciculus (left) | -0.004 | 0.009 | 0.006 | 0.005 | 1.34E-04 | 0.067 | -0.015 | 0.007 | -0.001 | 0.018 | 0 |
| Mean OD in uncinate fasciculus (right) | -0.008 | 0.010 | 0.006 | 0.005 | 2.13E-04 | 0.013 | -0.019 | 0.003 | 0.000 | 0.019 | 0 |
| Weighted-mean FA in tract acoustic radiation (left) | -0.014 | 0.002 | 0.005 | 0.004 | 1.90E-04 | 0.007 | -0.024 | -0.004 | -0.007 | 0.010 | 0 |
| Weighted-mean FA in tract acoustic radiation (right) | -0.011 | 0.004 | 0.005 | 0.004 | 1.47E-04 | 0.028 | -0.021 | -0.001 | -0.005 | 0.012 | 0 |
| Weighted-mean FA in tract anterior thalamic radiation (left) | -0.025 | -0.010 | 0.005 | 0.004 | 4.96E-04 | 1.70E-06 | -0.034 | -0.015 | -0.018 | -0.001 | 1 |
| Weighted-mean FA in tract anterior thalamic radiation (right) | -0.026 | -0.009 | 0.005 | 0.004 | 5.29E-04 | 8.21E-07 | -0.035 | -0.016 | -0.018 | -0.001 | 1 |
| Weighted-mean FA in tract cingulate gyrus part of cingulum (left) | -0.032 | -0.013 | 0.005 | 0.004 | 8.43E-04 | 1.81E-09 | -0.043 | -0.022 | -0.021 | -0.004 | 1 |
| Weighted-mean FA in tract cingulate gyrus part of cingulum (right) | -0.036 | -0.013 | 0.005 | 0.005 | 0.001 | 1.91E-11 | -0.047 | -0.026 | -0.022 | -0.004 | 1 |
| Weighted-mean FA in tract corticospinal tract (left) | -0.005 | 0.006 | 0.005 | 0.004 | 7.71E-05 | 0.133 | -0.015 | 0.005 | -0.003 | 0.014 | 0 |
| Weighted-mean FA in tract corticospinal tract (right) | -0.002 | 0.006 | 0.005 | 0.004 | 6.39E-04 | 0.198 | -0.012 | 0.008 | -0.002 | 0.015 | 0 |
| Weighted-mean FA in tract forceps major | -0.012 | -8.92E-04 | 0.005 | 0.004 | 1.26E-04 | 0.033 | -0.022 | -0.003 | -0.009 | 0.007 | 0 |
| Weighted-mean FA in tract forceps minor | -0.028 | -0.009 | 0.005 | 0.004 | 6.03E-04 | 5.44E-08 | -0.037 | -0.018 | -0.017 | -0.001 | 1 |
| Weighted-mean FA in tract inferior fronto-occipital | -0.018 | -0.008 | 0.005 | 0.004 | 2.54E-04 | 7.11E-04 | -0.027 | -0.008 | -0.016 | 0.000 | 0 |

| | | | | | | | | | | | |
|---|---|---|---|---|---|---|---|---|---|---|---|
| fasciculus (left) | | | | | | | | | | | |
| Weighted-mean FA in tract inferior fronto-occipital fasciculus (right) | -0.018 | -0.005 | 0.005 | 0.004 | 2.56E-04 | 7.22E-04 | -0.027 | -0.009 | -0.013 | 0.003 | 0 |
| Weighted-mean FA in tract inferior longitudinal fasciculus (left) | -0.020 | -0.006 | 0.005 | 0.004 | 3.12E-04 | 1.10E-04 | -0.029 | -0.011 | -0.014 | 0.002 | 1 |
| Weighted-mean FA in tract inferior longitudinal fasciculus (right) | -0.020 | -0.005 | 0.005 | 0.004 | 2.95E-04 | 2.01E-04 | -0.029 | -0.010 | -0.013 | 0.003 | 0 |
| Weighted-mean FA in tract medial lemniscus (left) | -0.011 | 0.003 | 0.005 | 0.004 | 1.53E-04 | 0.026 | -0.021 | -0.001 | -0.005 | 0.012 | 0 |
| Weighted-mean FA in tract medial lemniscus (right) | -0.007 | 0.002 | 0.005 | 0.004 | 5.44E-05 | 0.269 | -0.017 | 0.003 | -0.007 | 0.010 | 0 |
| Weighted-mean FA in tract middle cerebellar peduncle | -0.006 | 0.006 | 0.005 | 0.004 | 9.26E-05 | 0.113 | -0.016 | 0.004 | -0.003 | 0.014 | 0 |
| Weighted-mean FA in tract parahippocampal part of cingulum (left) | -0.020 | 0.004 | 0.006 | 0.005 | 4.45E-04 | 1.13E-04 | -0.031 | -0.009 | -0.005 | 0.014 | 1 |
| Weighted-mean FA in tract parahippocampal part of cingulum (right) | -0.016 | 0.003 | 0.006 | 0.005 | 2.77E-04 | 0.004 | -0.028 | -0.005 | -0.007 | 0.012 | 0 |
| Weighted-mean FA in tract posterior thalamic radiation (left) | -0.023 | -0.011 | 0.005 | 0.004 | 4.47E-04 | 2.82E-06 | -0.032 | -0.014 | -0.019 | -0.003 | 1 |
| Weighted-mean FA in tract posterior thalamic radiation (right) | -0.023 | -0.010 | 0.005 | 0.004 | 4.45E-04 | 4.81E-06 | -0.033 | -0.014 | -0.018 | -0.002 | 1 |
| Weighted-mean FA in tract superior longitudinal fasciculus (left) | -0.016 | -0.006 | 0.005 | 0.004 | 2.15E-04 | 0.002 | -0.026 | -0.007 | -0.014 | 0.002 | 0 |
| Weighted-mean FA in tract superior longitudinal fasciculus (right) | -0.019 | -0.007 | 0.005 | 0.004 | 2.80E-04 | 4.59E-04 | -0.028 | -0.009 | -0.015 | 0.001 | 0 |
| Weighted-mean FA in tract superior thalamic radiation (left) | -0.010 | 0.001 | 0.005 | 0.004 | 9.49-05 | 0.079 | -0.019 | 0.000 | -0.007 | 0.009 | 0 |
| Weighted-mean FA in tract superior thalamic radiation (right) | -0.007 | 0.003 | 0.005 | 0.004 | 6.43E-05 | 0.184 | -0.017 | 0.003 | -0.006 | 0.011 | 0 |
| Weighted-mean FA in tract uncinate fasciculus (left) | -0.018 | -0.006 | 0.005 | 0.004 | 2.54E-04 | 0.002 | -0.028 | -0.008 | -0.014 | 0.003 | 0 |

| | | | | | | | | | | | |
|---|---|---|---|---|---|---|---|---|---|---|---|
| Weighted-mean FA in tract uncinate fasciculus (right) | -0.023 | -0.008 | 0.005 | 0.004 | 4.11E-04 | 1.98E-05 | -0.032 | -0.013 | -0.016 | 0.001 | 1 |
| Weighted-mean ICVF in tract acoustic radiation (left) | -0.024 | -0.003 | 0.005 | 0.004 | 4.50E-04 | 8.44E-06 | -0.033 | -0.014 | -0.012 | 0.005 | 1 |
| Weighted-mean ICVF in tract acoustic radiation (right) | -0.022 | -0.002 | 0.005 | 0.004 | 4.12E-04 | 3.01E-05 | -0.032 | -0.012 | -0.010 | 0.007 | 1 |
| Weighted-mean ICVF in tract anterior thalamic radiation (left) | -0.026 | -0.007 | 0.005 | 0.004 | 5.31E-04 | 7.46E-07 | -0.036 | -0.016 | -0.015 | 0.001 | 1 |
| Weighted-mean ICVF in tract anterior thalamic radiation (right) | -0.028 | -0.009 | 0.005 | 0.004 | 6.19E-04 | 6.17E-08 | -0.038 | -0.018 | -0.017 | -0.001 | 1 |
| Weighted-mean ICVF in tract cingulate gyrus part of cingulum (left) | -0.029 | -0.011 | 0.005 | 0.004 | 6.50E-04 | 1.42E-07 | -0.039 | -0.018 | -0.020 | -0.002 | 1 |
| Weighted-mean ICVF in tract cingulate gyrus part of cingulum (right) | -0.029 | -0.007 | 0.005 | 0.004 | 6.41E-04 | 1.88E-07 | -0.039 | -0.018 | -0.015 | 0.002 | 1 |
| Weighted-mean ICVF in tract corticospinal tract (left) | -0.018 | 0.001 | 0.005 | 0.004 | 2.89E-04 | 3.48E-04 | -0.027 | -0.008 | -0.007 | 0.009 | 0 |
| Weighted-mean ICVF in tract corticospinal tract (right) | -0.018 | 3.01E-04 | 0.005 | 0.004 | 3.00E-04 | 3.07E-04 | -0.028 | -0.009 | -0.008 | 0.009 | 0 |
| Weighted-mean ICVF in tract forceps major | -0.018 | -0.003 | 0.005 | 0.004 | 2.68E-04 | 7.44E-04 | -0.028 | -0.009 | -0.011 | 0.005 | 0 |
| Weighted-mean ICVF in tract forceps minor | -0.029 | -0.006 | 0.005 | 0.004 | 6.48E-04 | 6.37E-08 | -0.038 | -0.019 | -0.014 | 0.002 | 1 |
| Weighted-mean ICVF in tract inferior fronto-occipital fasciculus (left) | -0.024 | -0.006 | 0.005 | 0.004 | 4.42E-04 | 9.29E-06 | -0.033 | -0.014 | -0.014 | 0.003 | 1 |
| Weighted-mean ICVF in tract inferior fronto-occipital fasciculus (right) | -0.025 | -0.004 | 0.005 | 0.004 | 4.81E-04 | 3.51E-06 | -0.034 | -0.015 | -0.013 | 0.004 | 1 |
| Weighted-mean ICVF in tract inferior longitudinal fasciculus (left) | -0.022 | -0.004 | 0.005 | 0.004 | 3.73E-04 | 4.97E-05 | -0.031 | -0.012 | -0.013 | 0.004 | 1 |
| Weighted-mean ICVF in tract inferior longitudinal fasciculus (right) | -0.022 | -0.003 | 0.005 | 0.004 | 3.74E-04 | 5.41E-05 | -0.031 | -0.012 | -0.012 | 0.005 | 1 |
| Weighted-mean ICVF in tract medial lemniscus (left) | -0.006 | 0.011 | 0.005 | 0.004 | 2.10E-04 | 0.003 | -0.015 | 0.004 | 0.003 | 0.019 | 0 |
| Weighted-mean ICVF in tract medial lemniscus (right) | -0.007 | 0.011 | 0.005 | 0.004 | 2.45E-04 | 0.002 | -0.016 | 0.003 | 0.003 | 0.020 | 0 |
| Weighted-mean ICVF in tract | -0.0155 | 0.004 | 0.005 | 0.004 | 2.72E-04 | 0.001 | -0.025 | -0.006 | -0.004 | 0.013 | 0 |

| | | | | | | | | | | | |
|---|---|---|---|---|---|---|---|---|---|---|---|
| middle cerebellar peduncle | | | | | | | | | | | |
| Weighted-mean ICVF in tract parahippocampal part of cingulum (left) | -0.017 | 0.004 | 0.005 | 0.004 | 3.34E-04 | 3.47E-04 | -0.027 | -0.007 | -0.005 | 0.013 | 0 |
| Weighted-mean ICVF in tract parahippocampal part of cingulum (right) | -0.017 | 0.003 | 0.005 | 0.005 | 2.99E-04 | 0.001 | -0.027 | -0.007 | -0.006 | 0.012 | 0 |
| Weighted-mean ICVF in tract posterior thalamic radiation (left) | -0.022 | -0.006 | 0.005 | 0.004 | 3.82E-04 | 3.22E-05 | -0.031 | -0.013 | -0.014 | 0.002 | 1 |
| Weighted-mean ICVF in tract posterior thalamic radiation (right) | -0.022 | -0.006 | 0.005 | 0.004 | 3.88E-04 | 3.37E-05 | -0.032 | -0.013 | -0.015 | 0.002 | 1 |
| Weighted-mean ICVF in tract superior longitudinal fasciculus (left) | -0.025 | -0.008 | 0.005 | 0.004 | 5.03E-04 | 1.30E-06 | -0.035 | -0.016 | -0.016 | 0.001 | 1 |
| Weighted-mean ICVF in tract superior longitudinal fasciculus (right) | -0.024 | -0.008 | 0.005 | 0.004 | 4.73E-04 | 3.62E-06 | -0.034 | -0.015 | -0.016 | 0.001 | 1 |
| Weighted-mean ICVF in tract superior thalamic radiation (left) | -0.027 | -0.007 | 0.005 | 0.004 | 5.59E-04 | 1.33E-07 | -0.036 | -0.017 | -0.015 | 0.001 | 1 |
| Weighted-mean ICVF in tract superior thalamic radiation (right) | -0.026 | -0.009 | 0.005 | 0.004 | 5.15E-04 | 5.19E-07 | -0.035 | -0.016 | -0.017 | -0.001 | 1 |
| Weighted-mean ICVF in tract uncinate fasciculus (left) | -0.025 | -0.007 | 0.005 | 0.004 | 4.78E-04 | 4.31E-06 | -0.034 | -0.015 | -0.016 | 0.001 | 1 |
| Weighted-mean ICVF in tract uncinate fasciculus (right) | -0.028 | -0.009 | 0.005 | 0.004 | 6.38E-04 | 5.91E-08 | -0.038 | -0.019 | -0.017 | -0.001 | 1 |
| Weighted-mean ISOVF in tract acoustic radiation (left) | -0.014 | 0.003 | 0.008 | 0.005 | 2.26E-04 | 0.016 | -0.026 | -0.003 | -0.007 | 0.013 | 0 |
| Weighted-mean ISOVF in tract acoustic radiation (right) | -0.008 | -0.001 | 0.006 | 0.005 | 5.39E-05 | 0.350 | -0.019 | 0.003 | -0.011 | 0.008 | 0 |
| Weighted-mean ISOVF in tract anterior thalamic radiation (left) | 0.021 | 0.028 | 0.006 | 0.005 | 9.33E-04 | 5.24E-09 | 0.010 | 0.032 | 0.019 | 0.038 | 1 |
| Weighted-mean ISOVF in tract anterior thalamic radiation (right) | 0.015 | 0.030 | 0.006 | 0.005 | 9.80E-04 | 2.12E-09 | 0.004 | 0.026 | 0.021 | 0.040 | 1 |
| Weighted-mean ISOVF in tract cingulate gyrus part of cingulum (left) | -0.016 | 0.002 | 0.006 | 0.005 | 2.40E-04 | 0.014 | -0.027 | -0.004 | -0.008 | 0.012 | 0 |
| Weighted-mean ISOVF in tract cingulate gyrus part of cingulum | -0.015 | 0.009 | 0.006 | 0.005 | 4.01E-04 | 0.001 | -0.027 | -0.004 | -0.001 | 0.019 | 0 |

| | | | | | | | | | | |
|---|---|---|---|---|---|---|---|---|---|---|
| (right) | | | | | | | | | | |
| Weighted-mean ISOVF in tract corticospinal tract (left) | 0.006 | 0.013 | 0.006 | 0.005 | 1.84E-04 | 0.024 | -0.010 | 0.012 | 0.003 | 0.022 | 0 |
| Weighted-mean ISOVF in tract corticospinal tract (right) | -0.005 | 0.011 | 0.006 | 0.005 | 1.97E-04 | 0.017 | -0.015 | 0.006 | 0.002 | 0.020 | 0 |
| Weighted-mean ISOVF in tract forceps major | 0.003 | 0.002 | 0.006 | 0.005 | 1.18E-05 | 0.793 | -0.008 | 0.015 | -0.007 | 0.012 | 0 |
| Weighted-mean ISOVF in tract forceps minor | -0.003 | 0.014 | 0.006 | 0.005 | 2.77E-04 | 0.005 | -0.014 | 0.009 | 0.005 | 0.024 | 0 |
| Weighted-mean ISOVF in tract inferior fronto-occipital fasciculus (left) | 0.017 | 0.017 | 0.006 | 0.005 | 4.20E-04 | 2.50E-04 | 0.006 | 0.028 | 0.008 | 0.027 | 0 |
| Weighted-mean ISOVF in tract inferior fronto-occipital fasciculus (right) | 0.019 | 0.023 | 0.006 | 0.005 | 6.27E-04 | 2.44E-06 | 0.008 | 0.029 | 0.013 | 0.032 | 1 |
| Weighted-mean ISOVF in tract inferior longitudinal fasciculus (left) | 0.027 | 0.013 | 0.006 | 0.005 | 6.16E-04 | 4.06E-06 | 0.016 | 0.038 | 0.003 | 0.022 | 1 |
| Weighted-mean ISOVF in tract inferior longitudinal fasciculus (right) | 0.033 | 0.018 | 0.005 | 0.005 | 9.37E-04 | 2.06E-09 | 0.022 | 0.044 | 0.009 | 0.027 | 1 |
| Weighted-mean ISOVF in tract medial lemniscus (left) | 0.009 | 0.007 | 0.006 | 0.005 | 8.85E-05 | 0.162 | -0.002 | 0.020 | -0.002 | 0.016 | 0 |
| Weighted-mean ISOVF in tract medial lemniscus (right) | -0.001 | 0.002 | 0.005 | 0.005 | 9.60E-05 | 0.812 | -0.012 | 0.009 | -0.007 | 0.011 | 0 |
| Weighted-mean ISOVF in tract middle cerebellar peduncle | 0.028 | 0.033 | 0.006 | 0.005 | 0.001 | 2.13E-11 | 0.016 | 0.040 | 0.023 | 0.043 | 1 |
| Weighted-mean ISOVF in tract parahippocampal part of cingulum (left) | 0.017 | -0.001 | 0.006 | 0.005 | 2.77E-04 | 0.006 | 0.006 | 0.029 | -0.011 | 0.009 | 0 |
| Weighted-mean ISOVF in tract parahippocampal part of cingulum (right) | 0.017 | 0.007 | 0.006 | 0.005 | 2.44E-04 | 0.010 | 0.006 | 0.029 | -0.002 | 0.017 | 0 |
| Weighted-mean ISOVF in tract posterior thalamic radiation (left) | 0.039 | 0.023 | 0.006 | 0.005 | 0.001 | 9.08E-13 | 0.028 | 0.050 | 0.014 | 0.033 | 1 |
| Weighted-mean ISOVF in tract posterior thalamic radiation (right) | 0.040 | 0.027 | 0.006 | 0.005 | 0.002 | 1.09E-14 | 0.029 | 0.051 | 0.017 | 0.036 | 1 |
| Weighted-mean ISOVF in tract superior longitudinal fasciculus (left) | 0.015 | 0.023 | 0.006 | 0.005 | 5.82E-04 | 5.00E-06 | 0.005 | 0.026 | 0.013 | 0.032 | 1 |

| | | | | | | | | | | | |
|---|---|---|---|---|---|---|---|---|---|---|---|
| Weighted-mean ISOVF in tract superior longitudinal fasciculus (right) | 0.018 | 0.027 | 0.005 | 0.005 | 8.11E-04 | 2.17E-08 | 0.007 | 0.028 | 0.018 | 0.036 | 1 |
| Weighted-mean ISOVF in tract superior thalamic radiation (left) | 0.016 | 0.024 | 0.005 | 0.005 | 6.68E-04 | 2.62E-07 | 0.006 | 0.027 | 0.015 | 0.033 | 1 |
| Weighted-mean ISOVF in tract superior thalamic radiation (right) | 0.012 | 0.022 | 0.005 | 0.005 | 5.256E-04 | 7.72E-06 | 0.001 | 0.022 | 0.013 | 0.031 | 1 |
| Weighted-mean ISOVF in tract uncinate fasciculus (left) | 0.006 | 0.015 | 0.006 | 0.005 | 2.34E-04 | 0.013 | -0.005 | 0.018 | 0.005 | 0.025 | 0 |
| Weighted-mean ISOVF in tract uncinate fasciculus (right) | 0.008 | 0.019 | 0.006 | 0.005 | 3.65E-04 | 0.001 | -0.004 | 0.019 | 0.009 | 0.028 | 0 |
| Weighted-mean MD in tract acoustic radiation (left) | -0.013 | 0.004 | 0.005 | 0.004 | 2.10E-04 | 0.003 | -0.022 | -0.004 | -0.004 | 0.012 | 0 |
| Weighted-mean MD in tract acoustic radiation (right) | -0.010 | 0.001 | 0.005 | 0.004 | 9.95E-05 | 0.057 | -0.019 | -0.001 | -0.007 | 0.009 | 0 |
| Weighted-mean MD in tract anterior thalamic radiation (left) | 0.002 | 0.015 | 0.005 | 0.004 | 2.52E-04 | 0.001 | -0.007 | 0.011 | 0.007 | 0.023 | 0 |
| Weighted-mean MD in tract anterior thalamic radiation (right) | 0.003 | 0.017 | 0.005 | 0.004 | 3.212E-04 | 9.04E-05 | -0.006 | 0.012 | 0.009 | 0.025 | 1 |
| Weighted-mean MD in tract cingulate gyrus part of cingulum (left) | -0.009 | 0.007 | 0.005 | 0.004 | 1.78E-04 | 0.005 | -0.018 | 0.000 | -0.001 | 0.015 | 0 |
| Weighted-mean MD in tract cingulate gyrus part of cingulum (right) | -0.009 | 0.008 | 0.005 | 0.004 | 1.96E-04 | 0.003 | -0.018 | 0.000 | 0.000 | 0.016 | 0 |
| Weighted-mean MD in tract corticospinal tract (left) | -0.008 | 0.007 | 0.005 | 0.004 | 1.46E-04 | 0.017 | -0.017 | 0.002 | -0.001 | 0.015 | 0 |
| Weighted-mean MD in tract corticospinal tract (right) | -0.009 | 0.007 | 0.005 | 0.004 | 1.65E-04 | 0.009 | -0.018 | 0.001 | -0.001 | 0.015 | 0 |
| Weighted-mean MD in tract forceps major | -0.004 | 0.005 | 0.005 | 0.004 | 4.90E-05 | 0.288 | -0.014 | 0.006 | -0.004 | 0.013 | 0 |
| Weighted-mean MD in tract forceps minor | -0.001 | 0.009 | 0.005 | 0.004 | 1.09E-04 | 0.046 | -0.010 | 0.009 | 0.001 | 0.017 | 0 |
| Weighted-mean MD in tract inferior fronto-occipital fasciculus (left) | -0.003 | 0.008 | 0.005 | 0.004 | 9.86E-05 | 0.061 | -0.012 | 0.006 | 0.000 | 0.016 | 0 |
| Weighted-mean MD in tract inferior fronto-occipital fasciculus (right) | 3.93E-06 | 0.012 | 0.005 | 0.004 | 1.58E-04 | 0.0117 | -0.009 | 0.009 | 0.004 | 0.020 | 0 |
| Weighted-mean MD in tract | 2.21E-04 | 0.008 | 0.005 | 0.004 | 7.77E-05 | 0.112 | -0.009 | 0.010 | 0.000 | 0.016 | 0 |

| | | | | | | | | | | |
|---|---|---|---|---|---|---|---|---|---|---|
| inferior longitudinal fasciculus (left) | | | | | | | | | | |
| Weighted-mean MD in tract inferior longitudinal fasciculus (right) | 0.002 | 0.010 | 0.005 | 0.004 | 1.02E-04 | 0.054 | -0.007 | 0.012 | 0.002 | 0.018 | 0 |
| Weighted-mean MD in tract medial lemniscus (left) | -0.007 | 0.002 | 0.005 | 0.004 | 5.56E-05 | 0.220 | -0.016 | 0.003 | -0.006 | 0.010 | 0 |
| Weighted-mean MD in tract medial lemniscus (right) | -0.012 | -4.02E-04 | 0.005 | 0.004 | 1.32E-04 | 0.025 | -0.022 | -0.003 | -0.008 | 0.008 | 0 |
| Weighted-mean MD in tract middle cerebellar peduncle | 0.012 | 0.023 | 0.005 | 0.005 | 5.67E-04 | 3.29E-06 | 0.001 | 0.022 | 0.014 | 0.032 | 1 |
| Weighted-mean MD in tract parahippocampal part of cingulum (left) | 0.001 | -0.002 | 0.005 | 0.005 | 3.81E-06 | 0.916 | -0.010 | 0.011 | -0.010 | 0.007 | 0 |
| Weighted-mean MD in tract parahippocampal part of cingulum (right) | 0.002 | 0.005 | 0.005 | 0.004 | 3.08E-04 | 0.485 | -0.008 | 0.012 | -0.003 | 0.014 | 0 |
| Weighted-mean MD in tract posterior thalamic radiation (left) | 0.008 | 0.015 | 0.005 | 0.004 | 2.34E-04 | 0.002 | -0.001 | 0.018 | 0.006 | 0.023 | 0 |
| Weighted-mean MD in tract posterior thalamic radiation (right) | 0.009 | 0.016 | 0.005 | 0.004 | 2.72E-04 | 6.47E-04 | -0.001 | 0.018 | 0.008 | 0.024 | 0 |
| Weighted-mean MD in tract superior longitudinal fasciculus (left) | 0.002 | 0.014 | 0.005 | 0.004 | 2.22E-04 | 0.001 | -0.007 | 0.011 | 0.006 | 0.022 | 0 |
| Weighted-mean MD in tract superior longitudinal fasciculus (right) | 0.002 | 0.016 | 0.005 | 0.004 | 2.71E-04 | 3.44E-04 | -0.007 | 0.011 | 0.008 | 0.024 | 0 |
| Weighted-mean MD in tract superior thalamic radiation (left) | 0.004 | 0.014 | 0.005 | 0.004 | 2.19E-04 | 0.001 | -0.005 | 0.013 | 0.007 | 0.022 | 0 |
| Weighted-mean MD in tract superior thalamic radiation (right) | 0.002 | 0.015 | 0.005 | 0.004 | 2.69E-04 | 3.12E-04 | -0.008 | 0.011 | 0.008 | 0.023 | 0 |
| Weighted-mean MD in tract uncinate fasciculus (left) | -0.004 | 0.011 | 0.005 | 0.004 | 1.72E-04 | 0.009 | -0.013 | 0.006 | 0.003 | 0.019 | 0 |
| Weighted-mean MD in tract uncinate fasciculus (right) | -0.001 | 0.013 | 0.005 | 0.004 | 2.18E-04 | 0.002 | -0.010 | 0.009 | 0.005 | 0.021 | 0 |
| Weighted-mean OD in tract acoustic radiation (left) | -0.018 | -0.006 | 0.005 | 0.004 | 2.44E-04 | 0.003 | -0.028 | -0.007 | -0.015 | 0.002 | 0 |
| Weighted-mean OD in tract acoustic radiation (right) | -0.020 | -0.007 | 0.005 | 0.004 | 3.19E-04 | 2.60E-04 | -0.030 | -0.010 | -0.015 | 0.001 | 0 |

| | | | | | | | | | | | |
|---|---|---|---|---|---|---|---|---|---|---|---|
| Weighted-mean OD in tract anterior thalamic radiation (left) | -0.010 | 0.003 | 0.005 | 0.004 | 1.16E-04 | 0.042 | -0.019 | 0.000 | -0.005 | 0.011 | 0 |
| Weighted-mean OD in tract anterior thalamic radiation (right) | -0.011 | -2.81E-04 | 0.005 | 0.004 | 1.12E-04 | 0.044 | -0.021 | -0.002 | -0.008 | 0.008 | 0 |
| Weighted-mean OD in tract cingulate gyrus part of cingulum (left) | 0.006 | 0.011 | 0.005 | 0.005 | 1.27E-04 | 0.066 | -0.005 | 0.016 | 0.002 | 0.020 | 0 |
| Weighted-mean OD in tract cingulate gyrus part of cingulum (right) | 0.012 | 0.015 | 0.006 | 0.005 | 2.71E-04 | 0.003 | 0.001 | 0.023 | 0.006 | 0.024 | 0 |
| Weighted-mean OD in tract corticospinal tract (left) | -0.033 | -0.015 | 0.005 | 0.004 | 8.94E-04 | 6.22E-11 | -0.043 | -0.023 | -0.023 | -0.007 | 1 |
| Weighted-mean OD in tract corticospinal tract (right) | -0.036 | -0.012 | 0.005 | 0.004 | 0.001 | 3.43E-12 | -0.045 | -0.026 | -0.020 | -0.004 | 1 |
| Weighted-mean OD in tract forceps major | -0.027 | -0.007 | 0.005 | 0.004 | 0.001 | 3.26E-07 | -0.037 | -0.017 | -0.016 | 0.001 | 1 |
| Weighted-mean OD in tract forceps minor | -0.014 | -0.001 | 0.005 | 0.004 | 1.70E-04 | 0.010 | -0.024 | -0.005 | -0.009 | 0.007 | 0 |
| Weighted-mean OD in tract inferior fronto-occipital fasciculus (left) | -0.022 | 0.003 | 0.005 | 0.004 | 0.001 | 4.05E-06 | -0.032 | -0.012 | -0.005 | 0.012 | 1 |
| Weighted-mean OD in tract inferior fronto-occipital fasciculus (right) | -0.024 | 0.001 | 0.005 | 0.004 | 0.001 | 2.28E-06 | -0.033 | -0.014 | -0.007 | 0.010 | 1 |
| Weighted-mean OD in tract inferior longitudinal fasciculus (left) | -0.017 | 0.004 | 0.005 | 0.004 | 3.06E-04 | 3.607E-04 | -0.026 | -0.007 | -0.004 | 0.012 | 0 |
| Weighted-mean OD in tract inferior longitudinal fasciculus (right) | -0.018 | 0.004 | 0.005 | 0.004 | 3.61E-04 | 5.68E-05 | -0.028 | -0.009 | -0.004 | 0.012 | 1 |
| Weighted-mean OD in tract medial lemniscus (left) | 0.003 | 0.010 | 0.005 | 0.004 | 9.69E-05 | 0.063 | -0.006 | 0.013 | 0.002 | 0.018 | 0 |
| Weighted-mean OD in tract medial lemniscus (right) | -0.005 | 0.008 | 0.005 | 0.004 | 1.29E-04 | 0.027 | -0.014 | 0.005 | 0.000 | 0.017 | 0 |
| Weighted-mean OD in tract middle cerebellar peduncle | -0.025 | -0.011 | 0.005 | 0.004 | 5.27E-04 | 2.91E-06 | -0.035 | -0.015 | -0.020 | -0.003 | 1 |
| Weighted-mean OD in tract parahippocampal part of cingulum (left) | -0.008 | 0.004 | 0.005 | 0.004 | 8.47E-05 | 0.132 | -0.018 | 0.002 | -0.005 | 0.012 | 0 |
| Weighted-mean OD in tract parahippocampal part of | -0.012 | 0.002 | 0.005 | 0.004 | 1.57E-04 | 0.022 | -0.022 | -0.002 | -0.007 | 0.010 | 0 |

| | | | | | | | | | | | |
|---|---|---|---|---|---|---|---|---|---|---|---|
| cingulum (right) | | | | | | | | | | | |
| Weighted-mean OD in tract posterior thalamic radiation (left) | -0.013 | 0.005 | 0.005 | 0.004 | 2.36E-04 | 0.002 | -0.023 | -0.004 | -0.003 | 0.014 | 0 |
| Weighted-mean OD in tract posterior thalamic radiation (right) | -0.016 | 0.004 | 0.005 | 0.004 | 2.67E-04 | 0.001 | -0.025 | -0.006 | -0.005 | 0.012 | 0 |
| Weighted-mean OD in tract superior longitudinal fasciculus (left) | -0.029 | -0.003 | 0.005 | 0.004 | 7.09E-04 | 6.79E-09 | -0.039 | -0.020 | -0.011 | 0.006 | 1 |
| Weighted-mean OD in tract superior longitudinal fasciculus (right) | -0.025 | -0.003 | 0.005 | 0.004 | 5.16E-04 | 8.33E-07 | -0.035 | -0.016 | -0.012 | 0.005 | 1 |
| Weighted-mean OD in tract superior thalamic radiation (left) | -0.028 | -0.010 | 0.005 | 0.004 | 6.13E-04 | 2.35E-08 | -0.037 | -0.019 | -0.018 | -0.002 | 1 |
| Weighted-mean OD in tract superior thalamic radiation (right) | -0.031 | -0.011 | 0.005 | 0.004 | 7.73E-04 | 2.80E-10 | -0.040 | -0.022 | -0.019 | -0.003 | 1 |
| Weighted-mean OD in tract uncinate fasciculus (left) | -0.013 | 0.001 | 0.005 | 0.004 | 1.65E-04 | 0.015 | -0.023 | -0.004 | -0.008 | 0.009 | 0 |
| Weighted-mean OD in tract uncinate fasciculus (right) | -0.012 | 0.002 | 0.005 | 0.004 | 1.57E-04 | 0.018 | -0.022 | -0.003 | -0.006 | 0.010 | 0 |

*Note*. Model: IDP: white matter IDP, normalized for head size, x: standardized log(1 + daily units)

IDP = b0 + b1 x + b2 x^2+ bc controls + error term

List of control variables: standardized age, squared standardized age, genetic principal components 1 to 40, standardized height, handedness, sex (female:0, male:1), standardized townsend, county of residence, current smoker, former light smoker, former heavy smoker

FA = fractional anisotropy, MD = mean diffusivity, ICVF = intracellular volume fraction, ISOVF = isotropic volume fraction, OD = orientation dispersion, beta_lin = regression coefficient b1, for standardized log(1 + daily units), beta_quad = regression coefficient b2, for squared standardized log(1 + daily units), beta_lin (se) = standard error for b1, beta_quad (se)= standard error for b2, delta_r2 = difference in r-squared values for the regressions with and without intake and squared intake, p-val = p-value of the F-test of joint significance for b1 and b2, c95_lin (low) = 95% confidence interval for b1 (lower bound), c95_lin (high) = 95% confidence interval for b1 (upper bound), c95_quad (low) = 95% confidence interval for b2 (lower bound), c95_quad (high) = 95% confidence interval for b2 (upper bound), Sig = family-wise error rate (FWER) corrected significance (Holm method).
