## Extended Data Figure 1 for "Multimodal brain imaging study of 36,678 participants reveals adverse effects of moderate drinking"

**Extended Data Figure 1.** Marginal effect between alcohol intake and regional gray matter volume imaging-derived phenotypes predicted from the regression model with standard controls, arranged into brain lobes along the x-axis. For these variables, the significance of the correlation is plotted vertically, in units of $-\log_{10}$(p-value). The dotted horizontal line indicates the threshold corresponding to multiple correction using the Holm method ($1.65 \times 10^{-4}$). The panels show the effect of increasing from zero to one unit (top), from one to two units (second), from two to three units (third) and from three to four units (bottom).

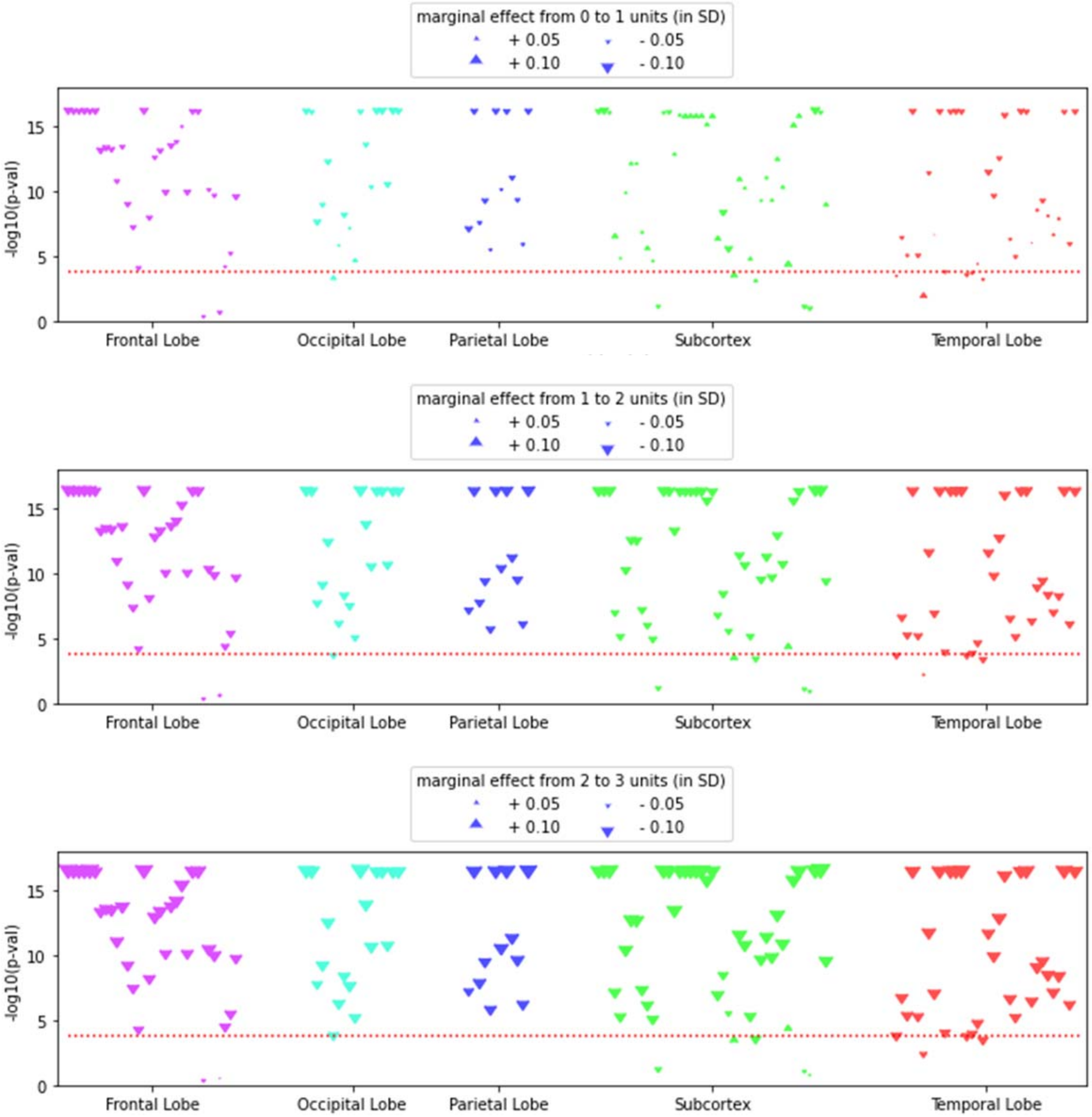

11

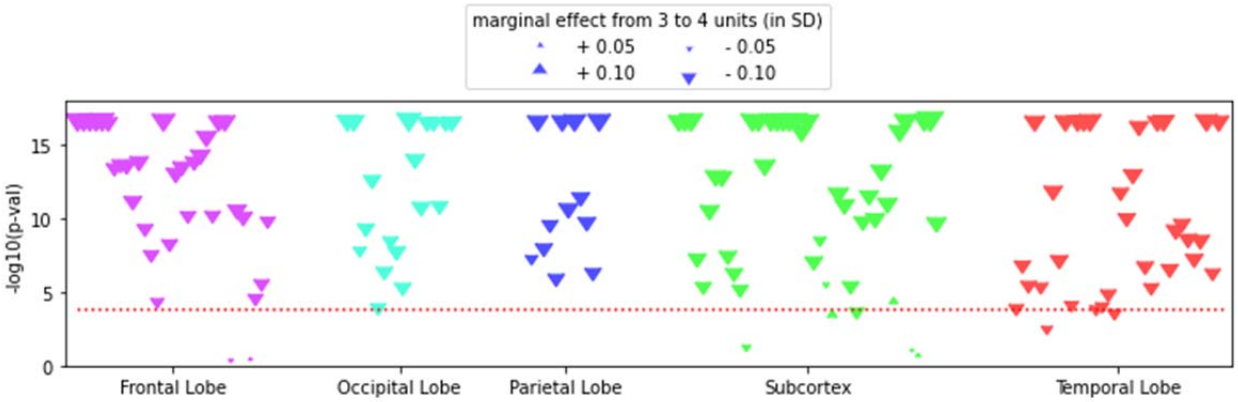

marginal effect from 3 to 4 units (in SD)
+ 0.05
+ 0.10
- 0.05
- 0.10
-log10(p-val)
15
10
5
0
Frontal Lobe
Occipital Lobe
Parietal Lobe
Subcortex
Temporal Lobe
