## Extended Data Figure 2 for "Multimodal brain imaging study of 36,678 participants reveals adverse effects of moderate drinking"

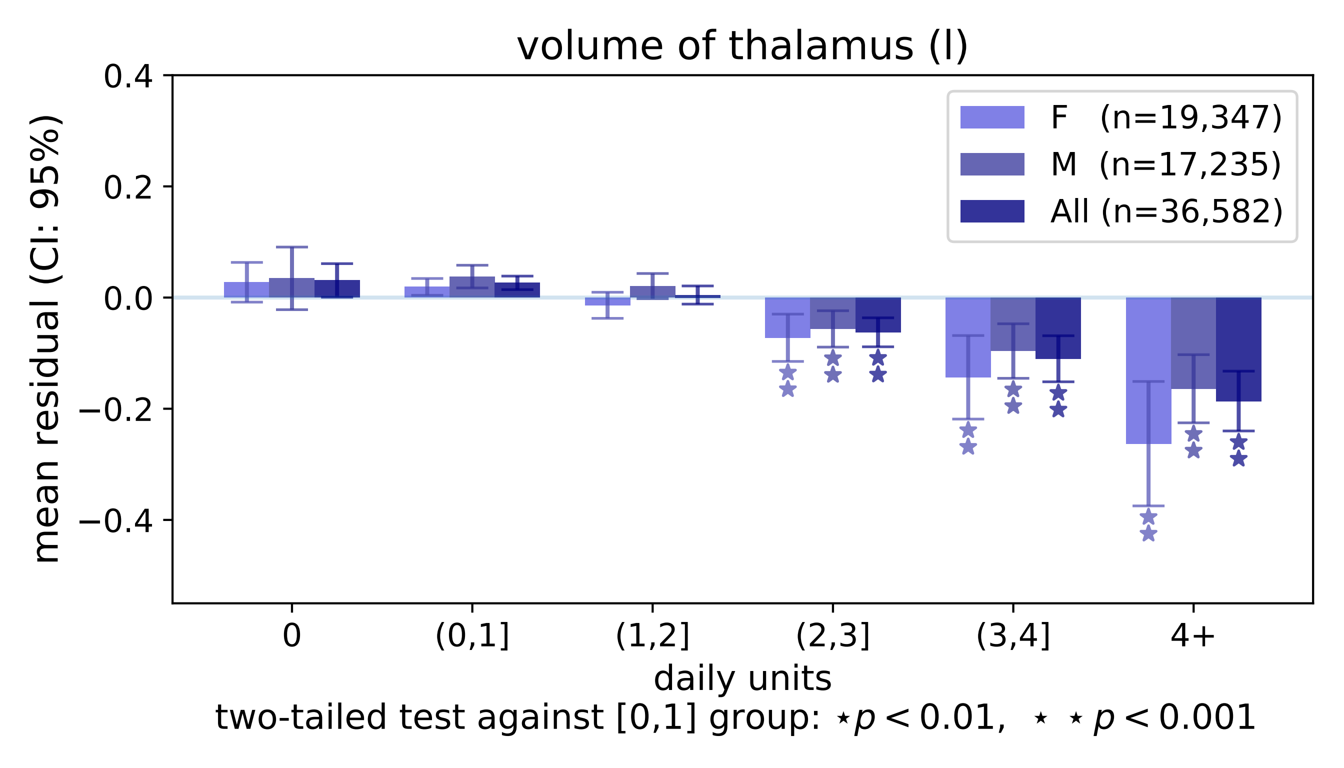

volume of thalamus (l)
mean residual (CI: 95%)
0.4
0.2
0.0
−0.2
−0.4
F (n=19,347)
M (n=17,235)
All (n=36,582)
0
(0,1]
(1,2]
(2,3]
(3,4]
4+
daily units
two-tailed test against [0,1] group: ⋆ p < 0.01, ⋆ ⋆ p < 0.001

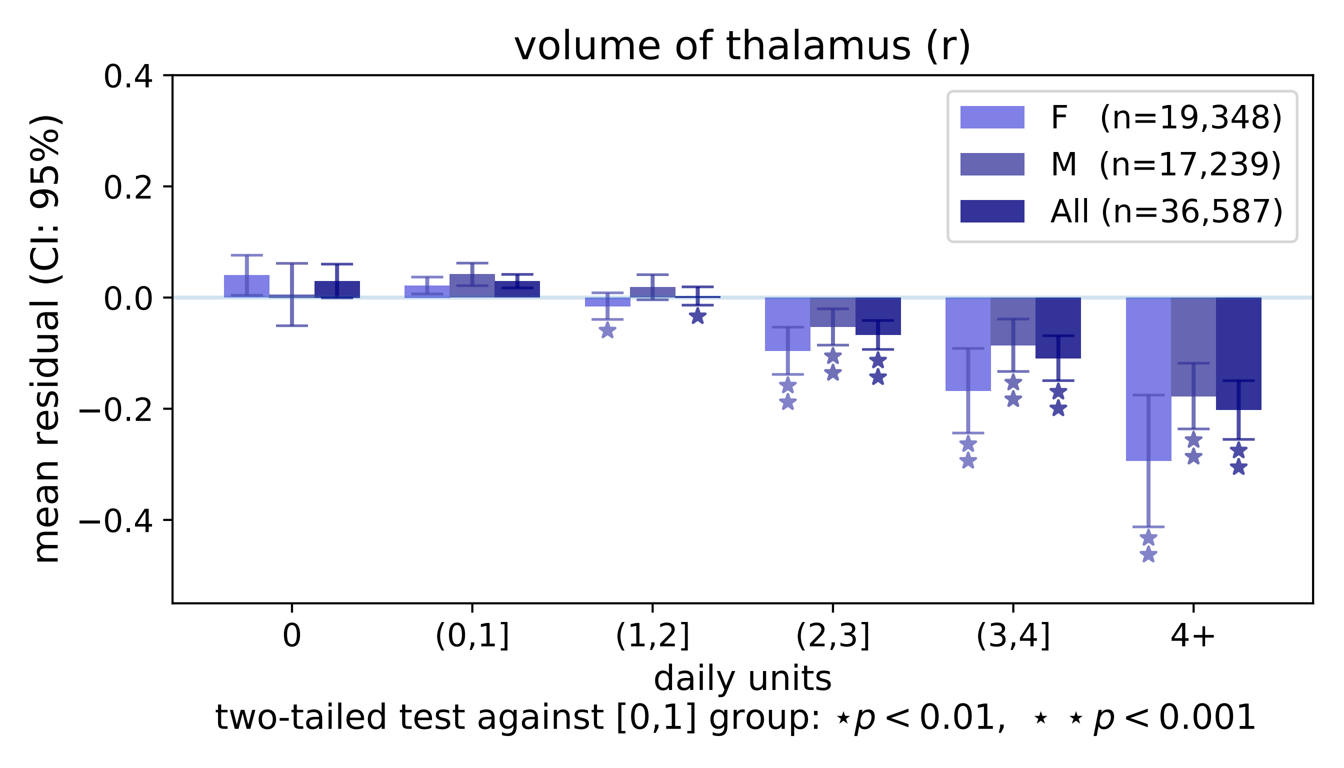
volume of thalamus (r)
mean residual (CI: 95%)
0.4
0.2
0.0
−0.2
−0.4
F (n=19,348)
M (n=17,239)
All (n=36,587)
0
(0,1]
(1,2]
(2,3]
(3,4]
4+
daily units
two-tailed test against [0,1] group: ⋆p < 0.01, ⋆ ⋆ p < 0.001

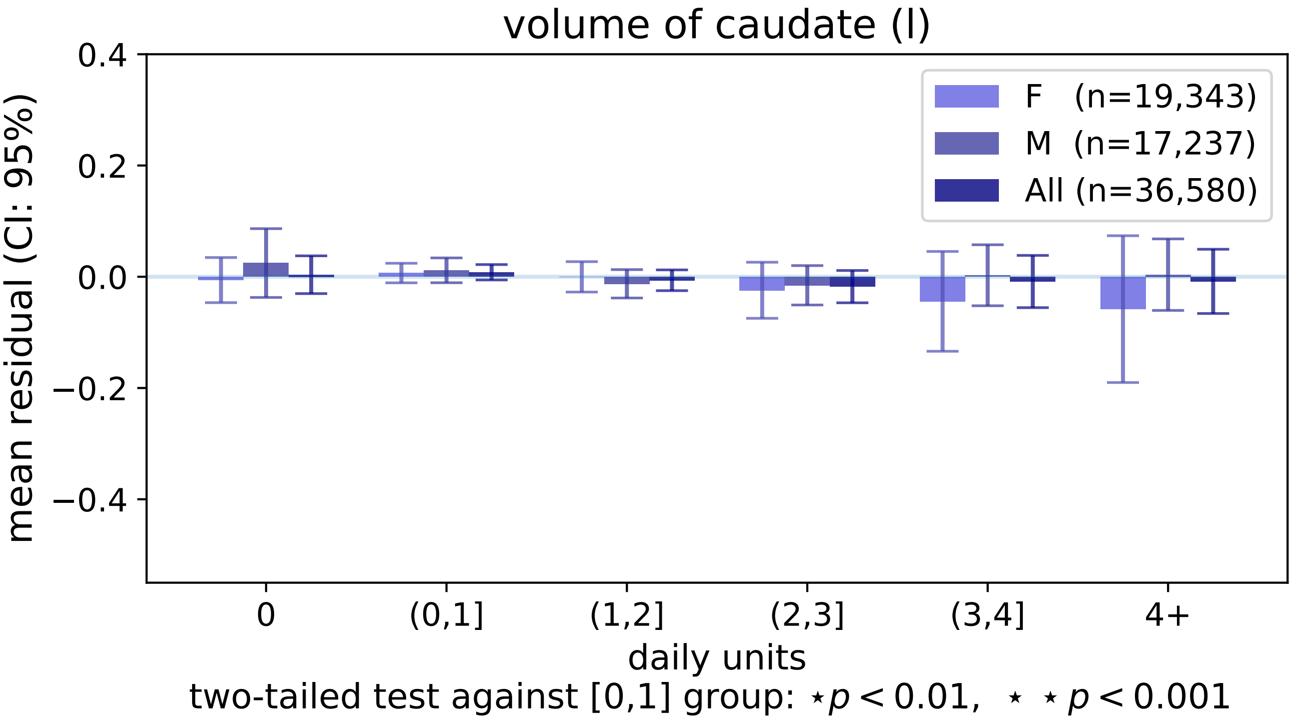
volume of caudate (l)
0.4
0.2
0.0
−0.2
−0.4
mean residual (CI: 95%)
F (n=19,343)
M (n=17,237)
All (n=36,580)
0
(0,1]
(1,2]
(2,3]
(3,4]
4+
daily units
two-tailed test against [0,1] group: ⋆p < 0.01, ⋆ ⋆ p < 0.001

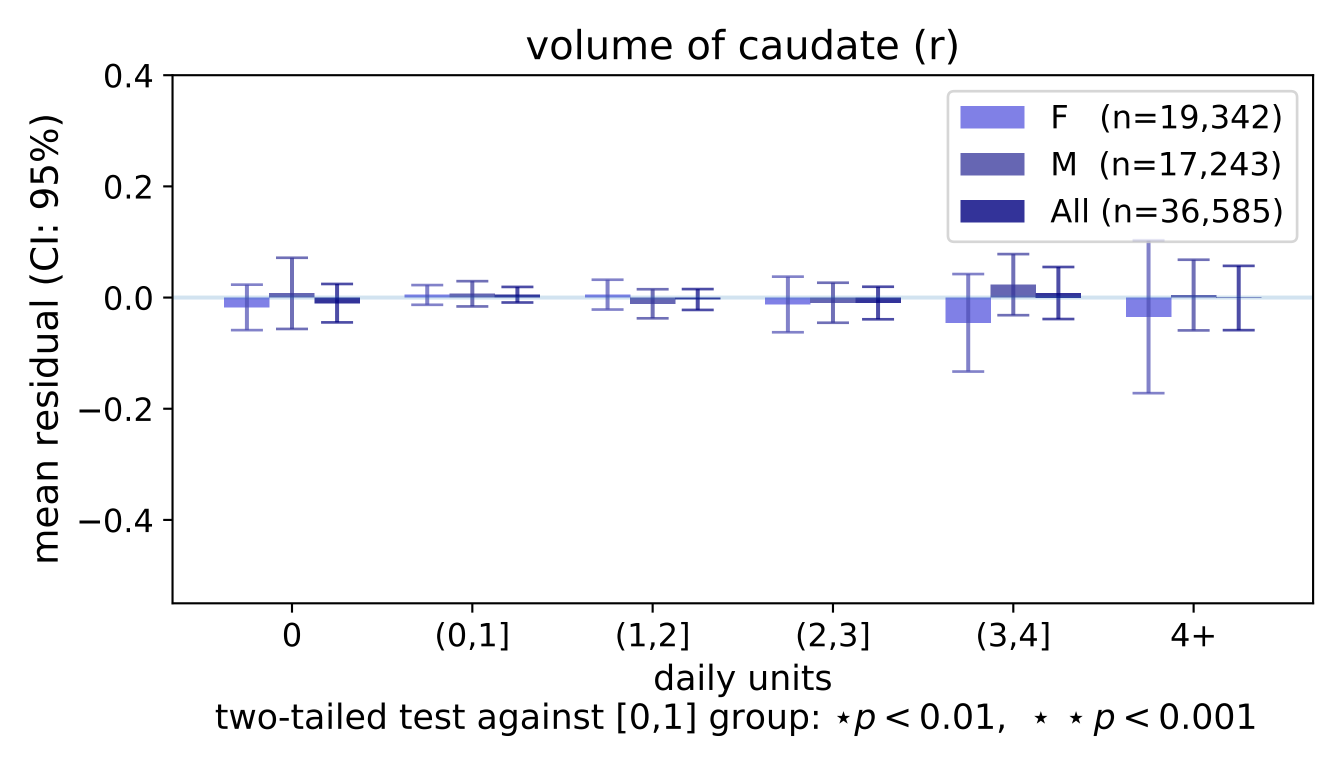
volume of caudate (r)
mean residual (CI: 95%)
0.4
0.2
0.0
−0.2
−0.4
F (n=19,342)
M (n=17,243)
All (n=36,585)
0
(0,1]
(1,2]
(2,3]
(3,4]
4+
daily units
two-tailed test against [0,1] group: ⋆p < 0.01, ⋆ ⋆ p < 0.001

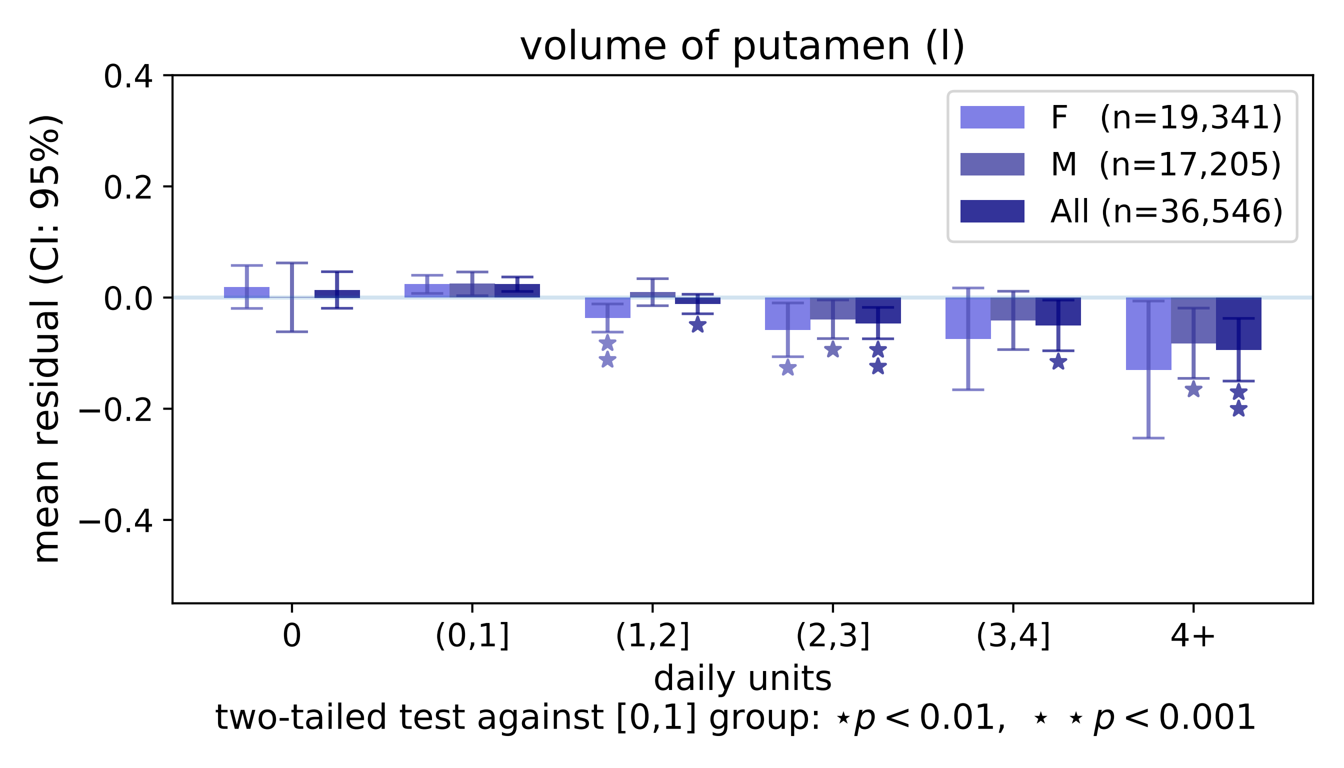

volume of putamen (l)
mean residual (CI: 95%)
0.4
0.2
0.0
−0.2
−0.4
F (n=19,341)
M (n=17,205)
All (n=36,546)
0
(0,1]
(1,2]
(2,3]
(3,4]
4+
daily units
two-tailed test against [0,1] group: ⋆ p < 0.01, ⋆ ⋆ p < 0.001

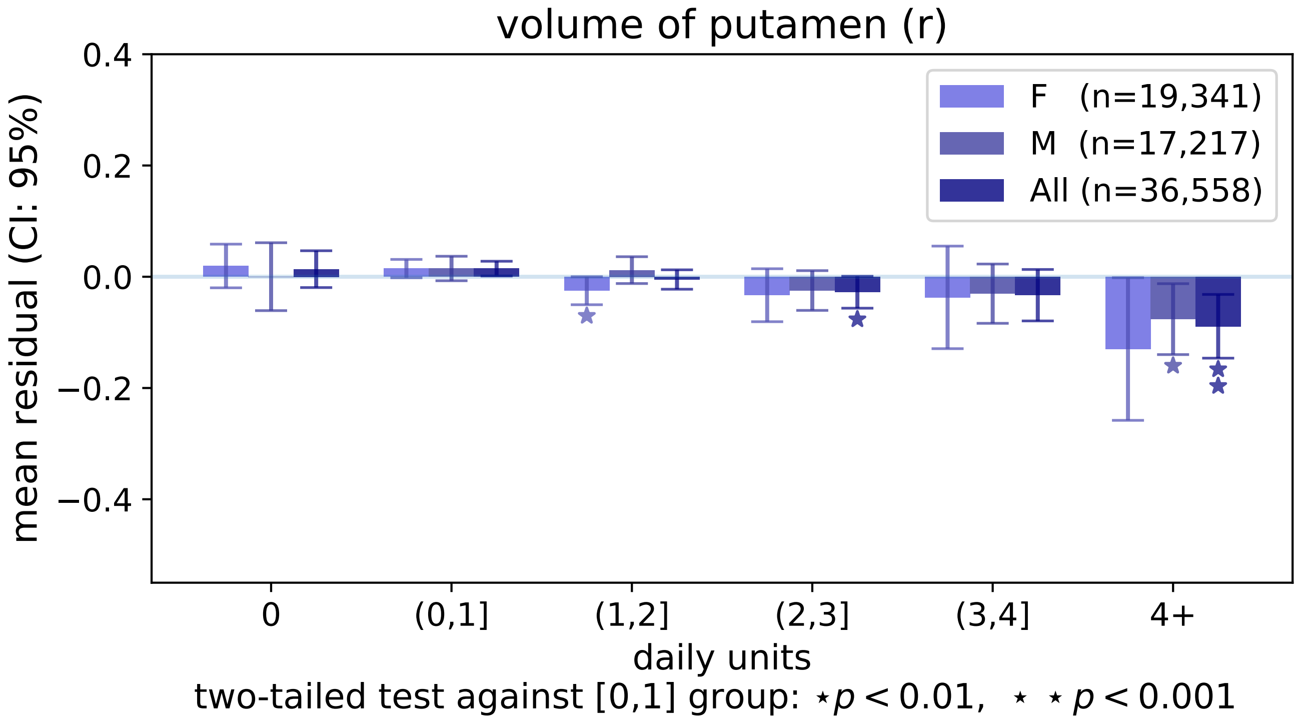

volume of putamen (r)
mean residual (CI: 95%)
0.4
0.2
0.0
−0.2
−0.4
F (n=19,341)
M (n=17,217)
All (n=36,558)
0
(0,1]
(1,2]
(2,3]
(3,4]
4+
daily units
two-tailed test against [0,1] group: ⋆ p < 0.01, ⋆ ⋆ p < 0.001

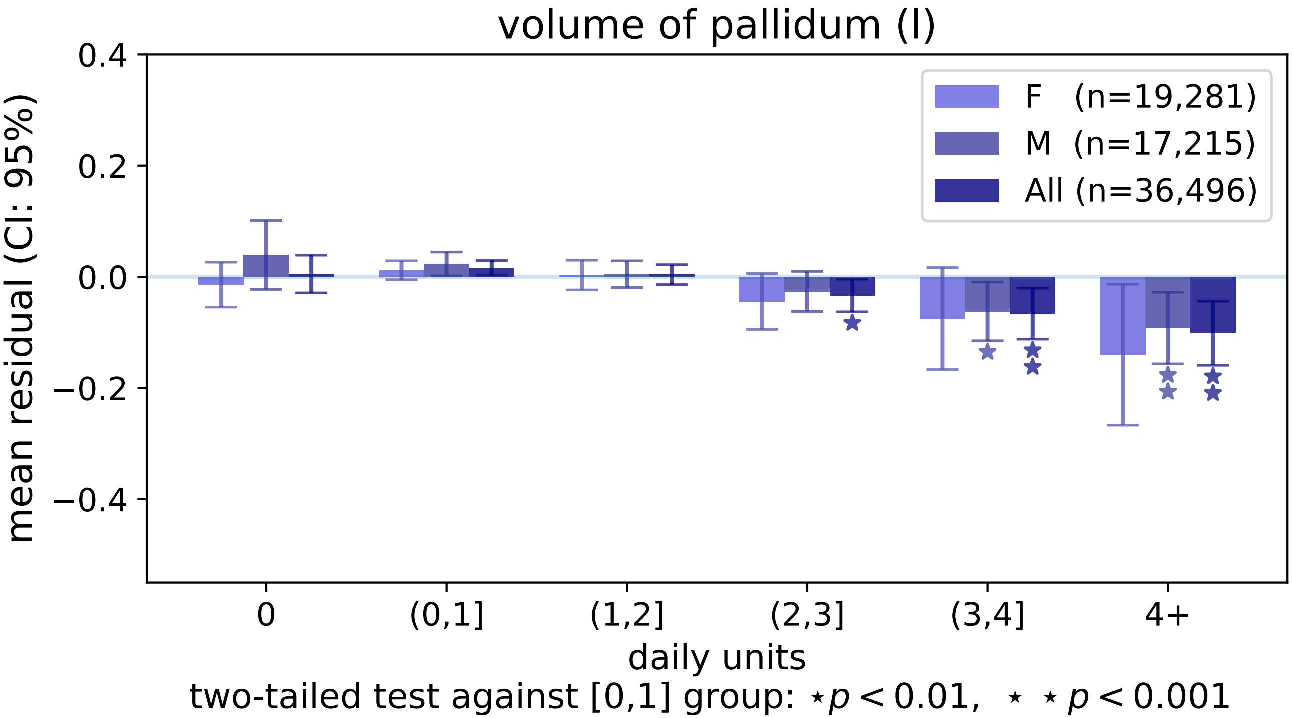

volume of pallidum (l)
0.4
0.2
0.0
−0.2
−0.4
mean residual (CI: 95%)
F (n=19,281)
M (n=17,215)
All (n=36,496)
0
(0,1]
(1,2]
(2,3]
(3,4]
4+
daily units
two-tailed test against [0,1] group: ⋆ $p < 0.01$, ⋆ ⋆ $p < 0.001$

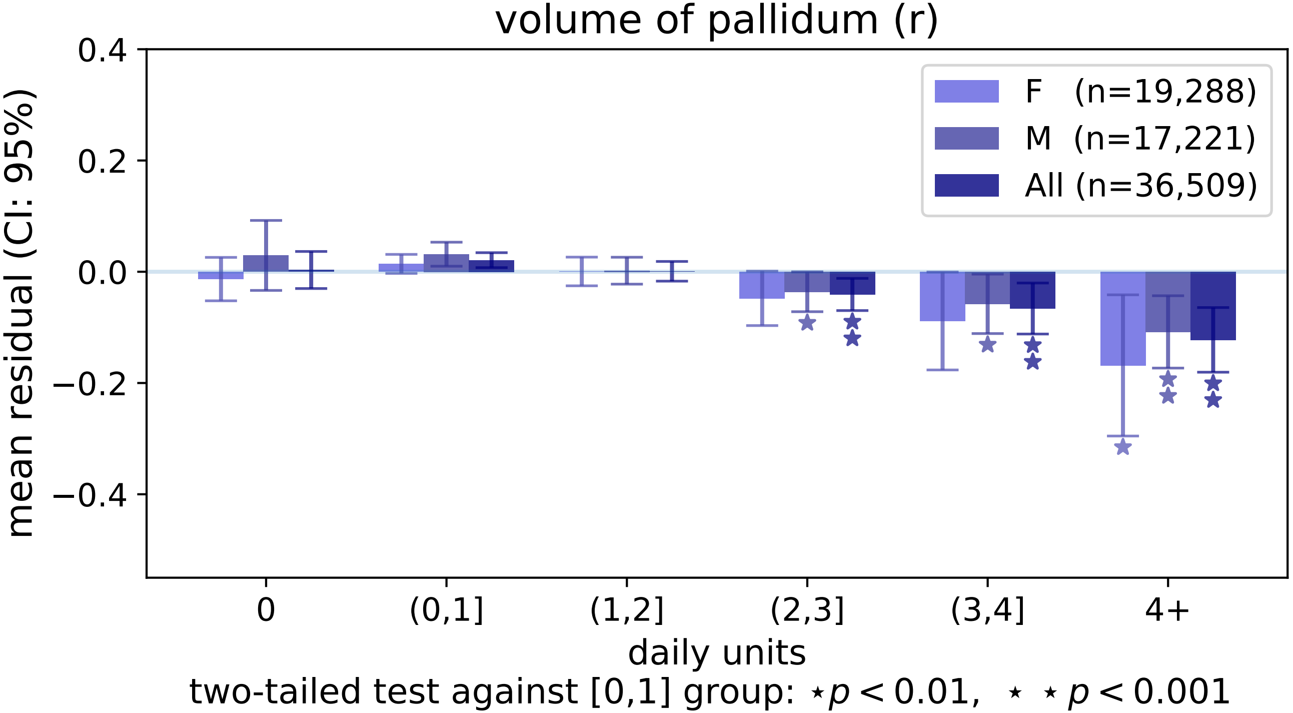

volume of pallidum (r)
0.4
0.2
0.0
−0.2
−0.4
mean residual (CI: 95%)
F (n=19,288)
M (n=17,221)
All (n=36,509)
0
(0,1]
(1,2]
(2,3]
(3,4]
4+
daily units
two-tailed test against [0,1] group: ⋆p < 0.01, ⋆ ⋆ p < 0.001

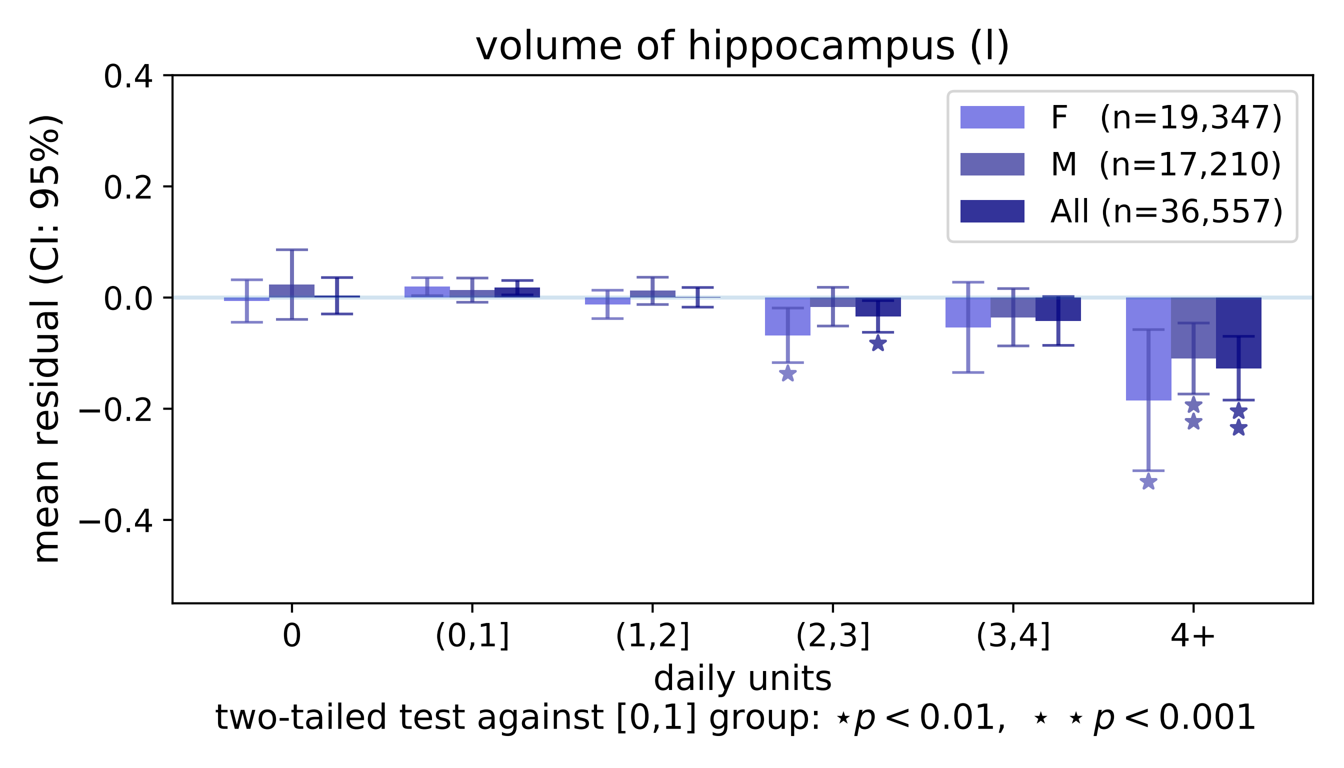
volume of hippocampus (l)
mean residual (CI: 95%)
0.4
0.2
0.0
−0.2
−0.4
F (n=19,347)
M (n=17,210)
All (n=36,557)
0
(0,1]
(1,2]
(2,3]
(3,4]
4+
daily units
two-tailed test against [0,1] group: ⋆ $p < 0.01$, ⋆ ⋆ $p < 0.001$

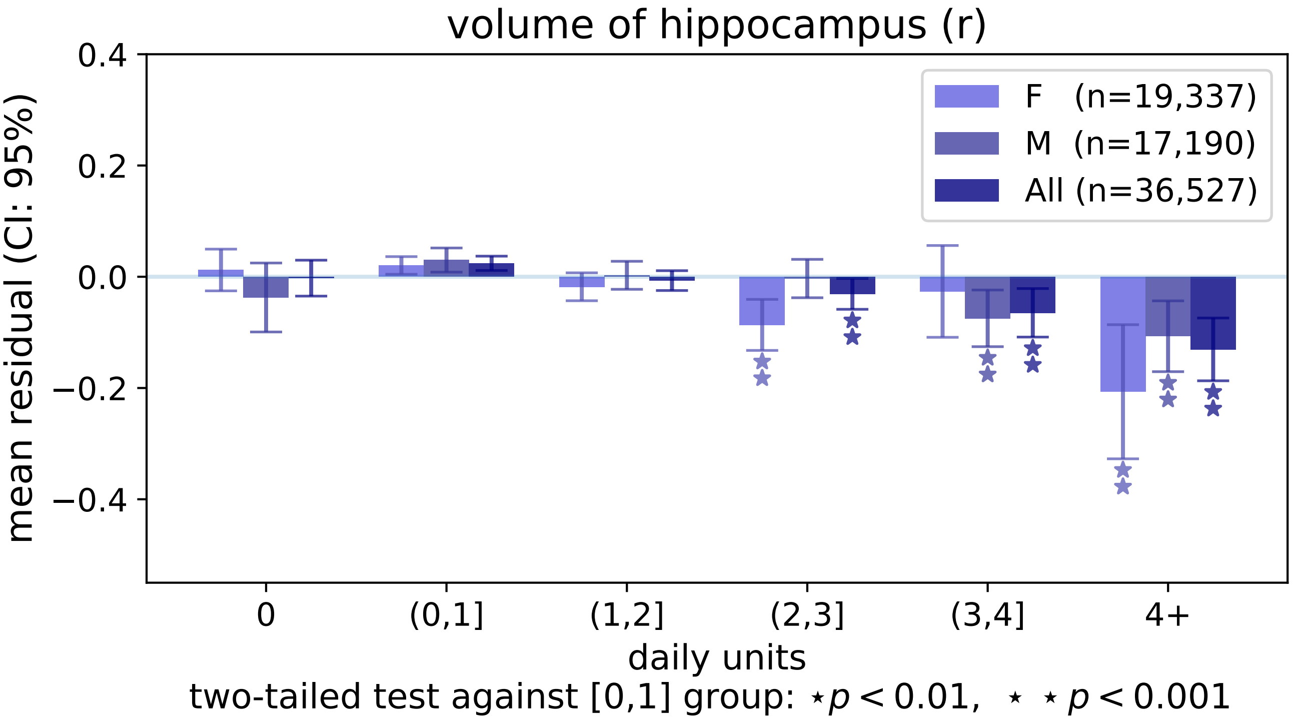

volume of hippocampus (r)
mean residual (CI: 95%)
0.4
0.2
0.0
−0.2
−0.4
F (n=19,337)
M (n=17,190)
All (n=36,527)
0
(0,1]
(1,2]
(2,3]
(3,4]
4+
daily units
two-tailed test against [0,1] group: ⋆ $p < 0.01$, ⋆ ⋆ $p < 0.001$

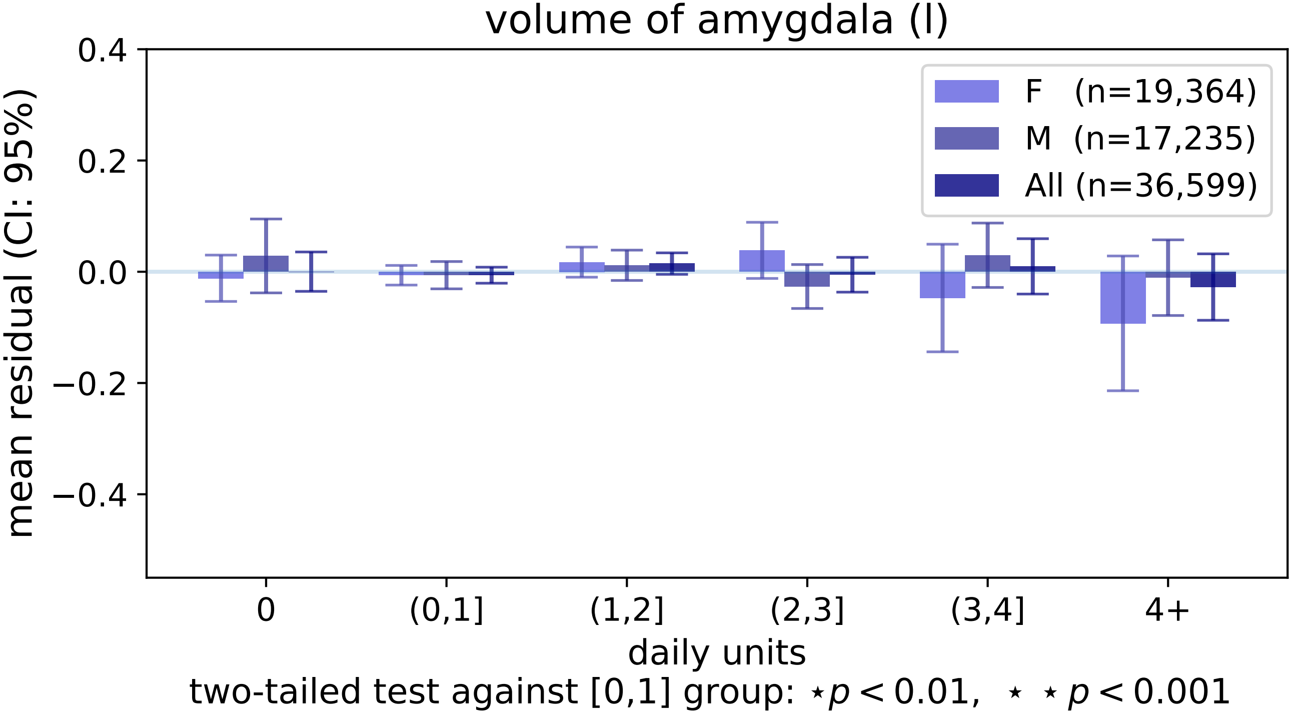

volume of amygdala (l)
0.4
0.2
0.0
−0.2
−0.4
mean residual (CI: 95%)
F (n=19,364)
M (n=17,235)
All (n=36,599)
0
(0,1]
(1,2]
(2,3]
(3,4]
4+
daily units
two-tailed test against [0,1] group: ⋆p < 0.01, ⋆ ⋆ p < 0.001

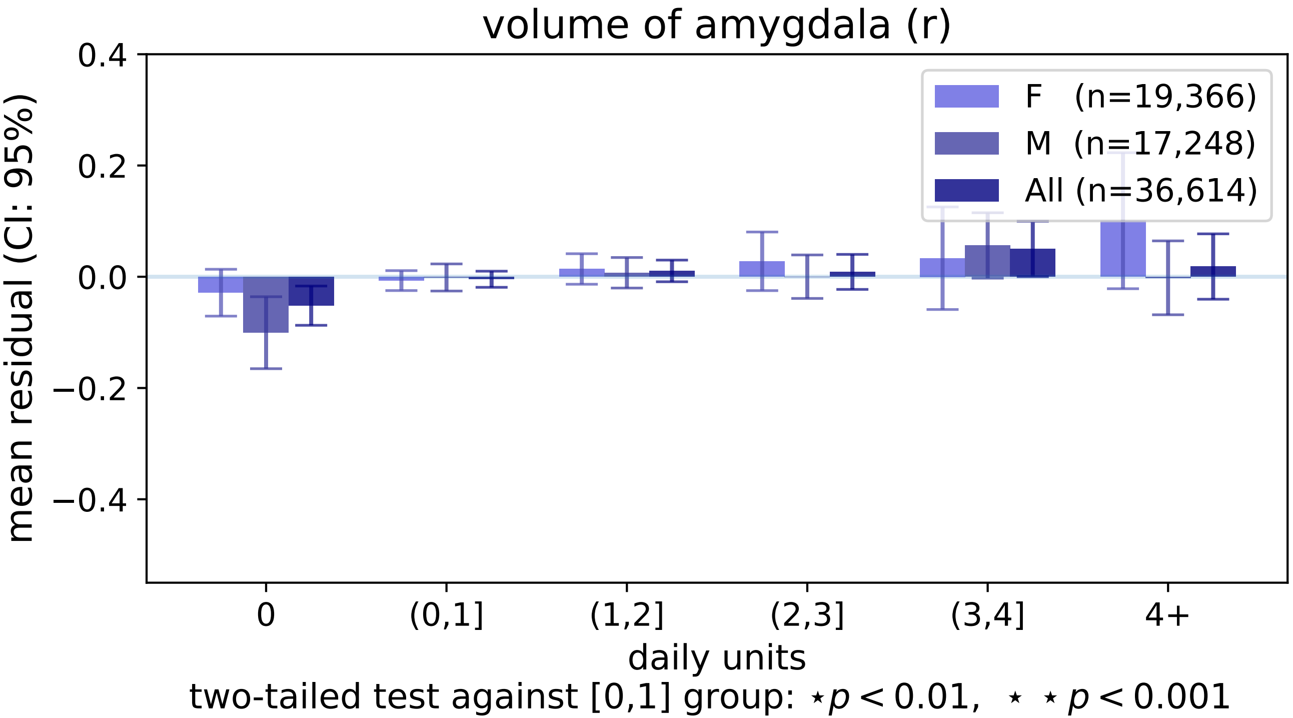
volume of amygdala (r)
mean residual (CI: 95%)
0.4
0.2
0.0
−0.2
−0.4
F (n=19,366)
M (n=17,248)
All (n=36,614)
0
(0,1]
(1,2]
(2,3]
(3,4]
4+
daily units
two-tailed test against [0,1] group: ⋆p < 0.01, ⋆ ⋆ p < 0.001

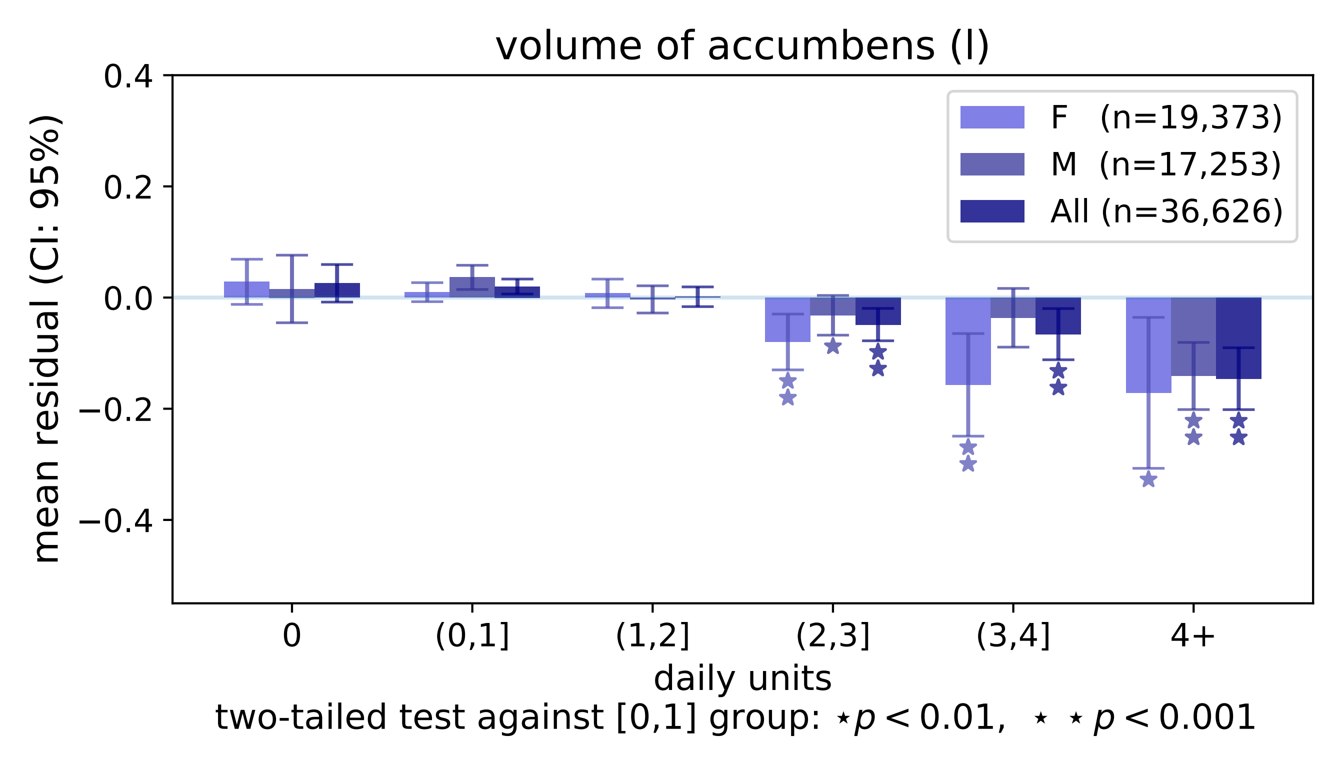
volume of accumbens (l)
0.4
0.2
0.0
−0.2
−0.4
mean residual (CI: 95%)
F (n=19,373)
M (n=17,253)
All (n=36,626)
0
(0,1]
(1,2]
(2,3]
(3,4]
4+
daily units
two-tailed test against [0,1] group: ⋆p < 0.01, ⋆ ⋆ p < 0.001

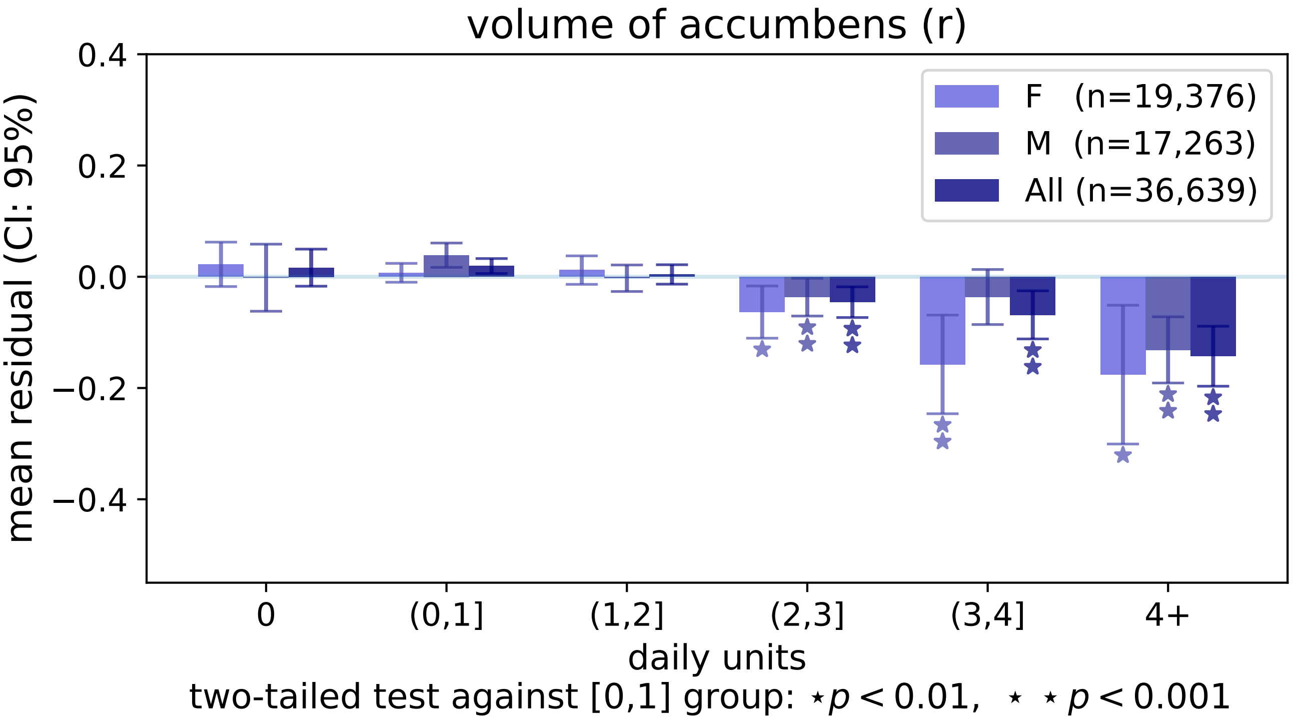

volume of accumbens (r)
0.4
0.2
0.0
−0.2
−0.4
mean residual (CI: 95%)
F (n=19,376)
M (n=17,263)
All (n=36,639)
0
(0,1]
(1,2]
(2,3]
(3,4]
4+
daily units
two-tailed test against [0,1] group: ⋆p < 0.01, ⋆ ⋆ p < 0.001

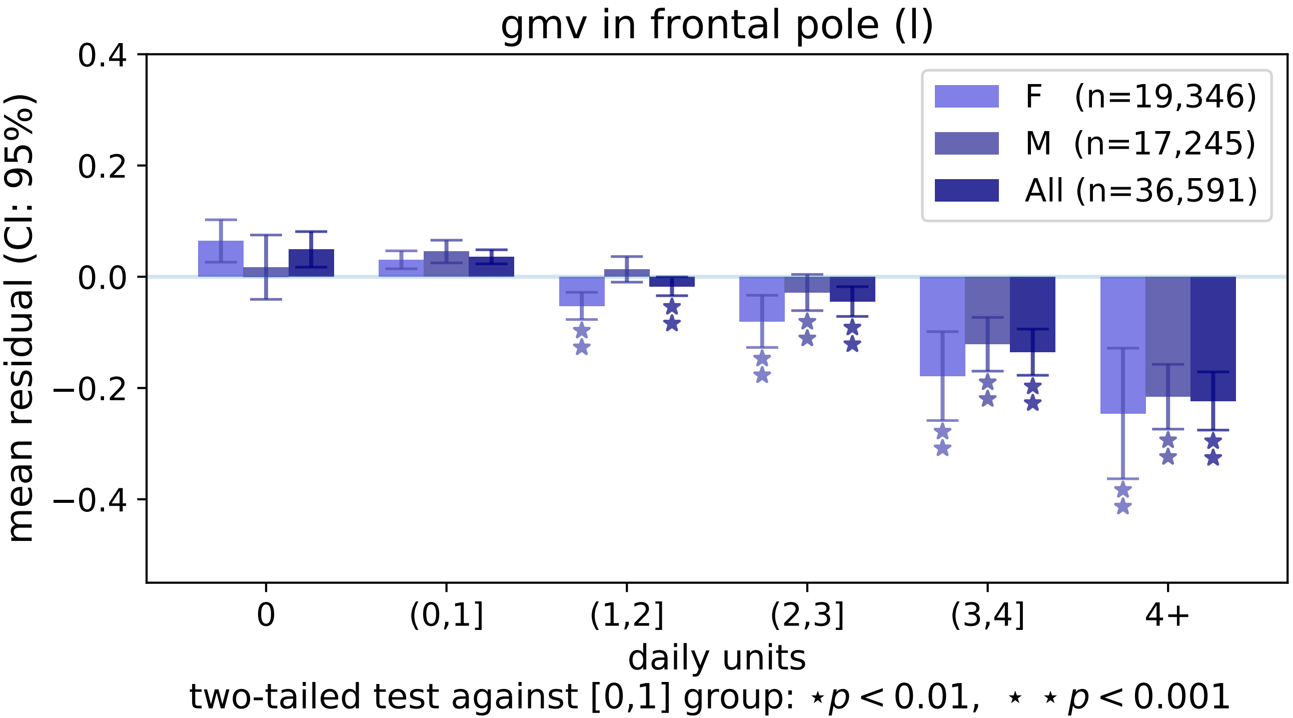
gmv in frontal pole (l)
mean residual (CI: 95%)
0.4
0.2
0.0
−0.2
−0.4
F (n=19,346)
M (n=17,245)
All (n=36,591)
0
(0,1]
(1,2]
(2,3]
(3,4]
4+
daily units
two-tailed test against [0,1] group: ⋆ p < 0.01, ⋆ ⋆ p < 0.001

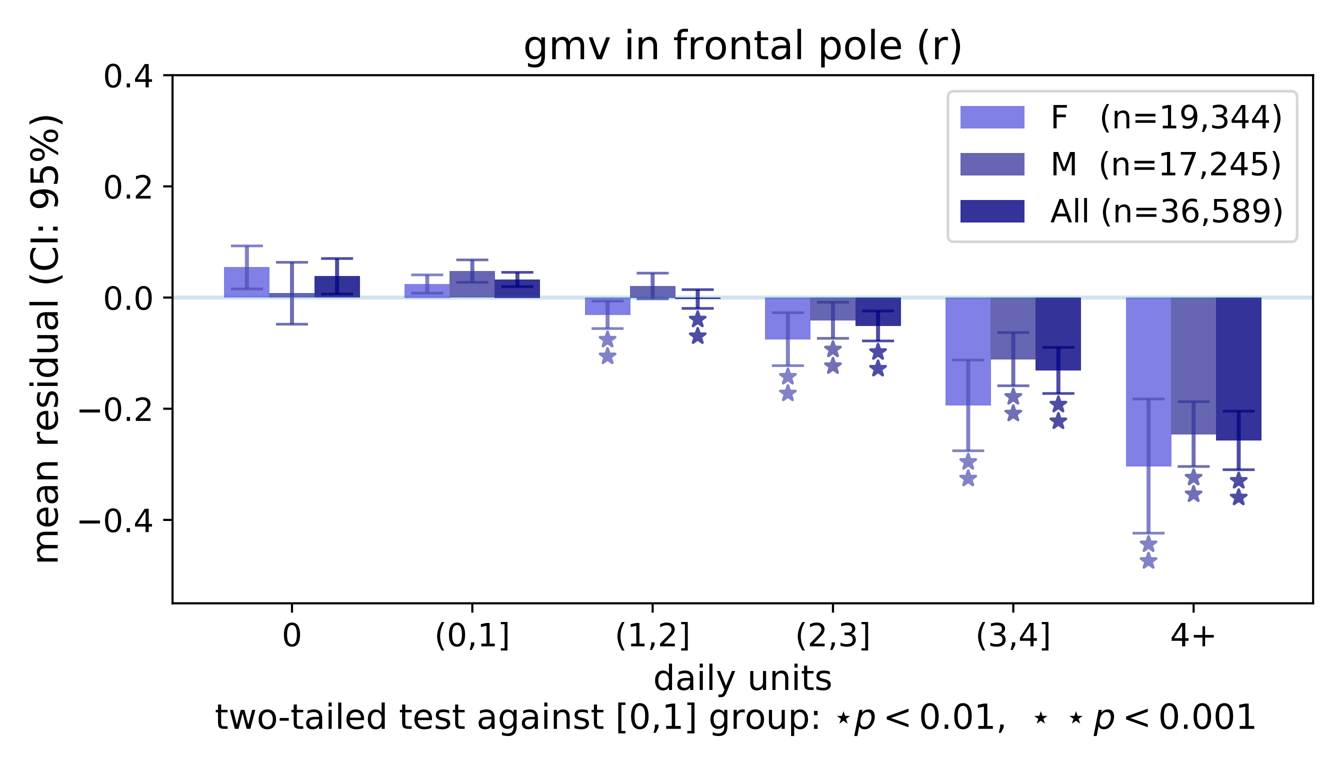

gmv in frontal pole (r)
mean residual (CI: 95%)
0.4
0.2
0.0
−0.2
−0.4
F (n=19,344)
M (n=17,245)
All (n=36,589)
0
(0,1]
(1,2]
(2,3]
(3,4]
4+
daily units
two-tailed test against [0,1] group: ⋆p < 0.01, ⋆ ⋆p < 0.001

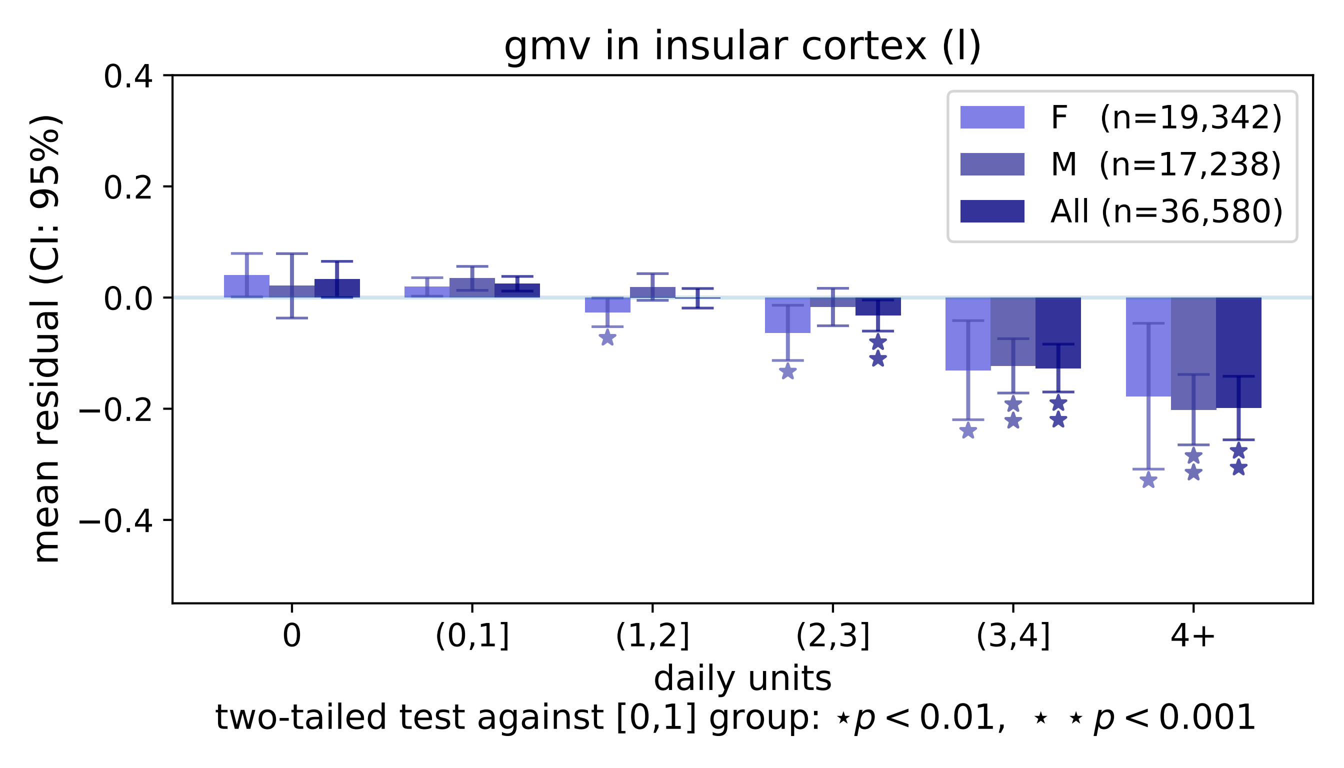
gmv in insular cortex (l)
mean residual (CI: 95%)
0.4
0.2
0.0
−0.2
−0.4
F (n=19,342)
M (n=17,238)
All (n=36,580)
0
(0,1]
(1,2]
(2,3]
(3,4]
4+
daily units
two-tailed test against [0,1] group: ⋆ p < 0.01, ⋆ ⋆ p < 0.001

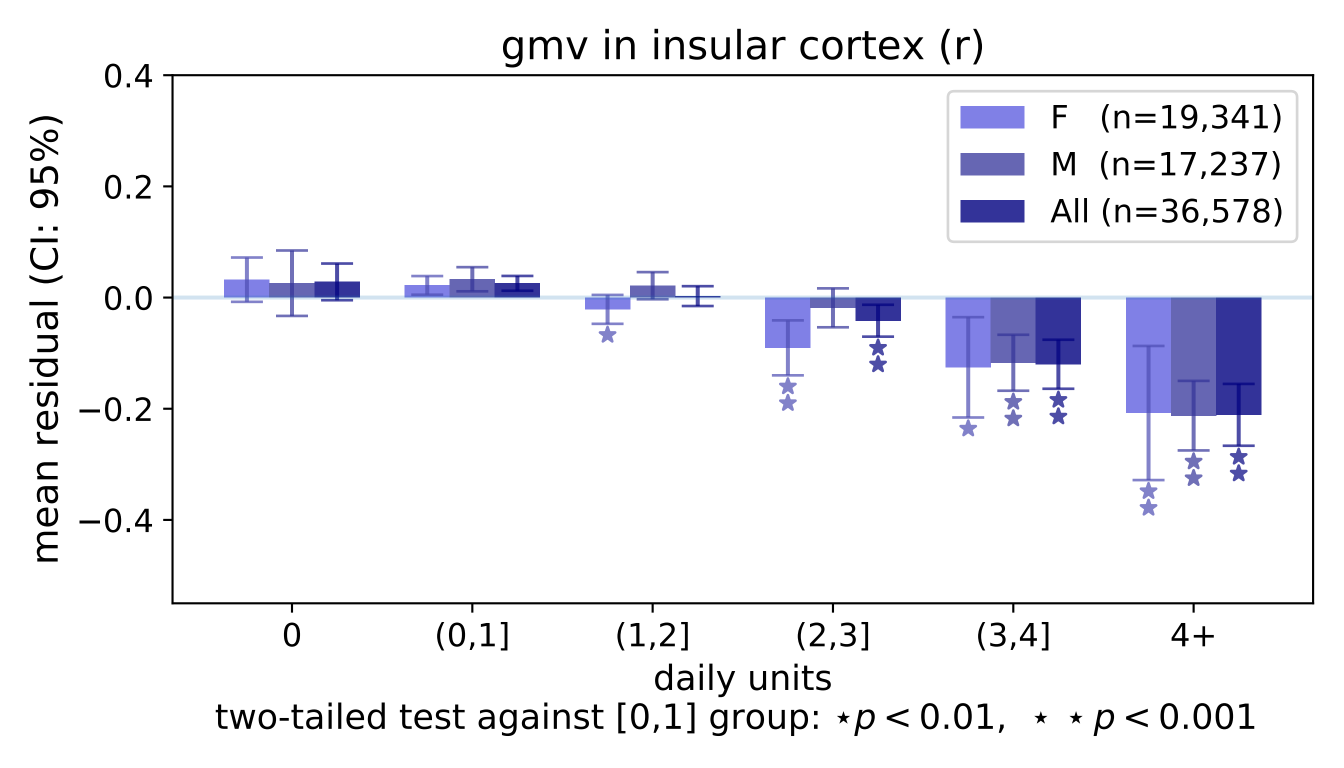

gmv in insular cortex (r)
mean residual (CI: 95%)
0.4
0.2
0.0
−0.2
−0.4
F (n=19,341)
M (n=17,237)
All (n=36,578)
0
(0,1]
(1,2]
(2,3]
(3,4]
4+
daily units
two-tailed test against [0,1] group: ⋆p < 0.01, ⋆ ⋆ p < 0.001

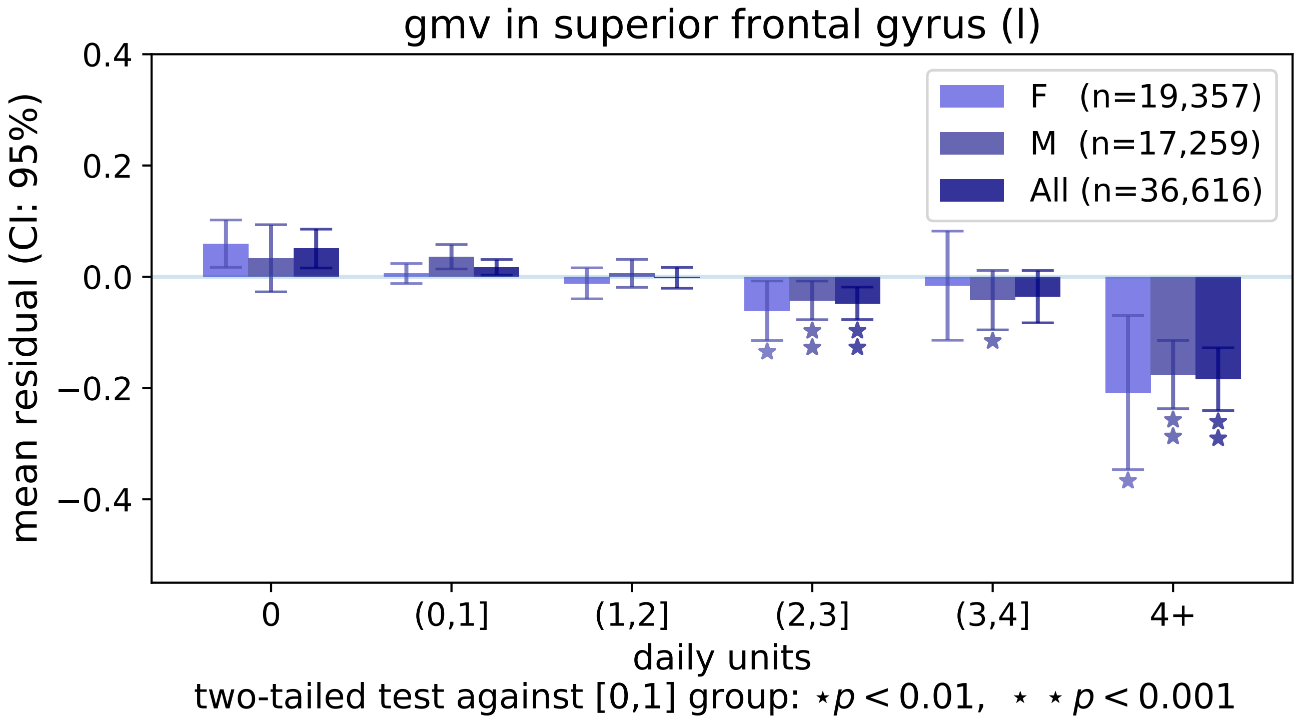
gmv in superior frontal gyrus (l)
0.4
0.2
0.0
−0.2
−0.4
mean residual (CI: 95%)
F (n=19,357)
M (n=17,259)
All (n=36,616)
0
(0,1]
(1,2]
(2,3]
(3,4]
4+
daily units
two-tailed test against [0,1] group: ⋆ $p < 0.01$, ⋆ ⋆ $p < 0.001$

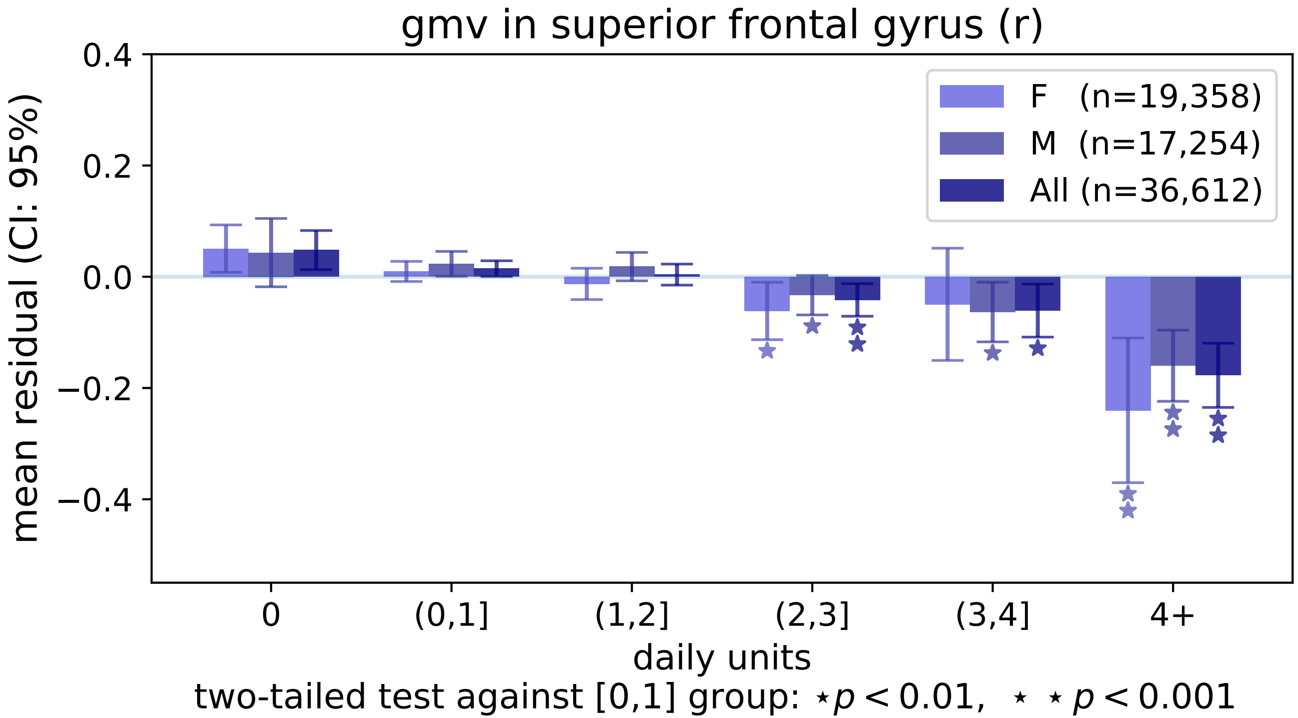
gmv in superior frontal gyrus (r)
mean residual (CI: 95%)
0.4
0.2
0.0
−0.2
−0.4
F (n=19,358)
M (n=17,254)
All (n=36,612)
0
(0,1]
(1,2]
(2,3]
(3,4]
4+
daily units
two-tailed test against [0,1] group: ⋆ p < 0.01, ⋆ ⋆ p < 0.001

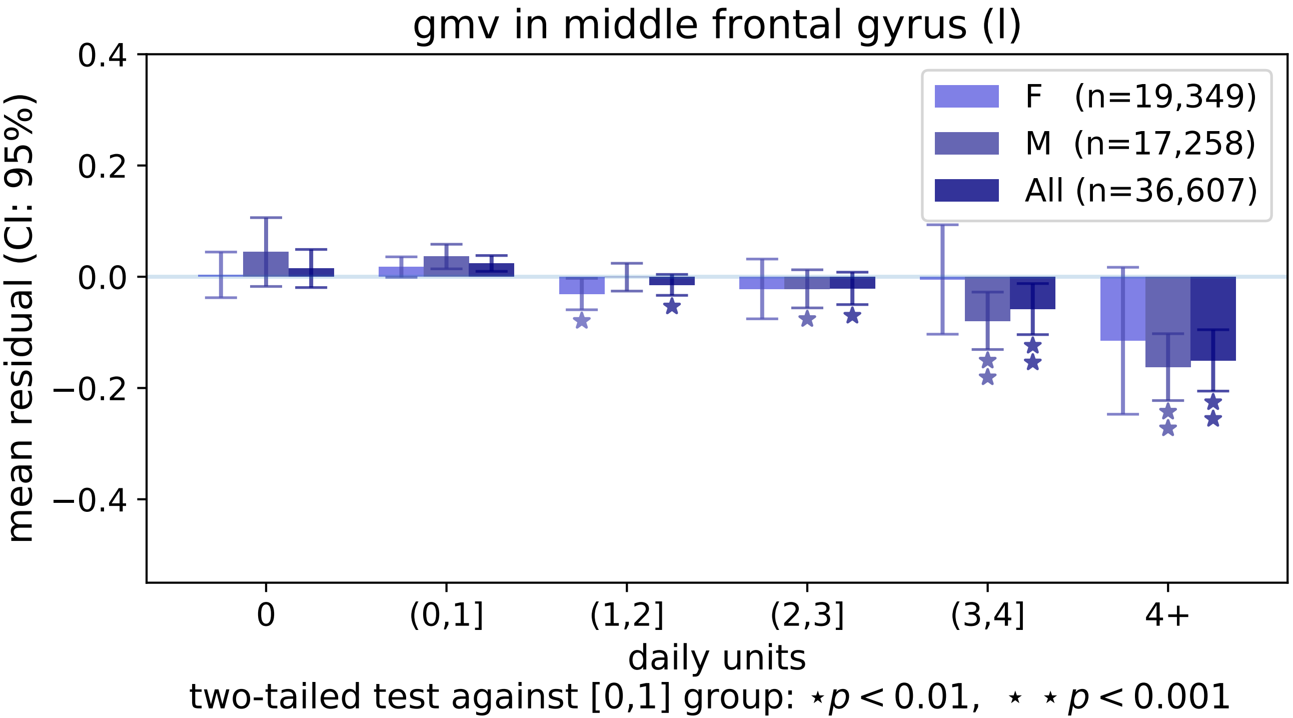

gmv in middle frontal gyrus (l)
mean residual (CI: 95%)
0.4
0.2
0.0
−0.2
−0.4
F (n=19,349)
M (n=17,258)
All (n=36,607)
0
(0,1]
(1,2]
(2,3]
(3,4]
4+
daily units
two-tailed test against [0,1] group: ⋆ $p < 0.01$, ⋆ ⋆ $p < 0.001$

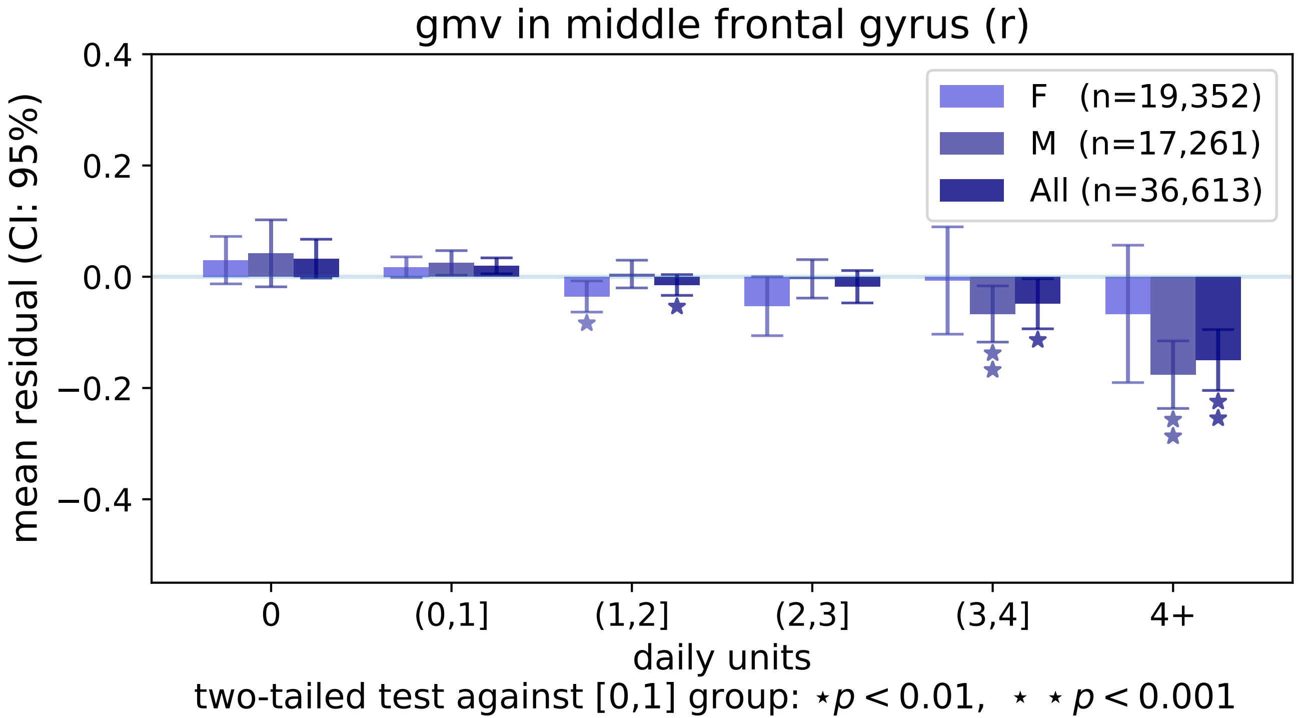

gmv in middle frontal gyrus (r)
0.4
0.2
0.0
−0.2
−0.4
mean residual (CI: 95%)
F (n=19,352)
M (n=17,261)
All (n=36,613)
0
(0,1]
(1,2]
(2,3]
(3,4]
4+
daily units
two-tailed test against [0,1] group: ⋆p < 0.01, ⋆ ⋆ p < 0.001

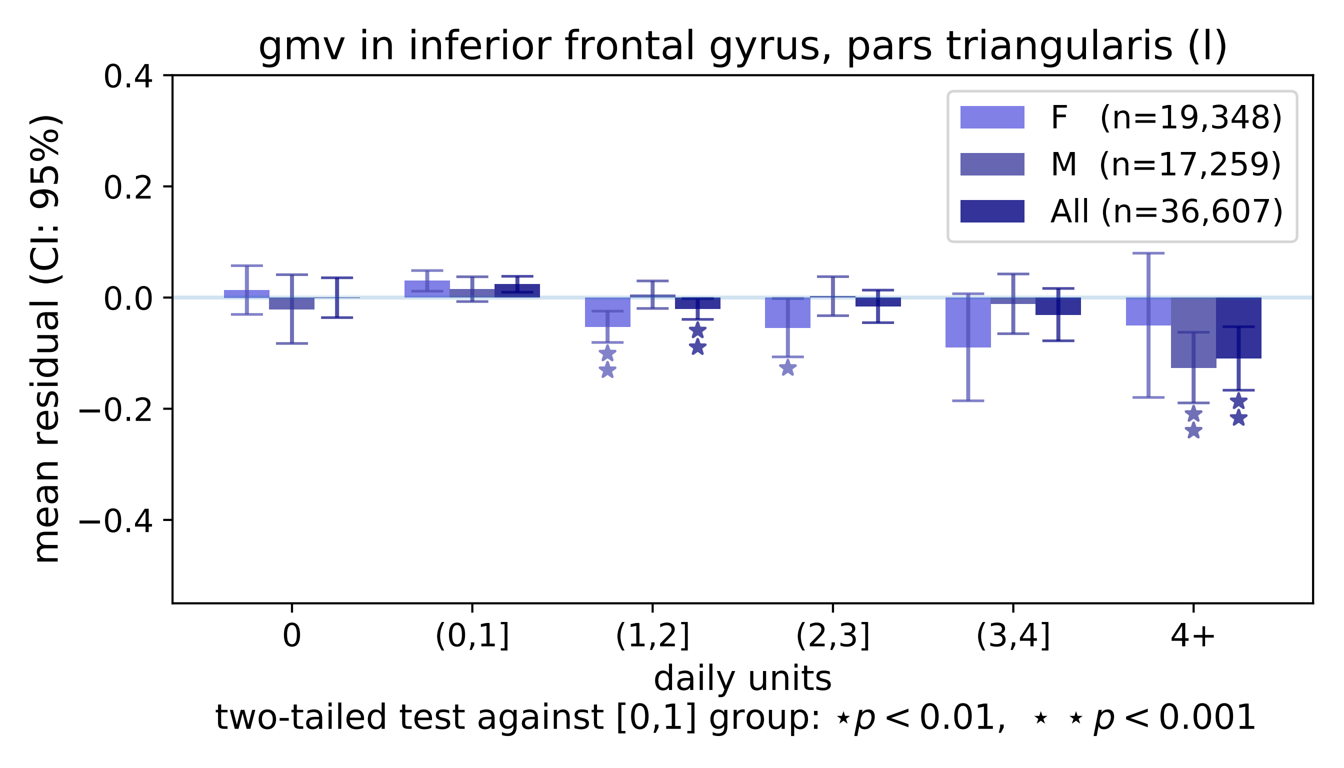

gmv in inferior frontal gyrus, pars triangularis (l)
mean residual (CI: 95%)
0.4
0.2
0.0
−0.2
−0.4
F (n=19,348)
M (n=17,259)
All (n=36,607)
0
(0,1]
(1,2]
(2,3]
(3,4]
4+
daily units
two-tailed test against [0,1] group: ⋆ p < 0.01, ⋆ ⋆ p < 0.001

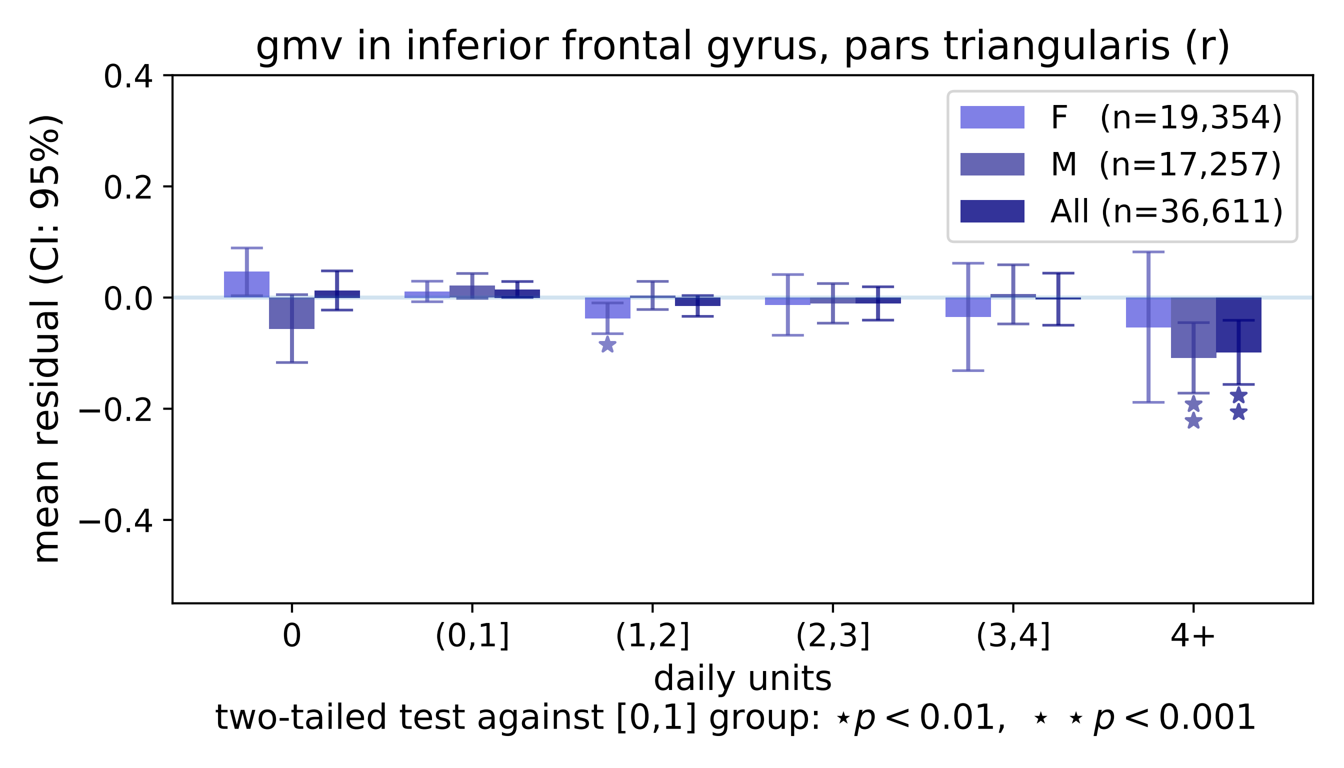
gmv in inferior frontal gyrus, pars triangularis (r)
0.4
0.2
0.0
−0.2
−0.4
mean residual (CI: 95%)
F (n=19,354)
M (n=17,257)
All (n=36,611)
0
(0,1]
(1,2]
(2,3]
(3,4]
4+
daily units
two-tailed test against [0,1] group: ⋆p < 0.01, ⋆ ⋆ p < 0.001

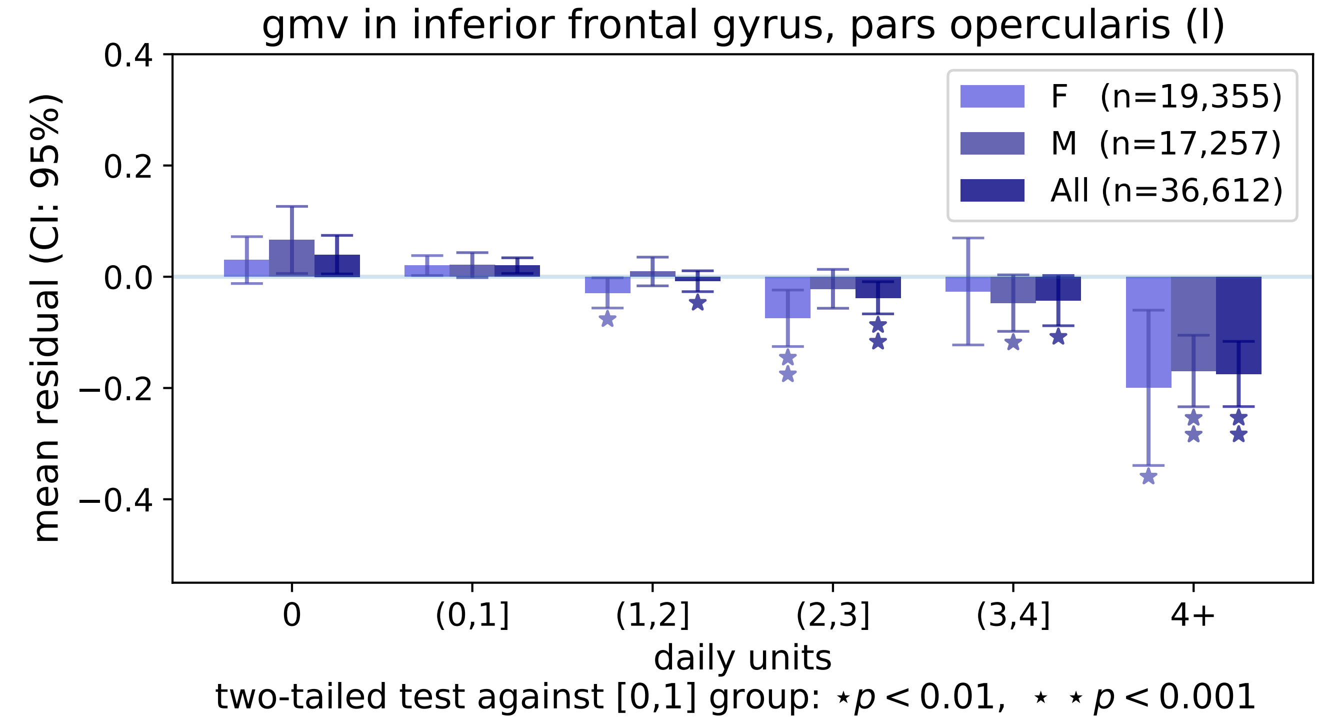
gmv in inferior frontal gyrus, pars opercularis (l)
mean residual (CI: 95%)
0.4
0.2
0.0
−0.2
−0.4
F (n=19,355)
M (n=17,257)
All (n=36,612)
0
(0,1]
(1,2]
(2,3]
(3,4]
4+
daily units
two-tailed test against [0,1] group: ⋆ p < 0.01, ⋆ ⋆ p < 0.001

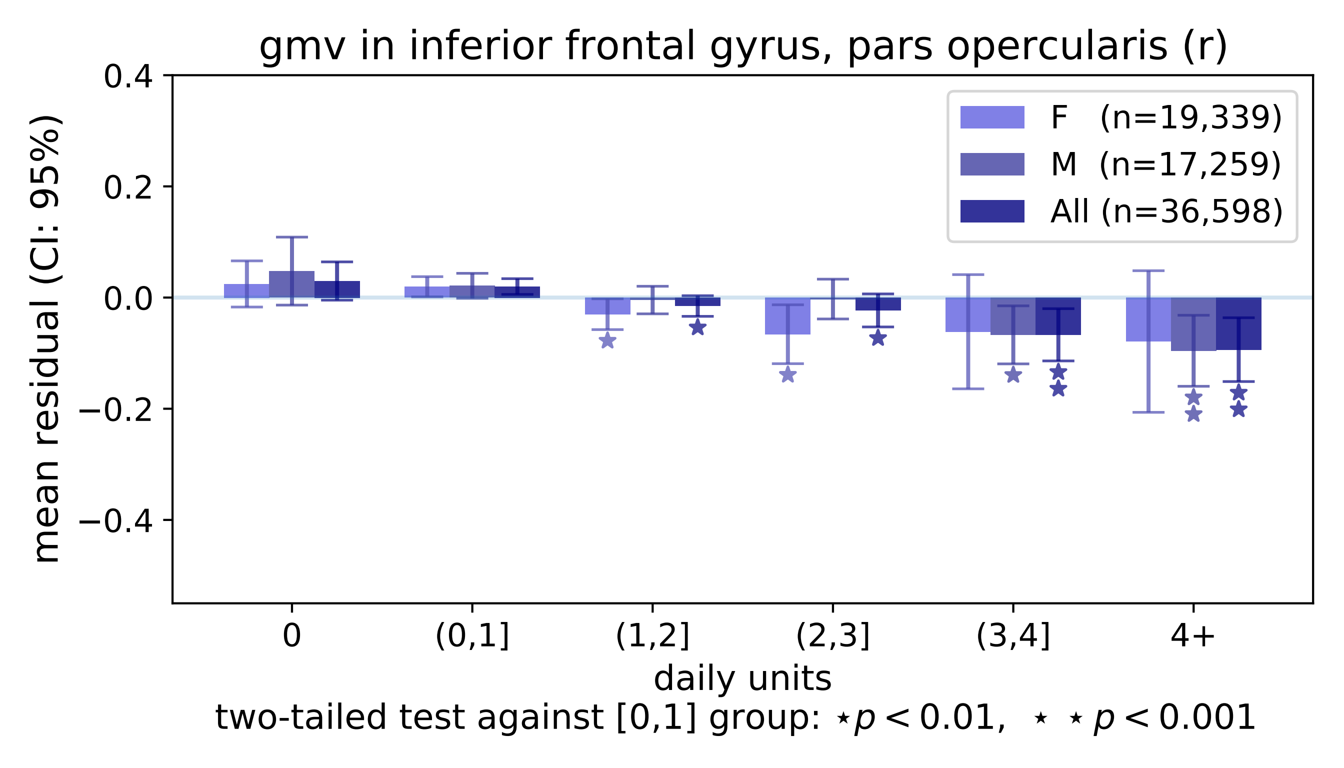

gmv in inferior frontal gyrus, pars opercularis (r)
F (n=19,339)
M (n=17,259)
All (n=36,598)
mean residual (CI: 95%)
0.4
0.2
0.0
−0.2
−0.4
0
(0,1]
(1,2]
(2,3]
(3,4]
4+
daily units
two-tailed test against [0,1] group: ⋆p < 0.01, ⋆ ⋆p < 0.001

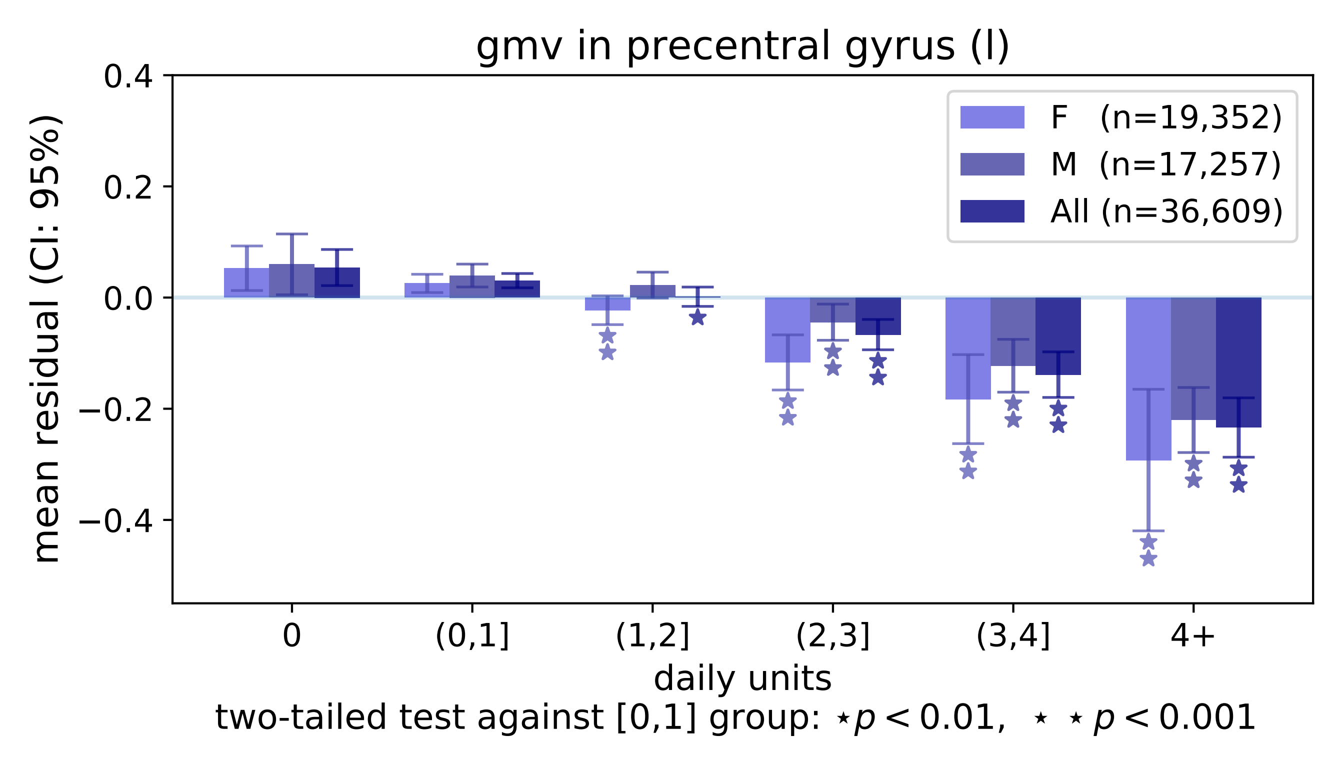

gmv in precentral gyrus (l)
mean residual (CI: 95%)
0.4
0.2
0.0
−0.2
−0.4
F (n=19,352)
M (n=17,257)
All (n=36,609)
0
(0,1]
(1,2]
(2,3]
(3,4]
4+
daily units
two-tailed test against [0,1] group: ⋆p < 0.01, ⋆ ⋆ p < 0.001

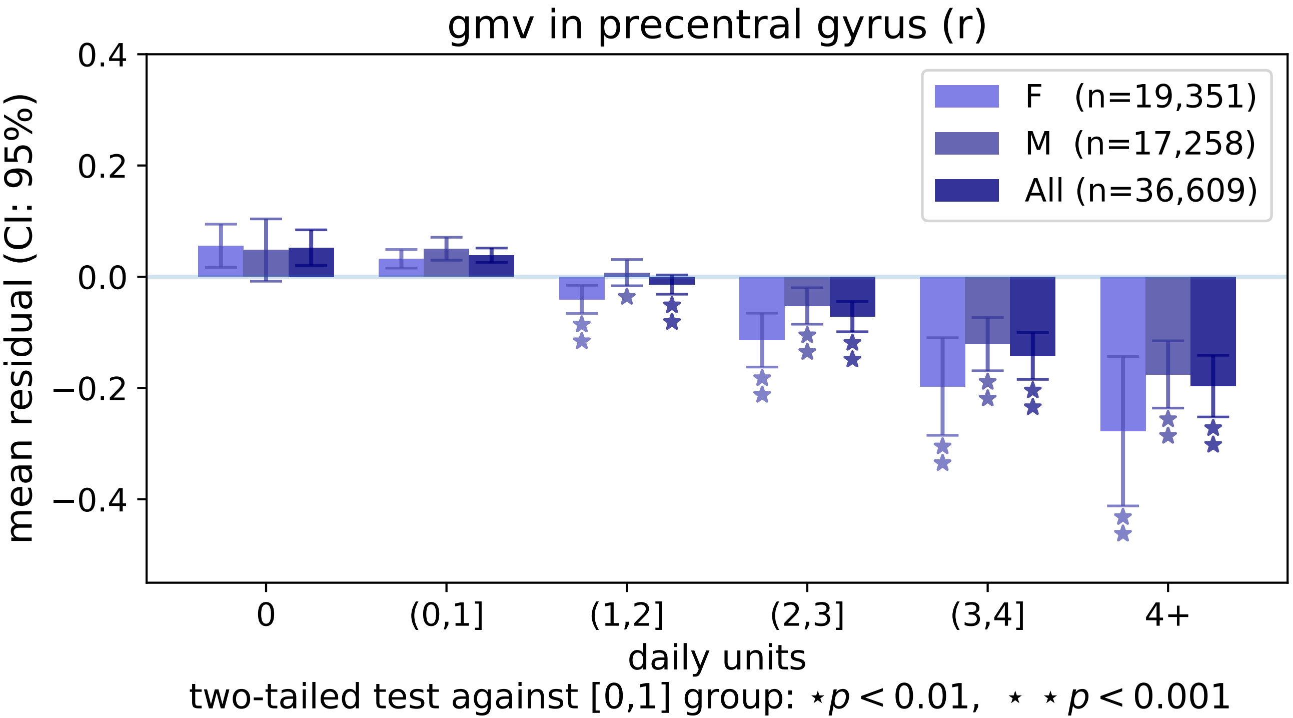
gmv in precentral gyrus (r)
mean residual (CI: 95%)
0.4
0.2
0.0
−0.2
−0.4
F (n=19,351)
M (n=17,258)
All (n=36,609)
0
(0,1]
(1,2]
(2,3]
(3,4]
4+
daily units
two-tailed test against [0,1] group: ⋆ $p < 0.01$, ⋆ ⋆ $p < 0.001$

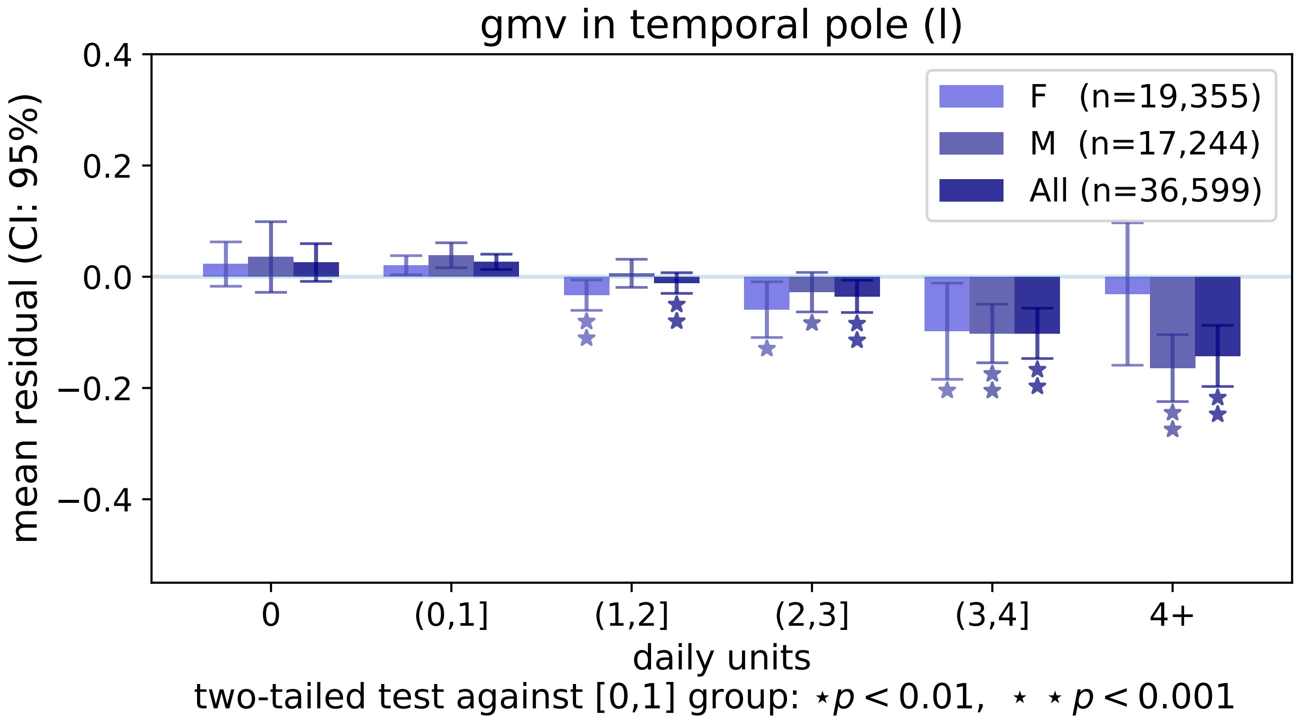
gmv in temporal pole (l)
mean residual (CI: 95%)
0.4
0.2
0.0
−0.2
−0.4
F (n=19,355)
M (n=17,244)
All (n=36,599)
0
(0,1]
(1,2]
(2,3]
(3,4]
4+
daily units
two-tailed test against [0,1] group: ⋆p < 0.01, ⋆ ⋆ p < 0.001

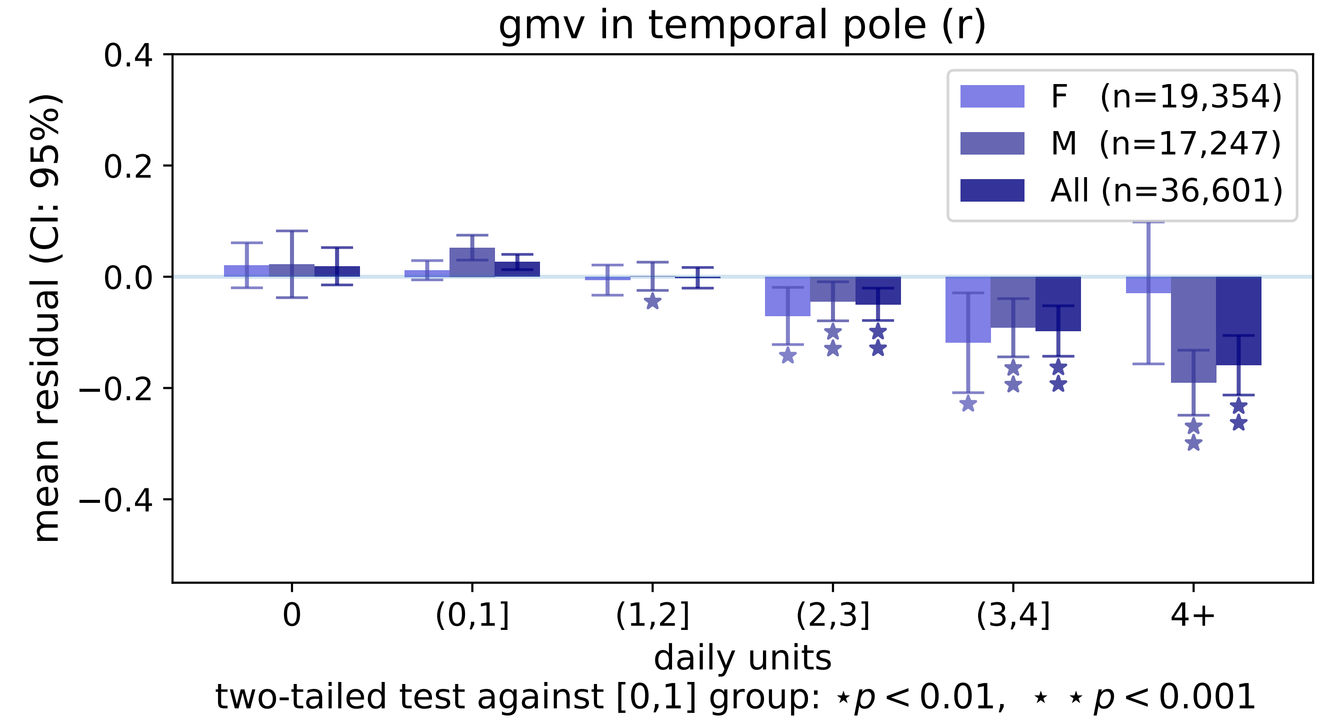

gmv in temporal pole (r)
F (n=19,354)
M (n=17,247)
All (n=36,601)
mean residual (CI: 95%)
0.4
0.2
0.0
−0.2
−0.4
0
(0,1]
(1,2]
(2,3]
(3,4]
4+
daily units
two-tailed test against [0,1] group: ⋆ p < 0.01, ⋆ ⋆ p < 0.001

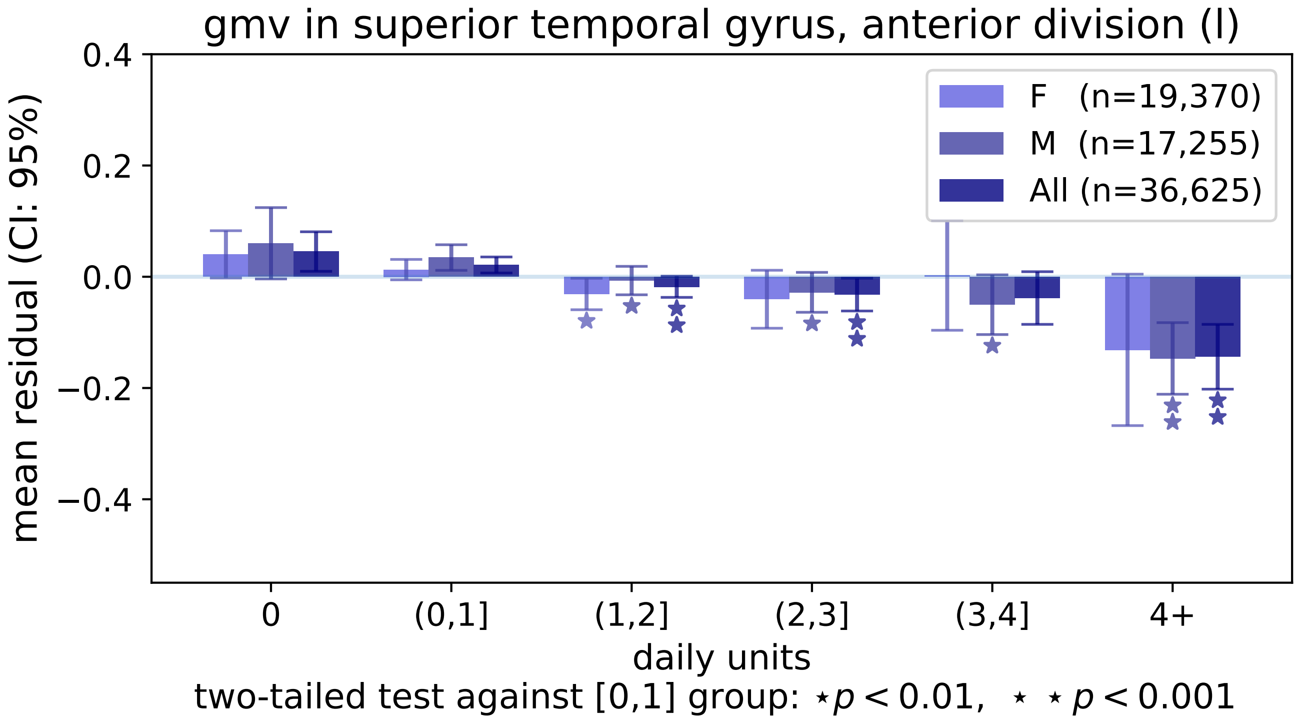

gmv in superior temporal gyrus, anterior division (l)
mean residual (CI: 95%)
0.4
0.2
0.0
−0.2
−0.4
F (n=19,370)
M (n=17,255)
All (n=36,625)
0
(0,1]
(1,2]
(2,3]
(3,4]
4+
daily units
two-tailed test against [0,1] group: ⋆ $p < 0.01$, ⋆ ⋆ $p < 0.001$

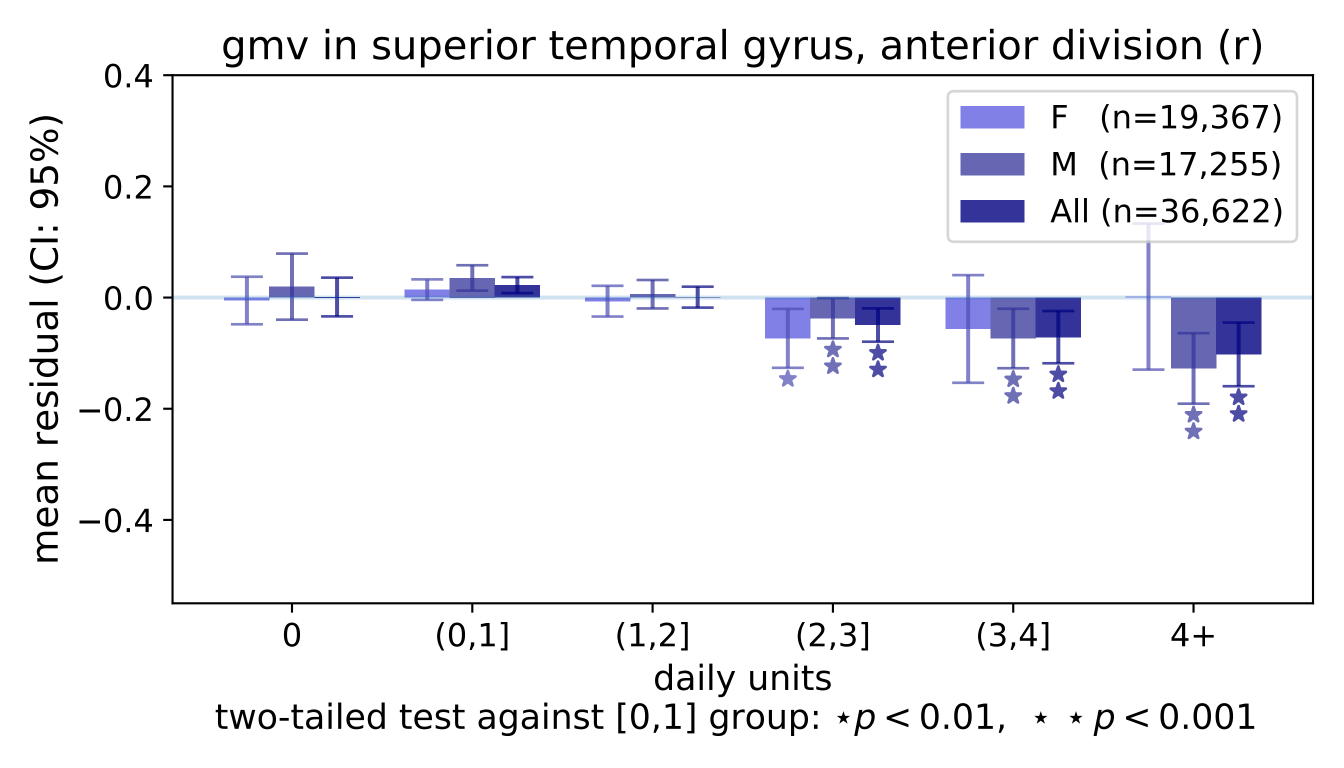
gmv in superior temporal gyrus, anterior division (r)
0.4
0.2
0.0
−0.2
−0.4
mean residual (CI: 95%)
F (n=19,367)
M (n=17,255)
All (n=36,622)
0
(0,1]
(1,2]
(2,3]
(3,4]
4+
daily units
two-tailed test against [0,1] group: ⋆ p < 0.01, ⋆ ⋆ p < 0.001

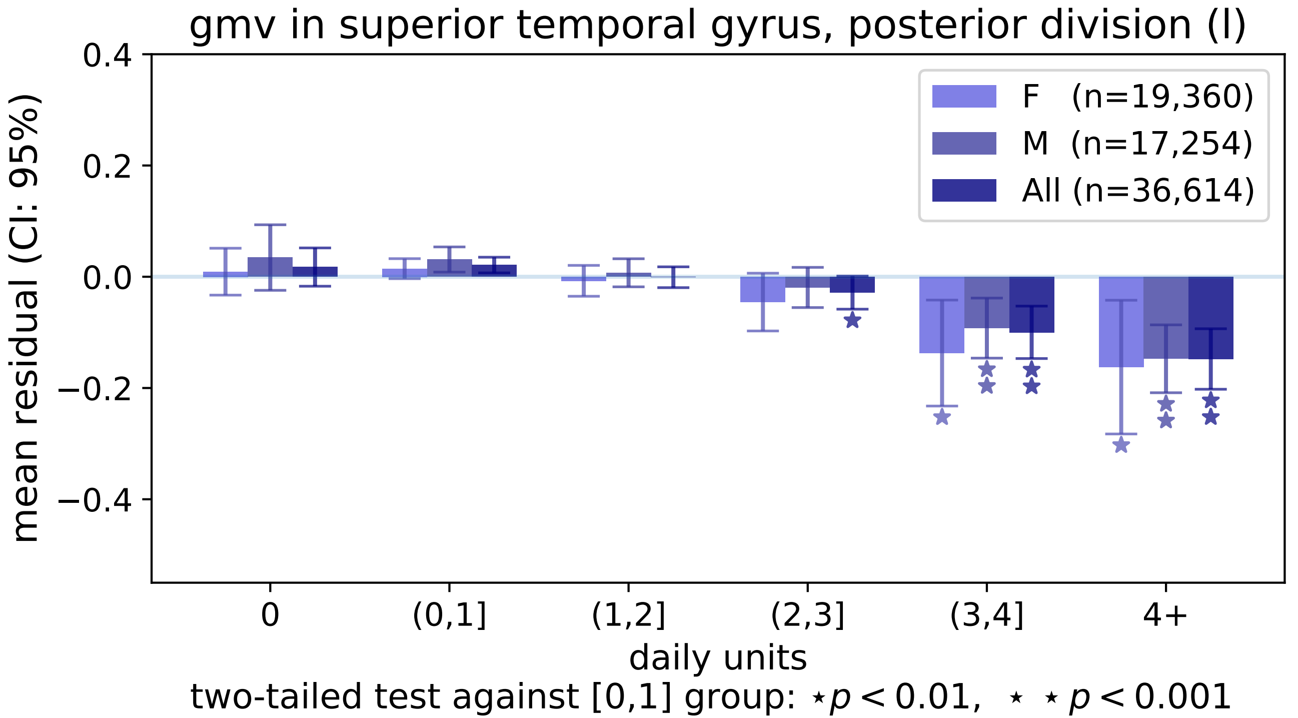

gmv in superior temporal gyrus, posterior division (l)
mean residual (CI: 95%)
0.4
0.2
0.0
−0.2
−0.4
F (n=19,360)
M (n=17,254)
All (n=36,614)
0
(0,1]
(1,2]
(2,3]
(3,4]
4+
daily units
two-tailed test against [0,1] group: ⋆ p < 0.01, ⋆ ⋆ p < 0.001

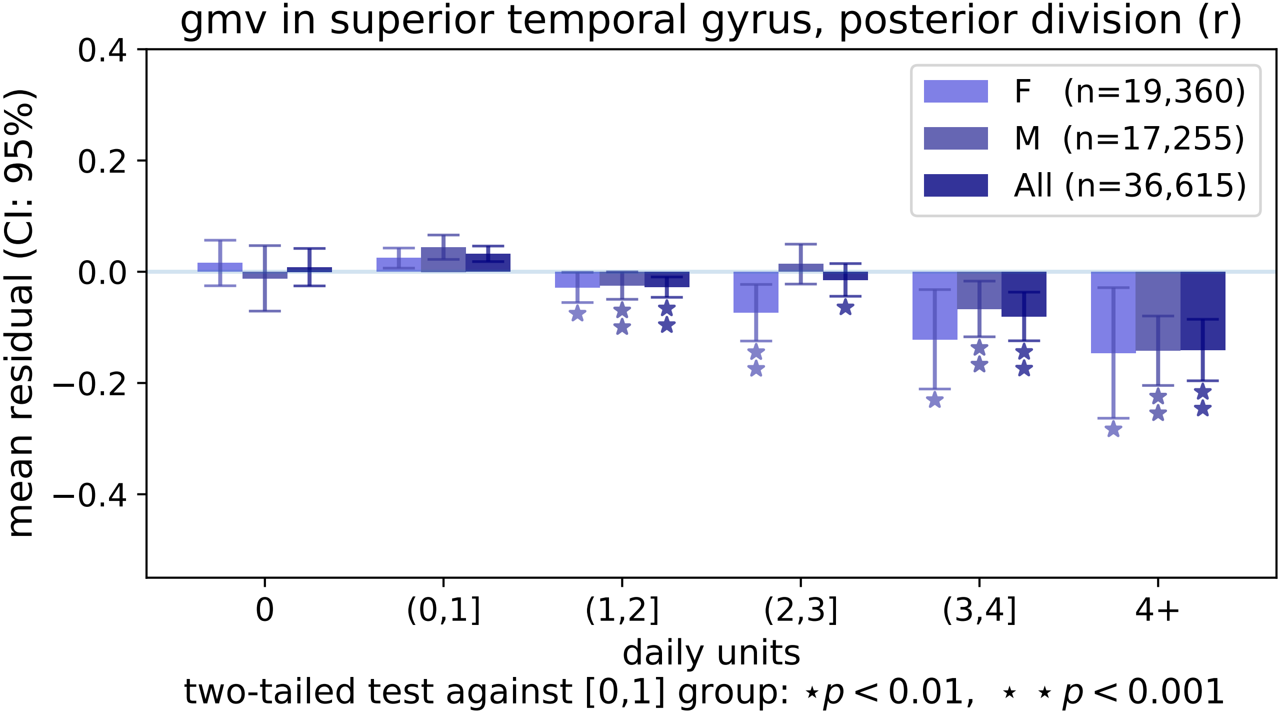
gmv in superior temporal gyrus, posterior division (r)
mean residual (CI: 95%)
0.4
0.2
0.0
−0.2
−0.4
F (n=19,360)
M (n=17,255)
All (n=36,615)
0
(0,1]
(1,2]
(2,3]
(3,4]
4+
daily units
two-tailed test against [0,1] group: ⋆p < 0.01, ⋆ ⋆ p < 0.001

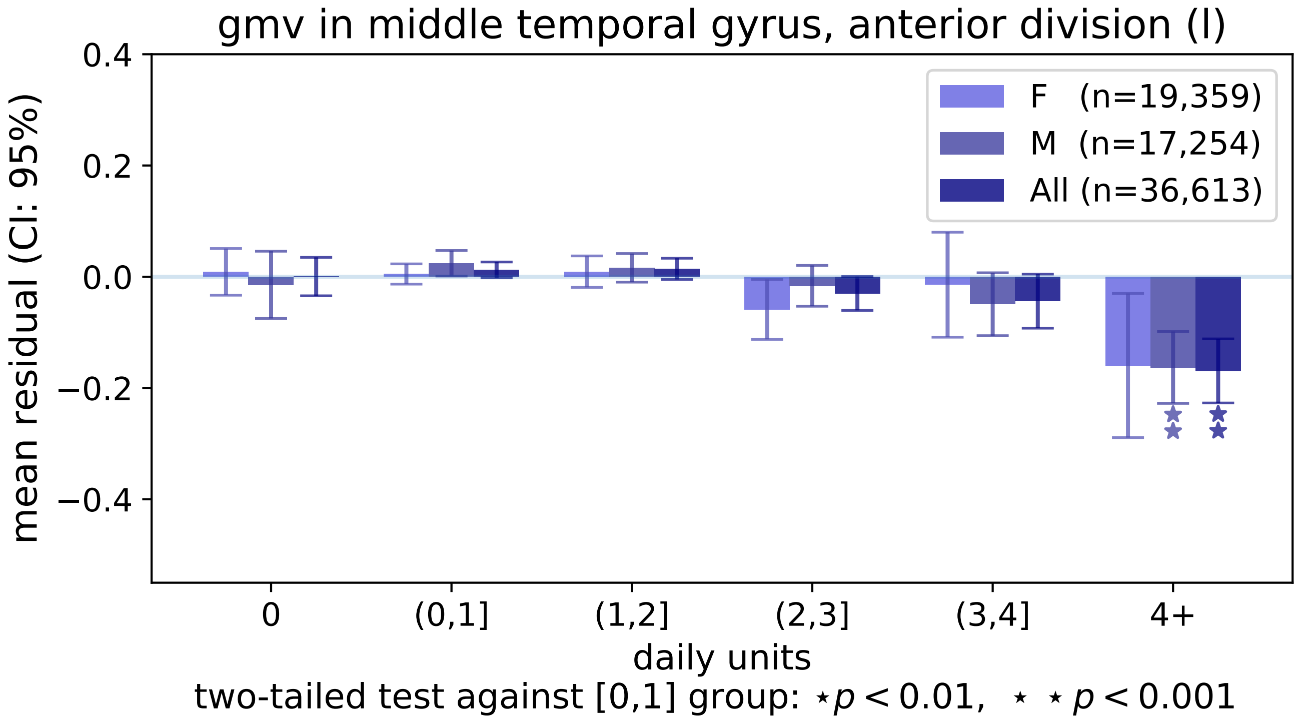

gmv in middle temporal gyrus, anterior division (l)
F (n=19,359)
M (n=17,254)
All (n=36,613)
mean residual (CI: 95%)
0.4
0.2
0.0
−0.2
−0.4
0
(0,1]
(1,2]
(2,3]
(3,4]
4+
daily units
two-tailed test against [0,1] group: ⋆p < 0.01, ⋆ ⋆ p < 0.001

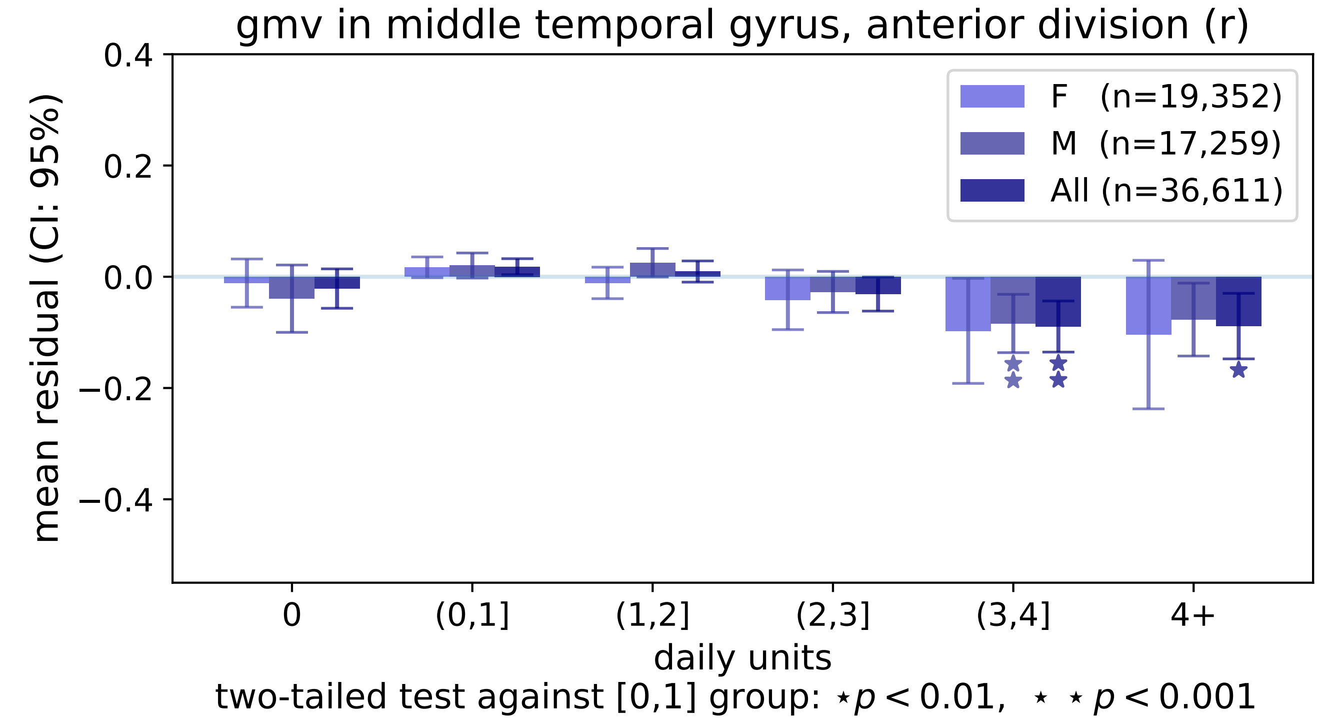
gmv in middle temporal gyrus, anterior division (r)
mean residual (CI: 95%)
0.4
0.2
0.0
−0.2
−0.4
F (n=19,352)
M (n=17,259)
All (n=36,611)
0
(0,1]
(1,2]
(2,3]
(3,4]
4+
daily units
two-tailed test against [0,1] group: ⋆ p < 0.01, ⋆ ⋆ p < 0.001

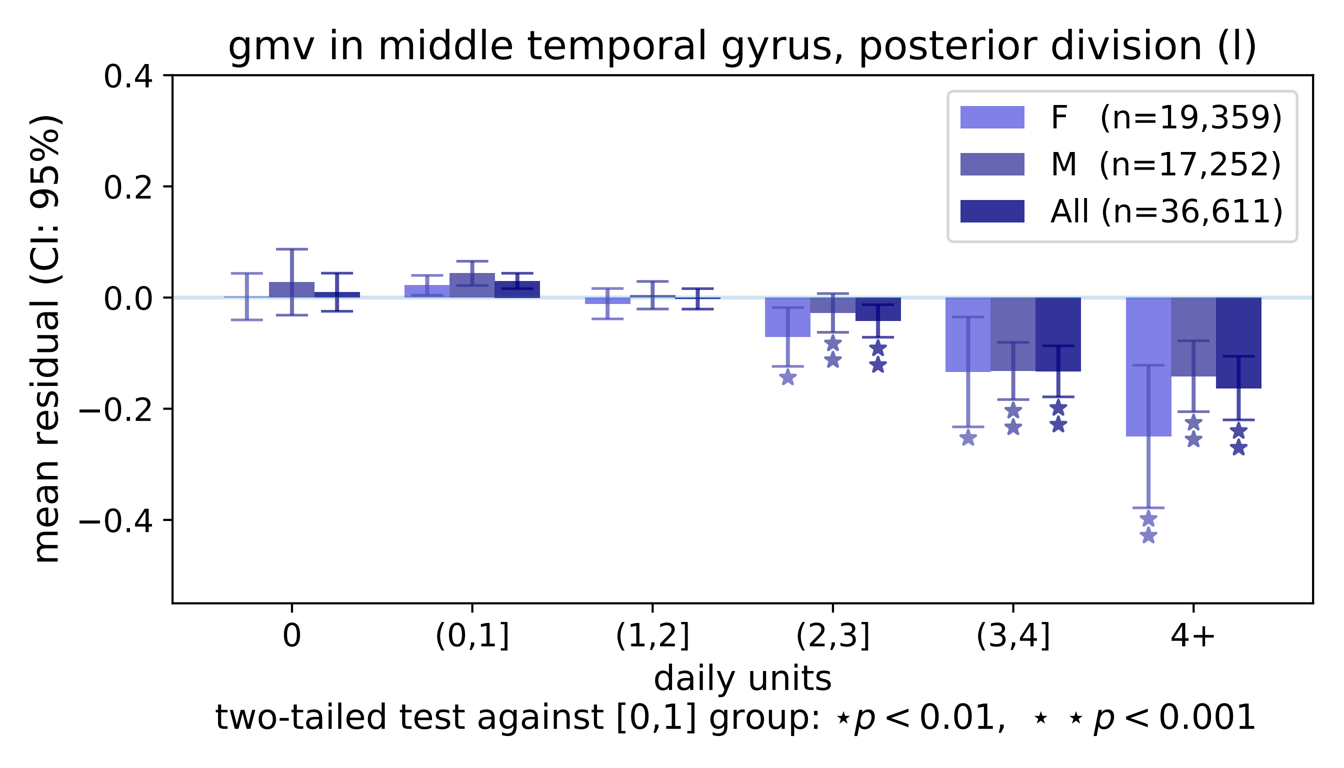
gmv in middle temporal gyrus, posterior division (l)
mean residual (CI: 95%)
0.4
0.2
0.0
−0.2
−0.4
F (n=19,359)
M (n=17,252)
All (n=36,611)
0
(0,1]
(1,2]
(2,3]
(3,4]
4+
daily units
two-tailed test against [0,1] group: ⋆p < 0.01, ⋆ ⋆ p < 0.001

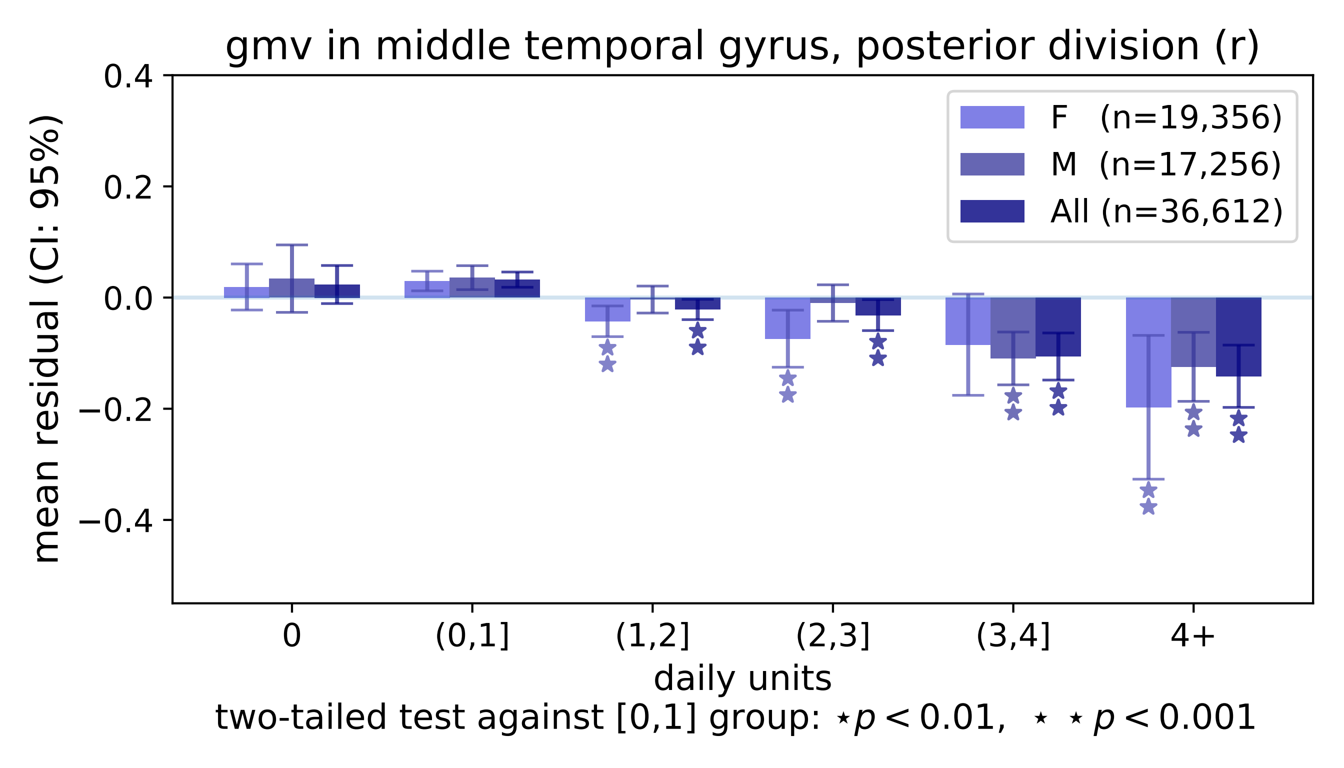
gmv in middle temporal gyrus, posterior division (r)
mean residual (CI: 95%)
0.4
0.2
0.0
−0.2
−0.4
F (n=19,356)
M (n=17,256)
All (n=36,612)
0
(0,1]
(1,2]
(2,3]
(3,4]
4+
daily units
two-tailed test against [0,1] group: ⋆$p < 0.01$, ⋆ ⋆ $p < 0.001$

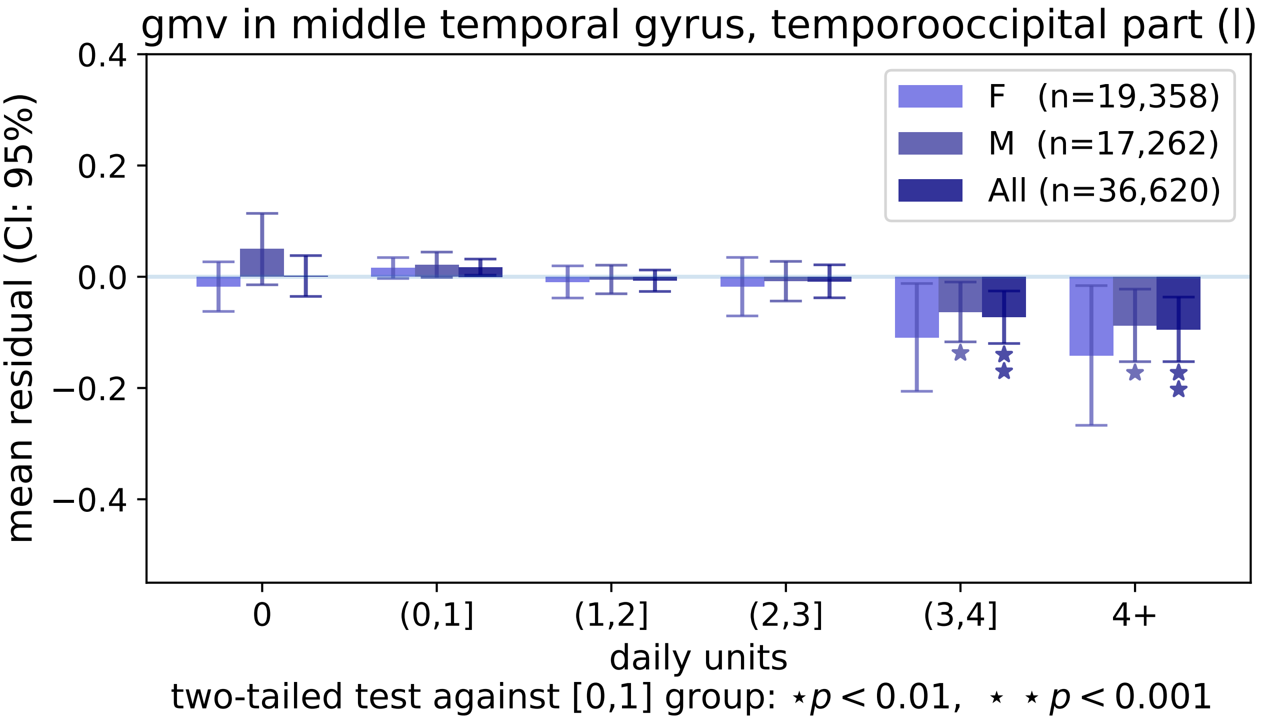

gmv in middle temporal gyrus, temporooccipital part (l)
mean residual (CI: 95%)
0.4
0.2
0.0
−0.2
−0.4
F (n=19,358)
M (n=17,262)
All (n=36,620)
0
(0,1]
(1,2]
(2,3]
(3,4]
4+
daily units
two-tailed test against [0,1] group: ⋆p < 0.01, ⋆ ⋆ p < 0.001

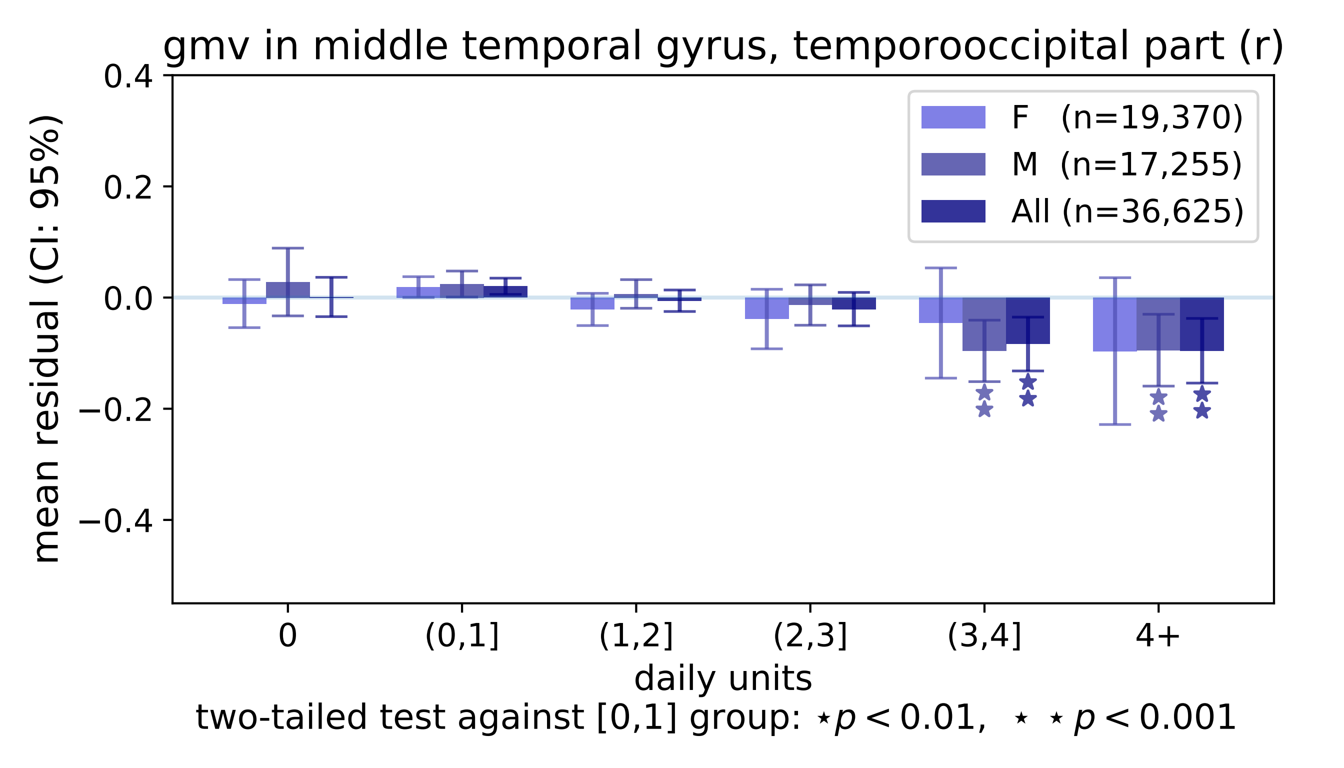

gmv in middle temporal gyrus, temporooccipital part (r)
mean residual (CI: 95%)
0.4
0.2
0.0
−0.2
−0.4
F (n=19,370)
M (n=17,255)
All (n=36,625)
0
(0,1]
(1,2]
(2,3]
(3,4]
4+
daily units
two-tailed test against [0,1] group: ⋆p < 0.01, ⋆ ⋆ p < 0.001

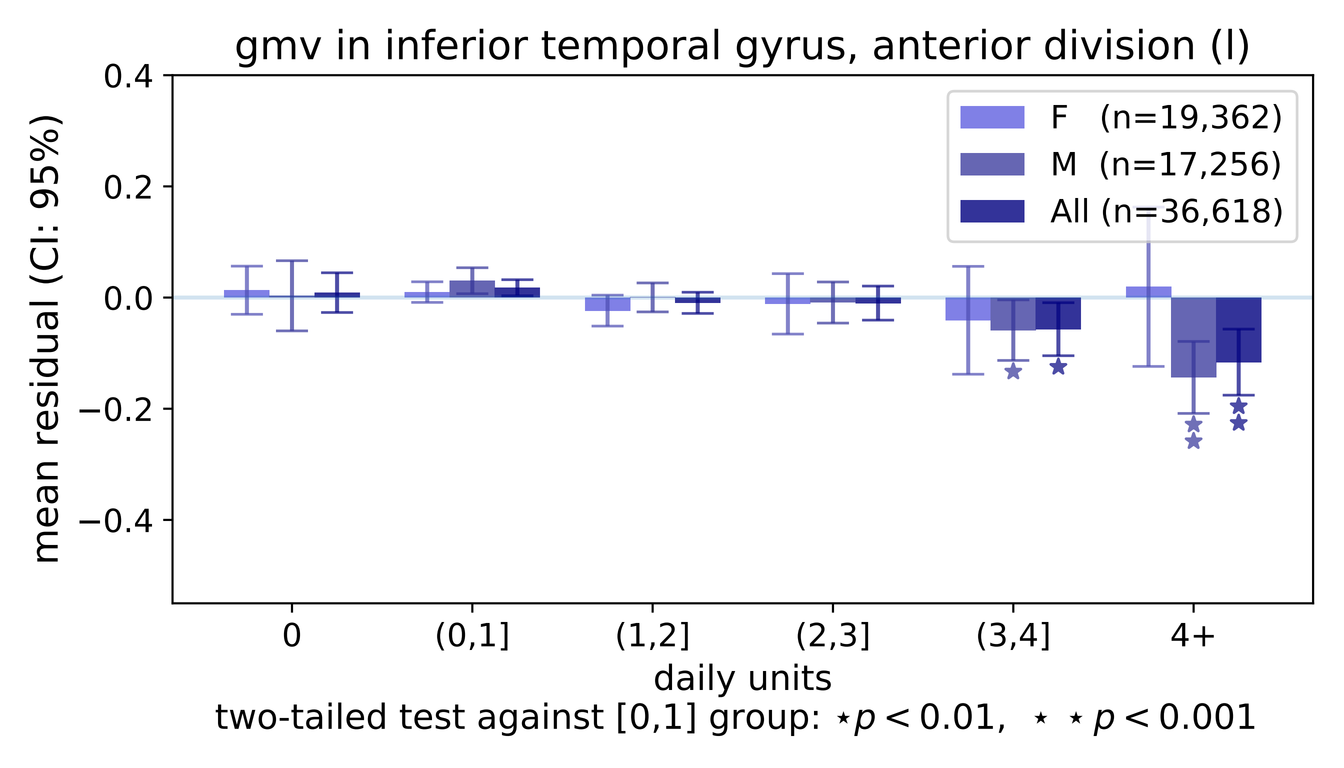

gmv in inferior temporal gyrus, anterior division (l)
mean residual (CI: 95%)
0.4
0.2
0.0
−0.2
−0.4
F (n=19,362)
M (n=17,256)
All (n=36,618)
0
(0,1]
(1,2]
(2,3]
(3,4]
4+
daily units
two-tailed test against [0,1] group: ⋆p < 0.01, ⋆ ⋆ p < 0.001

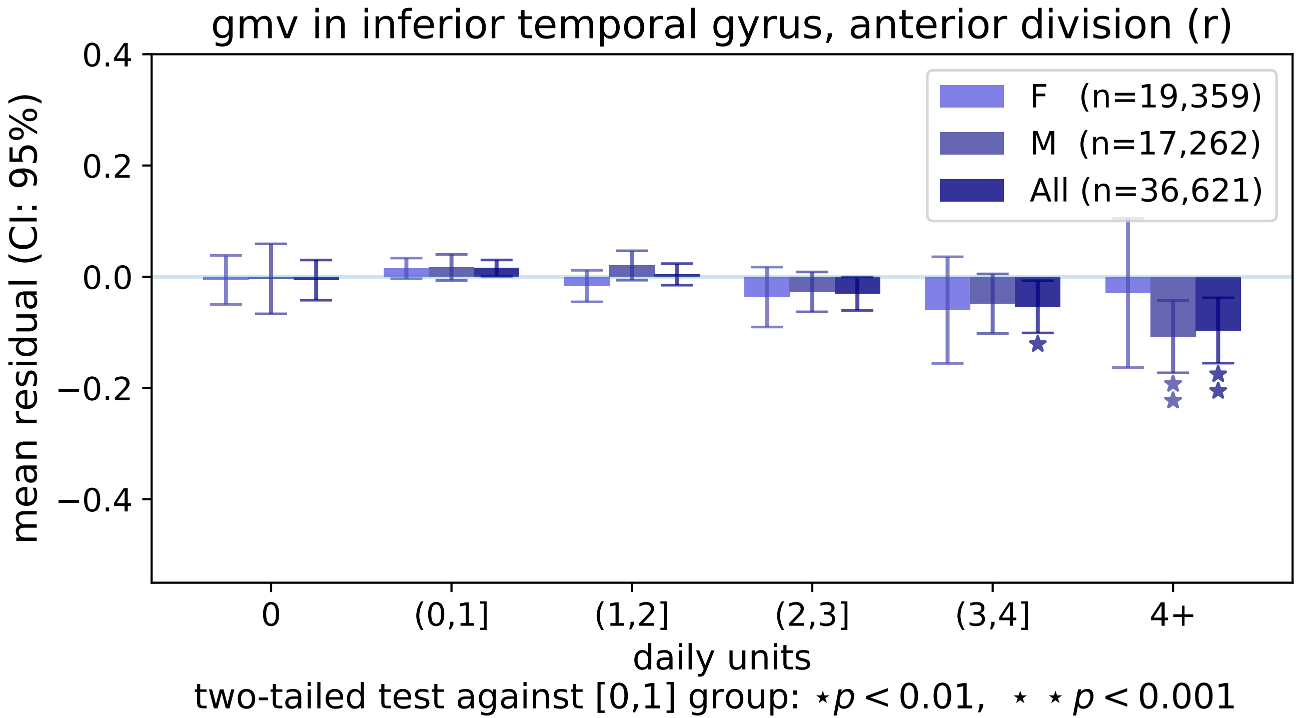
gmv in inferior temporal gyrus, anterior division (r)
0.4
0.2
0.0
−0.2
−0.4
mean residual (CI: 95%)
F (n=19,359)
M (n=17,262)
All (n=36,621)
0
(0,1]
(1,2]
(2,3]
(3,4]
4+
daily units
two-tailed test against [0,1] group: ⋆p < 0.01, ⋆ ⋆ p < 0.001

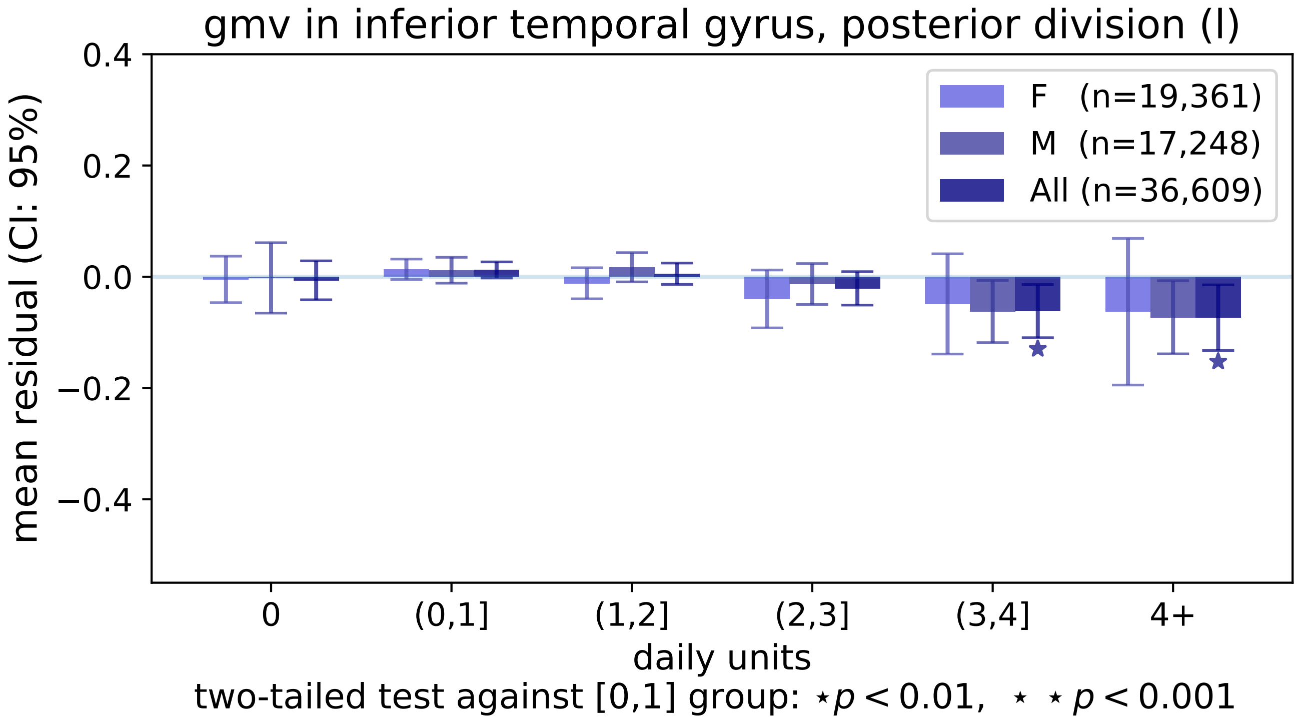

gmv in inferior temporal gyrus, posterior division (l)
mean residual (CI: 95%)
0.4
0.2
0.0
−0.2
−0.4
F (n=19,361)
M (n=17,248)
All (n=36,609)
0
(0,1]
(1,2]
(2,3]
(3,4]
4+
daily units
two-tailed test against [0,1] group: ⋆p < 0.01, ⋆ ⋆ p < 0.001

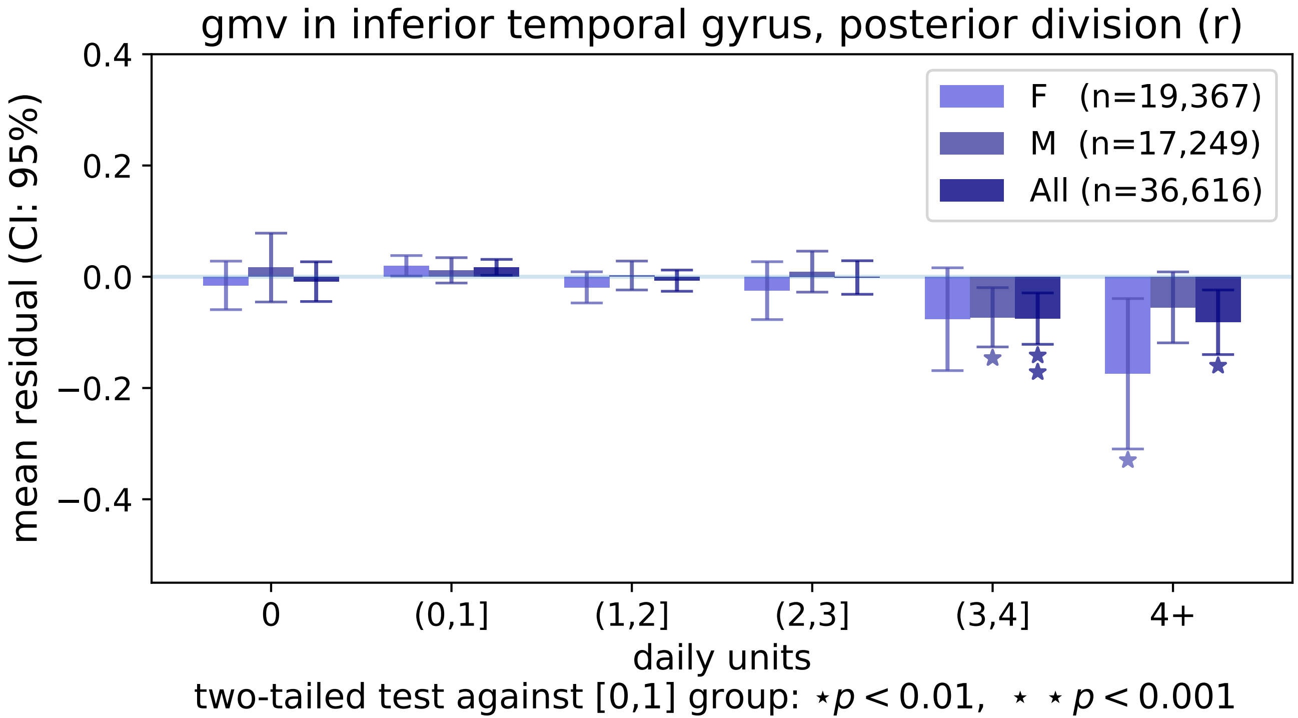

gmv in inferior temporal gyrus, posterior division (r)
mean residual (CI: 95%)
0.4
0.2
0.0
−0.2
−0.4
F (n=19,367)
M (n=17,249)
All (n=36,616)
0
(0,1]
(1,2]
(2,3]
(3,4]
4+
daily units
two-tailed test against [0,1] group: ⋆p < 0.01, ⋆ ⋆ p < 0.001

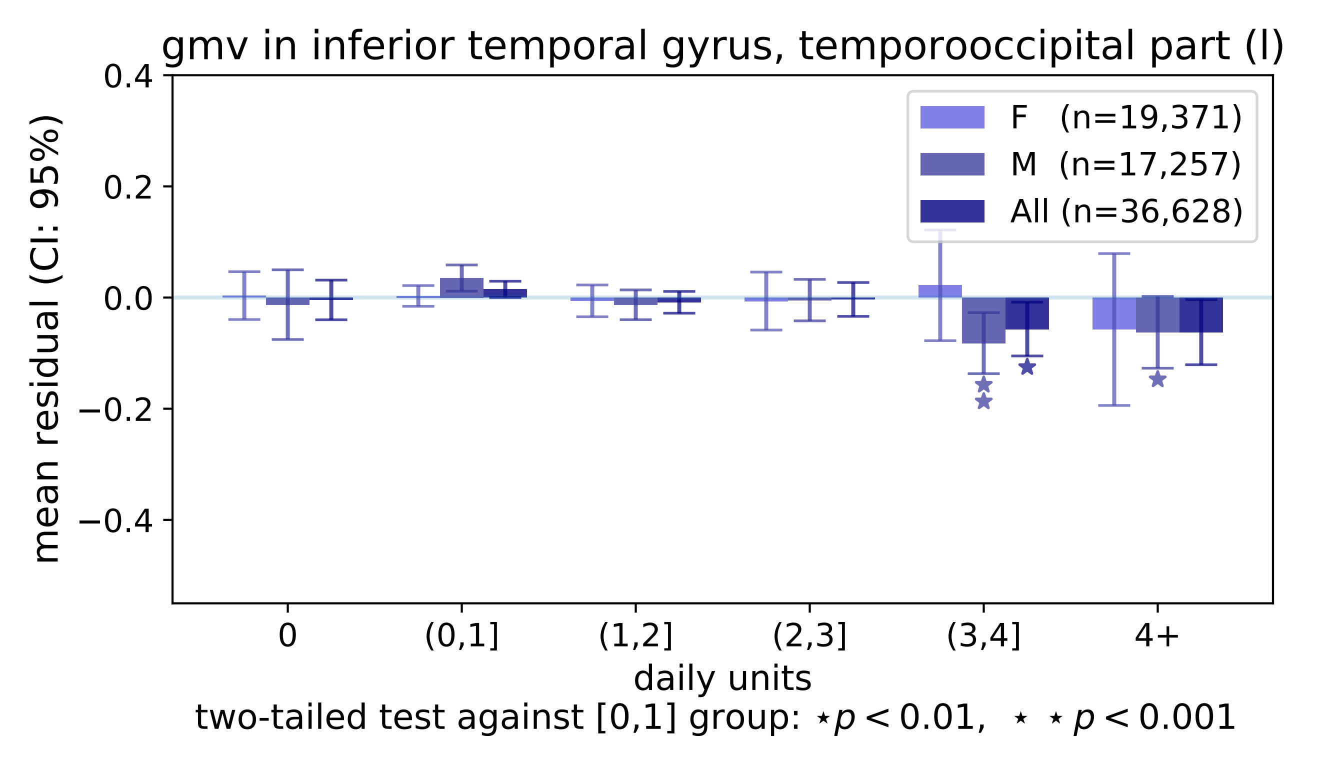

gmv in inferior temporal gyrus, temporooccipital part (l)
mean residual (CI: 95%)
0.4
0.2
0.0
−0.2
−0.4
F (n=19,371)
M (n=17,257)
All (n=36,628)
0
(0,1]
(1,2]
(2,3]
(3,4]
4+
daily units
two-tailed test against [0,1] group: ⋆p < 0.01, ⋆ ⋆p < 0.001

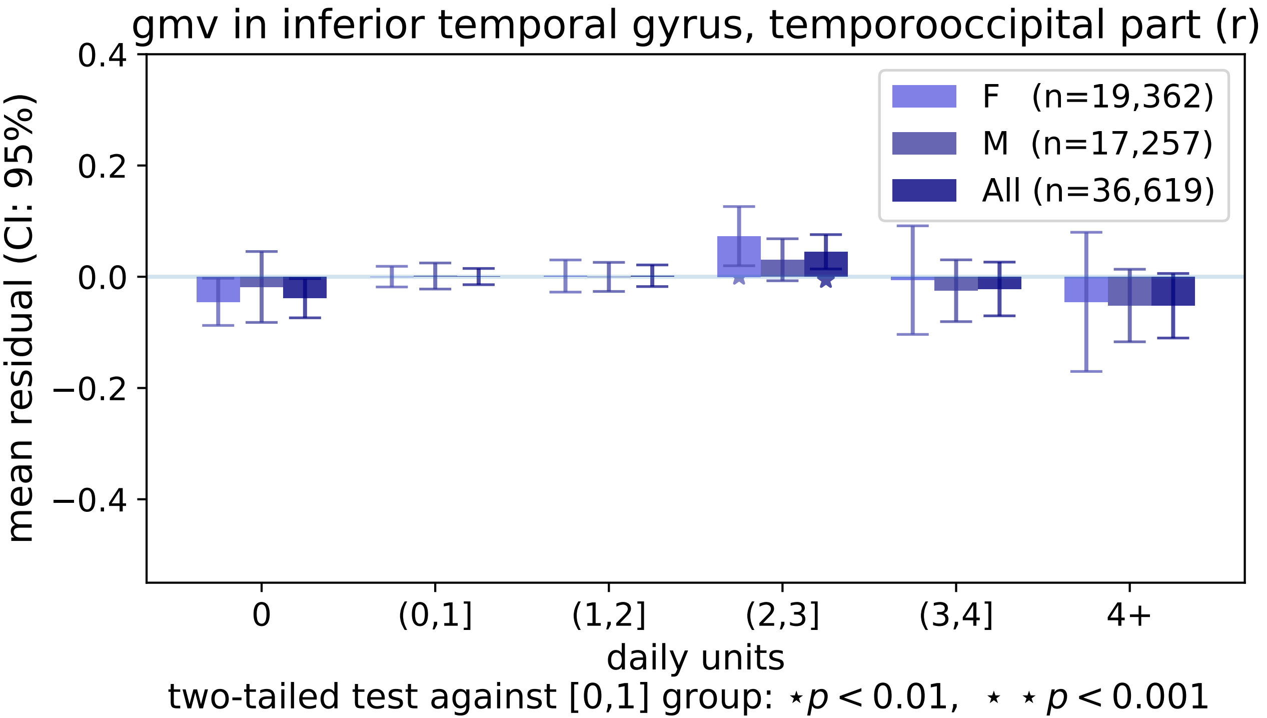
gmv in inferior temporal gyrus, temporooccipital part (r)
mean residual (CI: 95%)
0.4
0.2
0.0
−0.2
−0.4
F (n=19,362)
M (n=17,257)
All (n=36,619)
0
(0,1]
(1,2]
(2,3]
(3,4]
4+
daily units
two-tailed test against [0,1] group: ⋆p < 0.01, ⋆ ⋆p < 0.001

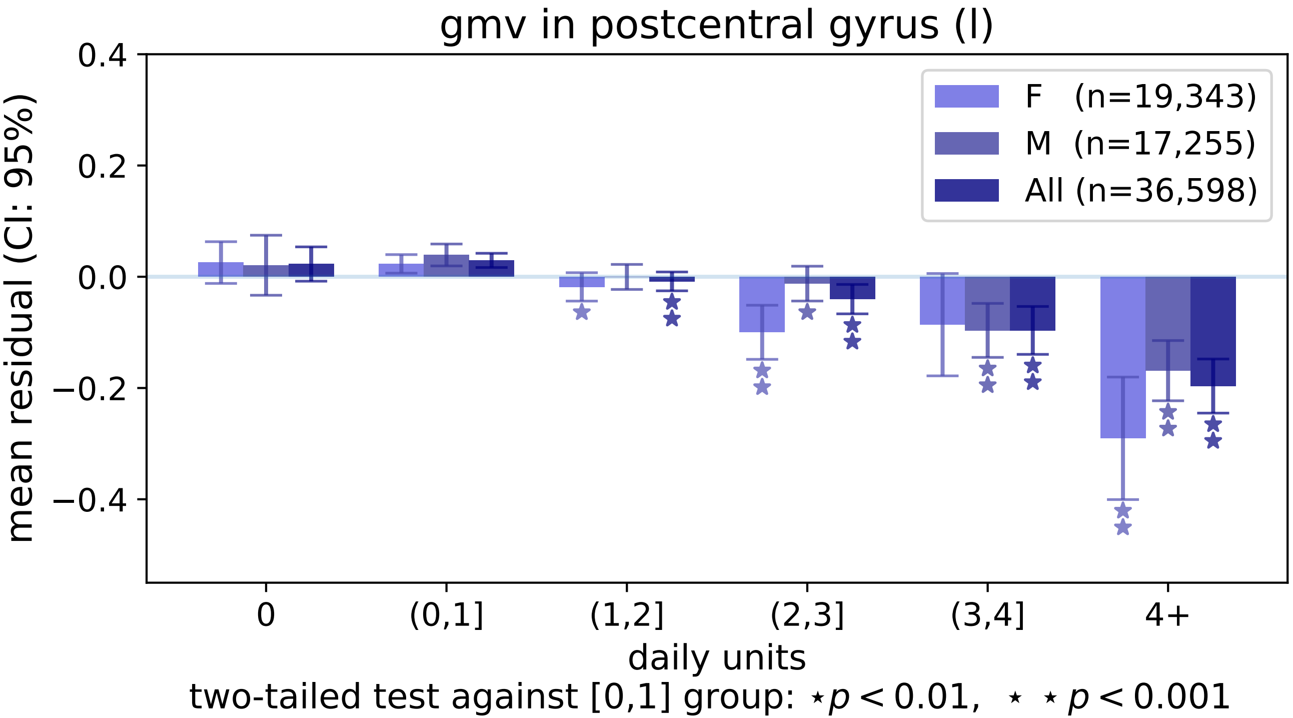

gmv in postcentral gyrus (l)
0.4
0.2
0.0
−0.2
−0.4
mean residual (CI: 95%)
F (n=19,343)
M (n=17,255)
All (n=36,598)
0
(0,1]
(1,2]
(2,3]
(3,4]
4+
daily units
two-tailed test against [0,1] group: ⋆ p < 0.01, ⋆ ⋆ p < 0.001

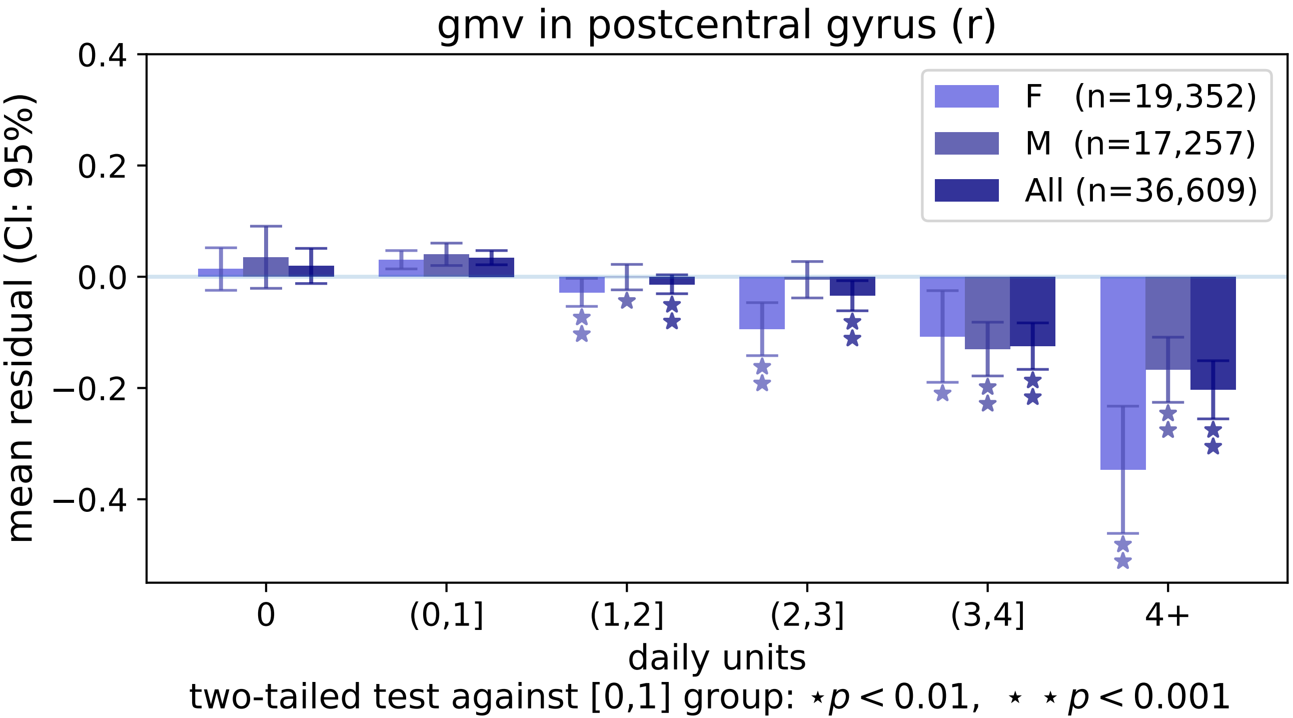
gmv in postcentral gyrus (r)
mean residual (CI: 95%)
0.4
0.2
0.0
−0.2
−0.4
F (n=19,352)
M (n=17,257)
All (n=36,609)
0
(0,1]
(1,2]
(2,3]
(3,4]
4+
daily units
two-tailed test against [0,1] group: ⋆p < 0.01, ⋆ ⋆ p < 0.001

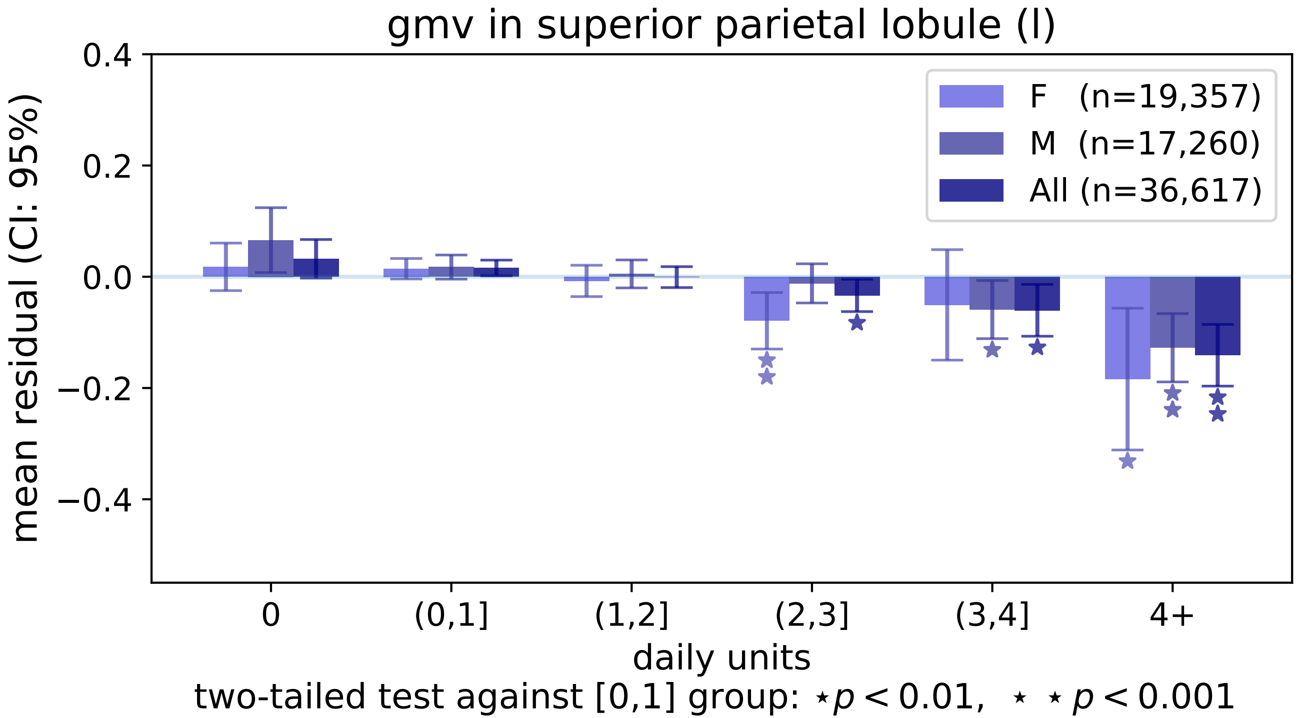

gmv in superior parietal lobule (l)
mean residual (CI: 95%)
0.4
0.2
0.0
−0.2
−0.4
F (n=19,357)
M (n=17,260)
All (n=36,617)
0
(0,1]
(1,2]
(2,3]
(3,4]
4+
daily units
two-tailed test against [0,1] group: ⋆ $p < 0.01$, ⋆ ⋆ $p < 0.001$

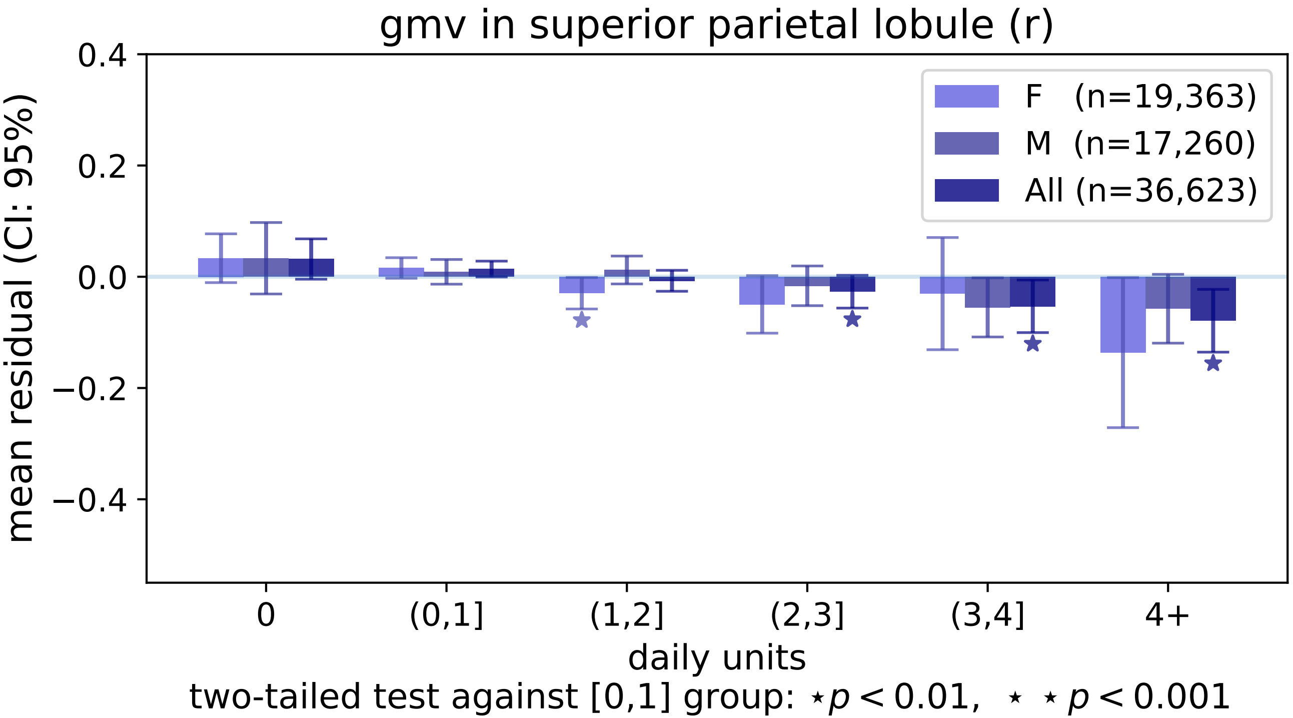
gmv in superior parietal lobule (r)
F (n=19,363)
M (n=17,260)
All (n=36,623)
mean residual (CI: 95%)
0.4
0.2
0.0
−0.2
−0.4
0
(0,1]
(1,2]
(2,3]
(3,4]
4+
daily units
two-tailed test against [0,1] group: ⋆ $p < 0.01$, ⋆ ⋆ $p < 0.001$

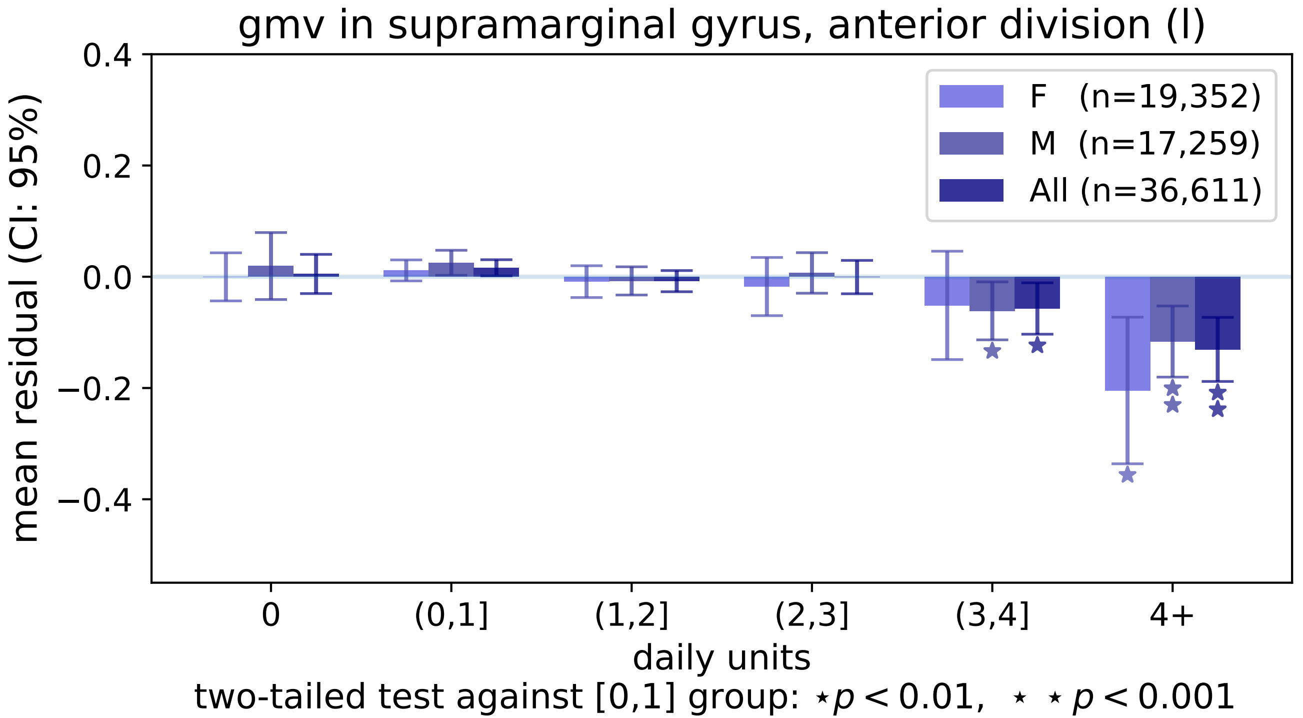
gmv in supramarginal gyrus, anterior division (l)
mean residual (CI: 95%)
0.4
0.2
0.0
−0.2
−0.4
F (n=19,352)
M (n=17,259)
All (n=36,611)
0
(0,1]
(1,2]
(2,3]
(3,4]
4+
daily units
two-tailed test against [0,1] group: ⋆ $p < 0.01$, ⋆ ⋆ $p < 0.001$

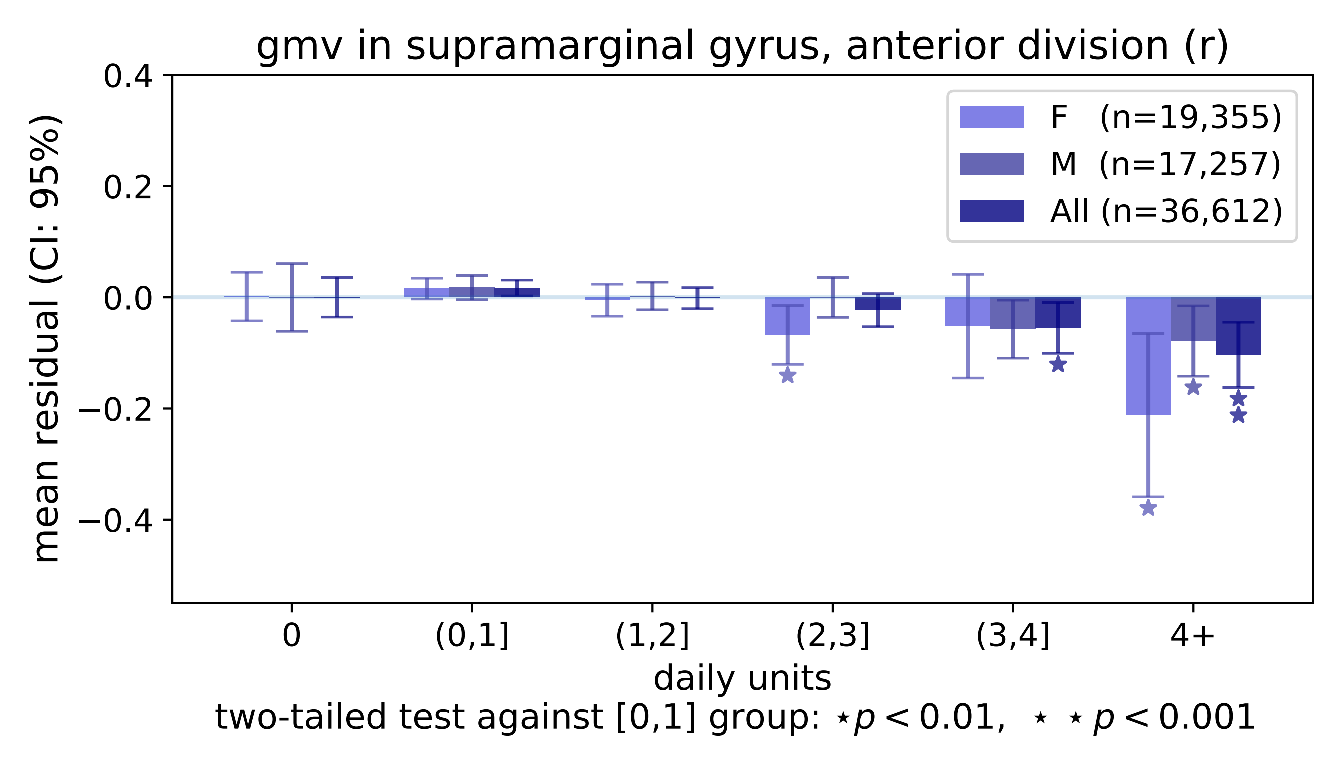

gmv in supramarginal gyrus, anterior division (r)
mean residual (CI: 95%)
0.4
0.2
0.0
−0.2
−0.4
F (n=19,355)
M (n=17,257)
All (n=36,612)
0
(0,1]
(1,2]
(2,3]
(3,4]
4+
daily units
two-tailed test against [0,1] group: ⋆p < 0.01, ⋆ ⋆ p < 0.001

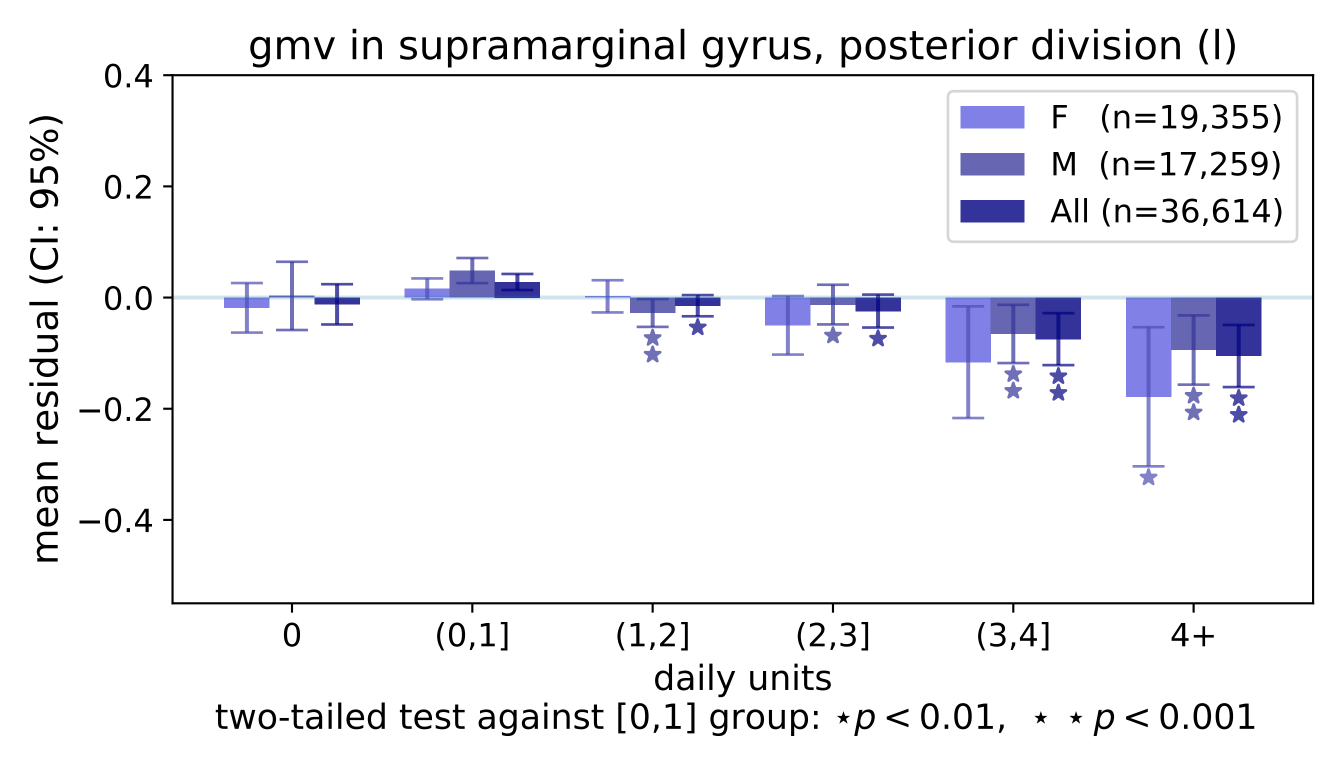
gmv in supramarginal gyrus, posterior division (l)
mean residual (CI: 95%)
0.4
0.2
0.0
−0.2
−0.4
F (n=19,355)
M (n=17,259)
All (n=36,614)
0
(0,1]
(1,2]
(2,3]
(3,4]
4+
daily units
two-tailed test against [0,1] group: ⋆p < 0.01, ⋆ ⋆ p < 0.001

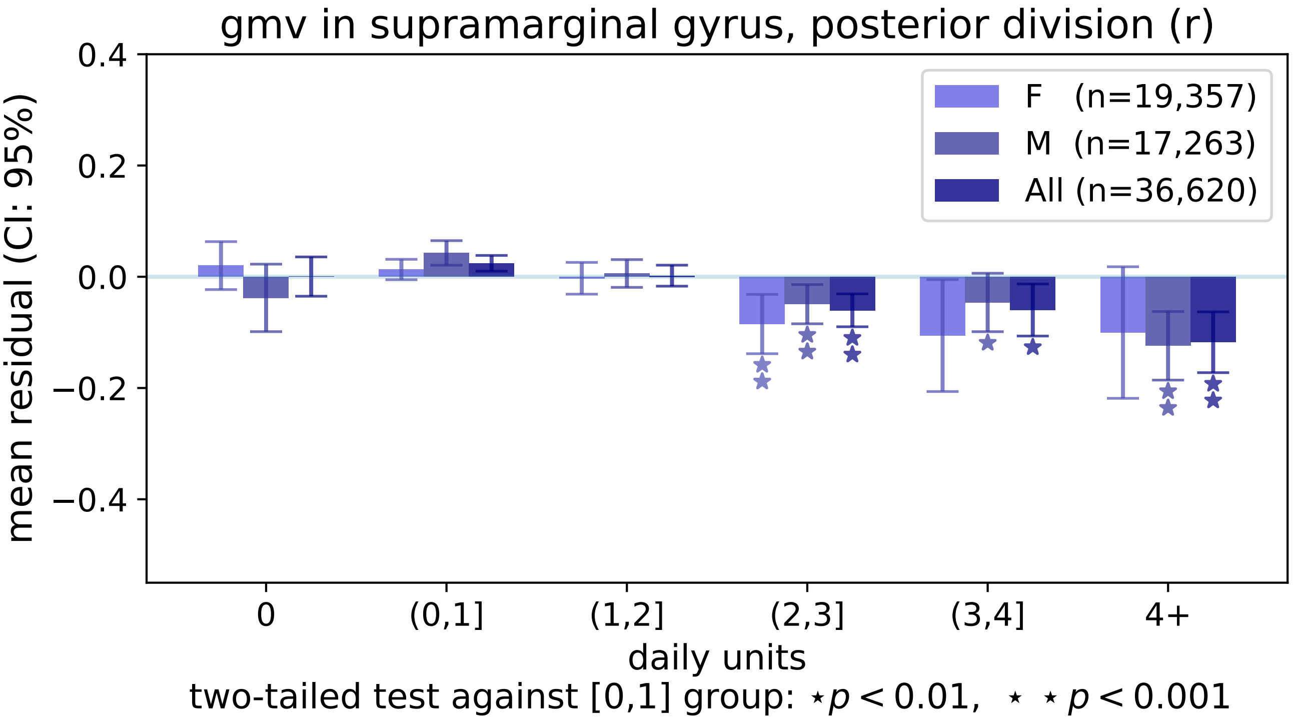
gmv in supramarginal gyrus, posterior division (r)
F (n=19,357)
M (n=17,263)
All (n=36,620)
0.4
0.2
0.0
−0.2
−0.4
mean residual (CI: 95%)
0
(0,1]
(1,2]
(2,3]
(3,4]
4+
daily units
two-tailed test against [0,1] group: ⋆p < 0.01, ⋆ ⋆ p < 0.001

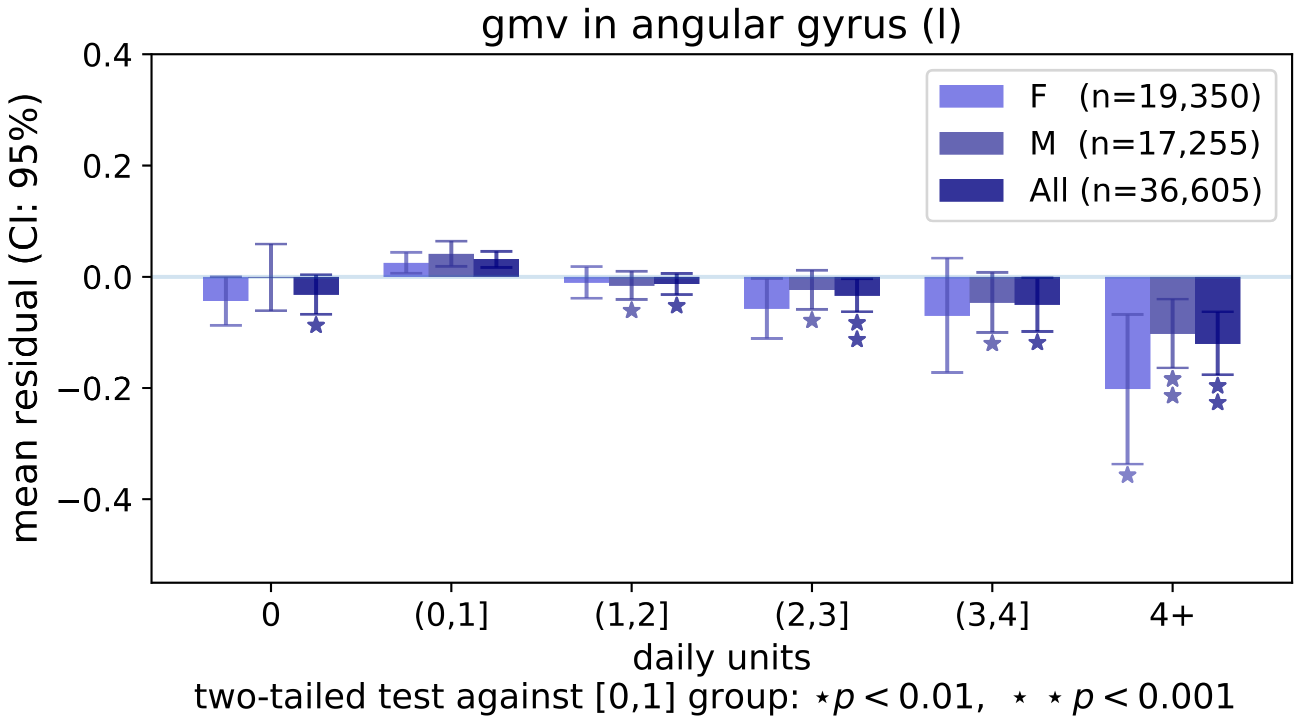

gmv in angular gyrus (l)
0.4
0.2
0.0
−0.2
−0.4
mean residual (CI: 95%)
F (n=19,350)
M (n=17,255)
All (n=36,605)
0
(0,1]
(1,2]
(2,3]
(3,4]
4+
daily units
two-tailed test against [0,1] group: ⋆ p < 0.01, ⋆ ⋆ p < 0.001

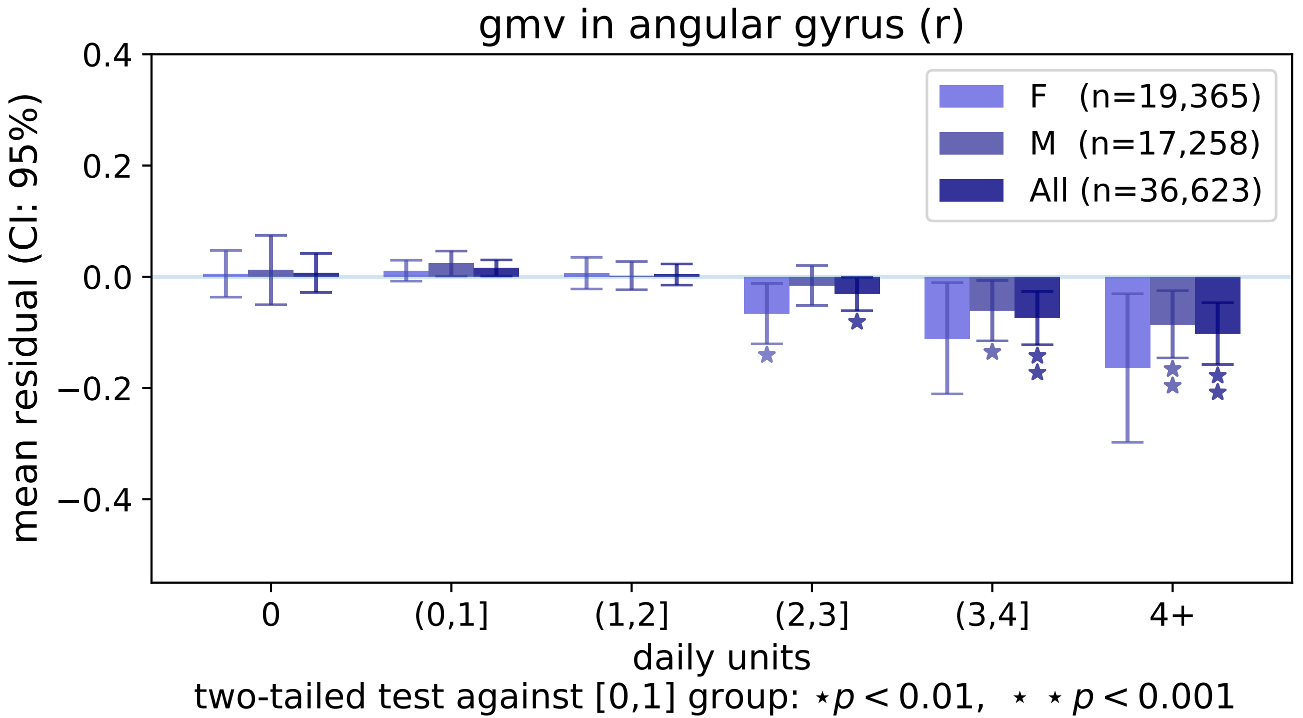

gmv in angular gyrus (r)
F (n=19,365)
M (n=17,258)
All (n=36,623)
mean residual (CI: 95%)
0.4
0.2
0.0
−0.2
−0.4
0
(0,1]
(1,2]
(2,3]
(3,4]
4+
daily units
two-tailed test against [0,1] group: ⋆ p < 0.01, ⋆ ⋆ p < 0.001

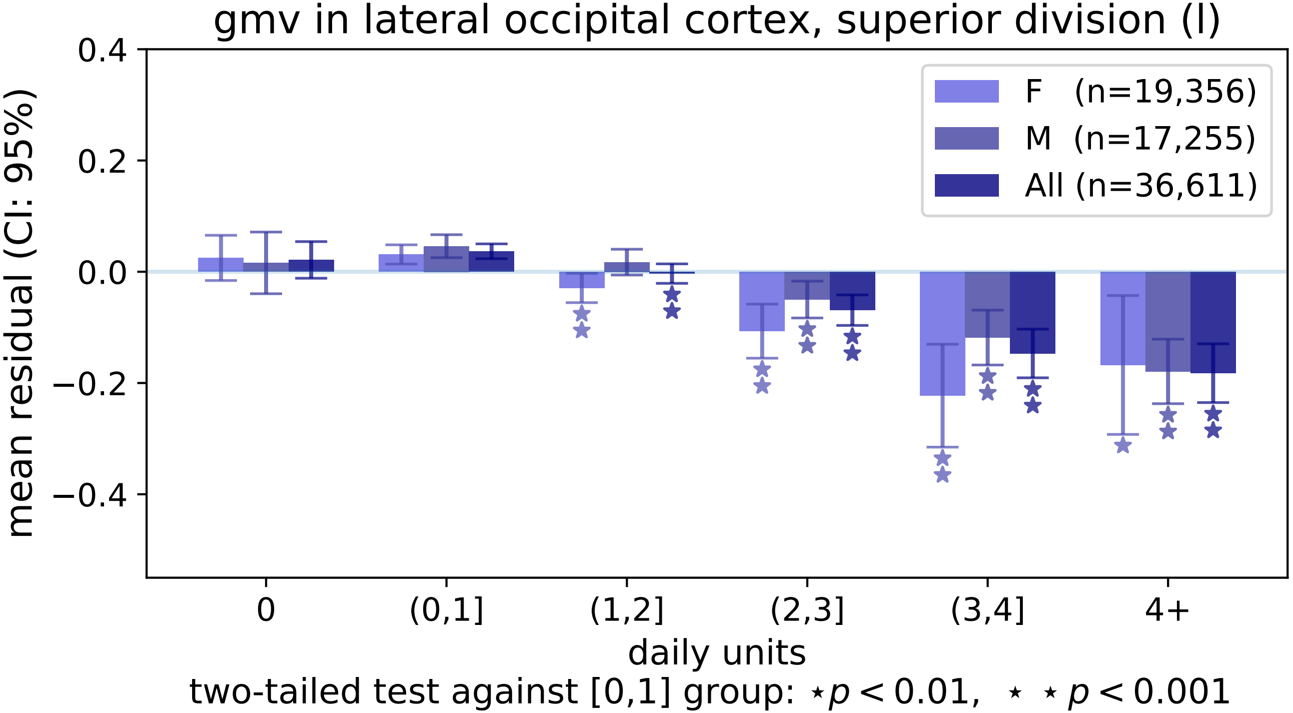

gmv in lateral occipital cortex, superior division (l)
mean residual (CI: 95%)
0.4
0.2
0.0
−0.2
−0.4
F (n=19,356)
M (n=17,255)
All (n=36,611)
0
(0,1]
(1,2]
(2,3]
(3,4]
4+
daily units
two-tailed test against [0,1] group: ⋆p < 0.01, ⋆ ⋆ p < 0.001

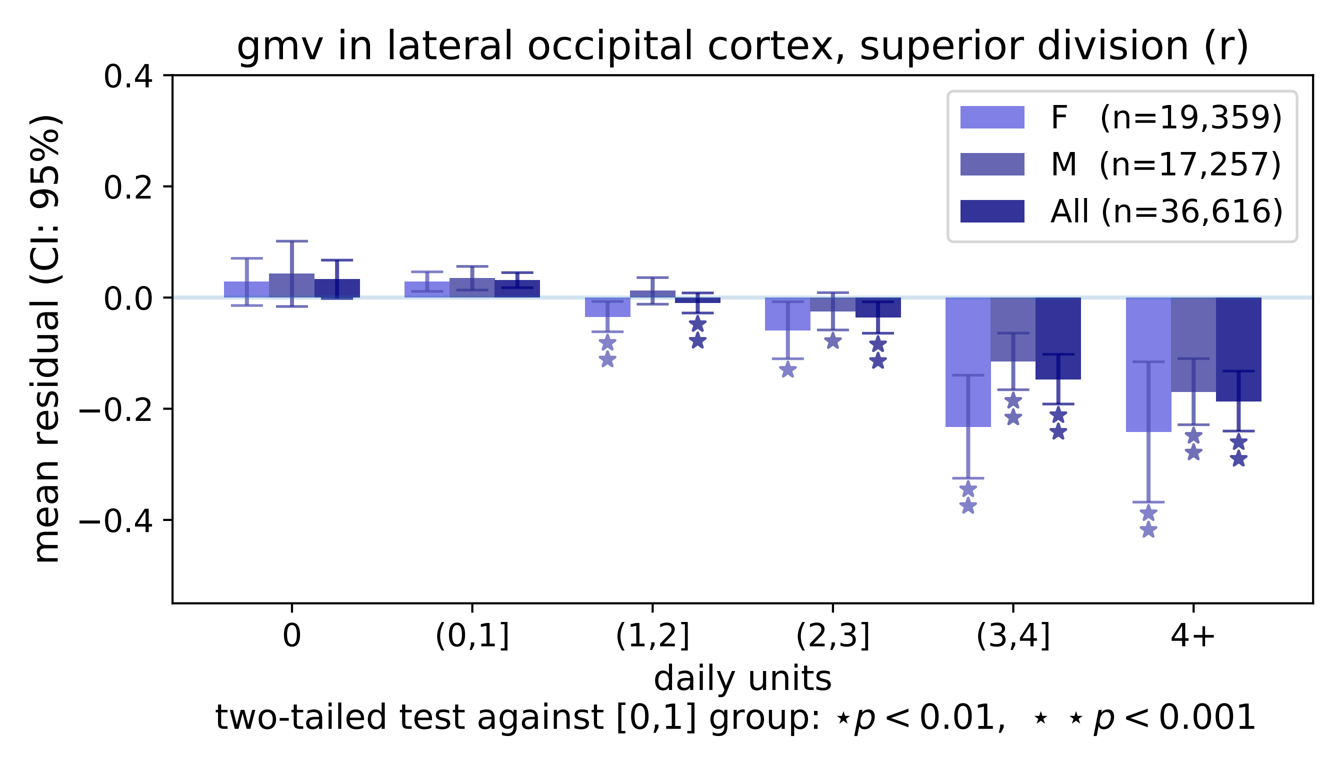
gmv in lateral occipital cortex, superior division (r)
mean residual (CI: 95%)
0.4
0.2
0.0
−0.2
−0.4
F (n=19,359)
M (n=17,257)
All (n=36,616)
0
(0,1]
(1,2]
(2,3]
(3,4]
4+
daily units
two-tailed test against [0,1] group: ⋆$p < 0.01$, ⋆ ⋆ $p < 0.001$

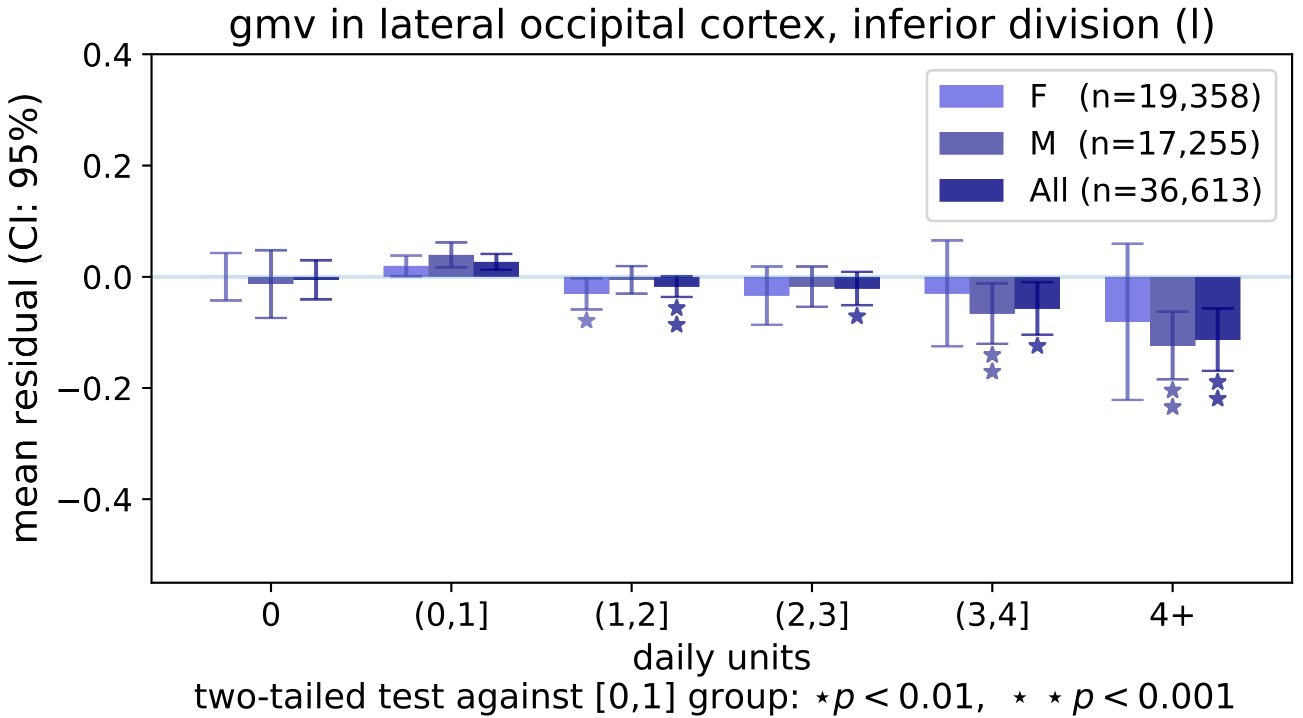
gmv in lateral occipital cortex, inferior division (l)
mean residual (CI: 95%)
0.4
0.2
0.0
−0.2
−0.4
F (n=19,358)
M (n=17,255)
All (n=36,613)
0
(0,1]
(1,2]
(2,3]
(3,4]
4+
daily units
two-tailed test against [0,1] group: ⋆p < 0.01, ⋆ ⋆ p < 0.001

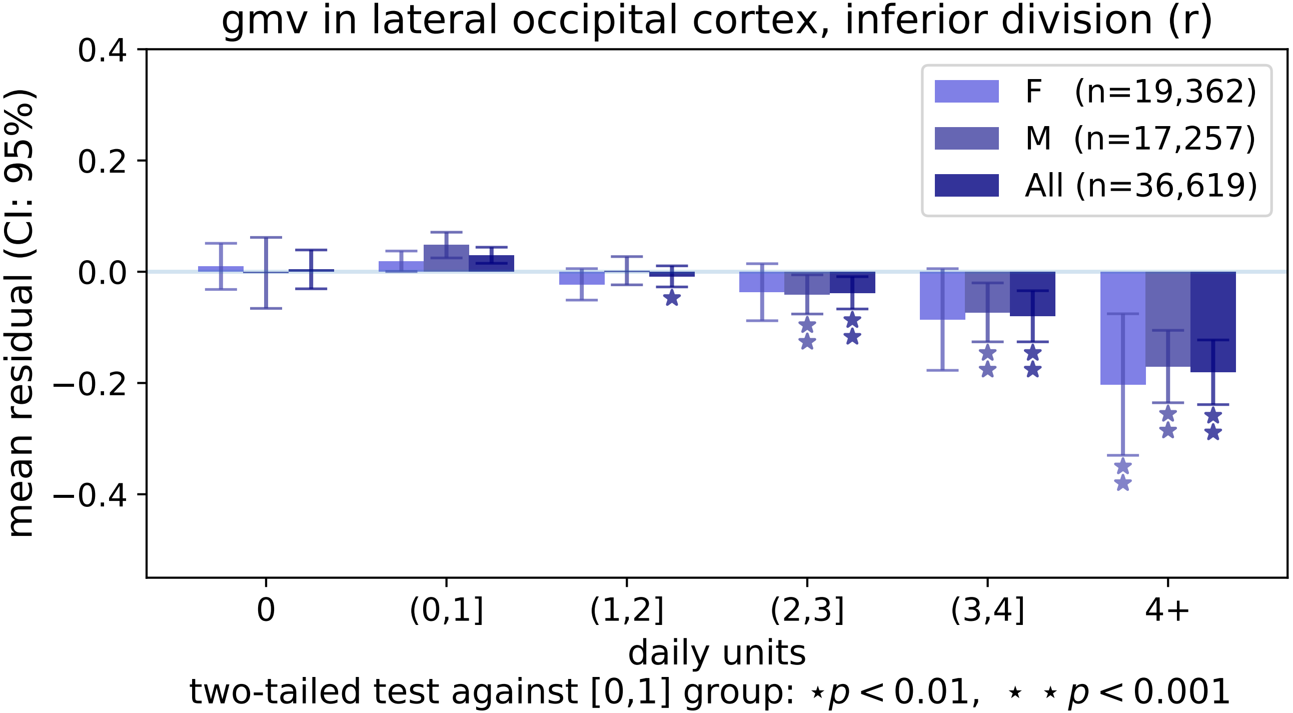
gmv in lateral occipital cortex, inferior division (r)
0.4
0.2
0.0
−0.2
−0.4
mean residual (CI: 95%)
F (n=19,362)
M (n=17,257)
All (n=36,619)
0
(0,1]
(1,2]
(2,3]
(3,4]
4+
daily units
two-tailed test against [0,1] group: ⋆ p < 0.01, ⋆ ⋆ p < 0.001

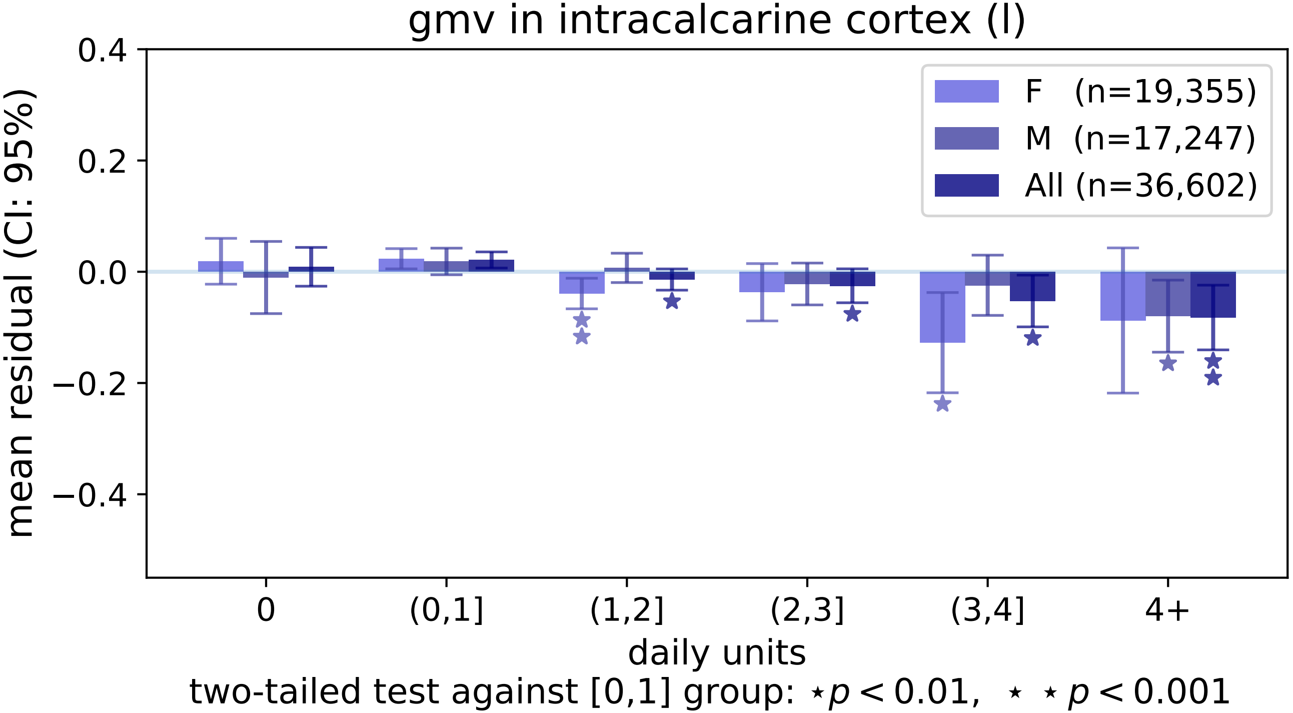
gmv in intracalcarine cortex (l)
mean residual (CI: 95%)
0.4
0.2
0.0
−0.2
−0.4
F (n=19,355)
M (n=17,247)
All (n=36,602)
0
(0,1]
(1,2]
(2,3]
(3,4]
4+
daily units
two-tailed test against [0,1] group: ⋆ $p < 0.01$, ⋆ ⋆ $p < 0.001$

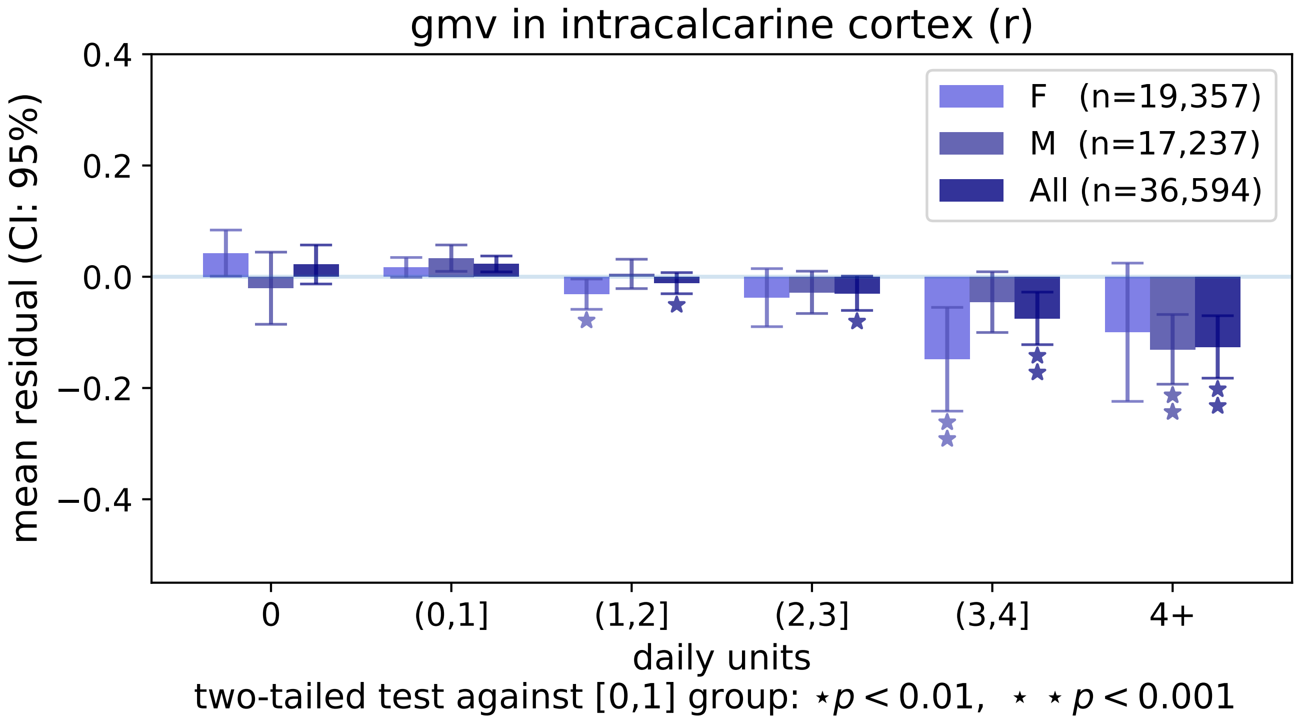
gmv in intracalcarine cortex (r)
mean residual (CI: 95%)
0.4
0.2
0.0
−0.2
−0.4
F (n=19,357)
M (n=17,237)
All (n=36,594)
0
(0,1]
(1,2]
(2,3]
(3,4]
4+
daily units
two-tailed test against [0,1] group: ⋆p < 0.01, ⋆ ⋆p < 0.001

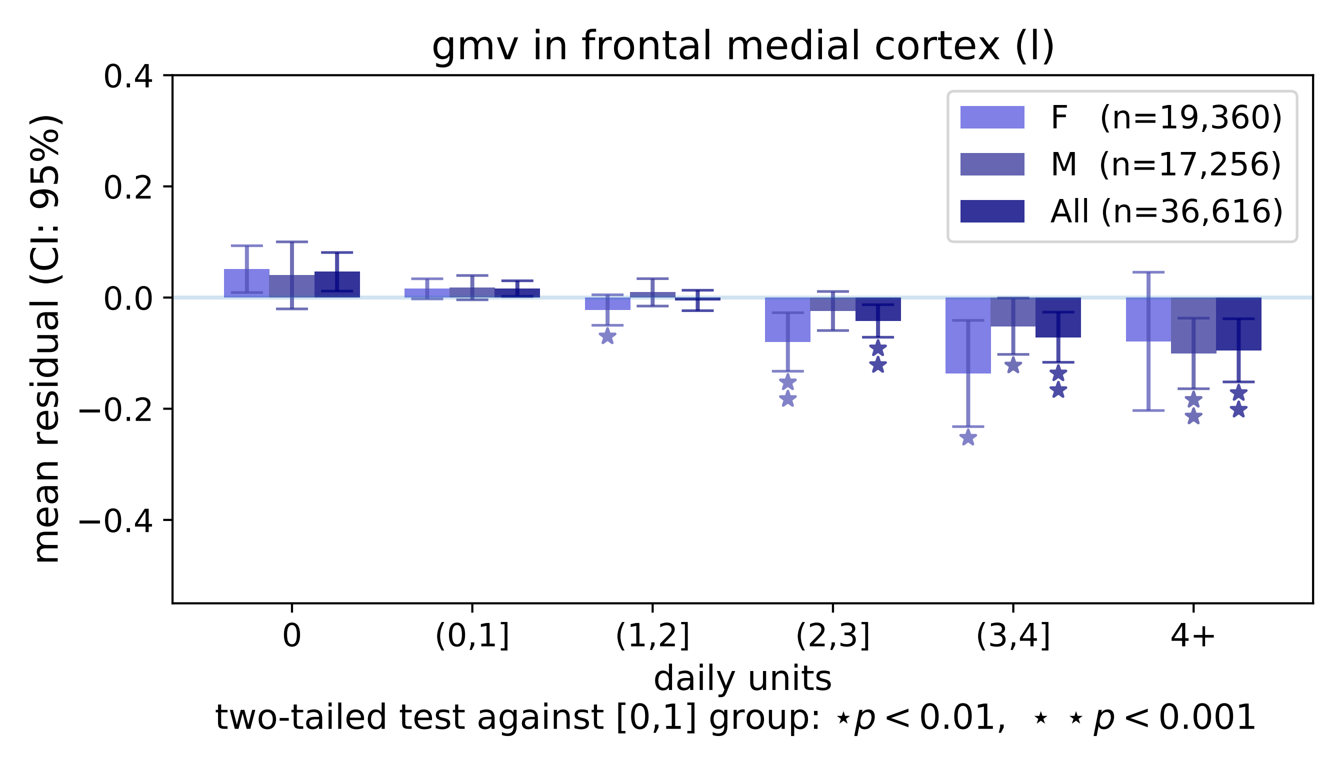

gmv in frontal medial cortex (l)
mean residual (CI: 95%)
0.4
0.2
0.0
−0.2
−0.4
F (n=19,360)
M (n=17,256)
All (n=36,616)
0
(0,1]
(1,2]
(2,3]
(3,4]
4+
daily units
two-tailed test against [0,1] group: ⋆p < 0.01, ⋆ ⋆ p < 0.001

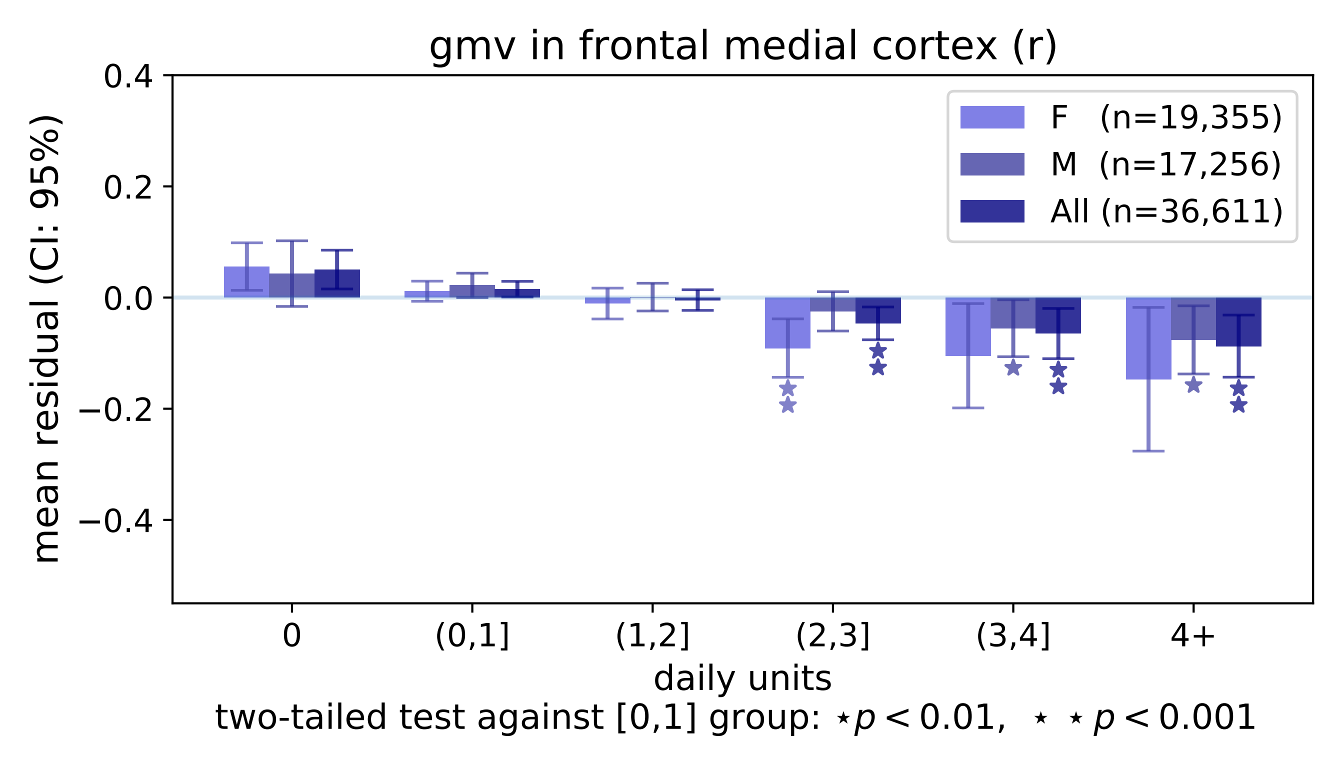
gmv in frontal medial cortex (r)
0.4
0.2
0.0
−0.2
−0.4
mean residual (CI: 95%)
F (n=19,355)
M (n=17,256)
All (n=36,611)
0
(0,1]
(1,2]
(2,3]
(3,4]
4+
daily units
two-tailed test against [0,1] group: ⋆ $p < 0.01$, ⋆ ⋆ $p < 0.001$

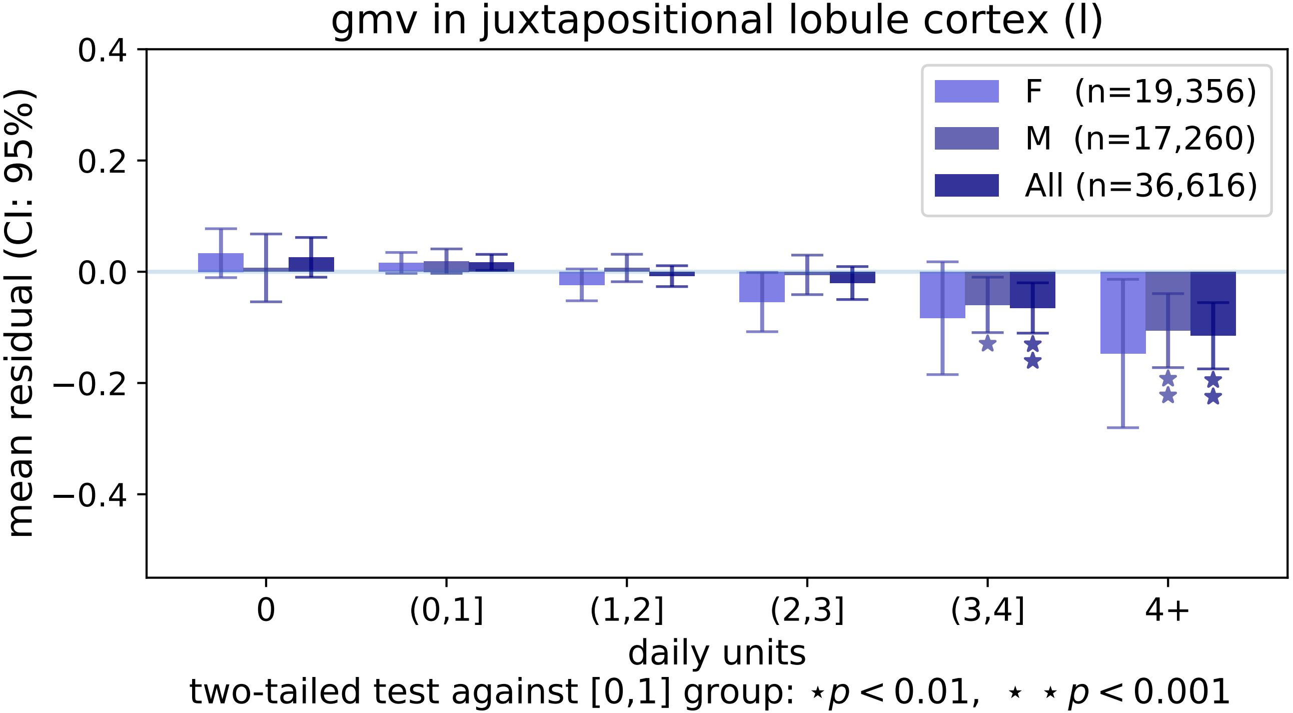
gmv in juxtapositional lobule cortex (l)
0.4
0.2
0.0
−0.2
−0.4
mean residual (CI: 95%)
F (n=19,356)
M (n=17,260)
All (n=36,616)
0
(0,1]
(1,2]
(2,3]
(3,4]
4+
daily units
two-tailed test against [0,1] group: ⋆p < 0.01, ⋆ ⋆ p < 0.001

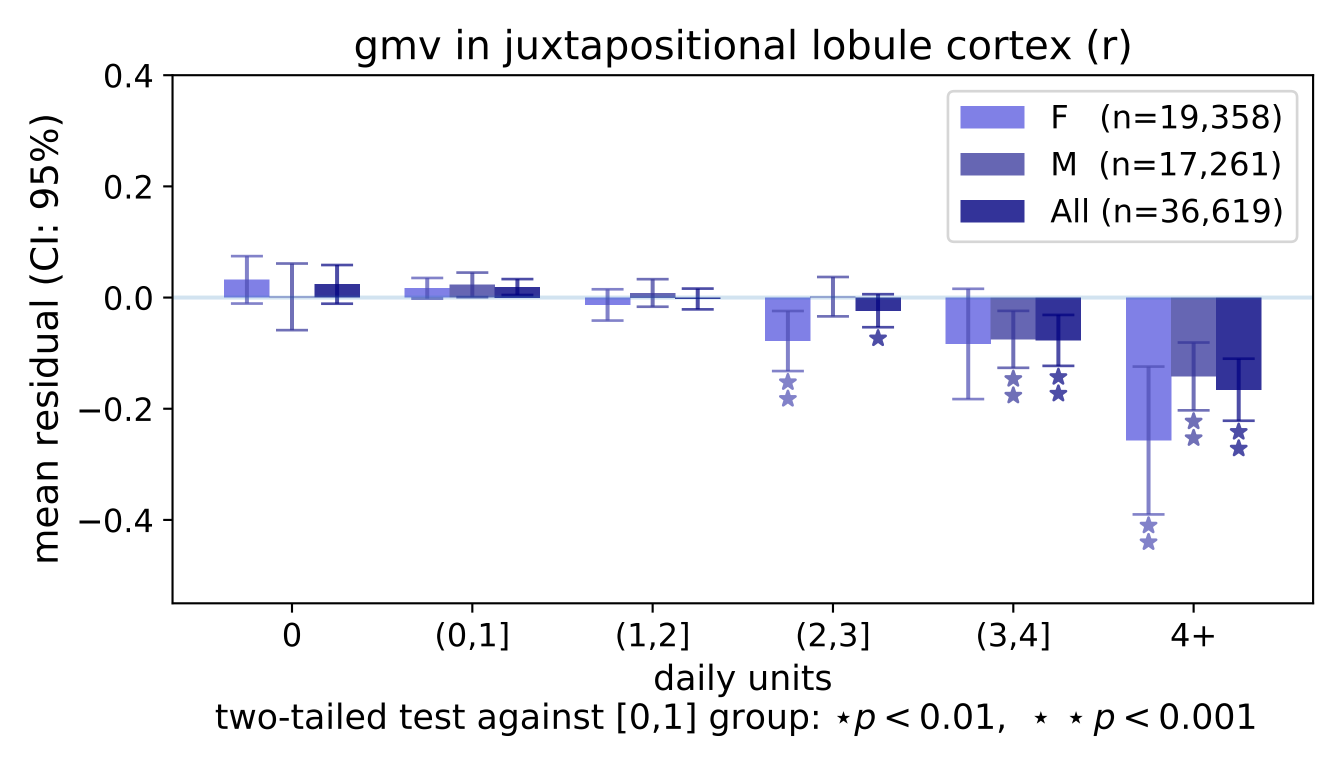
gmv in juxtapositional lobule cortex (r)
mean residual (CI: 95%)
0.4
0.2
0.0
−0.2
−0.4
F (n=19,358)
M (n=17,261)
All (n=36,619)
0
(0,1]
(1,2]
(2,3]
(3,4]
4+
daily units
two-tailed test against [0,1] group: ⋆ $p < 0.01$, ⋆ ⋆ $p < 0.001$

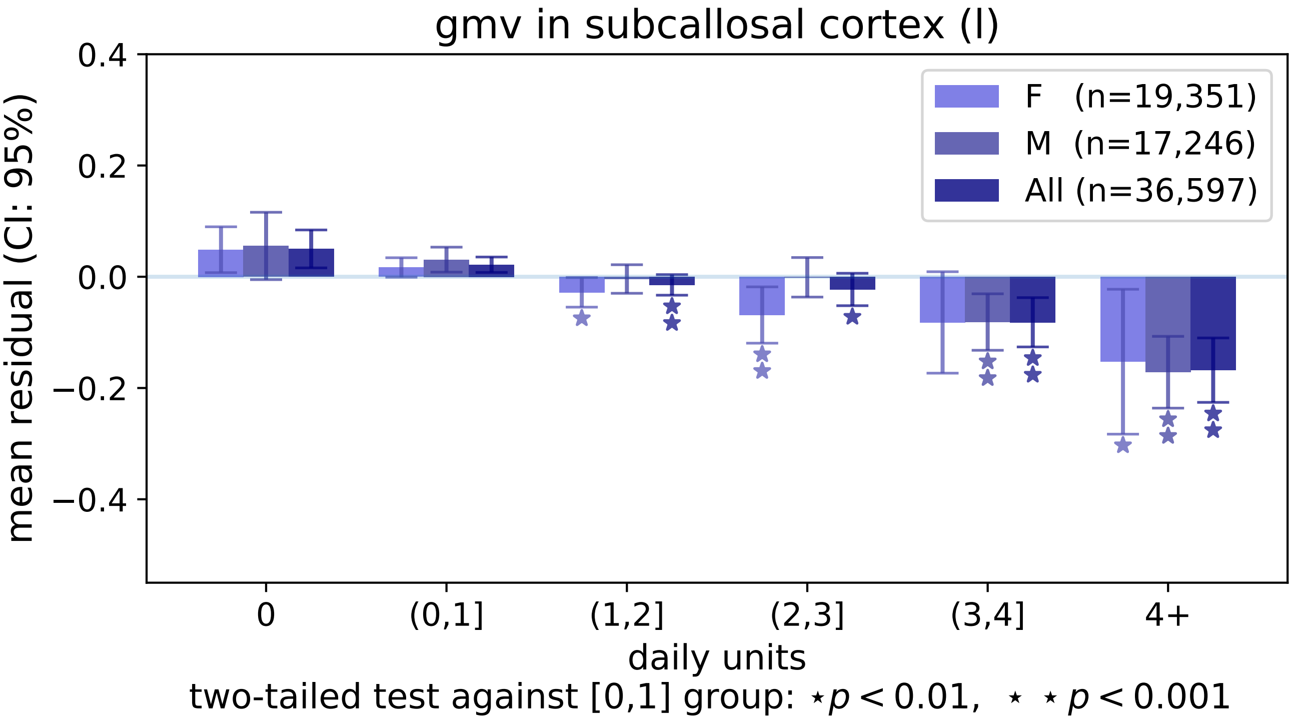

gmv in subcallosal cortex (l)
mean residual (CI: 95%)
0.4
0.2
0.0
−0.2
−0.4
F (n=19,351)
M (n=17,246)
All (n=36,597)
0
(0,1]
(1,2]
(2,3]
(3,4]
4+
daily units
two-tailed test against [0,1] group: ⋆ $p < 0.01$, ⋆ ⋆ $p < 0.001$

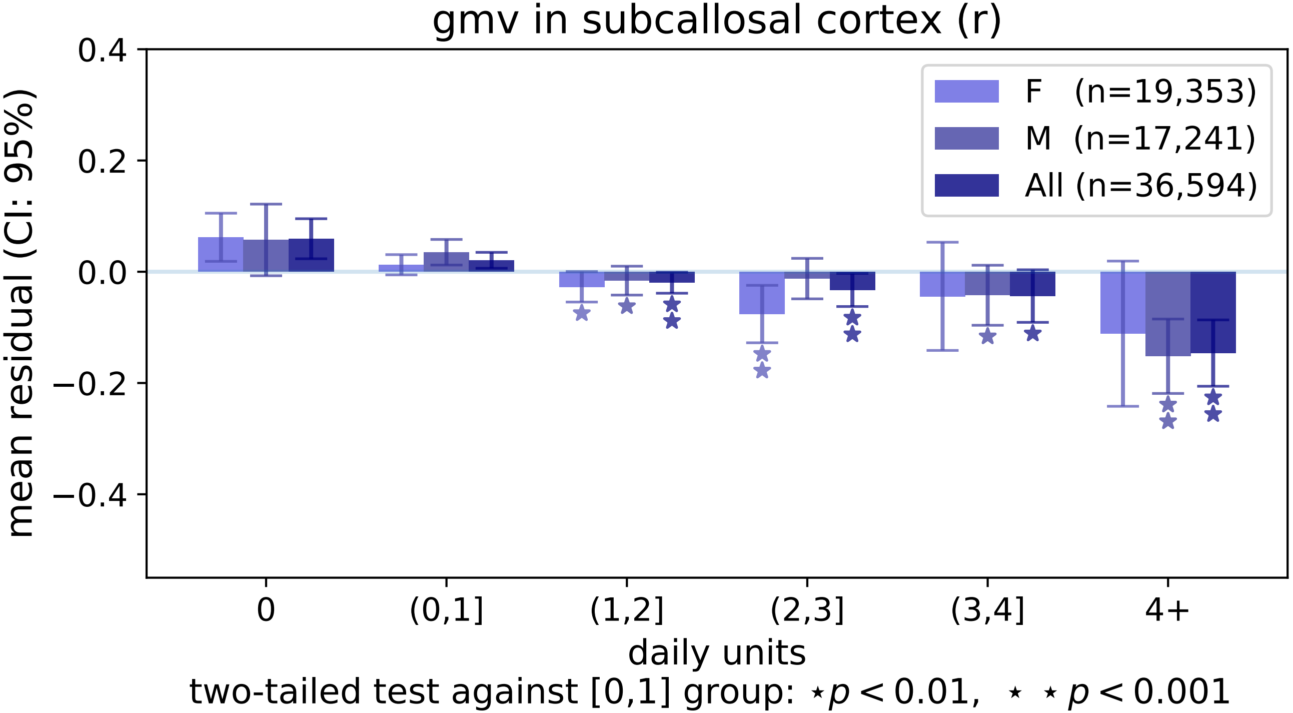

gmv in subcallosal cortex (r)
mean residual (CI: 95%)
0.4
0.2
0.0
−0.2
−0.4
F (n=19,353)
M (n=17,241)
All (n=36,594)
0
(0,1]
(1,2]
(2,3]
(3,4]
4+
daily units
two-tailed test against [0,1] group: ⋆p < 0.01, ⋆ ⋆ p < 0.001

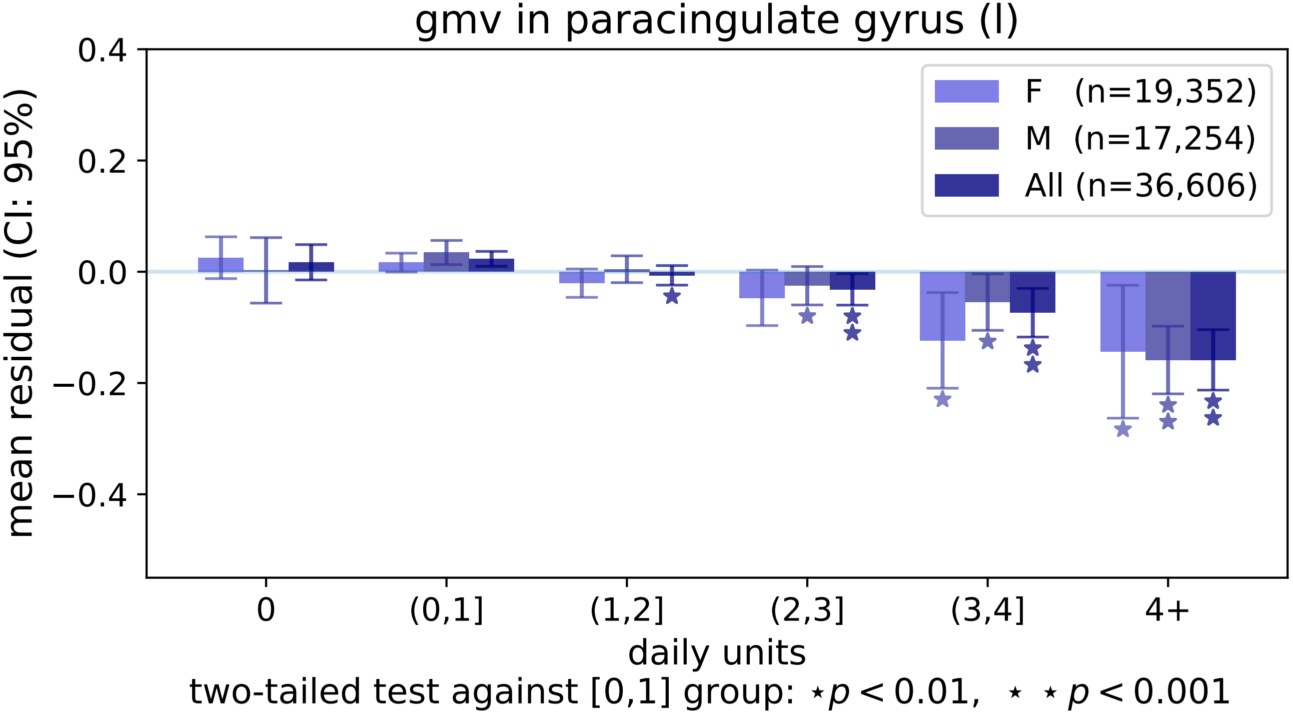
gmv in paracingulate gyrus (l)
mean residual (CI: 95%)
0.4
0.2
0.0
−0.2
−0.4
F (n=19,352)
M (n=17,254)
All (n=36,606)
0
(0,1]
(1,2]
(2,3]
(3,4]
4+
daily units
two-tailed test against [0,1] group: ⋆ $p < 0.01$, ⋆ ⋆ $p < 0.001$

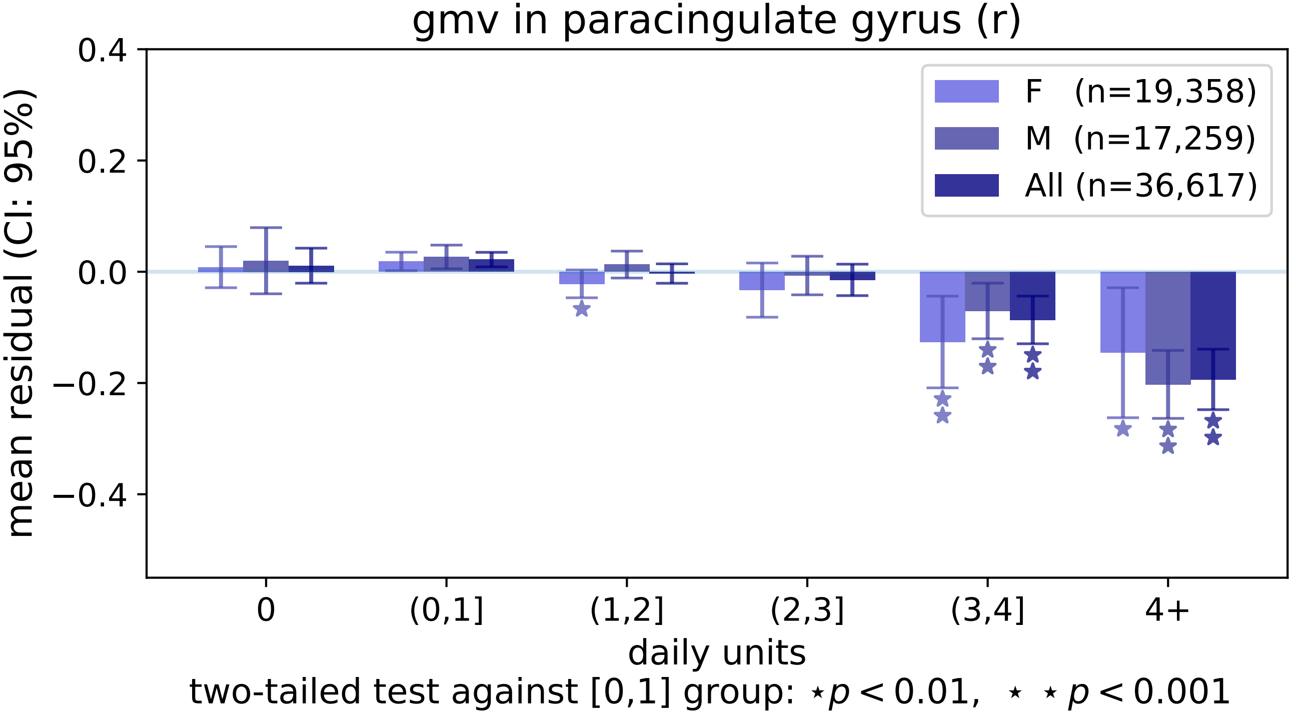
gmv in paracingulate gyrus (r)
mean residual (CI: 95%)
0.4
0.2
0.0
−0.2
−0.4
F (n=19,358)
M (n=17,259)
All (n=36,617)
0
(0,1]
(1,2]
(2,3]
(3,4]
4+
daily units
two-tailed test against [0,1] group: ⋆ p < 0.01, ⋆ ⋆ p < 0.001

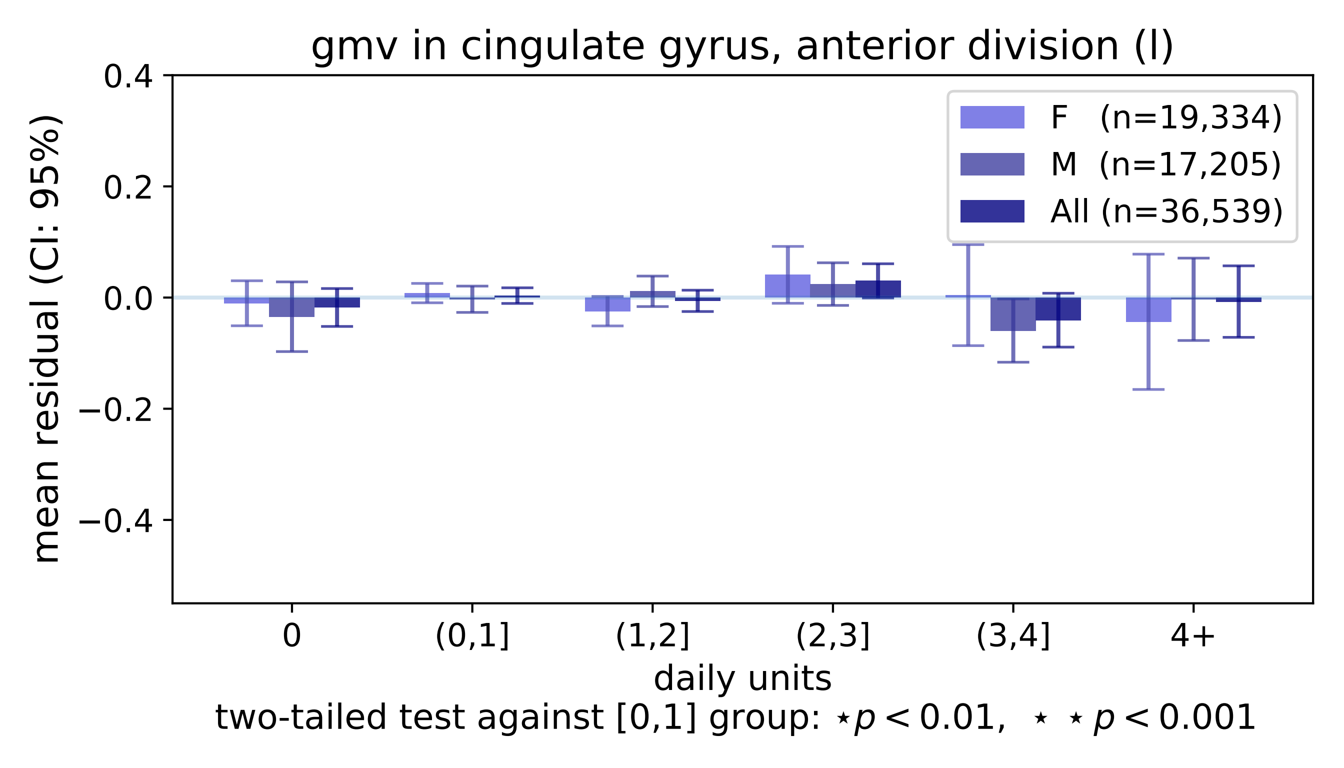

gmv in cingulate gyrus, anterior division (l)
F (n=19,334)
M (n=17,205)
All (n=36,539)
0.4
0.2
0.0
−0.2
−0.4
mean residual (CI: 95%)
0
(0,1]
(1,2]
(2,3]
(3,4]
4+
daily units
two-tailed test against [0,1] group: ⋆p < 0.01, ⋆ ⋆ p < 0.001

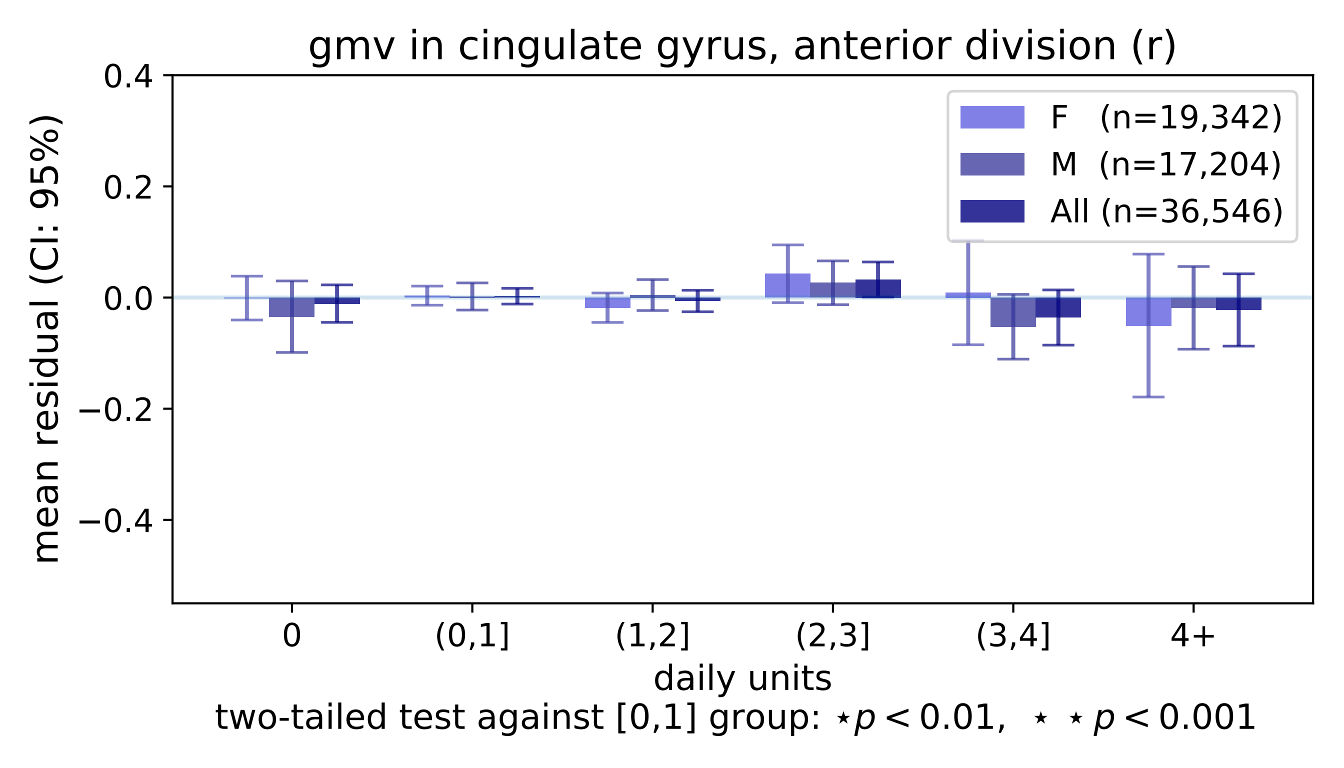

gmv in cingulate gyrus, anterior division (r)
mean residual (CI: 95%)
0.4
0.2
0.0
−0.2
−0.4
F (n=19,342)
M (n=17,204)
All (n=36,546)
0
(0,1]
(1,2]
(2,3]
(3,4]
4+
daily units
two-tailed test against [0,1] group: ⋆p < 0.01, ⋆ ⋆ p < 0.001

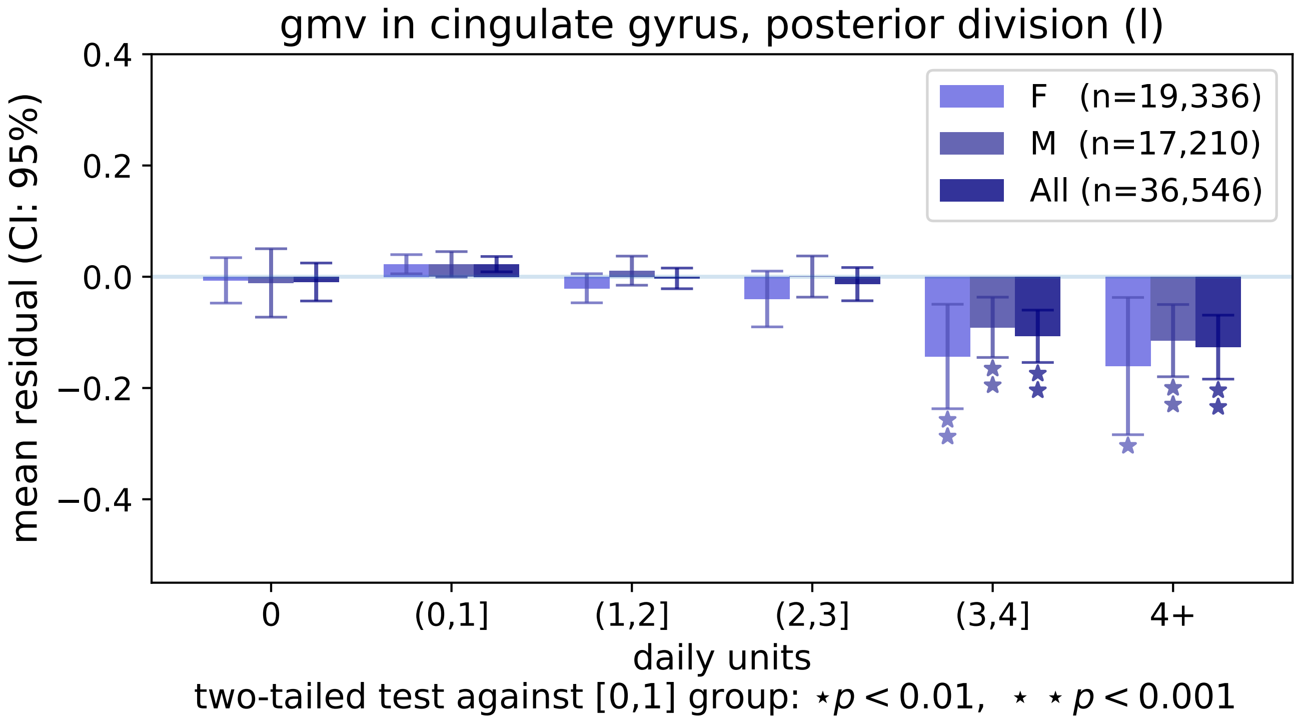
gmv in cingulate gyrus, posterior division (l)
mean residual (CI: 95%)
0.4
0.2
0.0
−0.2
−0.4
F (n=19,336)
M (n=17,210)
All (n=36,546)
0
(0,1]
(1,2]
(2,3]
(3,4]
4+
daily units
two-tailed test against [0,1] group: ⋆p < 0.01, ⋆ ⋆ p < 0.001

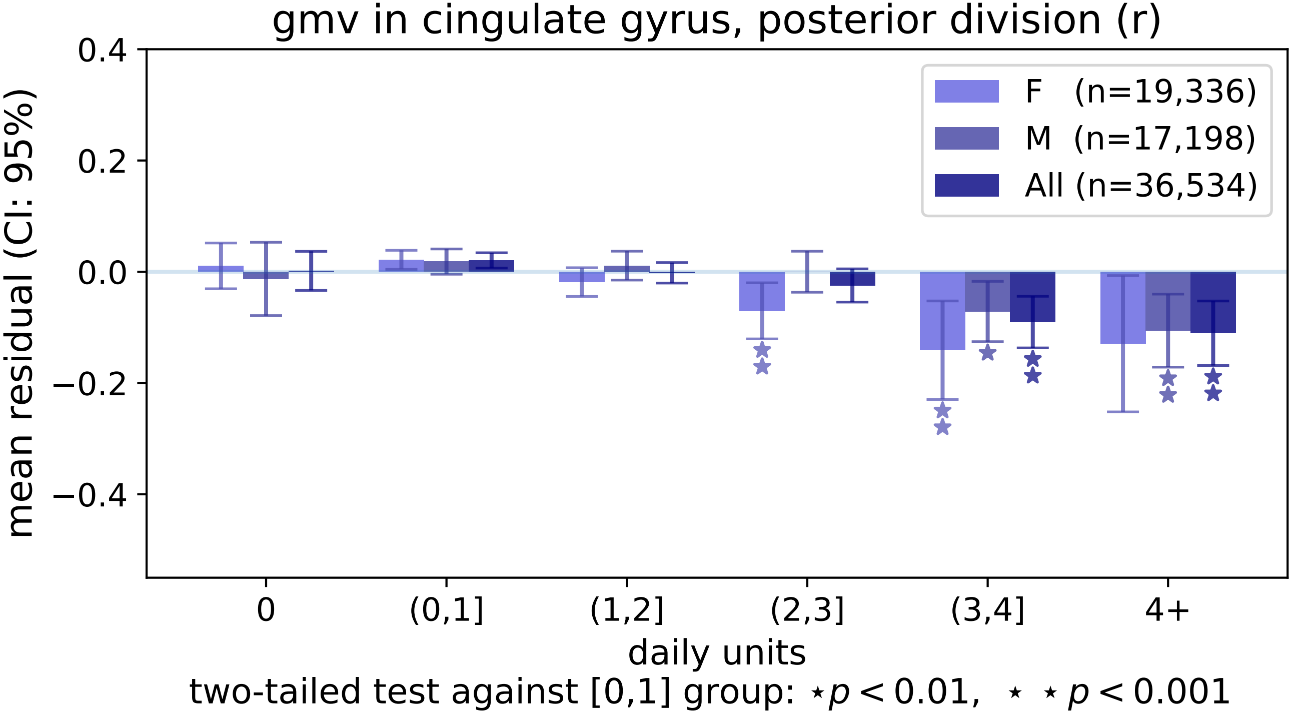

gmv in cingulate gyrus, posterior division (r)
mean residual (CI: 95%)
0.4
0.2
0.0
−0.2
−0.4
F (n=19,336)
M (n=17,198)
All (n=36,534)
0
(0,1]
(1,2]
(2,3]
(3,4]
4+
daily units
two-tailed test against [0,1] group: ⋆p < 0.01, ⋆ ⋆ p < 0.001

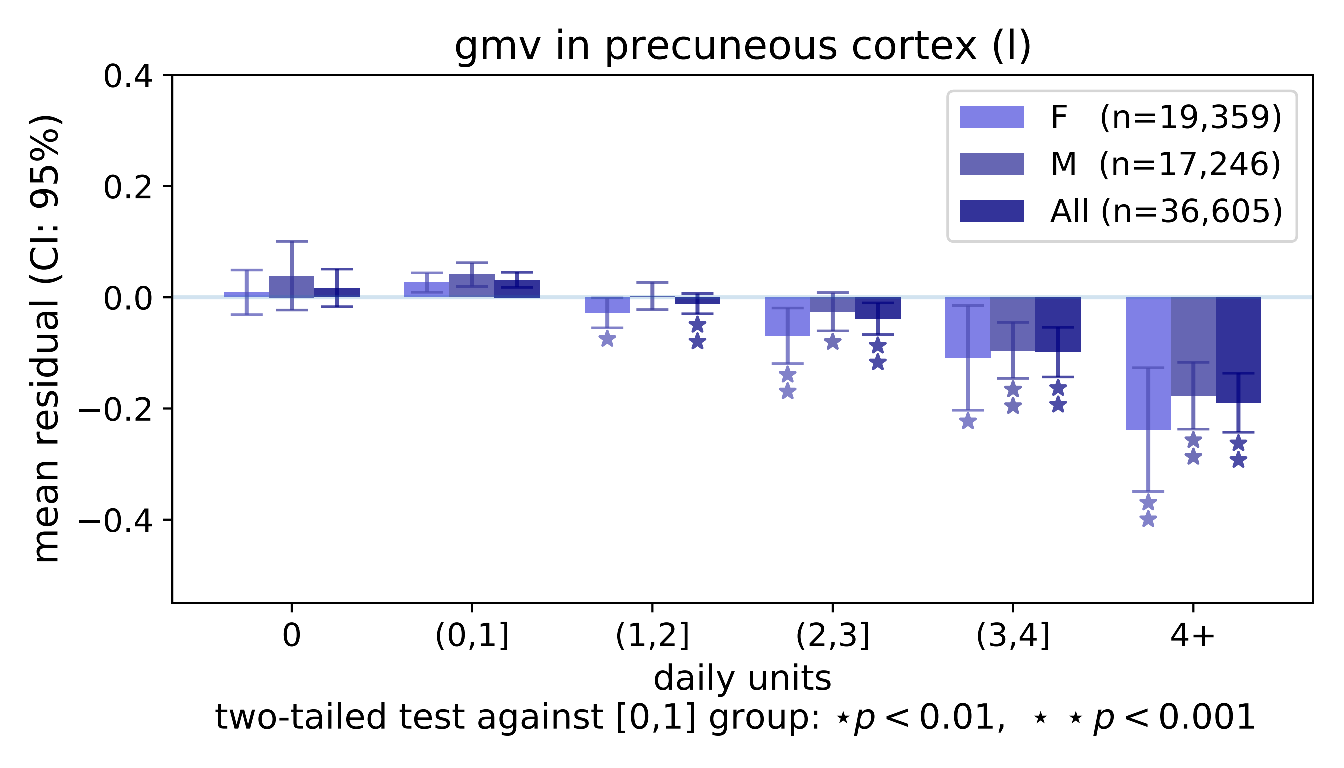

gmv in precuneous cortex (l)
mean residual (CI: 95%)
0.4
0.2
0.0
−0.2
−0.4
F (n=19,359)
M (n=17,246)
All (n=36,605)
0
(0,1]
(1,2]
(2,3]
(3,4]
4+
daily units
two-tailed test against [0,1] group: ⋆ p < 0.01, ⋆ ⋆ p < 0.001

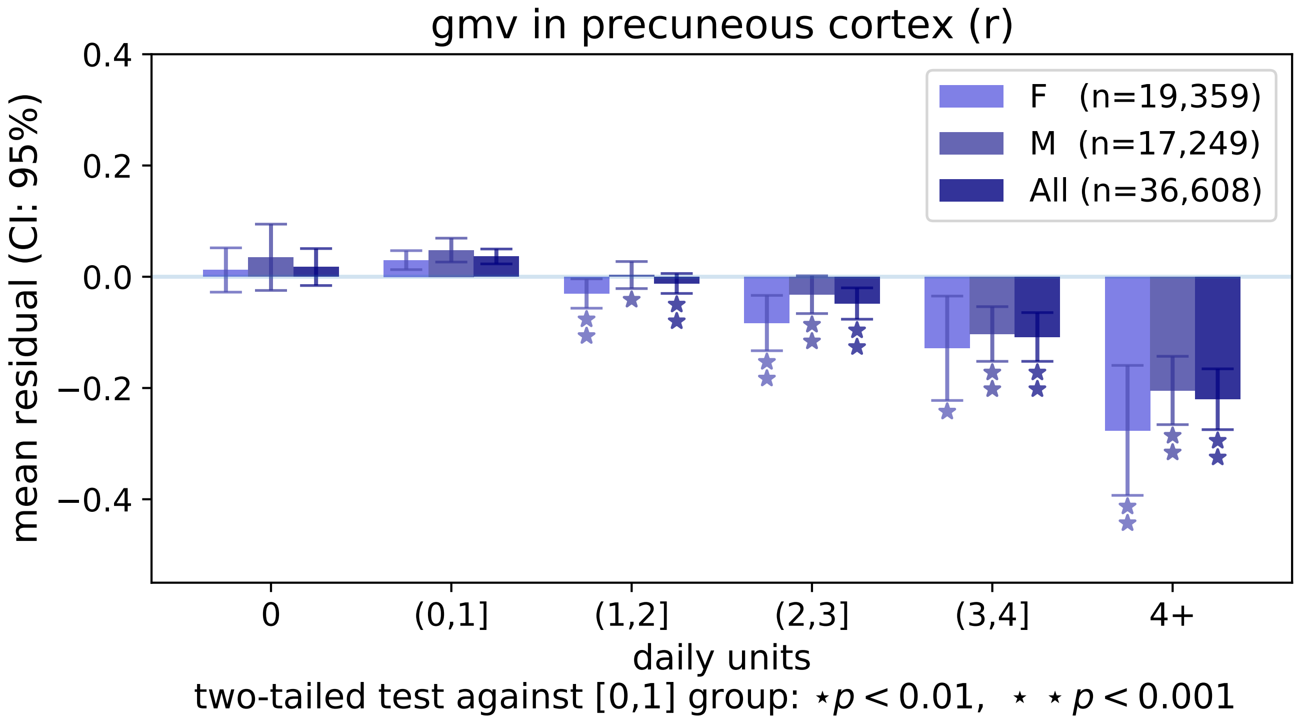
gmv in precuneous cortex (r)
0.4
0.2
0.0
−0.2
−0.4
mean residual (CI: 95%)
F (n=19,359)
M (n=17,249)
All (n=36,608)
0
(0,1]
(1,2]
(2,3]
(3,4]
4+
daily units
two-tailed test against [0,1] group: ⋆p < 0.01, ⋆ ⋆ p < 0.001

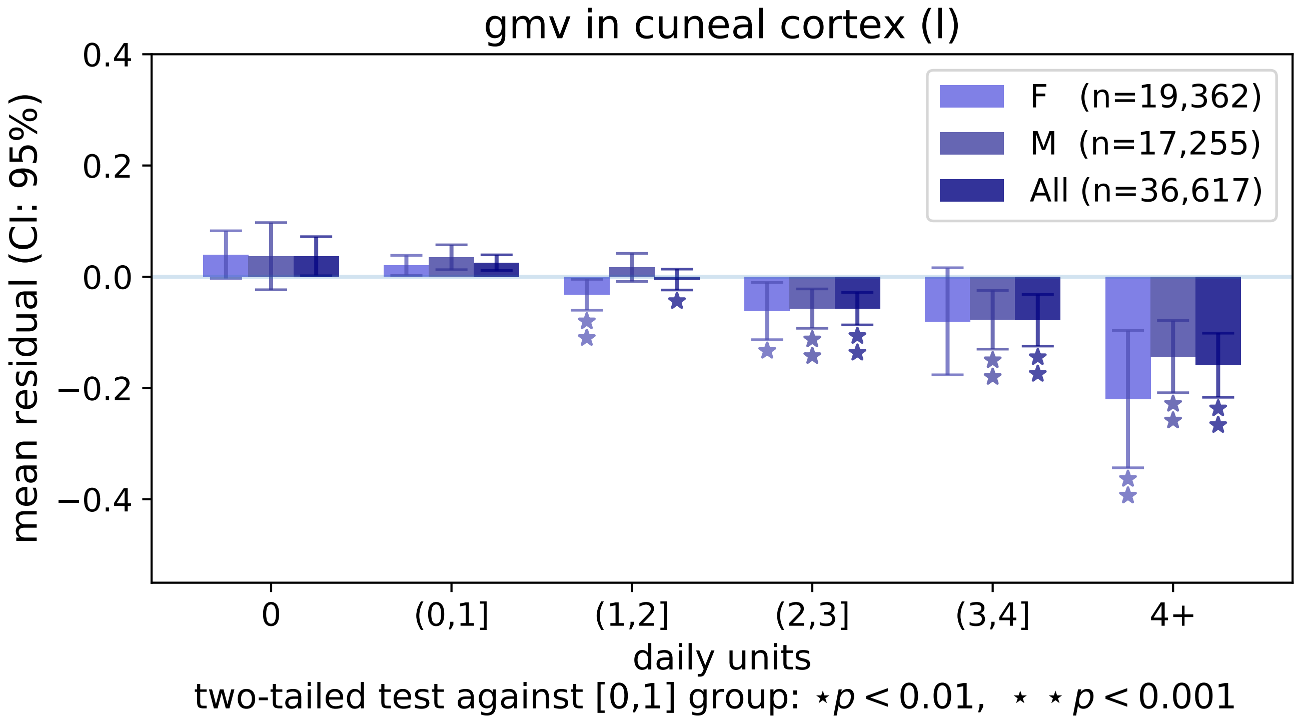

gmv in cuneal cortex (l)
mean residual (CI: 95%)
0.4
0.2
0.0
−0.2
−0.4
F (n=19,362)
M (n=17,255)
All (n=36,617)
0
(0,1]
(1,2]
(2,3]
(3,4]
4+
daily units
two-tailed test against [0,1] group: ⋆ p < 0.01, ⋆ ⋆ p < 0.001

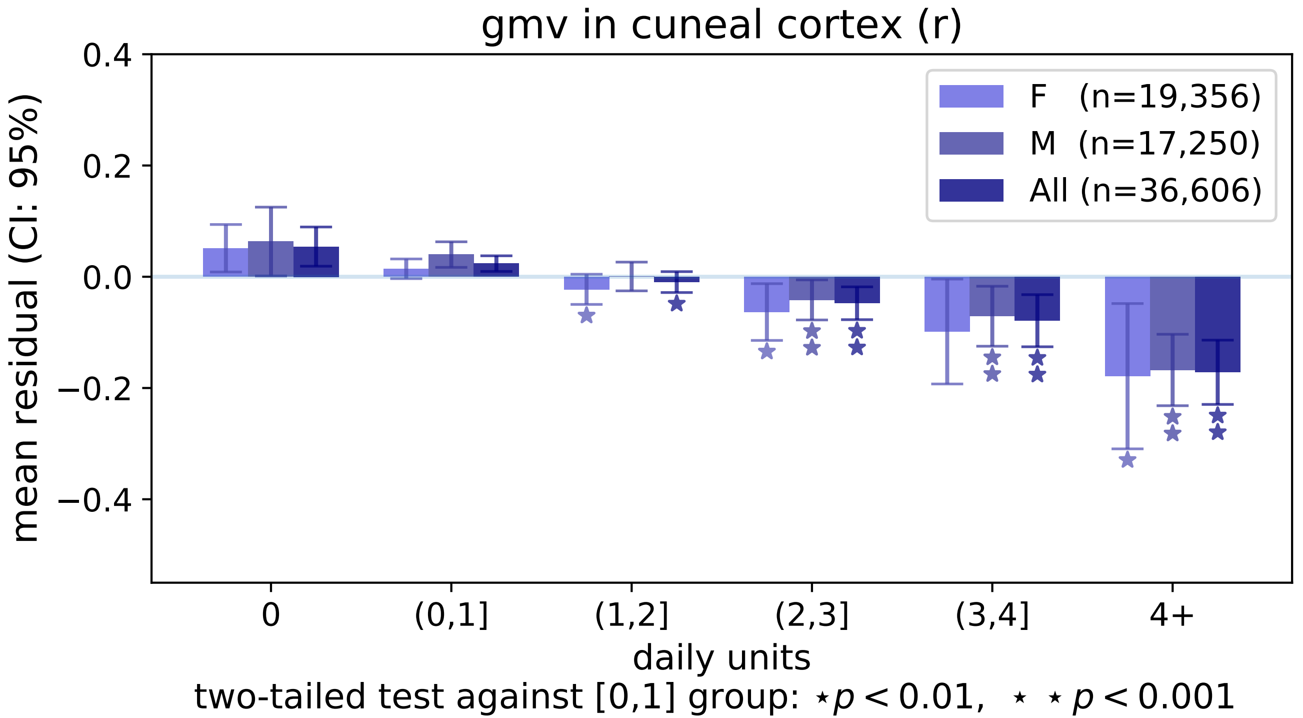

gmv in cuneal cortex (r)
0.4
0.2
0.0
−0.2
−0.4
mean residual (CI: 95%)
F (n=19,356)
M (n=17,250)
All (n=36,606)
0
(0,1]
(1,2]
(2,3]
(3,4]
4+
daily units
two-tailed test against [0,1] group: ⋆ p < 0.01, ⋆ ⋆ p < 0.001

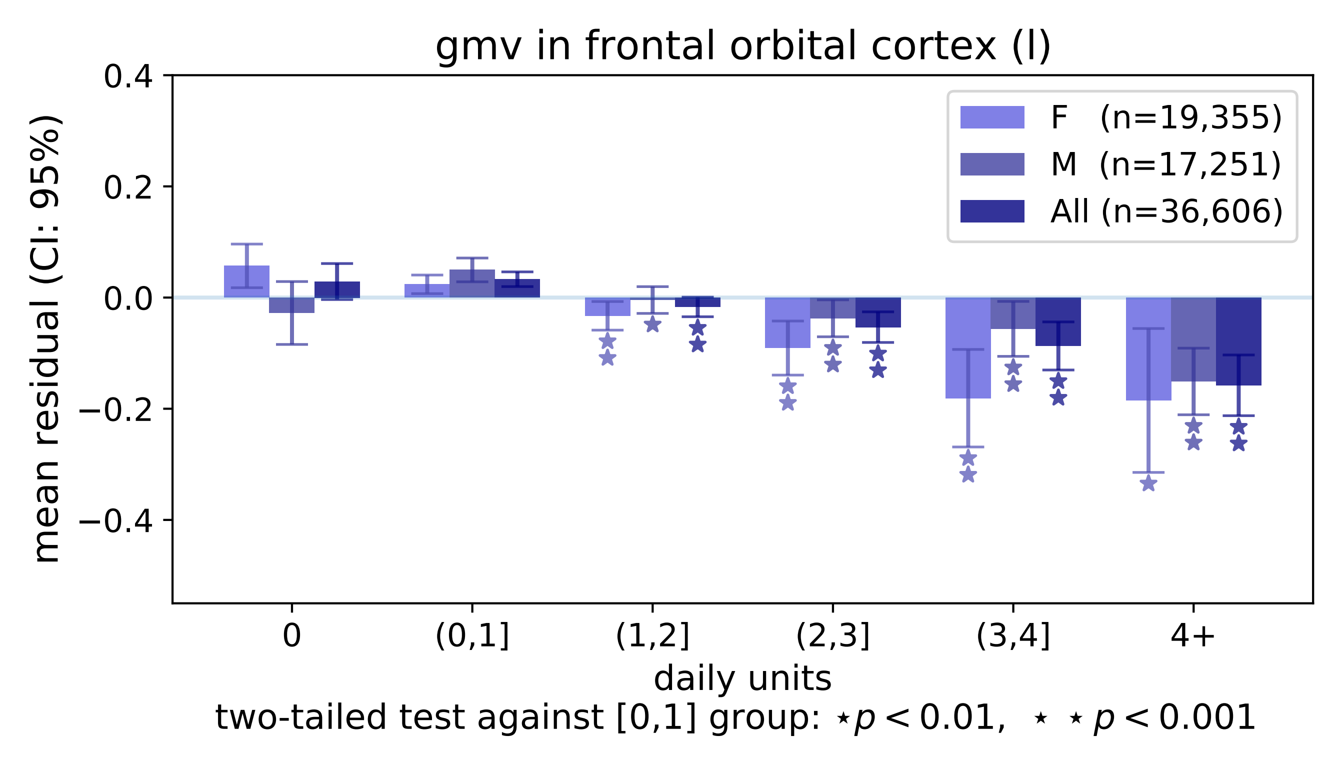

gmv in frontal orbital cortex (l)
mean residual (CI: 95%)
0.4
0.2
0.0
−0.2
−0.4
F (n=19,355)
M (n=17,251)
All (n=36,606)
0
(0,1]
(1,2]
(2,3]
(3,4]
4+
daily units
two-tailed test against [0,1] group: ⋆p < 0.01, ⋆ ⋆ p < 0.001

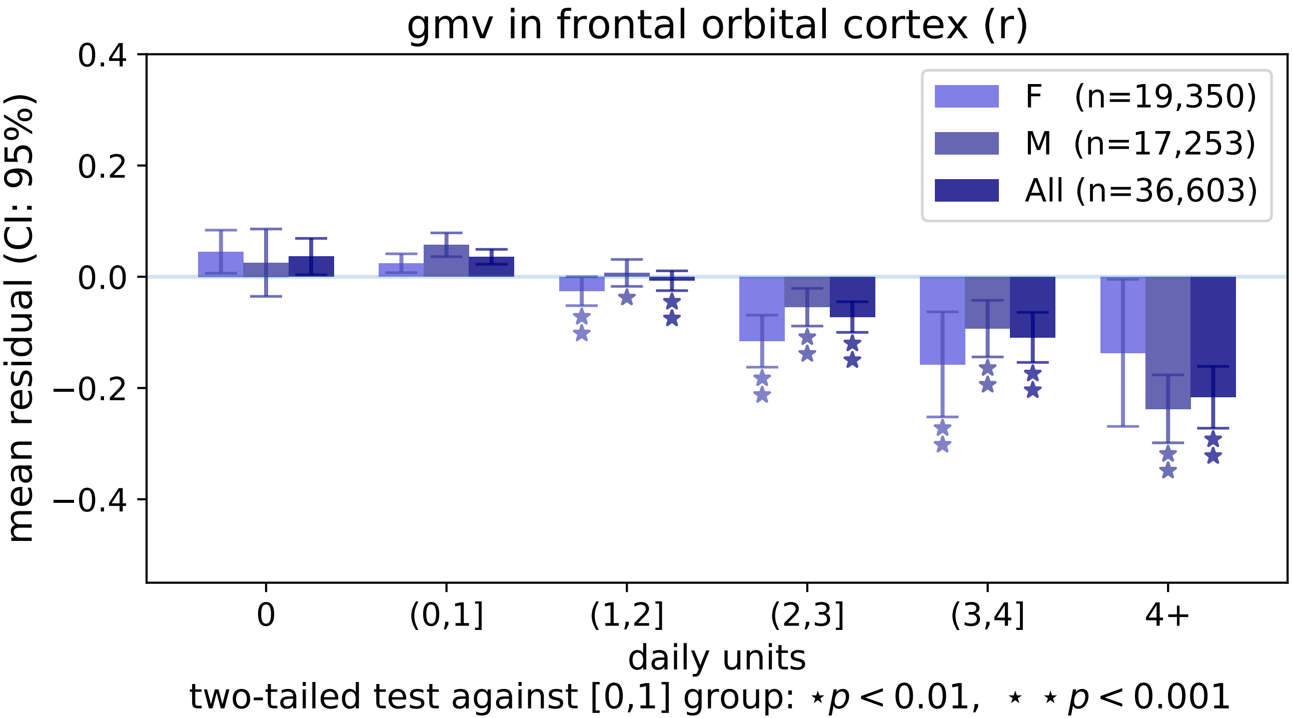
gmv in frontal orbital cortex (r)
0.4
0.2
0.0
−0.2
−0.4
mean residual (CI: 95%)
F (n=19,350)
M (n=17,253)
All (n=36,603)
0
(0,1]
(1,2]
(2,3]
(3,4]
4+
daily units
two-tailed test against [0,1] group: ⋆p < 0.01, ⋆ ⋆ p < 0.001

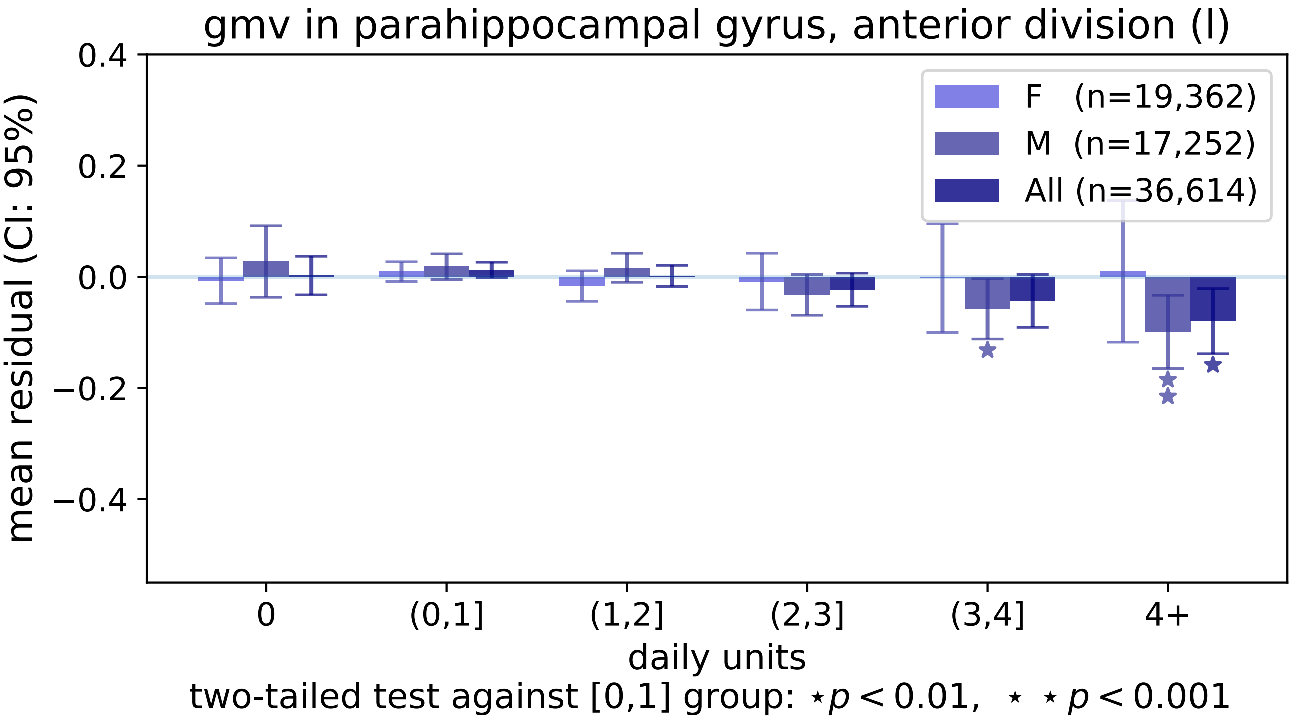
gmv in parahippocampal gyrus, anterior division (l)
0.4
0.2
0.0
−0.2
−0.4
mean residual (CI: 95%)
F (n=19,362)
M (n=17,252)
All (n=36,614)
0
(0,1]
(1,2]
(2,3]
(3,4]
4+
daily units
two-tailed test against [0,1] group: ⋆p < 0.01, ⋆ ⋆ p < 0.001

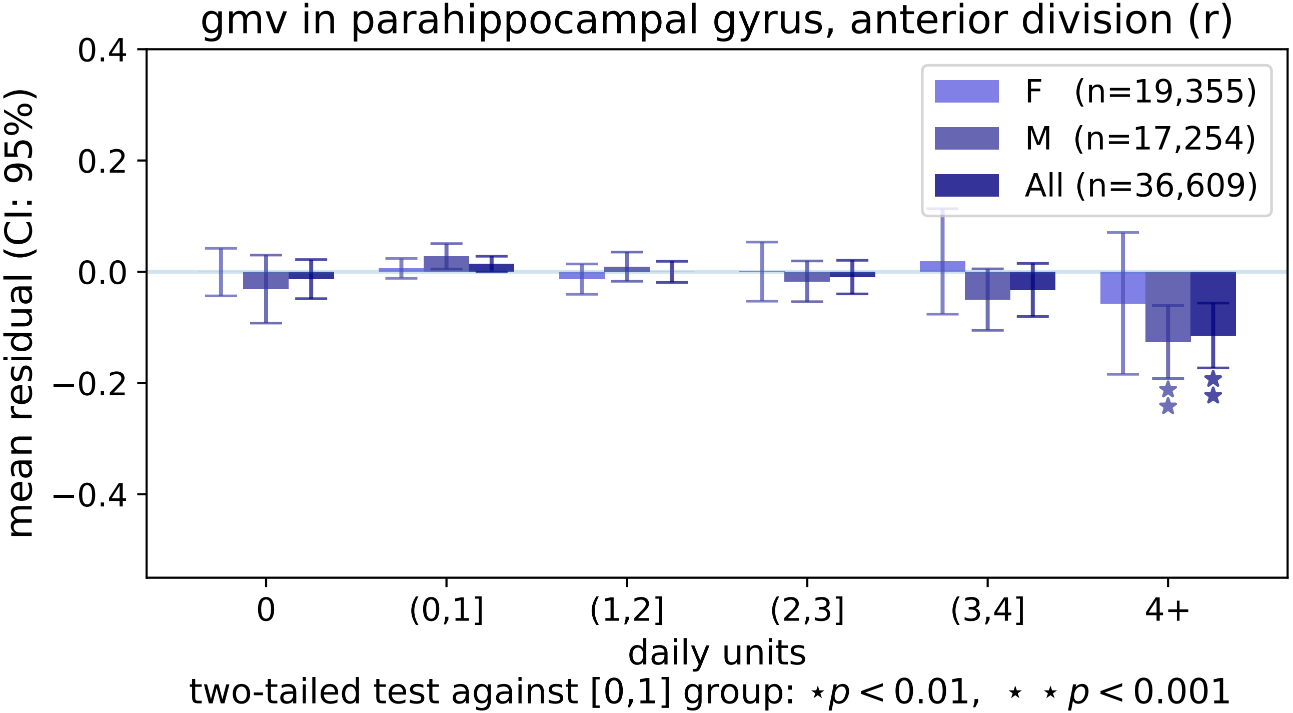
gmv in parahippocampal gyrus, anterior division (r)
mean residual (CI: 95%)
0.4
0.2
0.0
−0.2
−0.4
F (n=19,355)
M (n=17,254)
All (n=36,609)
0
(0,1]
(1,2]
(2,3]
(3,4]
4+
daily units
two-tailed test against [0,1] group: ⋆p < 0.01, ⋆ ⋆ p < 0.001

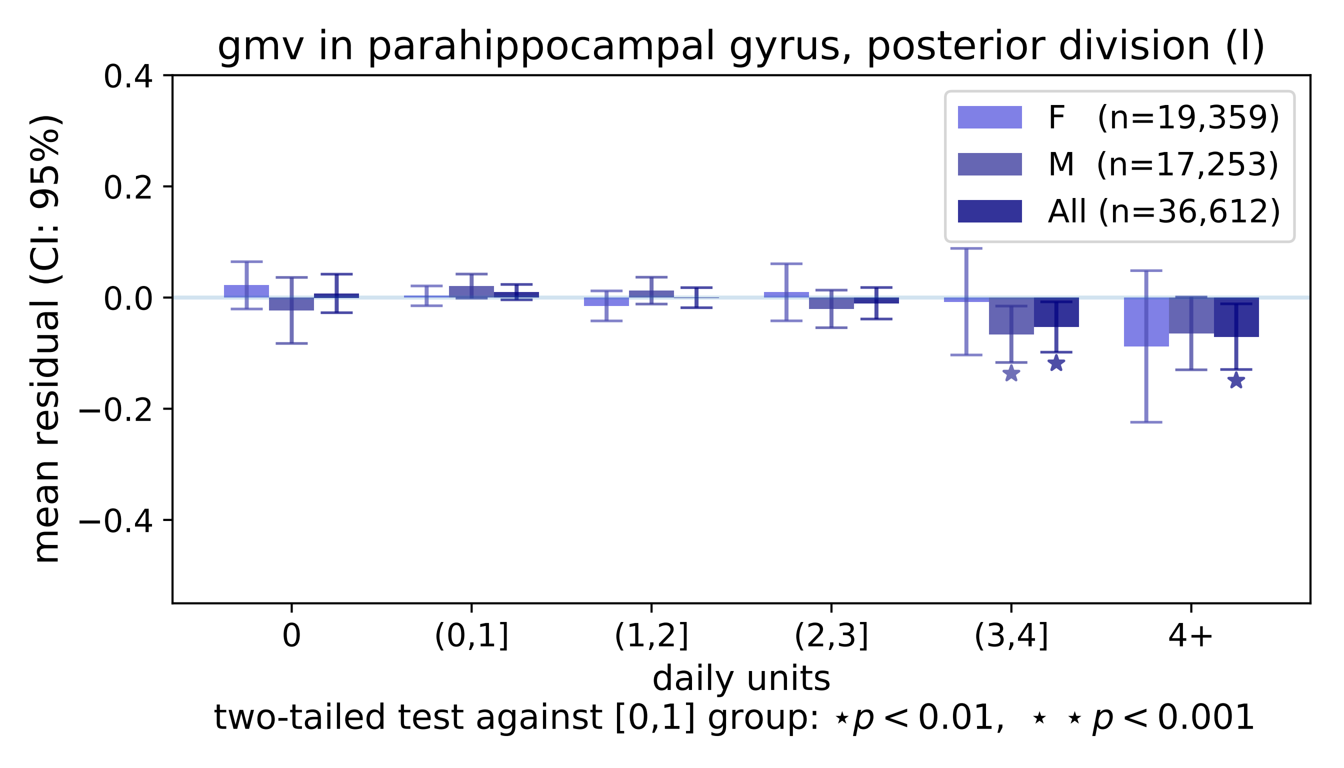
gmv in parahippocampal gyrus, posterior division (l)
0.4
0.2
0.0
−0.2
−0.4
mean residual (CI: 95%)
F (n=19,359)
M (n=17,253)
All (n=36,612)
0
(0,1]
(1,2]
(2,3]
(3,4]
4+
daily units
two-tailed test against [0,1] group: ⋆p < 0.01, ⋆ ⋆p < 0.001

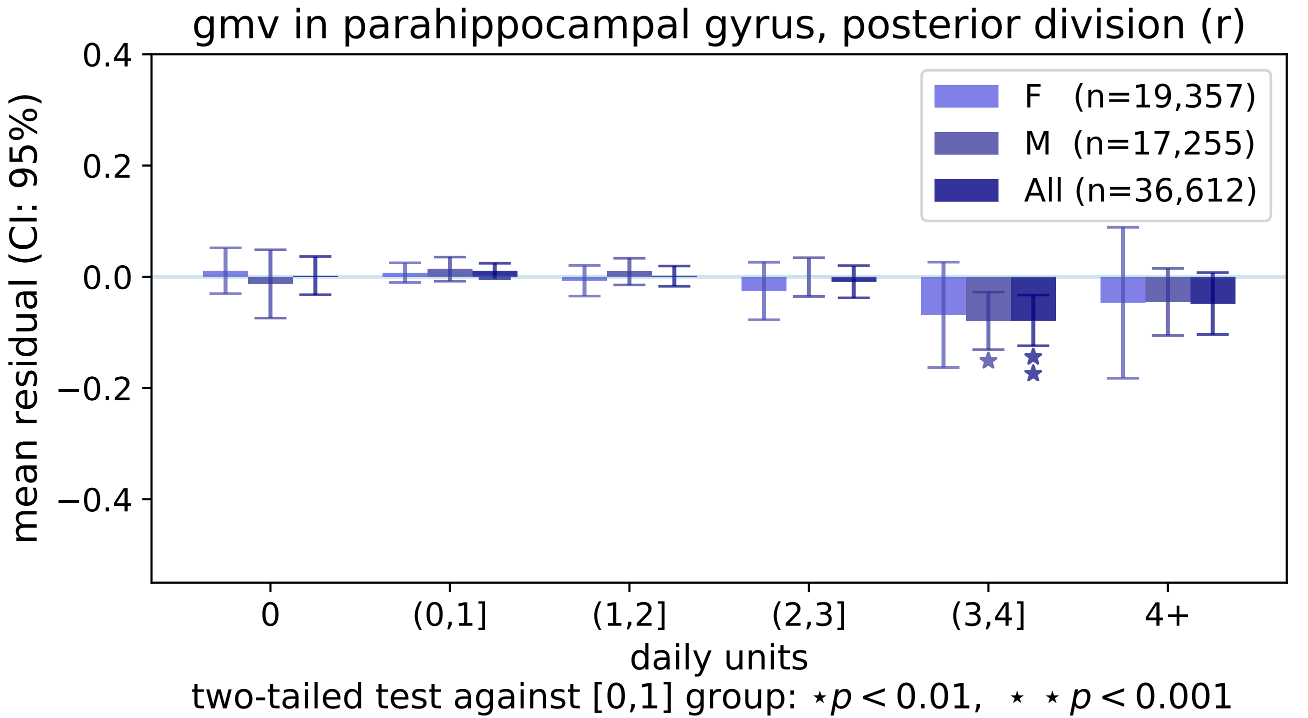

gmv in parahippocampal gyrus, posterior division (r)
0.4
0.2
0.0
−0.2
−0.4
mean residual (CI: 95%)
F (n=19,357)
M (n=17,255)
All (n=36,612)
0
(0,1]
(1,2]
(2,3]
(3,4]
4+
daily units
two-tailed test against [0,1] group: ⋆p < 0.01, ⋆ ⋆ p < 0.001

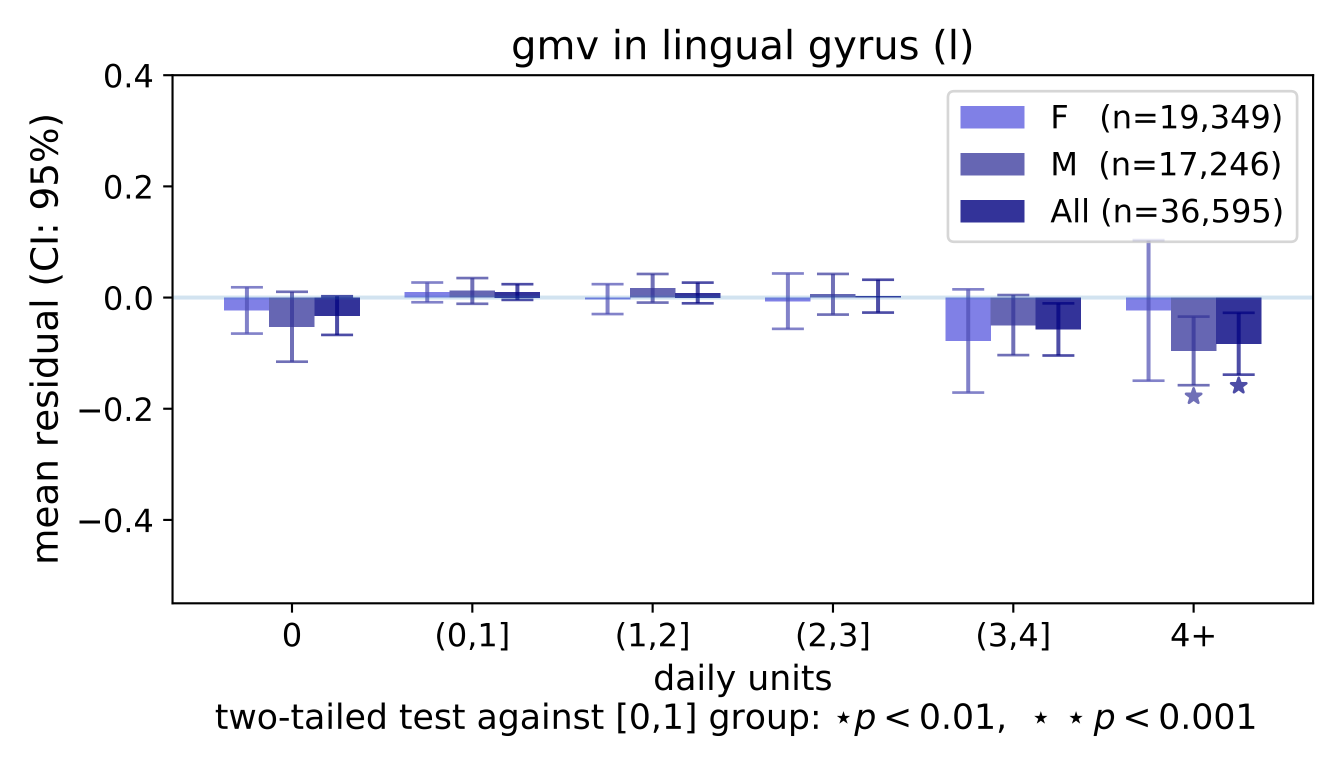

gmv in lingual gyrus (l)
mean residual (CI: 95%)
0.4
0.2
0.0
−0.2
−0.4
F (n=19,349)
M (n=17,246)
All (n=36,595)
0
(0,1]
(1,2]
(2,3]
(3,4]
4+
daily units
two-tailed test against [0,1] group: ⋆p < 0.01, ⋆ ⋆p < 0.001

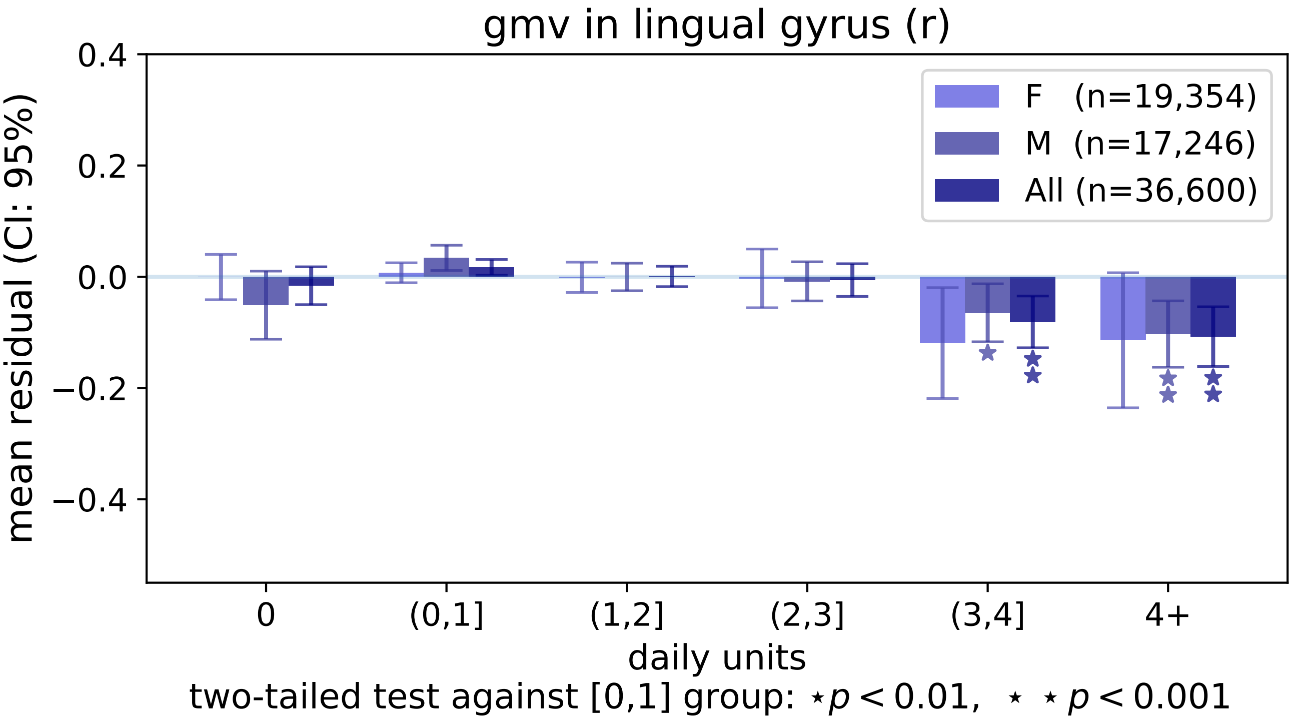
gmv in lingual gyrus (r)
0.4
0.2
0.0
−0.2
−0.4
mean residual (CI: 95%)
F (n=19,354)
M (n=17,246)
All (n=36,600)
0
(0,1]
(1,2]
(2,3]
(3,4]
4+
daily units
two-tailed test against [0,1] group: ⋆$p < 0.01$, ⋆ ⋆$p < 0.001$

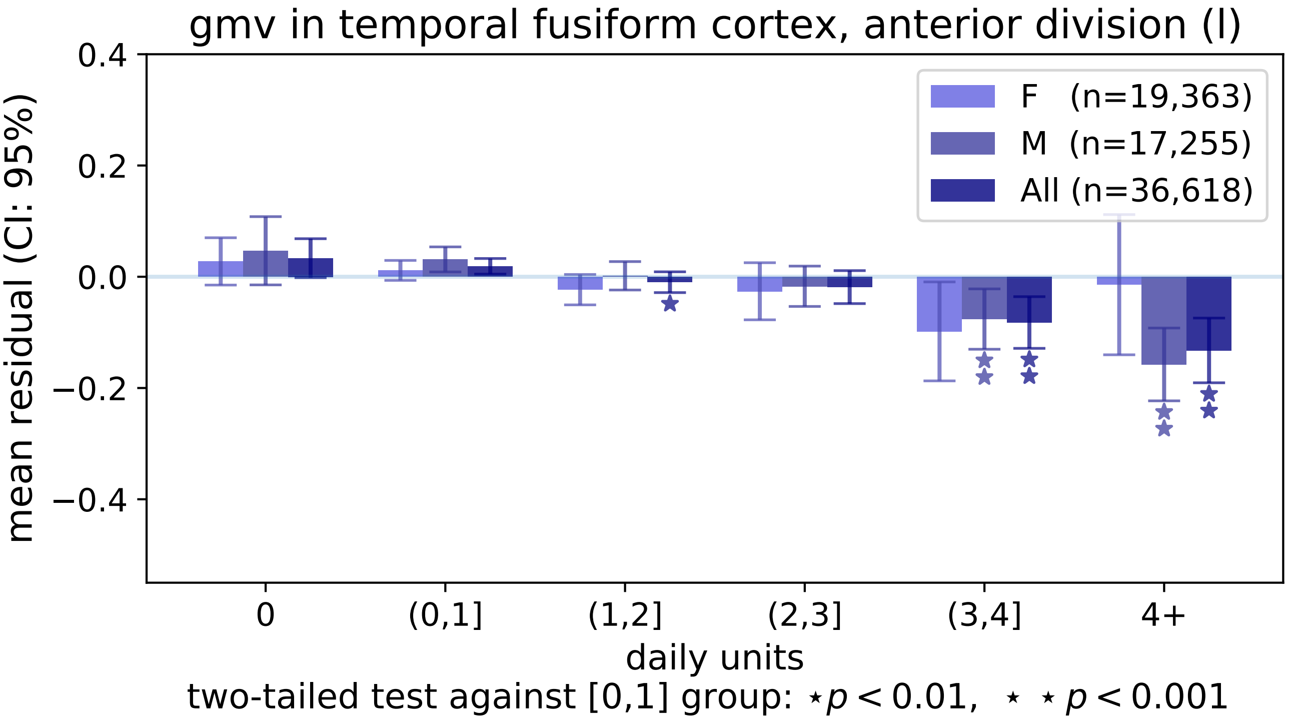

gmv in temporal fusiform cortex, anterior division (l)
0.4
0.2
0.0
−0.2
−0.4
mean residual (CI: 95%)
F (n=19,363)
M (n=17,255)
All (n=36,618)
0
(0,1]
(1,2]
(2,3]
(3,4]
4+
daily units
two-tailed test against [0,1] group: ⋆ p < 0.01, ⋆ ⋆ p < 0.001

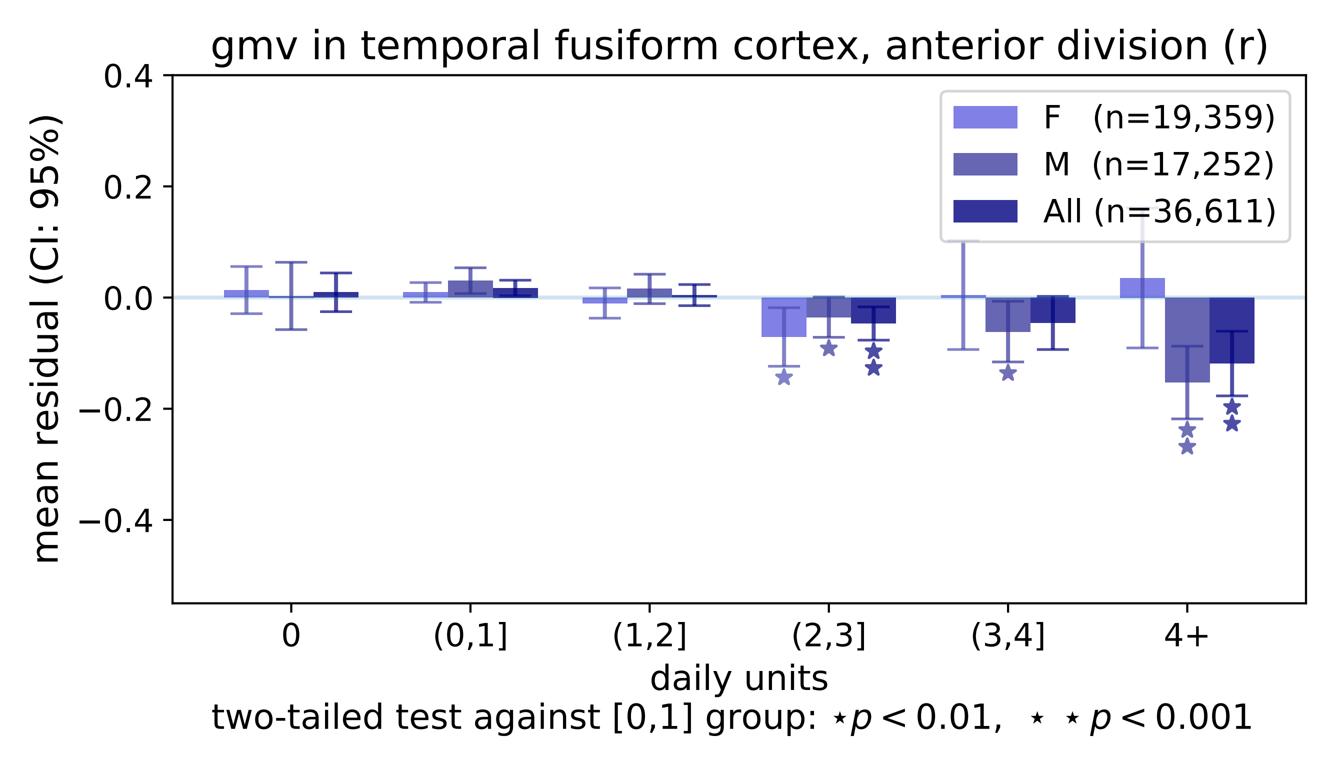

gmv in temporal fusiform cortex, anterior division (r)
0.4
0.2
0.0
−0.2
−0.4
mean residual (CI: 95%)
F (n=19,359)
M (n=17,252)
All (n=36,611)
0
(0,1]
(1,2]
(2,3]
(3,4]
4+
daily units
two-tailed test against [0,1] group: ⋆p < 0.01, ⋆ ⋆ p < 0.001

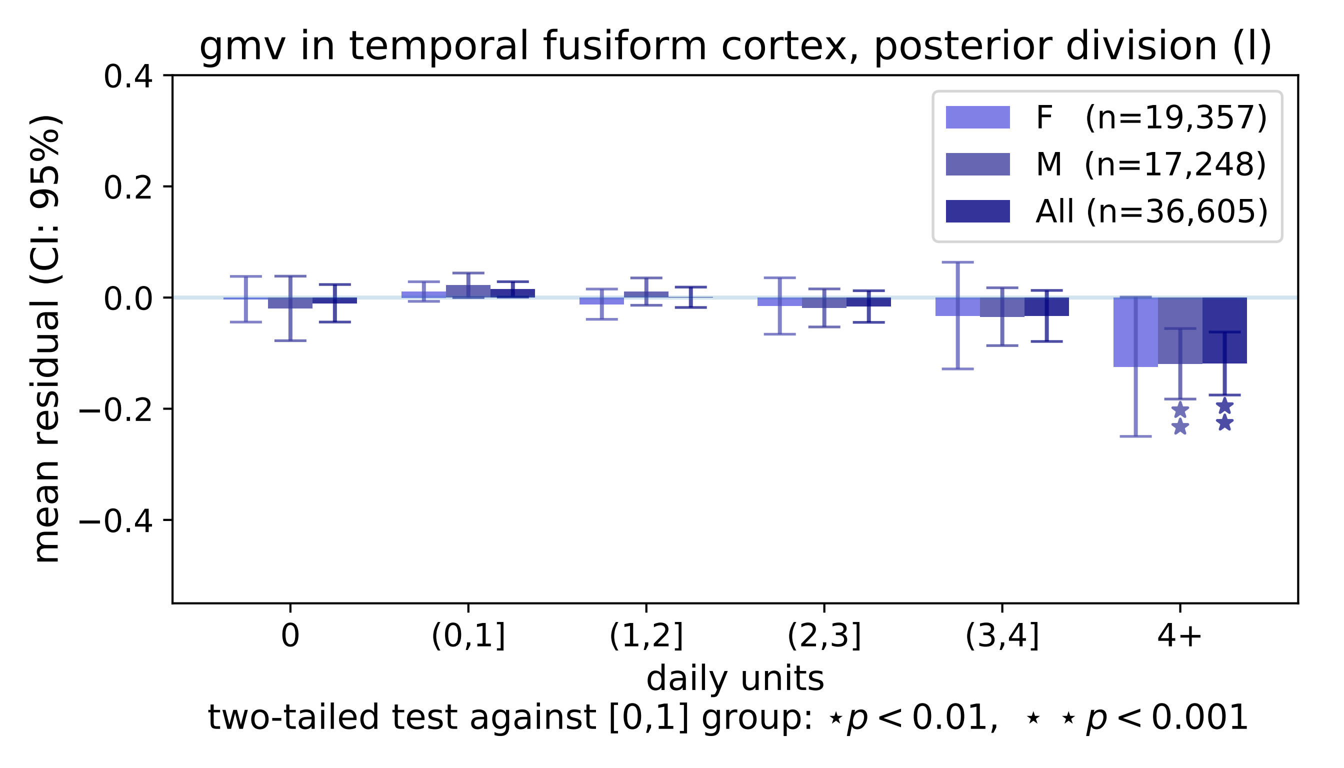
gmv in temporal fusiform cortex, posterior division (l)
0.4
0.2
0.0
−0.2
−0.4
mean residual (CI: 95%)
F (n=19,357)
M (n=17,248)
All (n=36,605)
0
(0,1]
(1,2]
(2,3]
(3,4]
4+
daily units
two-tailed test against [0,1] group: ⋆p < 0.01, ⋆ ⋆ p < 0.001

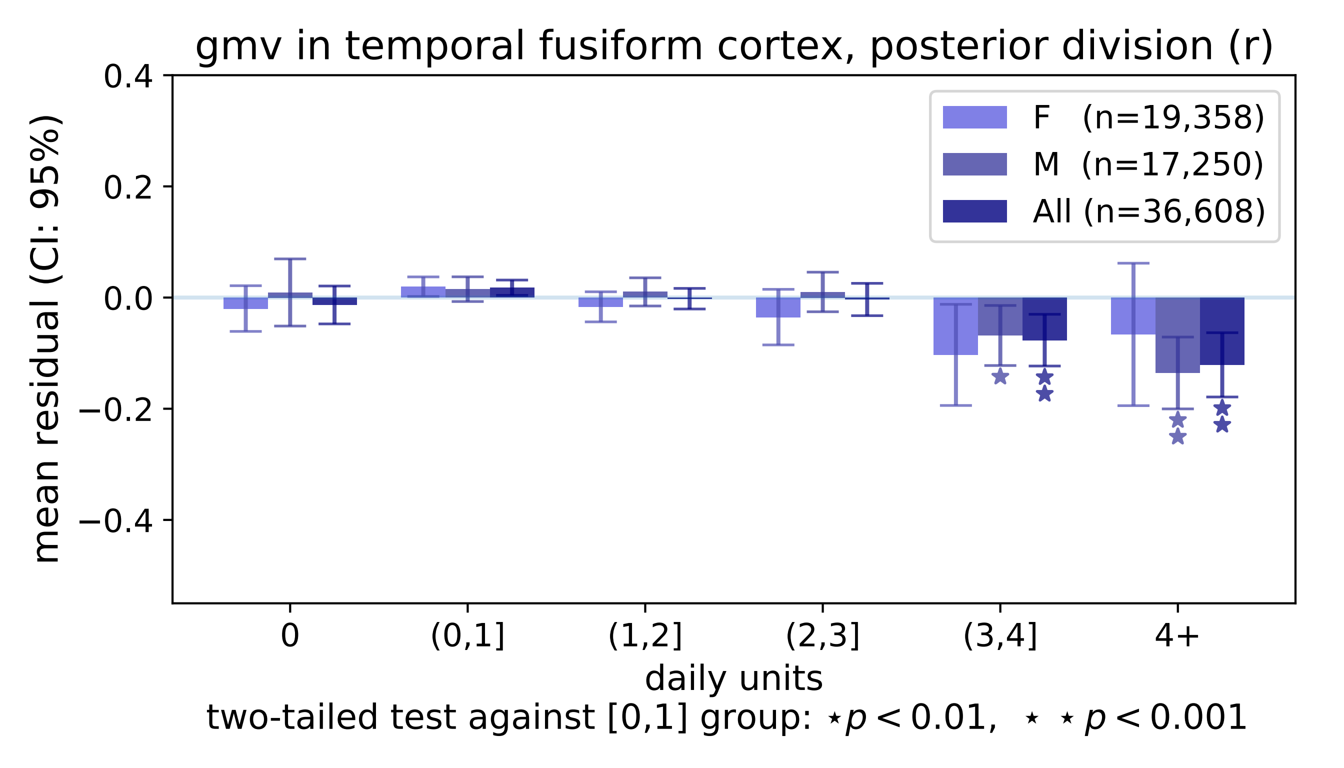

gmv in temporal fusiform cortex, posterior division (r)
mean residual (CI: 95%)
0.4
0.2
0.0
−0.2
−0.4
F (n=19,358)
M (n=17,250)
All (n=36,608)
0
(0,1]
(1,2]
(2,3]
(3,4]
4+
daily units
two-tailed test against [0,1] group: ⋆p < 0.01, ⋆ ⋆ p < 0.001

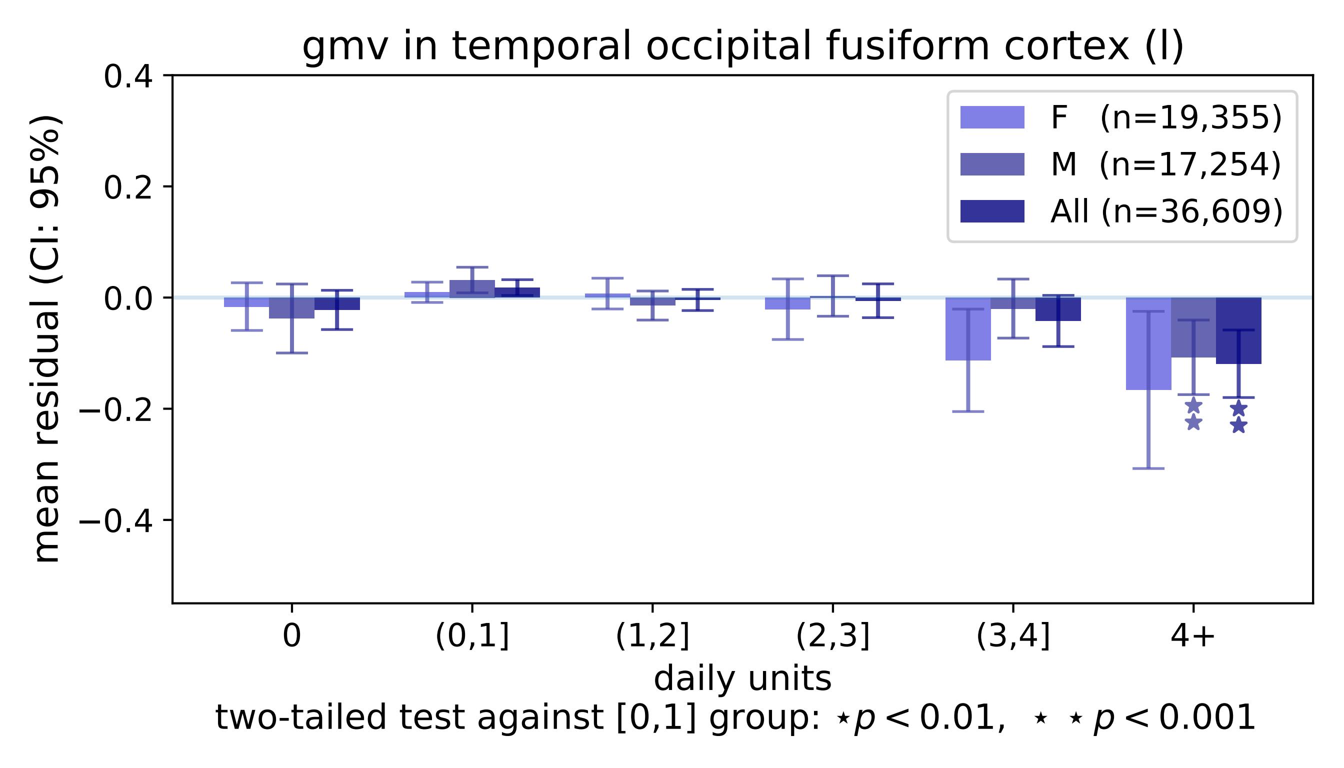
gmv in temporal occipital fusiform cortex (l)
mean residual (CI: 95%)
0.4
0.2
0.0
−0.2
−0.4
F (n=19,355)
M (n=17,254)
All (n=36,609)
0
(0,1]
(1,2]
(2,3]
(3,4]
4+
daily units
two-tailed test against [0,1] group: ⋆ $p < 0.01$, ⋆ ⋆ $p < 0.001$

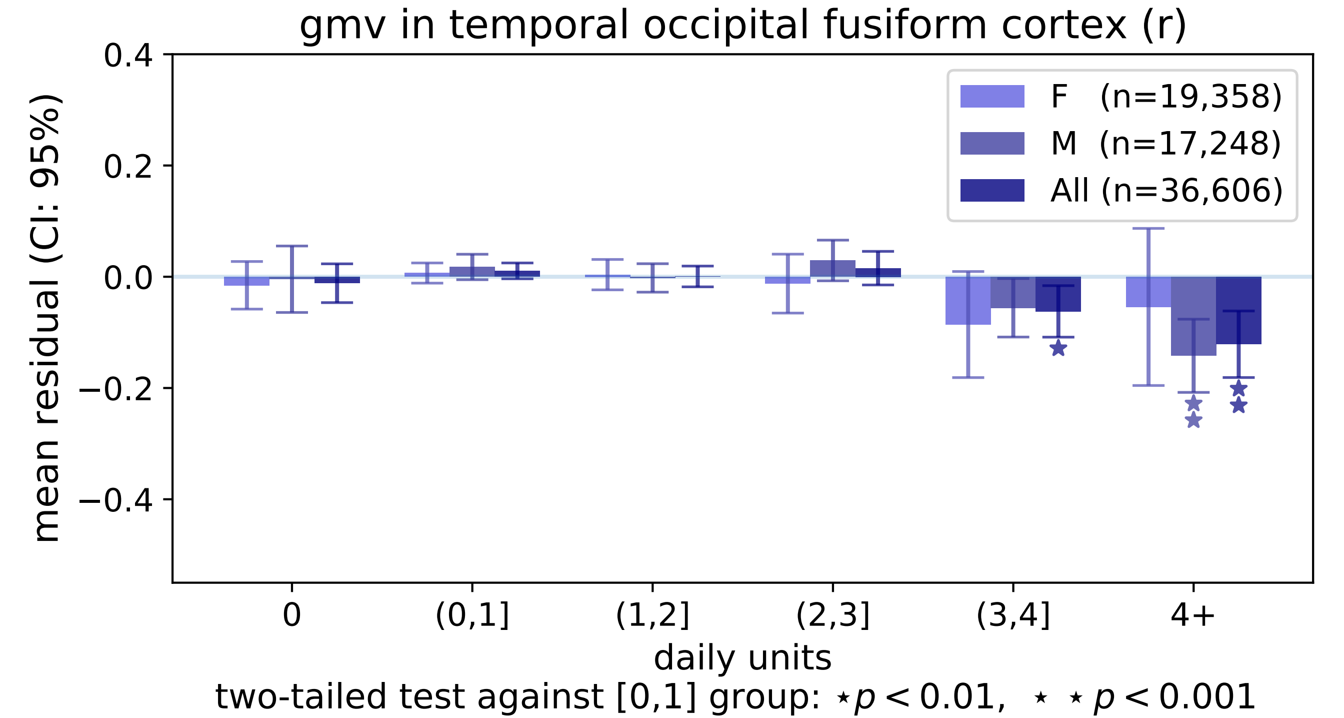
gmv in temporal occipital fusiform cortex (r)
0.4
0.2
0.0
−0.2
−0.4
mean residual (CI: 95%)
F (n=19,358)
M (n=17,248)
All (n=36,606)
0
(0,1]
(1,2]
(2,3]
(3,4]
4+
daily units
two-tailed test against [0,1] group: ⋆ p < 0.01, ⋆ ⋆ p < 0.001

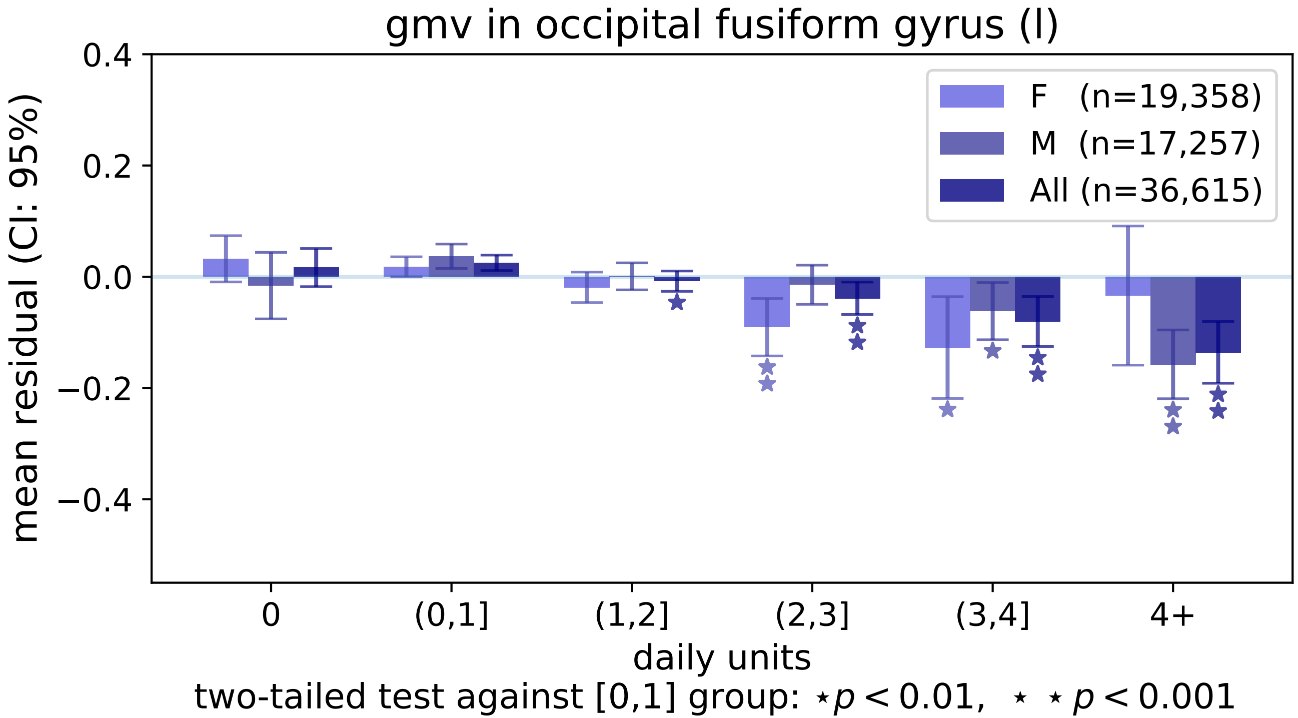

gmv in occipital fusiform gyrus (l)
mean residual (CI: 95%)
0.4
0.2
0.0
−0.2
−0.4
F (n=19,358)
M (n=17,257)
All (n=36,615)
0
(0,1]
(1,2]
(2,3]
(3,4]
4+
daily units
two-tailed test against [0,1] group: ⋆p < 0.01, ⋆ ⋆ p < 0.001

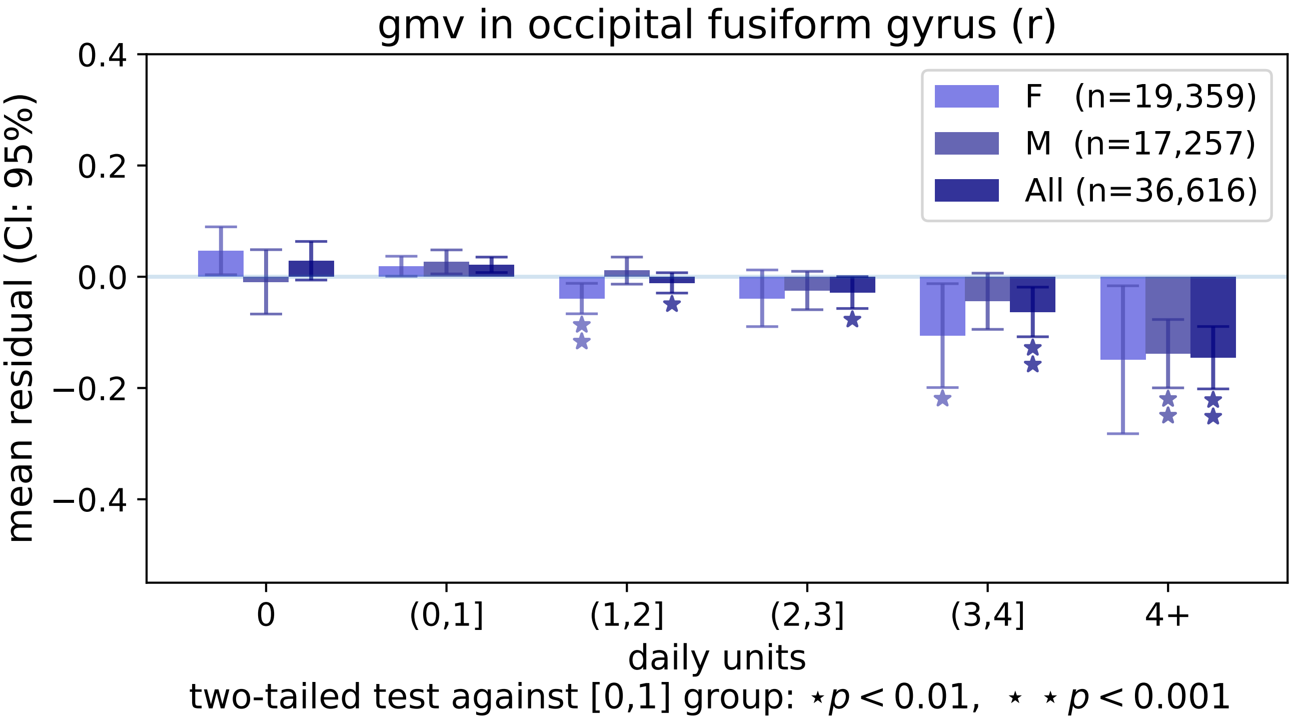

gmv in occipital fusiform gyrus (r)
mean residual (CI: 95%)
0.4
0.2
0.0
−0.2
−0.4
F (n=19,359)
M (n=17,257)
All (n=36,616)
0
(0,1]
(1,2]
(2,3]
(3,4]
4+
daily units
two-tailed test against [0,1] group: ⋆p < 0.01, ⋆ ⋆p < 0.001

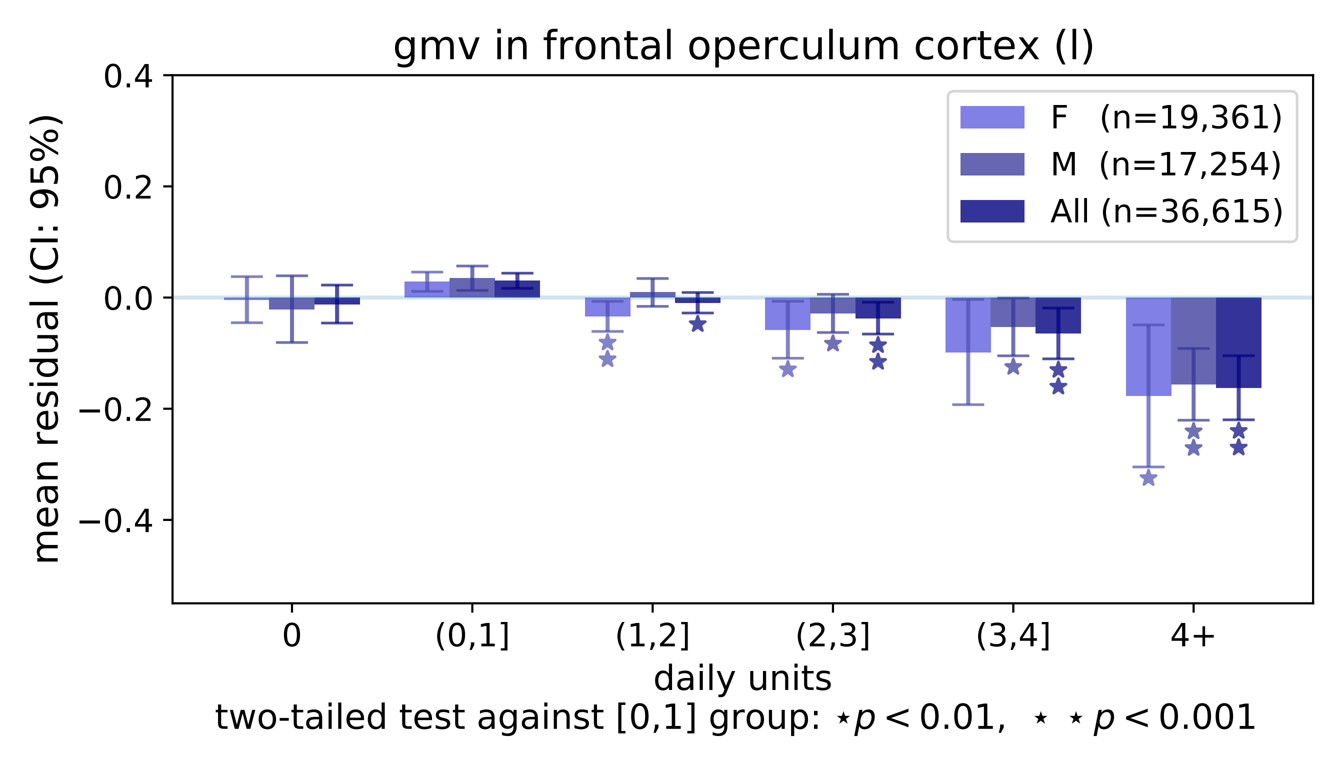
gmv in frontal operculum cortex (l)
mean residual (CI: 95%)
0.4
0.2
0.0
−0.2
−0.4
F (n=19,361)
M (n=17,254)
All (n=36,615)
0
(0,1]
(1,2]
(2,3]
(3,4]
4+
daily units
two-tailed test against [0,1] group: ⋆p < 0.01, ⋆ ⋆ p < 0.001

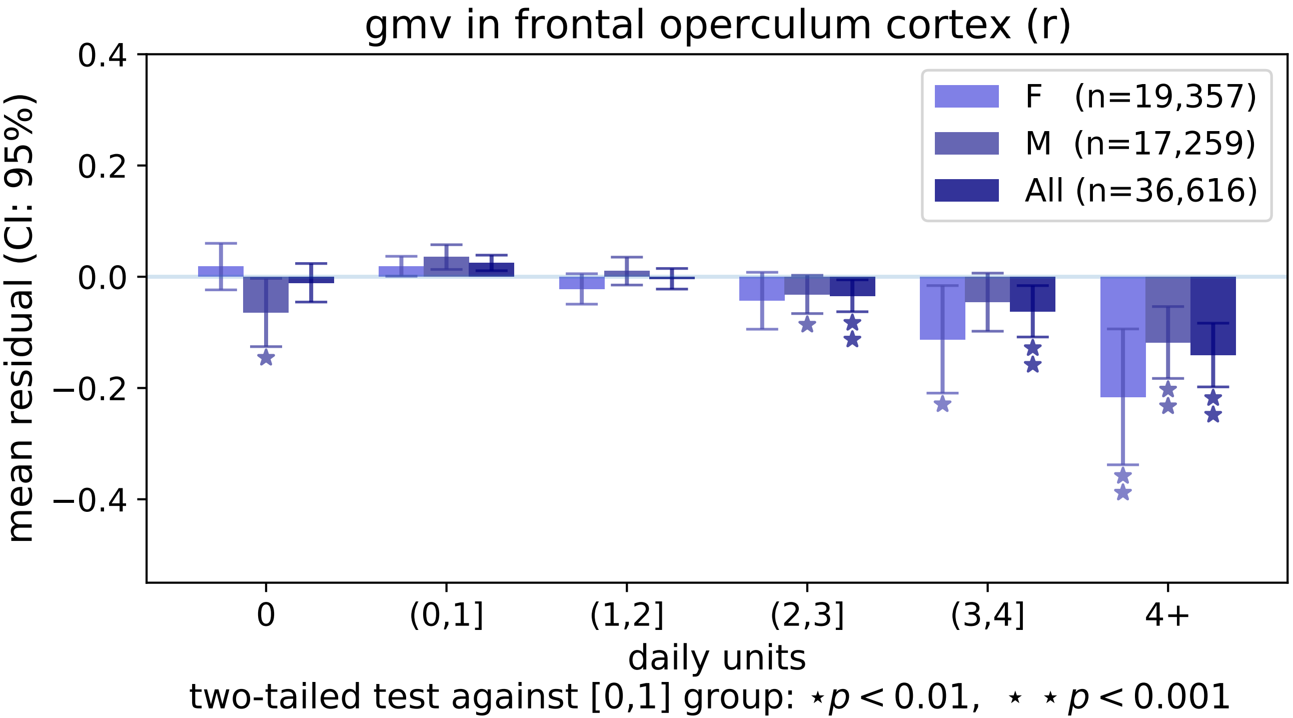

gmv in frontal operculum cortex (r)
F (n=19,357)
M (n=17,259)
All (n=36,616)
mean residual (CI: 95%)
0.4
0.2
0.0
−0.2
−0.4
0
(0,1]
(1,2]
(2,3]
(3,4]
4+
daily units
two-tailed test against [0,1] group: ⋆p < 0.01, ⋆ ⋆ p < 0.001

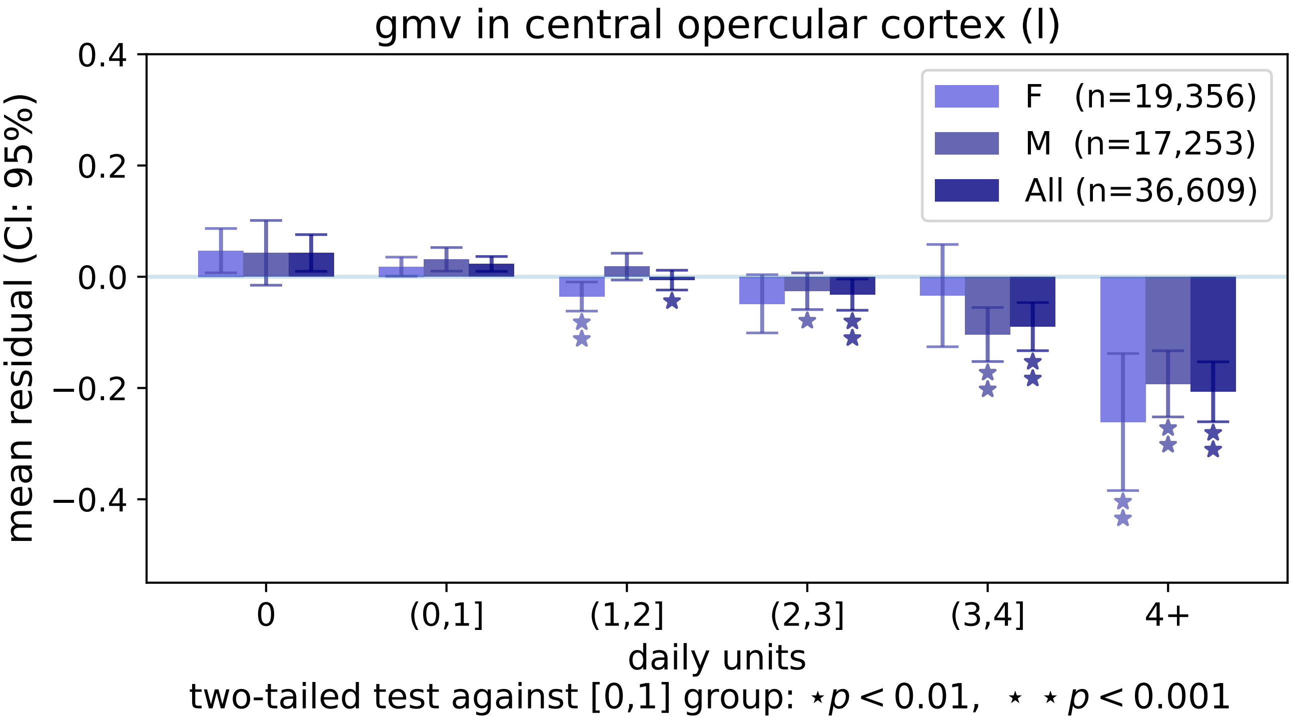

gmv in central opercular cortex (l)
0.4
0.2
0.0
−0.2
−0.4
mean residual (CI: 95%)
F (n=19,356)
M (n=17,253)
All (n=36,609)
0
(0,1]
(1,2]
(2,3]
(3,4]
4+
daily units
two-tailed test against [0,1] group: ⋆p < 0.01, ⋆ ⋆ p < 0.001

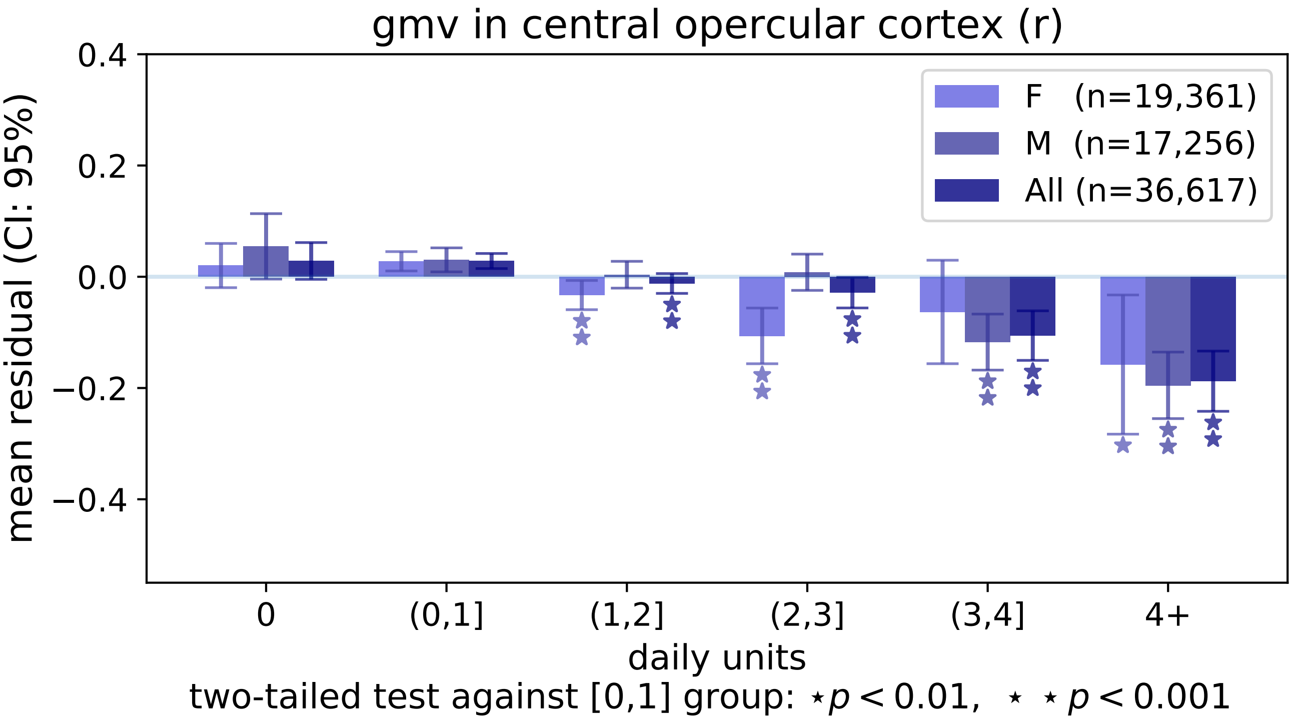

gmv in central opercular cortex (r)
0.4
0.2
0.0
−0.2
−0.4
mean residual (CI: 95%)
F (n=19,361)
M (n=17,256)
All (n=36,617)
0
(0,1]
(1,2]
(2,3]
(3,4]
4+
daily units
two-tailed test against [0,1] group: ⋆ p < 0.01, ⋆ ⋆ p < 0.001

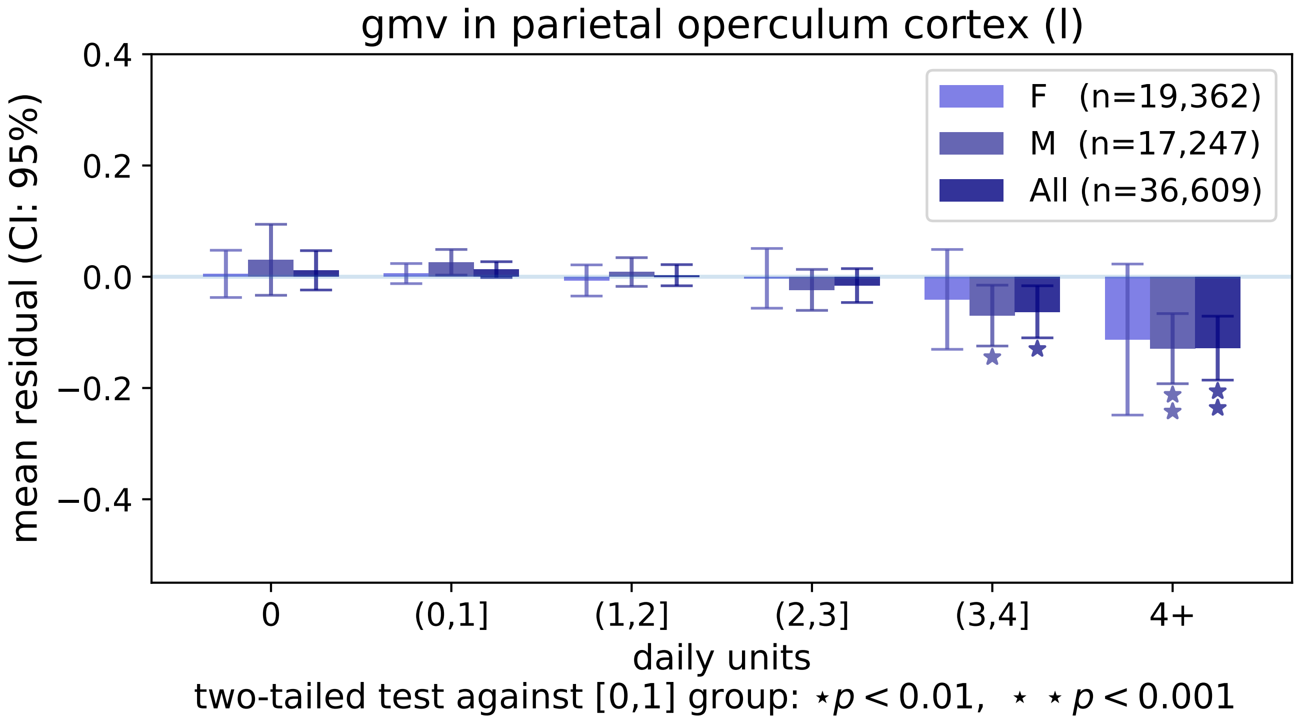
gmv in parietal operculum cortex (l)
mean residual (CI: 95%)
0.4
0.2
0.0
−0.2
−0.4
F (n=19,362)
M (n=17,247)
All (n=36,609)
0
(0,1]
(1,2]
(2,3]
(3,4]
4+
daily units
two-tailed test against [0,1] group: ⋆ $p < 0.01$, ⋆ ⋆ $p < 0.001$

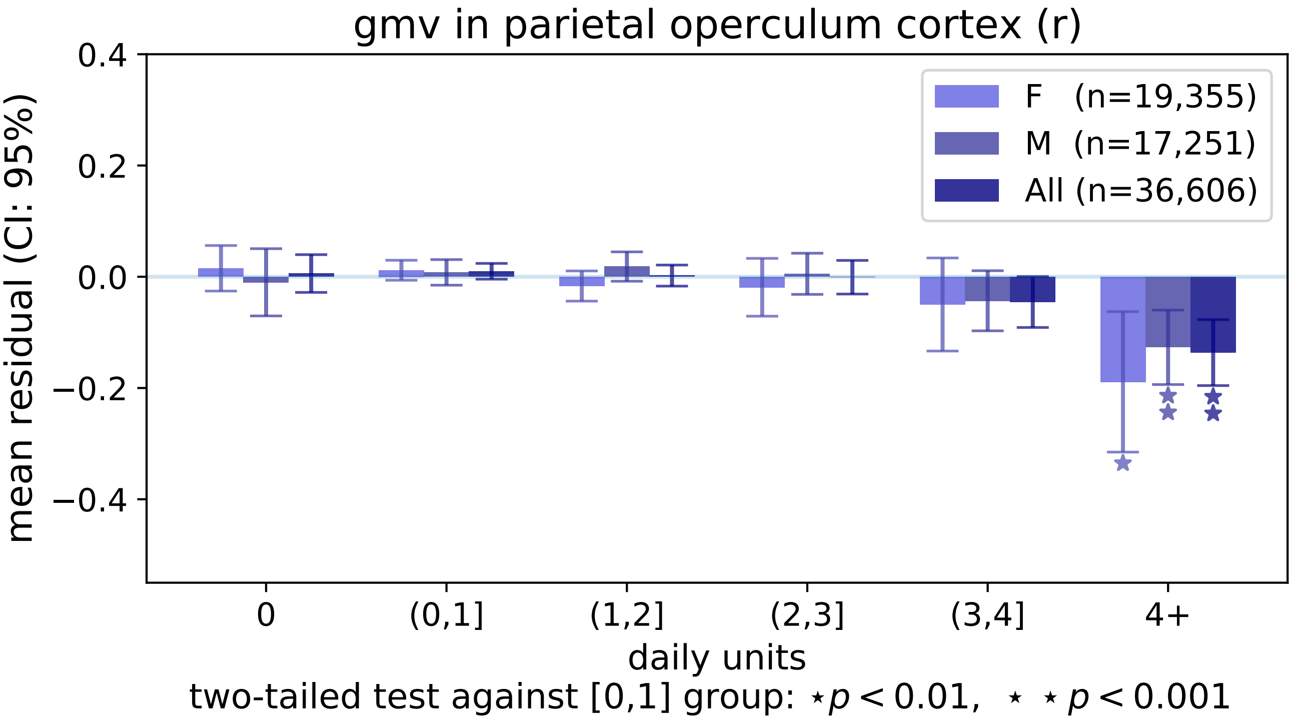
gmv in parietal operculum cortex (r)
F (n=19,355)
M (n=17,251)
All (n=36,606)
mean residual (CI: 95%)
0.4
0.2
0.0
−0.2
−0.4
0
(0,1]
(1,2]
(2,3]
(3,4]
4+
daily units
two-tailed test against [0,1] group: ⋆p < 0.01, ⋆ ⋆ p < 0.001

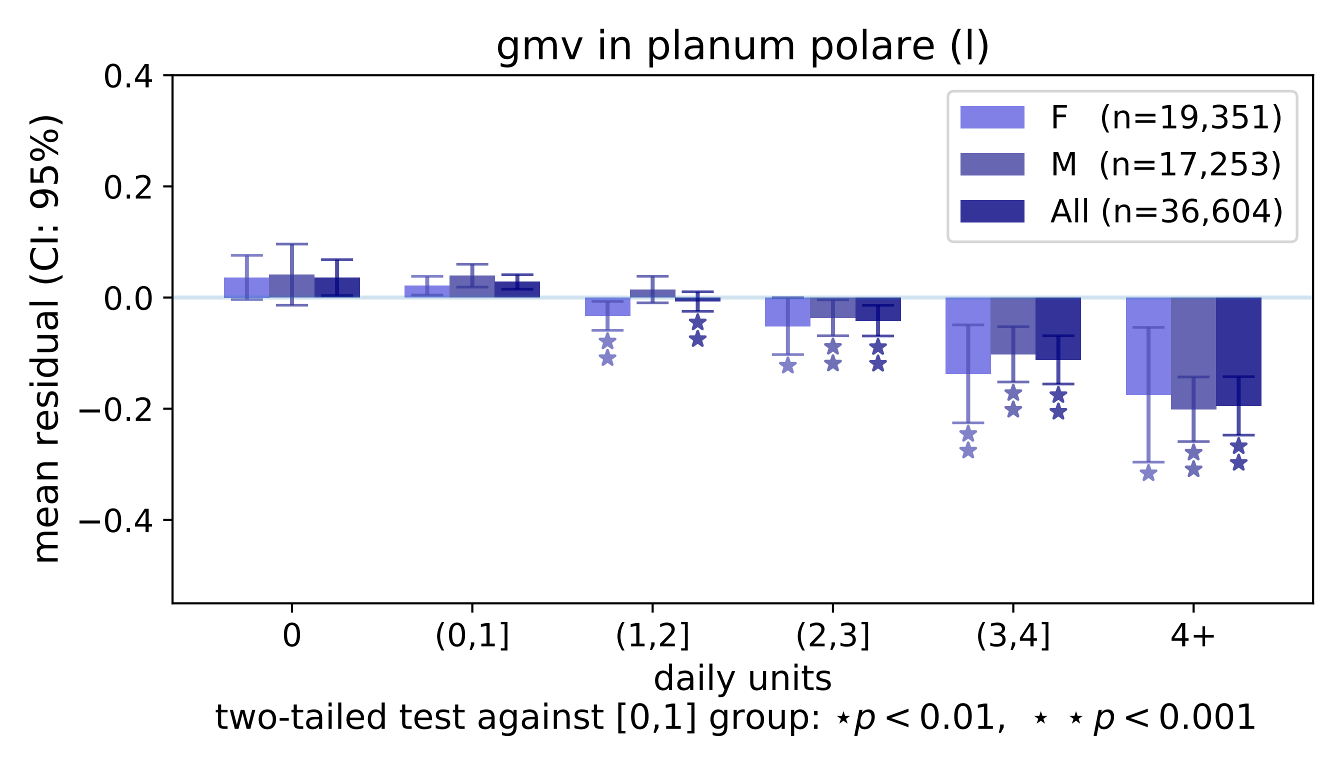

gmv in planum polare (l)
mean residual (CI: 95%)
0.4
0.2
0.0
−0.2
−0.4
F (n=19,351)
M (n=17,253)
All (n=36,604)
0
(0,1]
(1,2]
(2,3]
(3,4]
4+
daily units
two-tailed test against [0,1] group: ⋆p < 0.01, ⋆ ⋆ p < 0.001

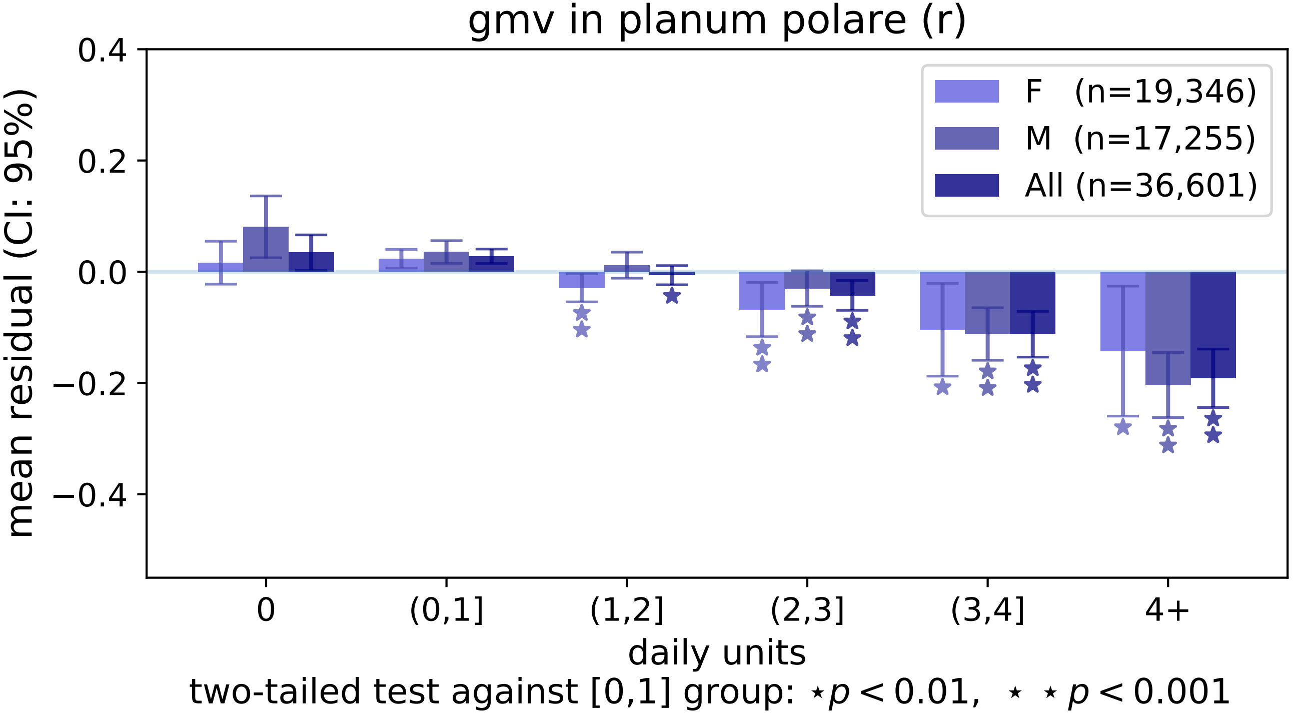
gmv in planum polare (r)
mean residual (CI: 95%)
0.4
0.2
0.0
−0.2
−0.4
F (n=19,346)
M (n=17,255)
All (n=36,601)
0
(0,1]
(1,2]
(2,3]
(3,4]
4+
daily units
two-tailed test against [0,1] group: ⋆p < 0.01, ⋆ ⋆ p < 0.001

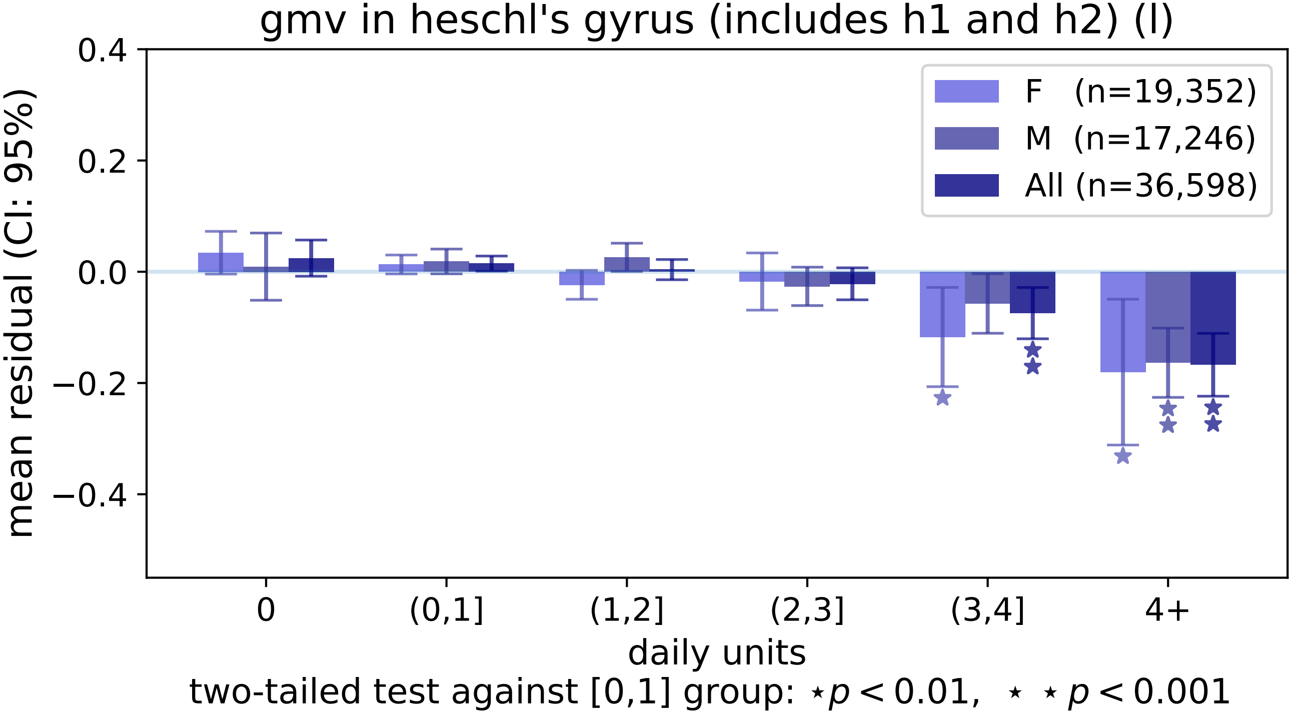

gmv in heschl's gyrus (includes h1 and h2) (l)
0.4
0.2
0.0
−0.2
−0.4
mean residual (CI: 95%)
F (n=19,352)
M (n=17,246)
All (n=36,598)
0
(0,1]
(1,2]
(2,3]
(3,4]
4+
daily units
two-tailed test against [0,1] group: ⋆p < 0.01, ⋆ ⋆ p < 0.001

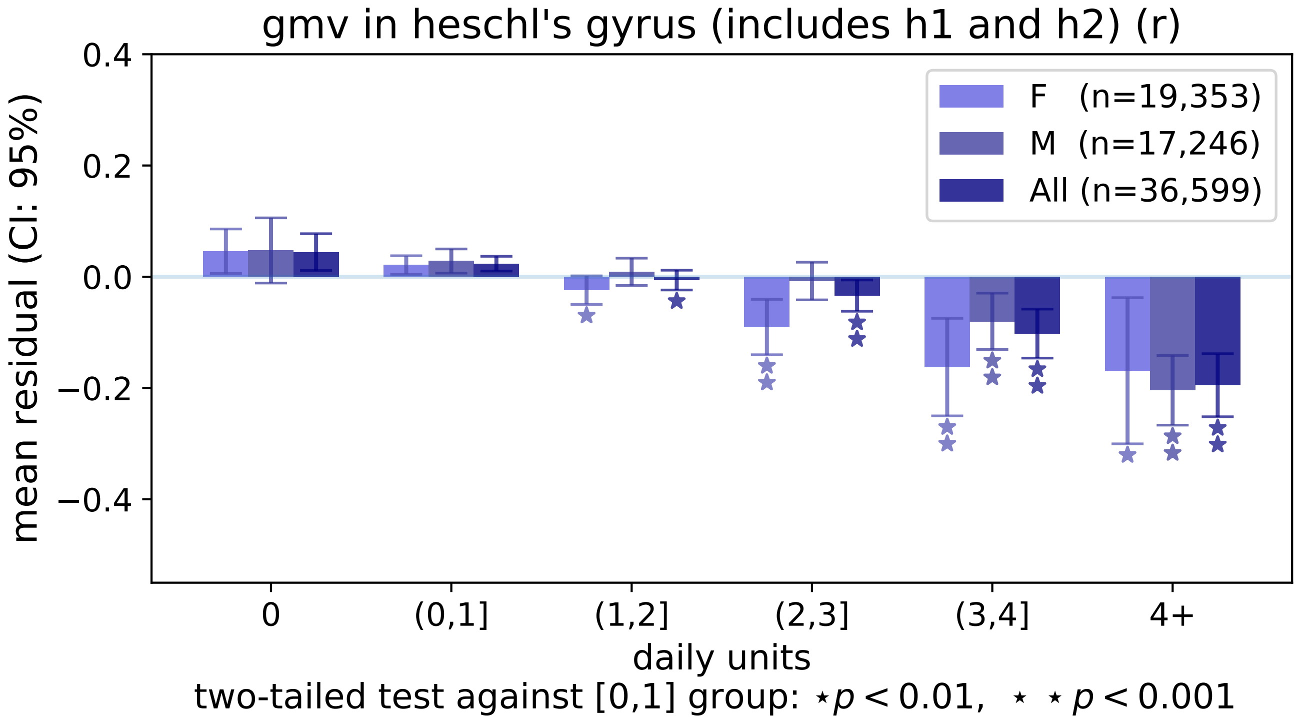
gmv in heschl's gyrus (includes h1 and h2) (r)
F (n=19,353)
M (n=17,246)
All (n=36,599)
mean residual (CI: 95%)
0.4
0.2
0.0
−0.2
−0.4
0
(0,1]
(1,2]
(2,3]
(3,4]
4+
daily units
two-tailed test against [0,1] group: ⋆ p < 0.01, ⋆ ⋆ p < 0.001

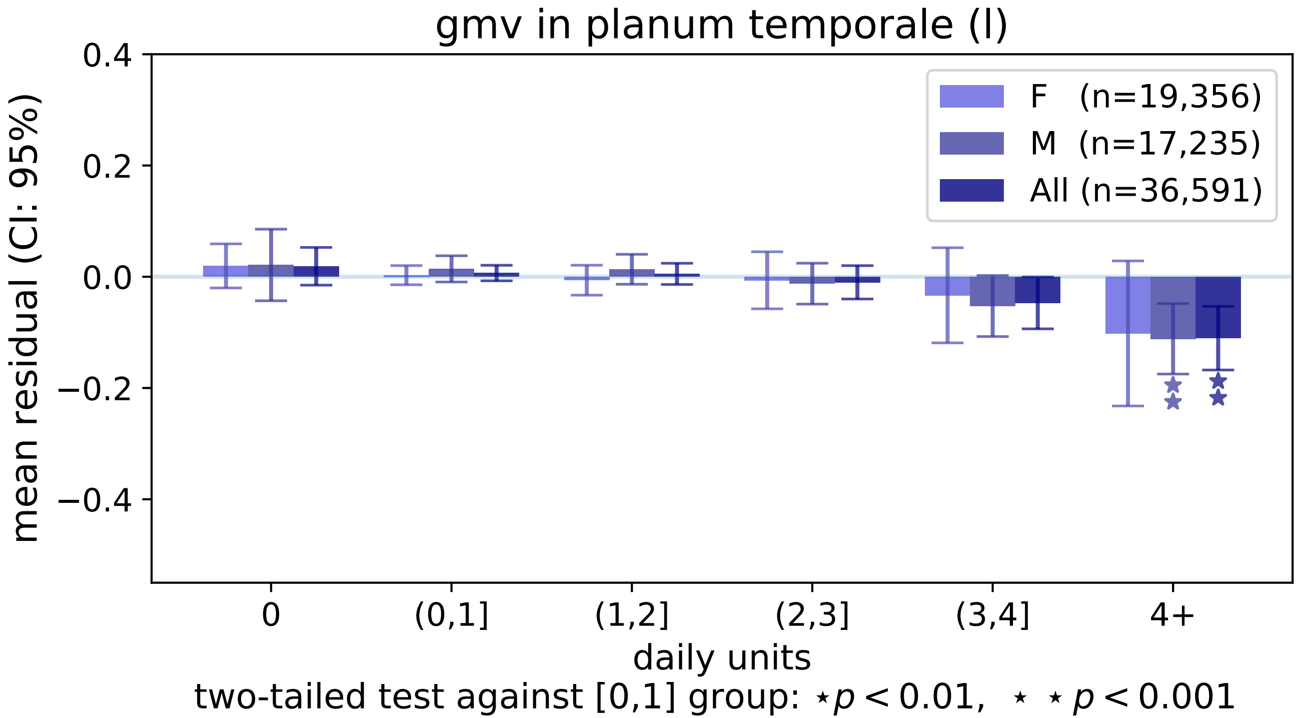
gmv in planum temporale (l)
0.4
0.2
0.0
−0.2
−0.4
mean residual (CI: 95%)
F (n=19,356)
M (n=17,235)
All (n=36,591)
0
(0,1]
(1,2]
(2,3]
(3,4]
4+
daily units
two-tailed test against [0,1] group: ⋆p < 0.01, ⋆ ⋆ p < 0.001

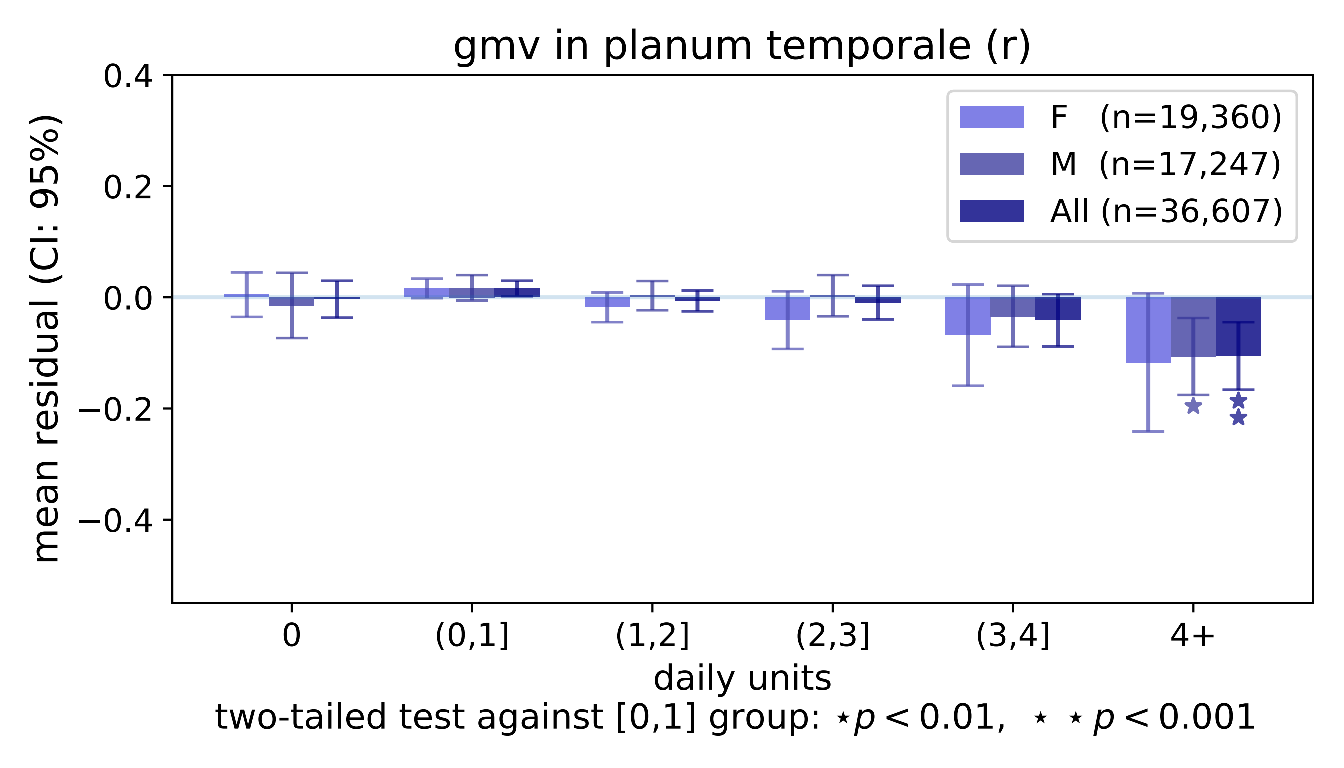
gmv in planum temporale (r)
0.4
0.2
0.0
−0.2
−0.4
mean residual (CI: 95%)
F (n=19,360)
M (n=17,247)
All (n=36,607)
0
(0,1]
(1,2]
(2,3]
(3,4]
4+
daily units
two-tailed test against [0,1] group: ⋆p < 0.01, ⋆ ⋆ p < 0.001

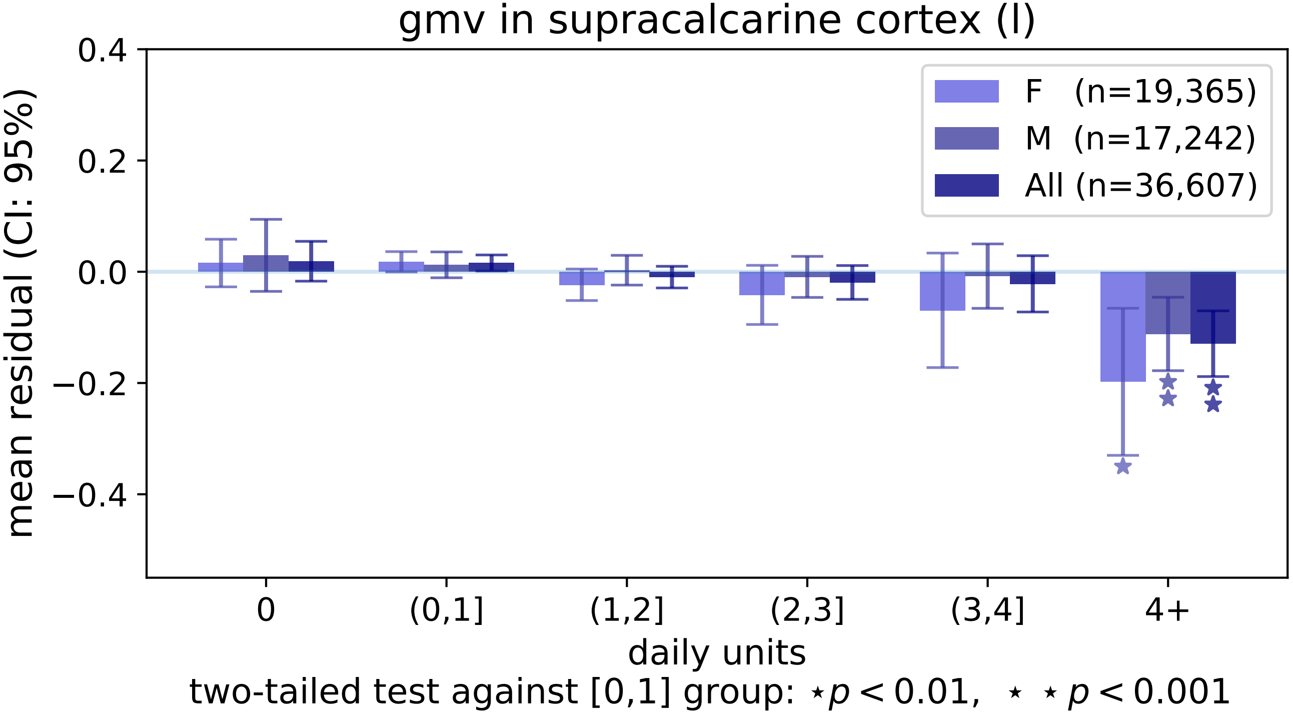
gmv in supracalcarine cortex (l)
mean residual (CI: 95%)
0.4
0.2
0.0
−0.2
−0.4
F (n=19,365)
M (n=17,242)
All (n=36,607)
0
(0,1]
(1,2]
(2,3]
(3,4]
4+
daily units
two-tailed test against [0,1] group: ⋆p < 0.01, ⋆ ⋆ p < 0.001

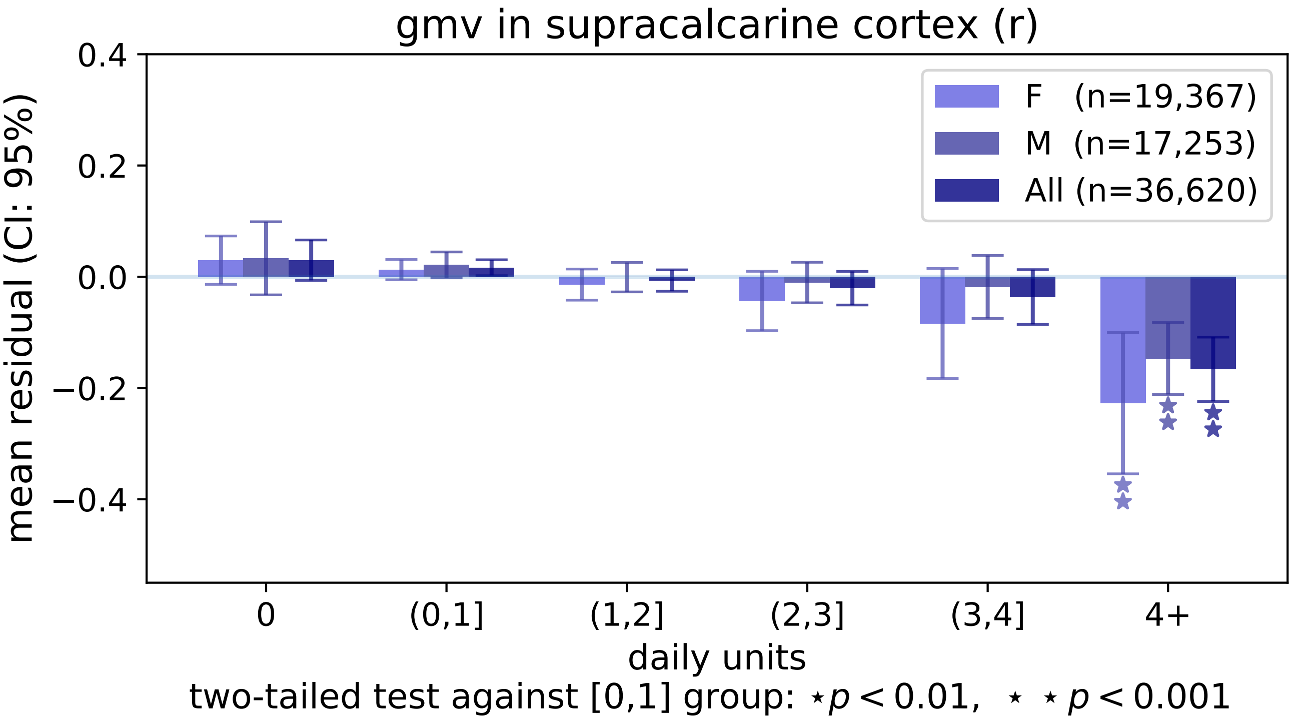
gmv in supracalcarine cortex (r)
mean residual (CI: 95%)
0.4
0.2
0.0
−0.2
−0.4
F (n=19,367)
M (n=17,253)
All (n=36,620)
0
(0,1]
(1,2]
(2,3]
(3,4]
4+
daily units
two-tailed test against [0,1] group: ⋆ $p < 0.01$, ⋆ ⋆ $p < 0.001$

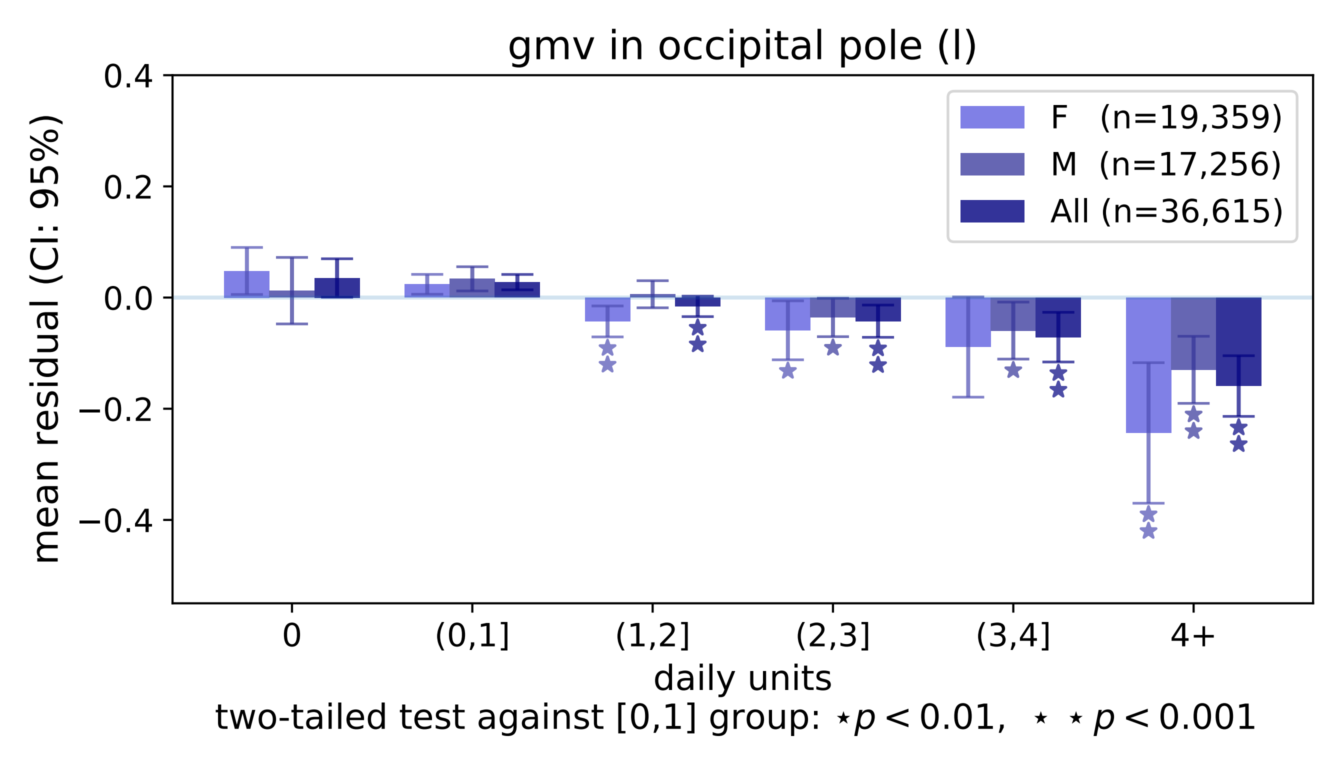
gmv in occipital pole (l)
0.4
0.2
0.0
−0.2
−0.4
mean residual (CI: 95%)
F (n=19,359)
M (n=17,256)
All (n=36,615)
0
(0,1]
(1,2]
(2,3]
(3,4]
4+
daily units
two-tailed test against [0,1] group: ⋆p < 0.01, ⋆ ⋆p < 0.001

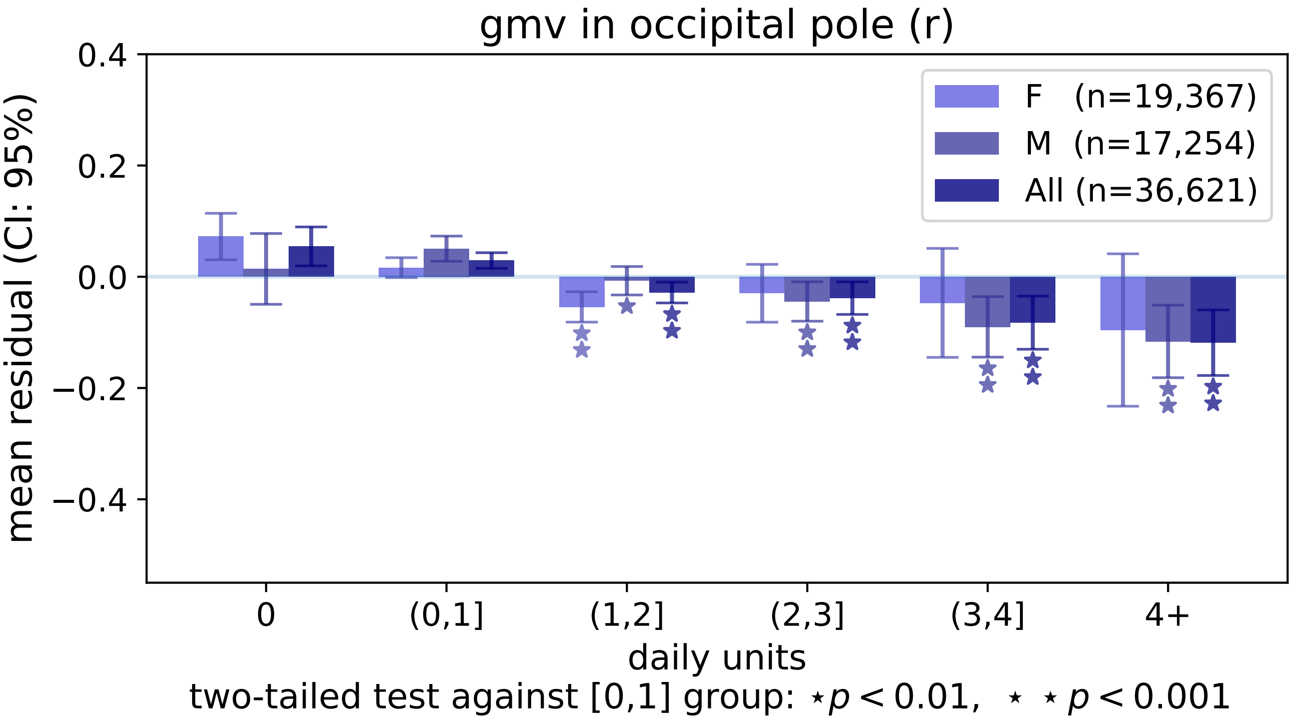

gmv in occipital pole (r)
0.4
0.2
0.0
−0.2
−0.4
mean residual (CI: 95%)
F (n=19,367)
M (n=17,254)
All (n=36,621)
0
(0,1]
(1,2]
(2,3]
(3,4]
4+
daily units
two-tailed test against [0,1] group: ⋆p < 0.01, ⋆ ⋆ p < 0.001

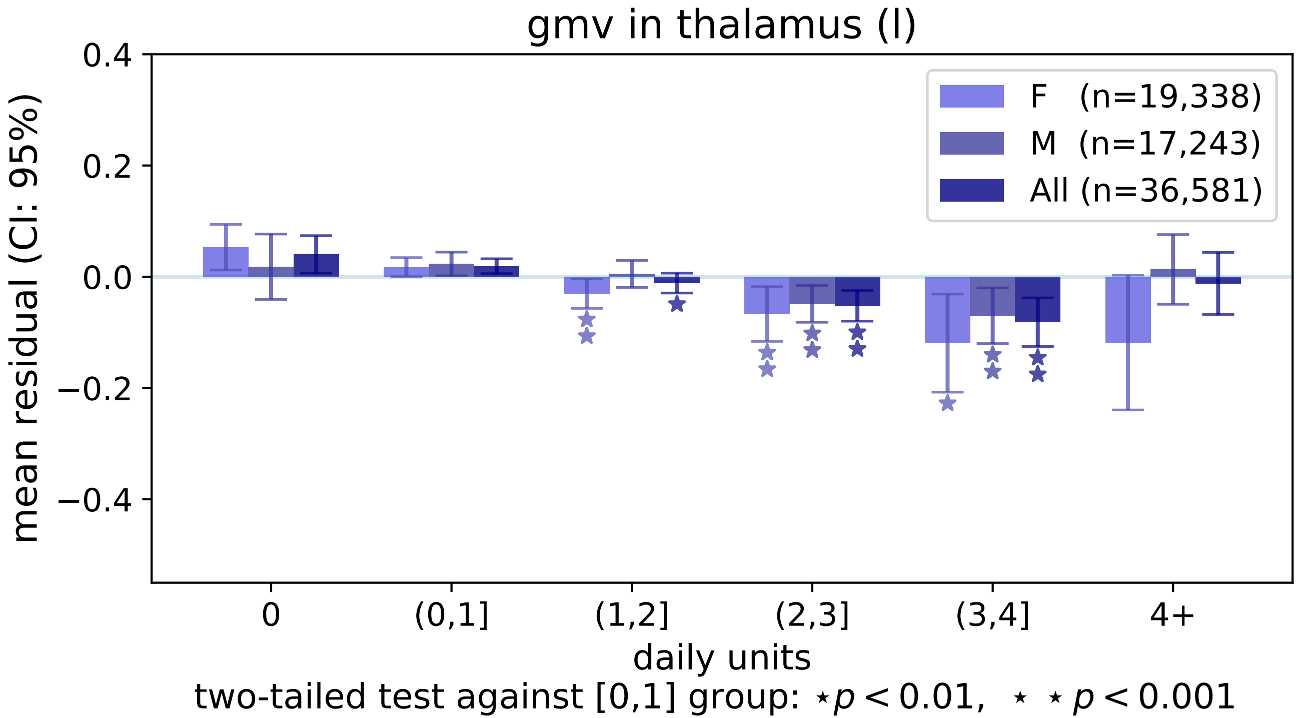
gmv in thalamus (l)
F (n=19,338)
M (n=17,243)
All (n=36,581)
mean residual (CI: 95%)
0.4
0.2
0.0
−0.2
−0.4
0
(0,1]
(1,2]
(2,3]
(3,4]
4+
daily units
two-tailed test against [0,1] group: ⋆p < 0.01, ⋆ ⋆ p < 0.001

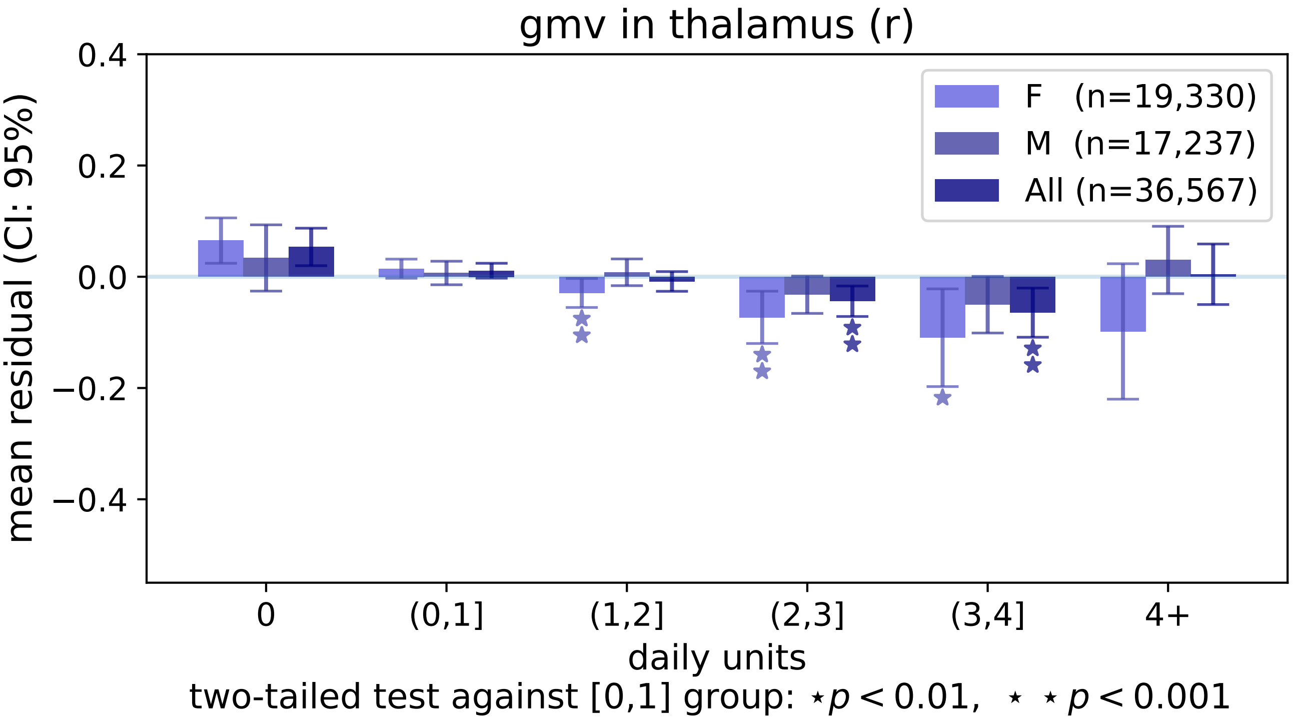

gmv in thalamus (r)
0.4
0.2
0.0
−0.2
−0.4
mean residual (CI: 95%)
F (n=19,330)
M (n=17,237)
All (n=36,567)
0
(0,1]
(1,2]
(2,3]
(3,4]
4+
daily units
two-tailed test against [0,1] group: ⋆p < 0.01, ⋆ ⋆ p < 0.001

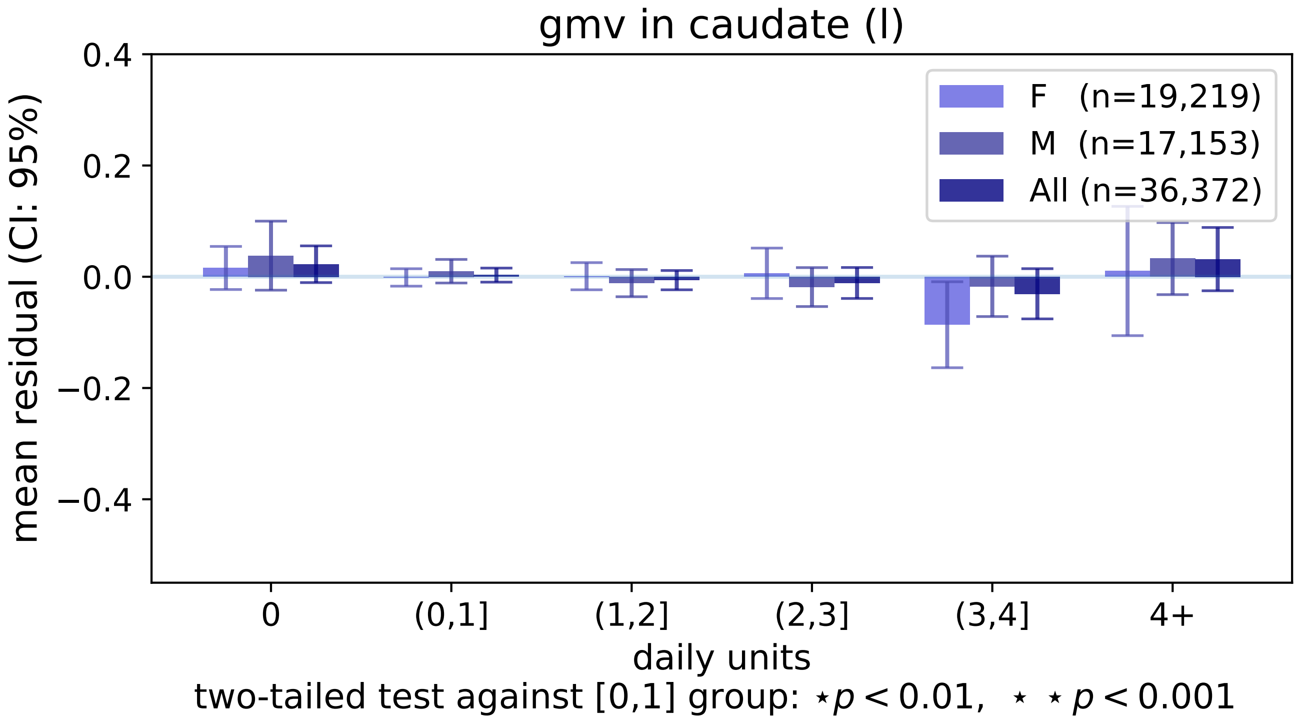

gmv in caudate (l)
0.4
0.2
0.0
−0.2
−0.4
mean residual (CI: 95%)
F (n=19,219)
M (n=17,153)
All (n=36,372)
0
(0,1]
(1,2]
(2,3]
(3,4]
4+
daily units
two-tailed test against [0,1] group: ⋆ p < 0.01, ⋆ ⋆ p < 0.001

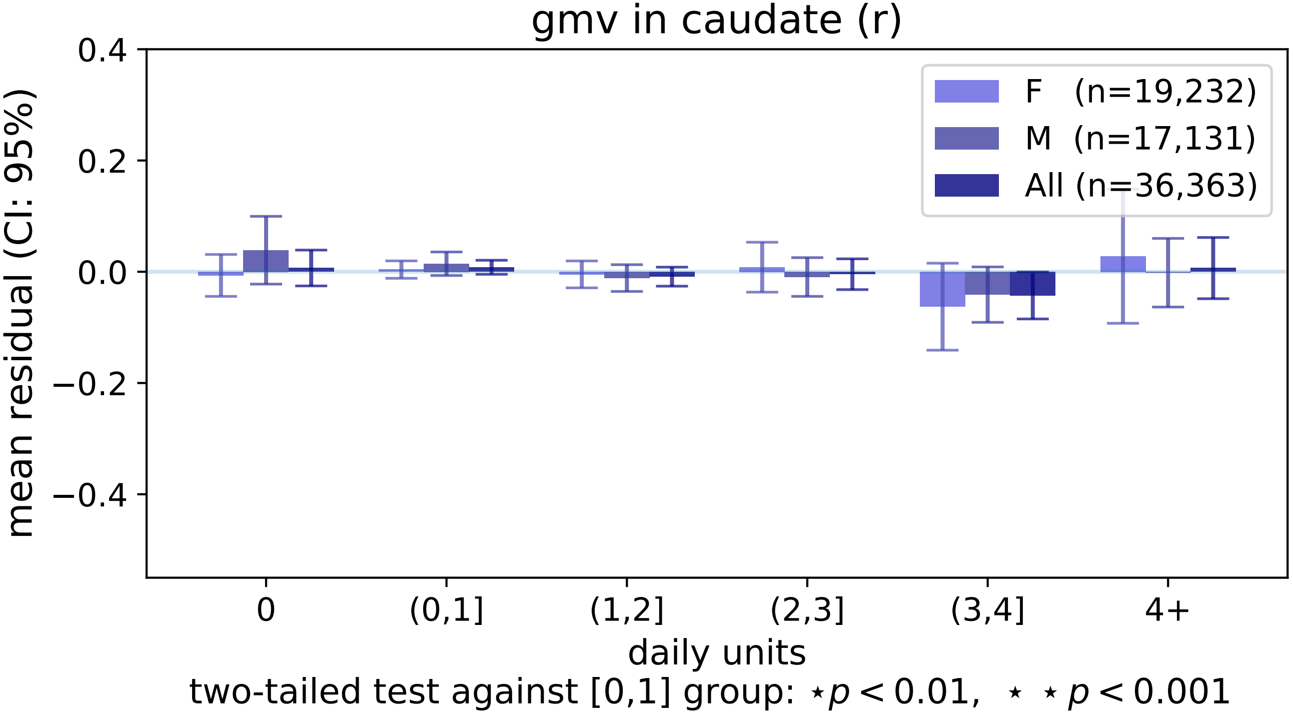

gmv in caudate (r)
0.4
0.2
0.0
−0.2
−0.4
mean residual (CI: 95%)
F (n=19,232)
M (n=17,131)
All (n=36,363)
0
(0,1]
(1,2]
(2,3]
(3,4]
4+
daily units
two-tailed test against [0,1] group: ⋆p < 0.01, ⋆ ⋆ p < 0.001

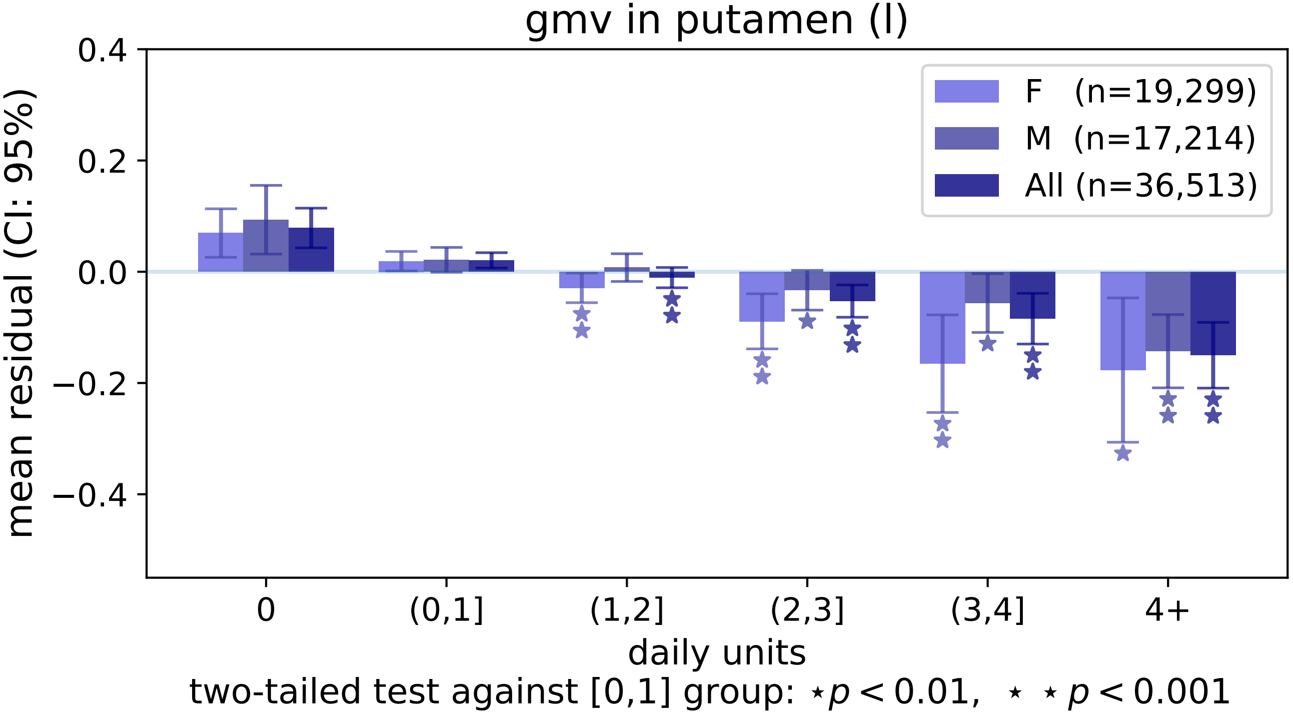

gmv in putamen (l)
mean residual (CI: 95%)
0.4
0.2
0.0
−0.2
−0.4
F (n=19,299)
M (n=17,214)
All (n=36,513)
0
(0,1]
(1,2]
(2,3]
(3,4]
4+
daily units
two-tailed test against [0,1] group: ⋆ $p < 0.01$, ⋆ ⋆ $p < 0.001$

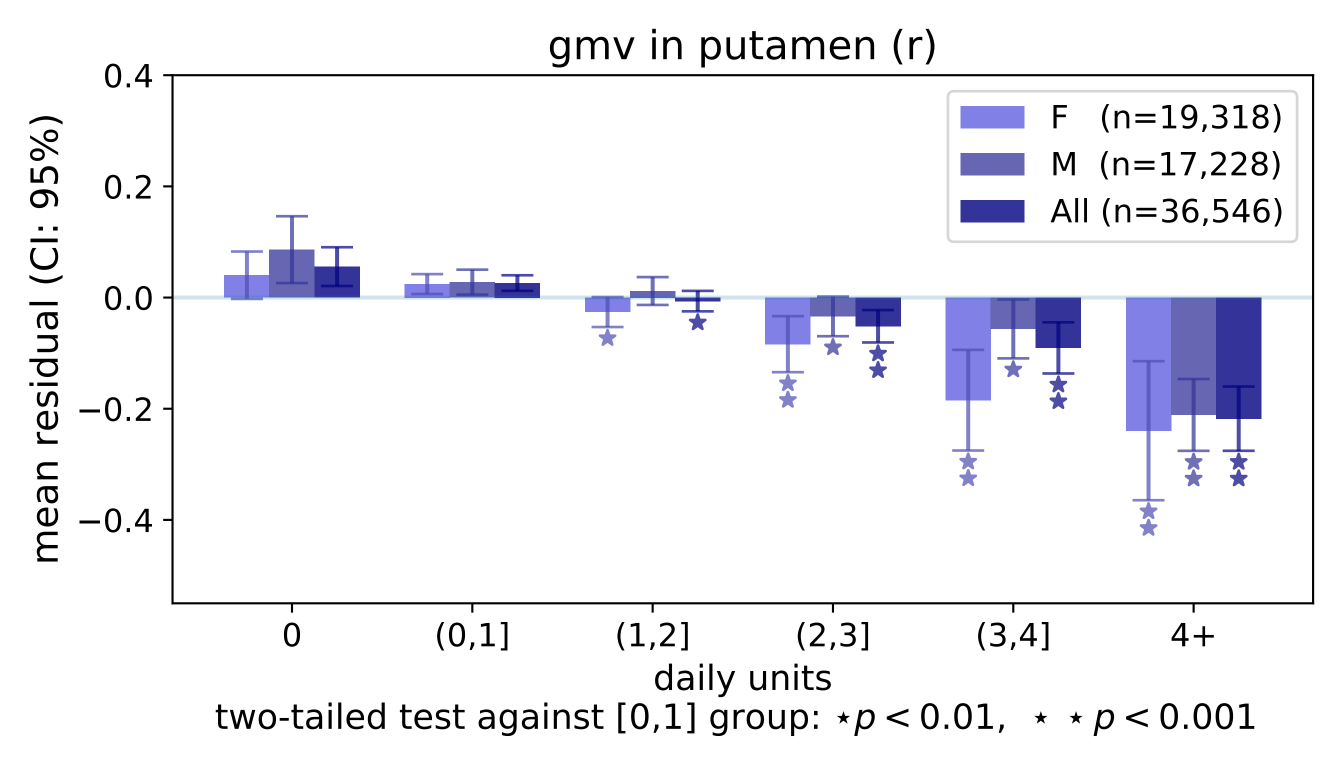
gmv in putamen (r)
mean residual (CI: 95%)
0.4
0.2
0.0
−0.2
−0.4
F (n=19,318)
M (n=17,228)
All (n=36,546)
0
(0,1]
(1,2]
(2,3]
(3,4]
4+
daily units
two-tailed test against [0,1] group: ⋆ $p < 0.01$, ⋆ ⋆ $p < 0.001$

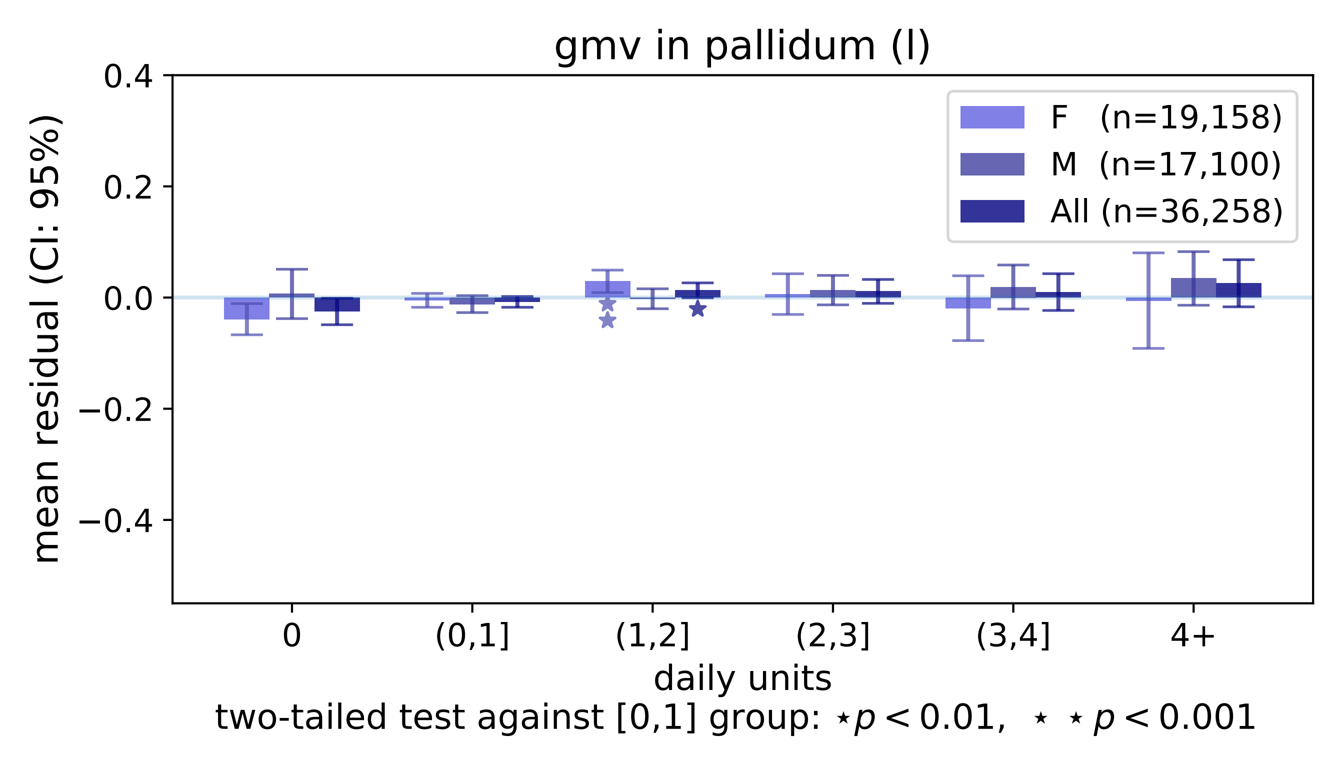

gmv in pallidum (l)
0.4
0.2
0.0
−0.2
−0.4
mean residual (CI: 95%)
F (n=19,158)
M (n=17,100)
All (n=36,258)
0
(0,1]
(1,2]
(2,3]
(3,4]
4+
daily units
two-tailed test against [0,1] group: ⋆p < 0.01, ⋆ ⋆ p < 0.001

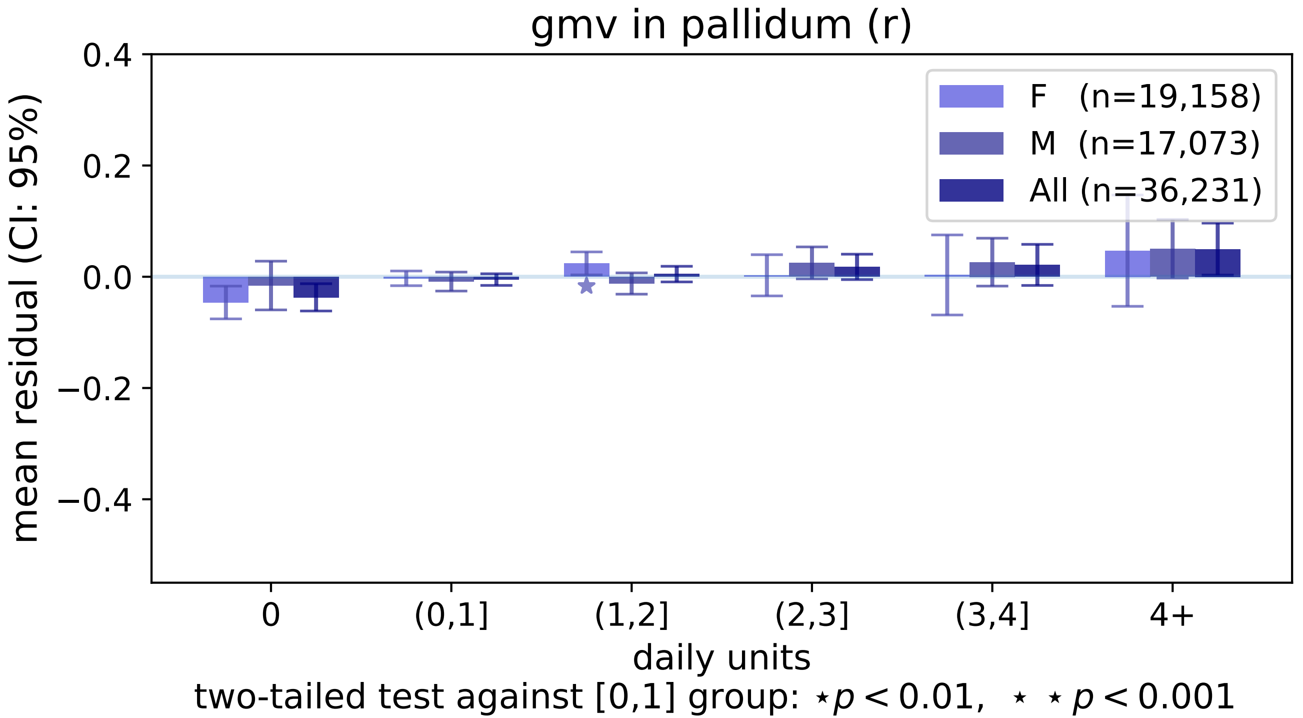

gmv in pallidum (r)
0.4
0.2
0.0
−0.2
−0.4
mean residual (CI: 95%)
F (n=19,158)
M (n=17,073)
All (n=36,231)
0
(0,1]
(1,2]
(2,3]
(3,4]
4+
daily units
two-tailed test against [0,1] group: ⋆p < 0.01, ⋆ ⋆ p < 0.001

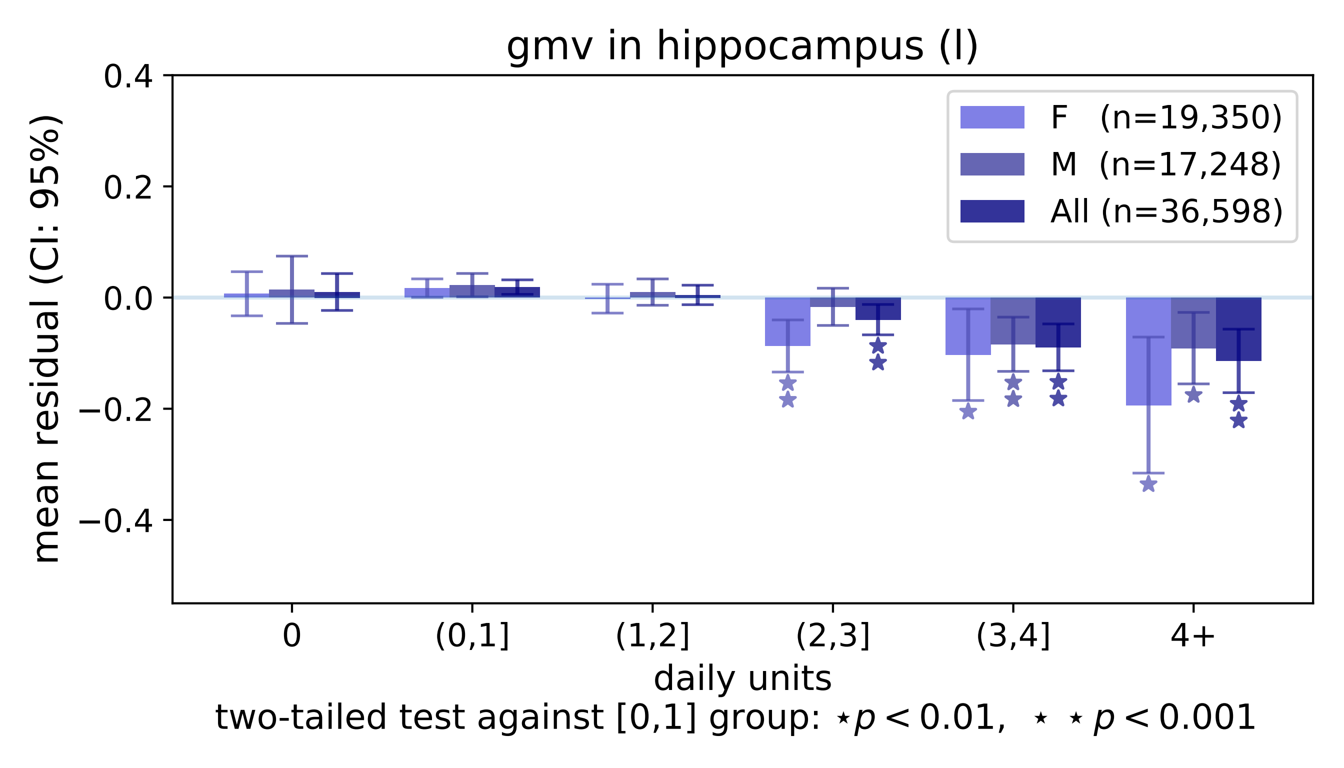

gmv in hippocampus (l)
mean residual (CI: 95%)
0.4
0.2
0.0
−0.2
−0.4
F (n=19,350)
M (n=17,248)
All (n=36,598)
0
(0,1]
(1,2]
(2,3]
(3,4]
4+
daily units
two-tailed test against [0,1] group: ⋆p < 0.01, ⋆ ⋆ p < 0.001

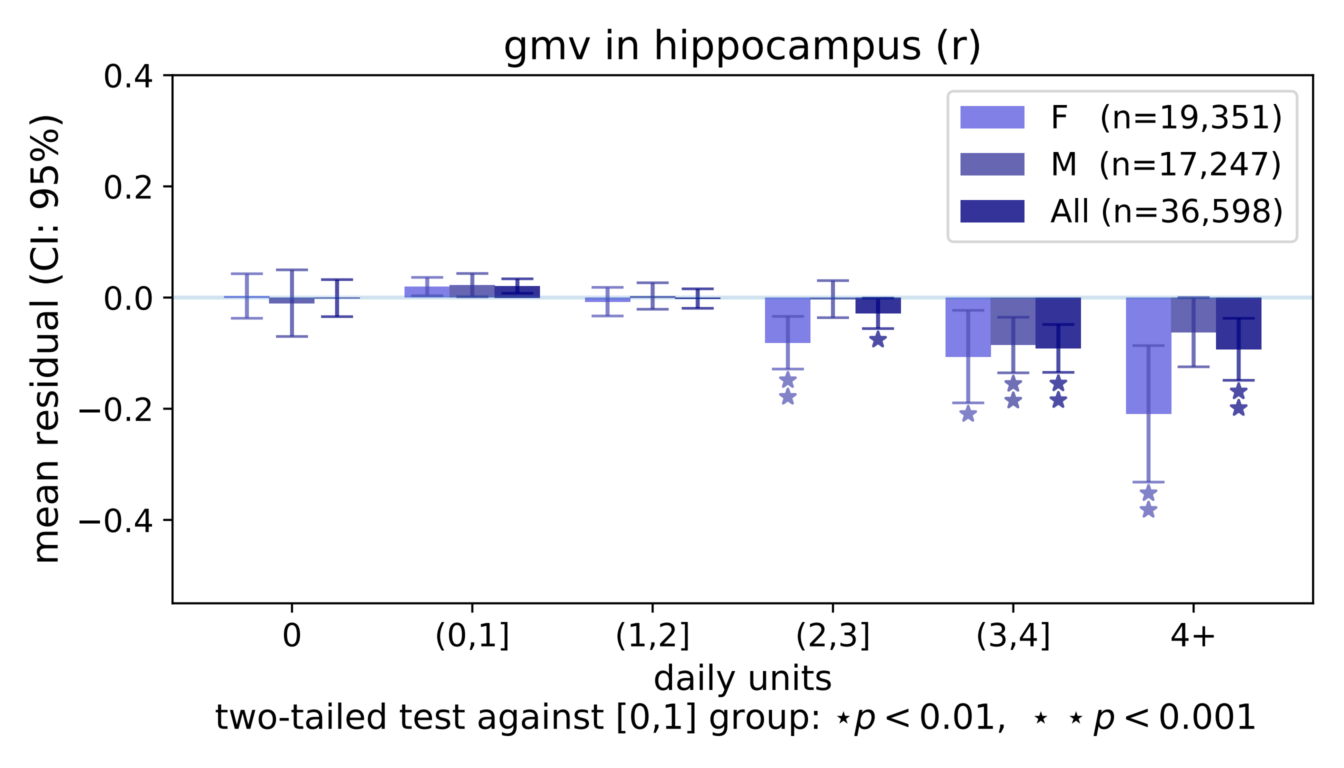

gmv in hippocampus (r)
F (n=19,351)
M (n=17,247)
All (n=36,598)
mean residual (CI: 95%)
0.4
0.2
0.0
−0.2
−0.4
0
(0,1]
(1,2]
(2,3]
(3,4]
4+
daily units
two-tailed test against [0,1] group: ⋆p < 0.01, ⋆ ⋆ p < 0.001

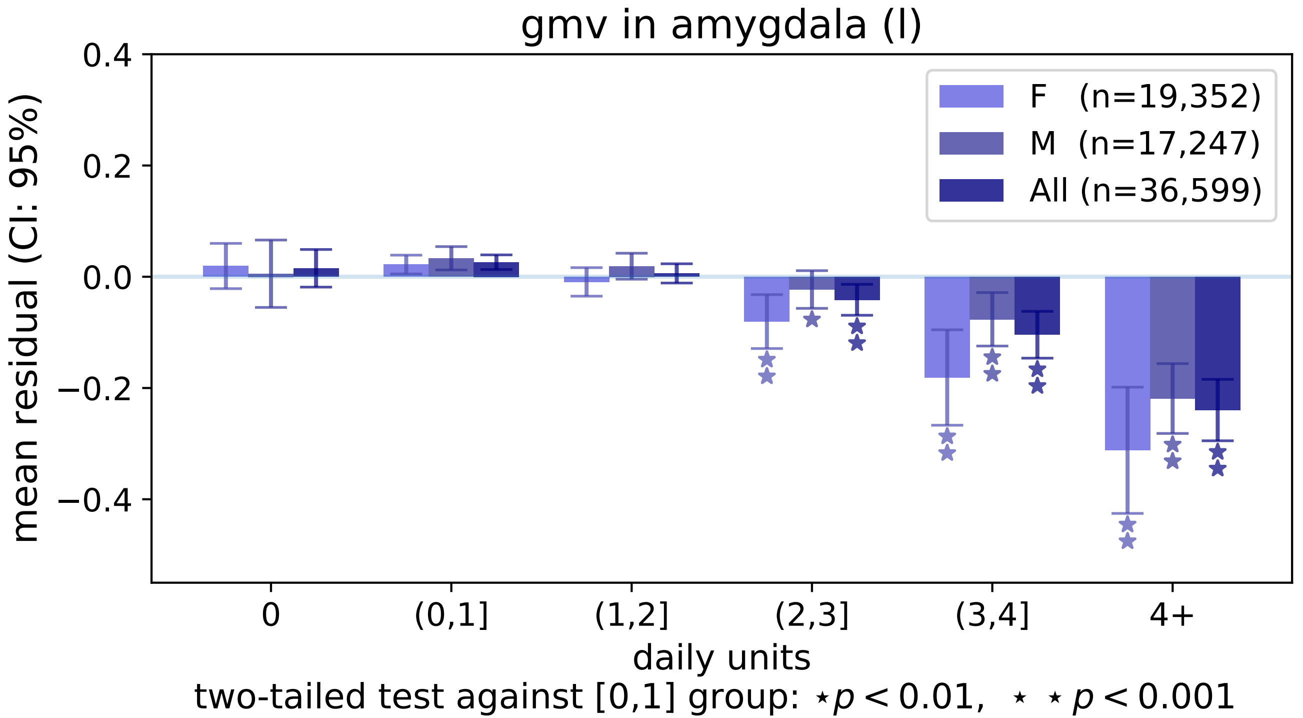
gmv in amygdala (l)
0.4
0.2
0.0
−0.2
−0.4
mean residual (CI: 95%)
F (n=19,352)
M (n=17,247)
All (n=36,599)
0
(0,1]
(1,2]
(2,3]
(3,4]
4+
daily units
two-tailed test against [0,1] group: ⋆p < 0.01, ⋆ ⋆ p < 0.001

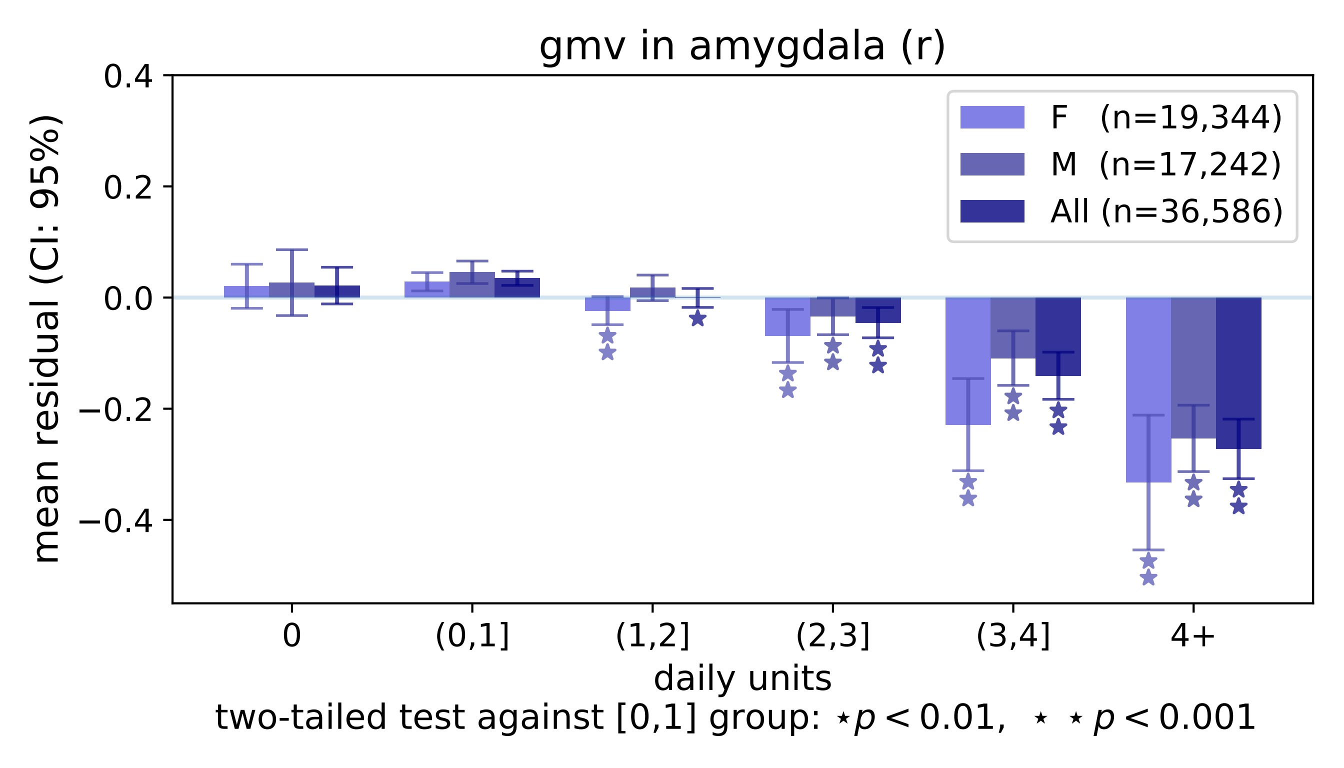
gmv in amygdala (r)
F (n=19,344)
M (n=17,242)
All (n=36,586)
mean residual (CI: 95%)
0.4
0.2
0.0
−0.2
−0.4
0
(0,1]
(1,2]
(2,3]
(3,4]
4+
daily units
two-tailed test against [0,1] group: ⋆p < 0.01, ⋆ ⋆ p < 0.001

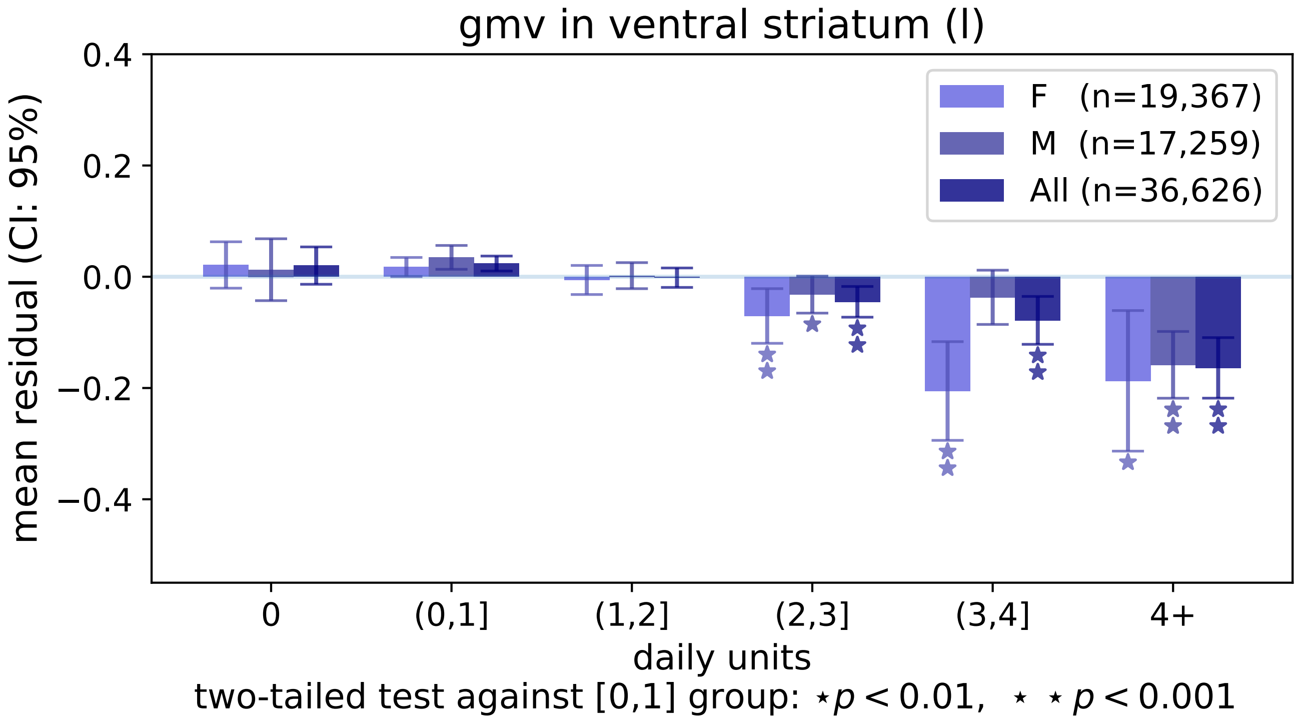

gmv in ventral striatum (l)
mean residual (CI: 95%)
0.4
0.2
0.0
−0.2
−0.4
F (n=19,367)
M (n=17,259)
All (n=36,626)
0
(0,1]
(1,2]
(2,3]
(3,4]
4+
daily units
two-tailed test against [0,1] group: ⋆ p < 0.01, ⋆ ⋆ p < 0.001

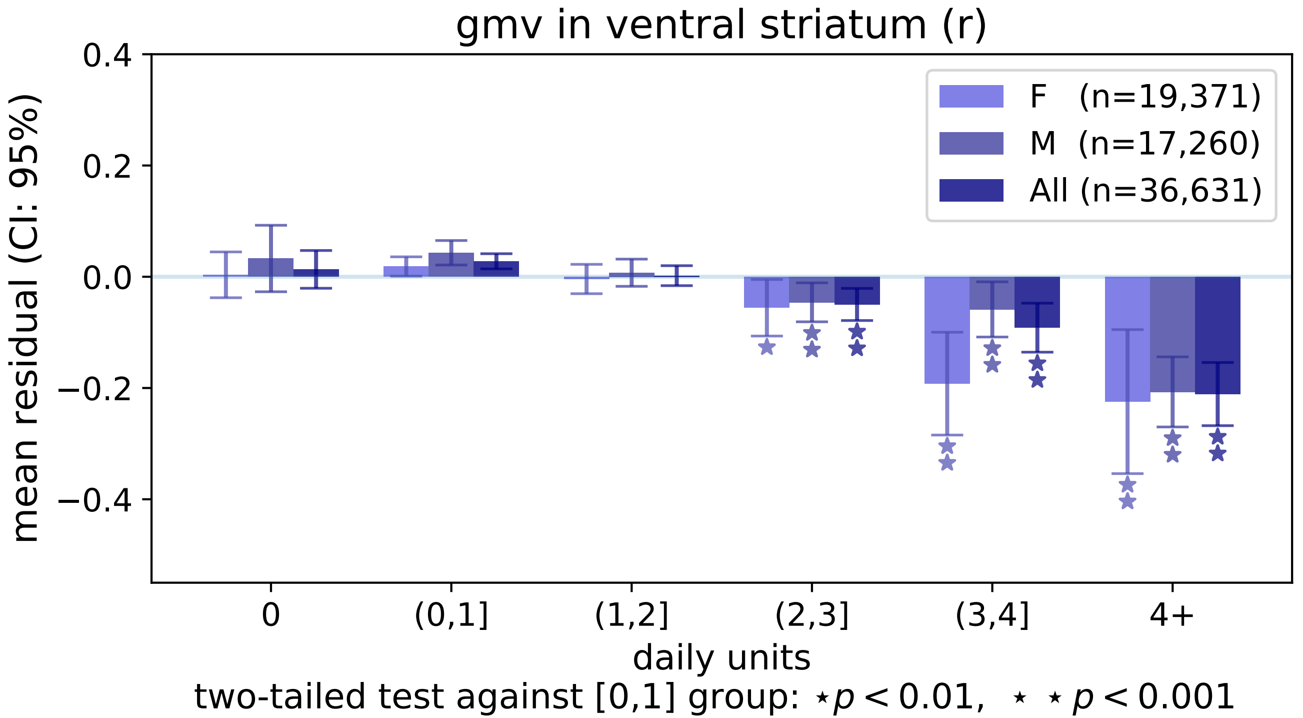
gmv in ventral striatum (r)
mean residual (CI: 95%)
0.4
0.2
0.0
−0.2
−0.4
F (n=19,371)
M (n=17,260)
All (n=36,631)
0
(0,1]
(1,2]
(2,3]
(3,4]
4+
daily units
two-tailed test against [0,1] group: ⋆ p < 0.01, ⋆ ⋆ p < 0.001

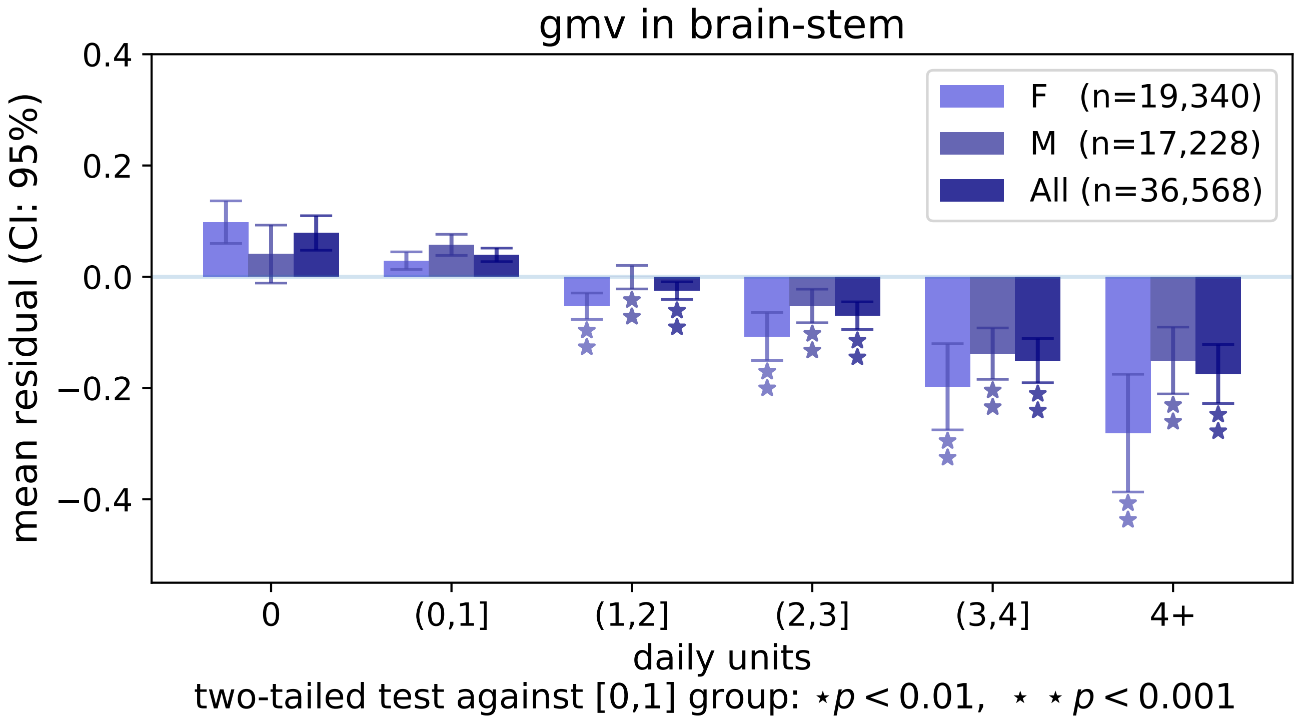
gmv in brain-stem
F (n=19,340)
M (n=17,228)
All (n=36,568)
mean residual (CI: 95%)
0.4
0.2
0.0
−0.2
−0.4
0
(0,1]
(1,2]
(2,3]
(3,4]
4+
daily units
two-tailed test against [0,1] group: ⋆p < 0.01, ⋆ ⋆ p < 0.001

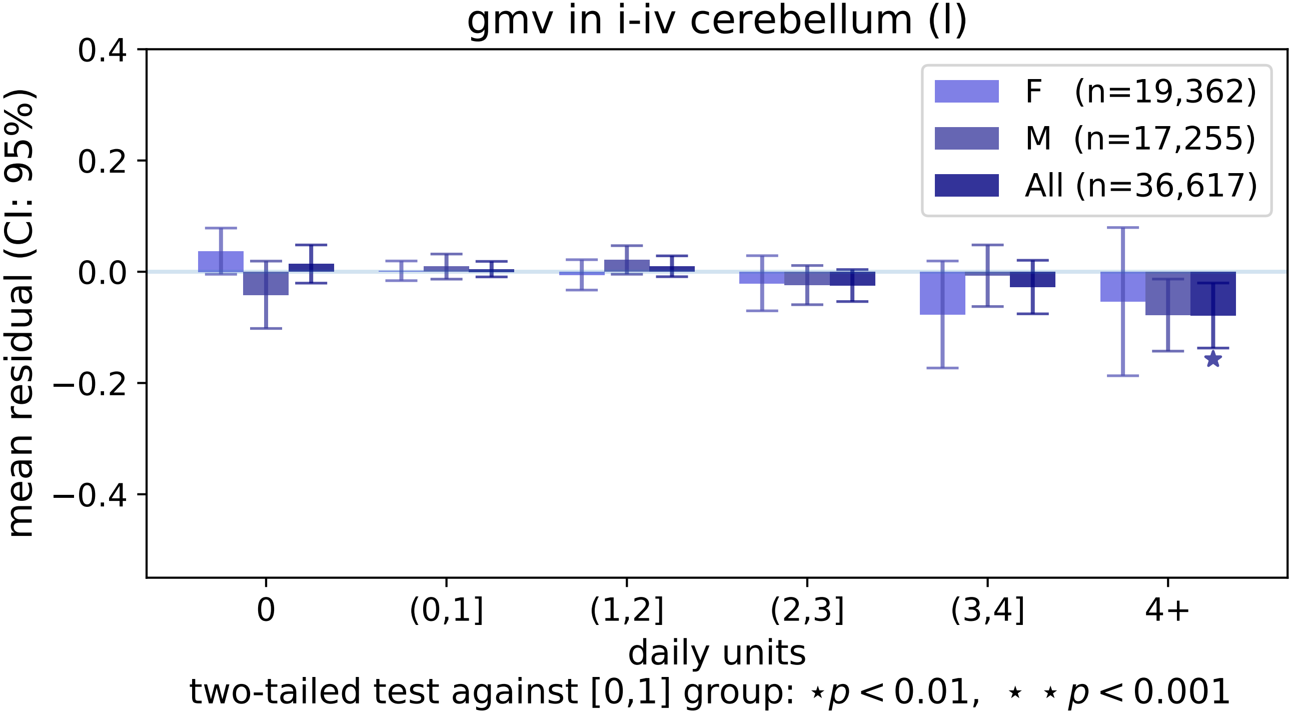

gmv in i-iv cerebellum (l)
mean residual (CI: 95%)
0.4
0.2
0.0
−0.2
−0.4
F (n=19,362)
M (n=17,255)
All (n=36,617)
0
(0,1]
(1,2]
(2,3]
(3,4]
4+
daily units
two-tailed test against [0,1] group: ⋆ p < 0.01, ⋆ ⋆ p < 0.001

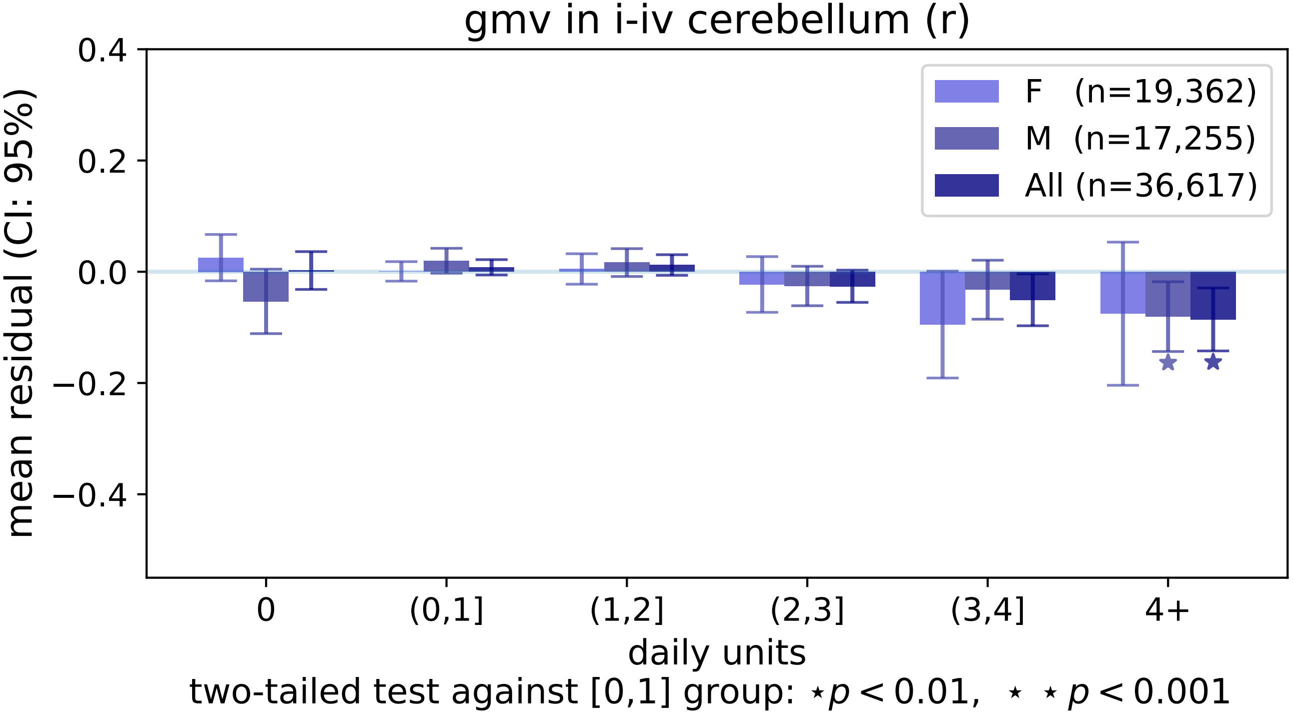
gmv in i-iv cerebellum (r)
mean residual (CI: 95%)
0.4
0.2
0.0
−0.2
−0.4
F (n=19,362)
M (n=17,255)
All (n=36,617)
0
(0,1]
(1,2]
(2,3]
(3,4]
4+
daily units
two-tailed test against [0,1] group: ⋆p < 0.01, ⋆ ⋆ p < 0.001

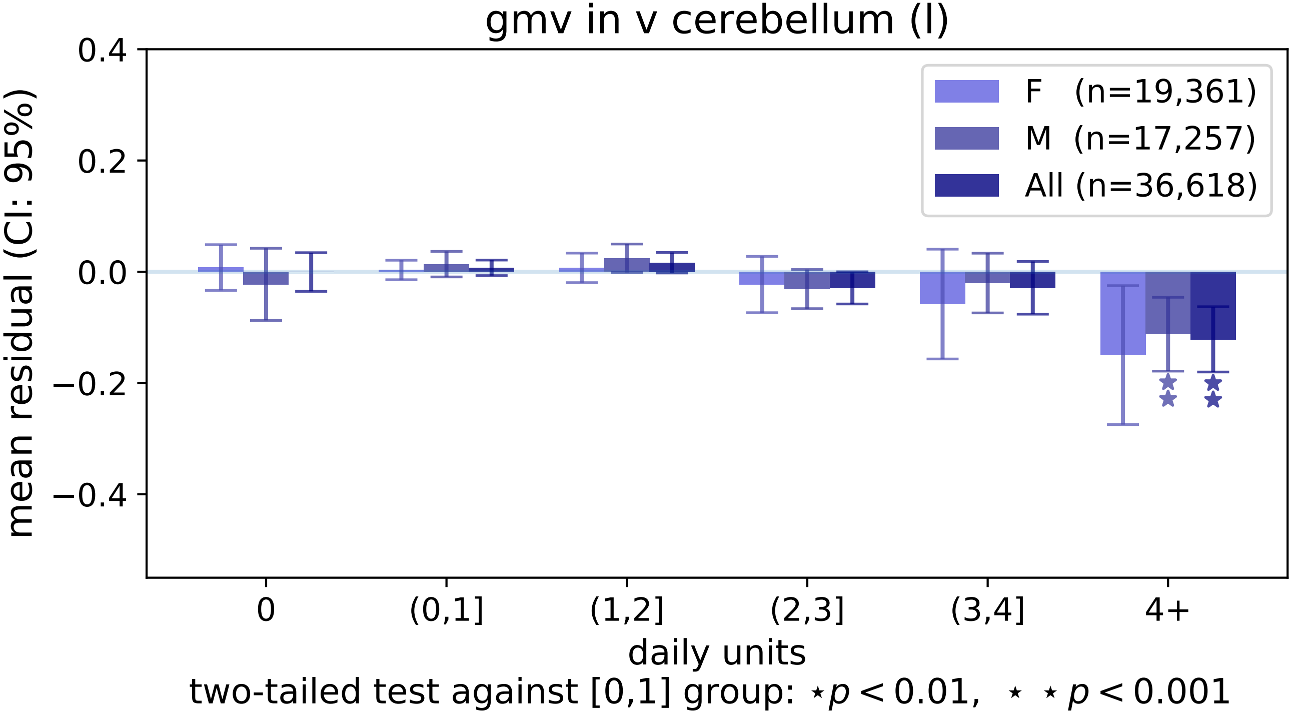

gmv in v cerebellum (l)
mean residual (CI: 95%)
0.4
0.2
0.0
−0.2
−0.4
F (n=19,361)
M (n=17,257)
All (n=36,618)
0
(0,1]
(1,2]
(2,3]
(3,4]
4+
daily units
two-tailed test against [0,1] group: ⋆p < 0.01, ⋆ ⋆ p < 0.001

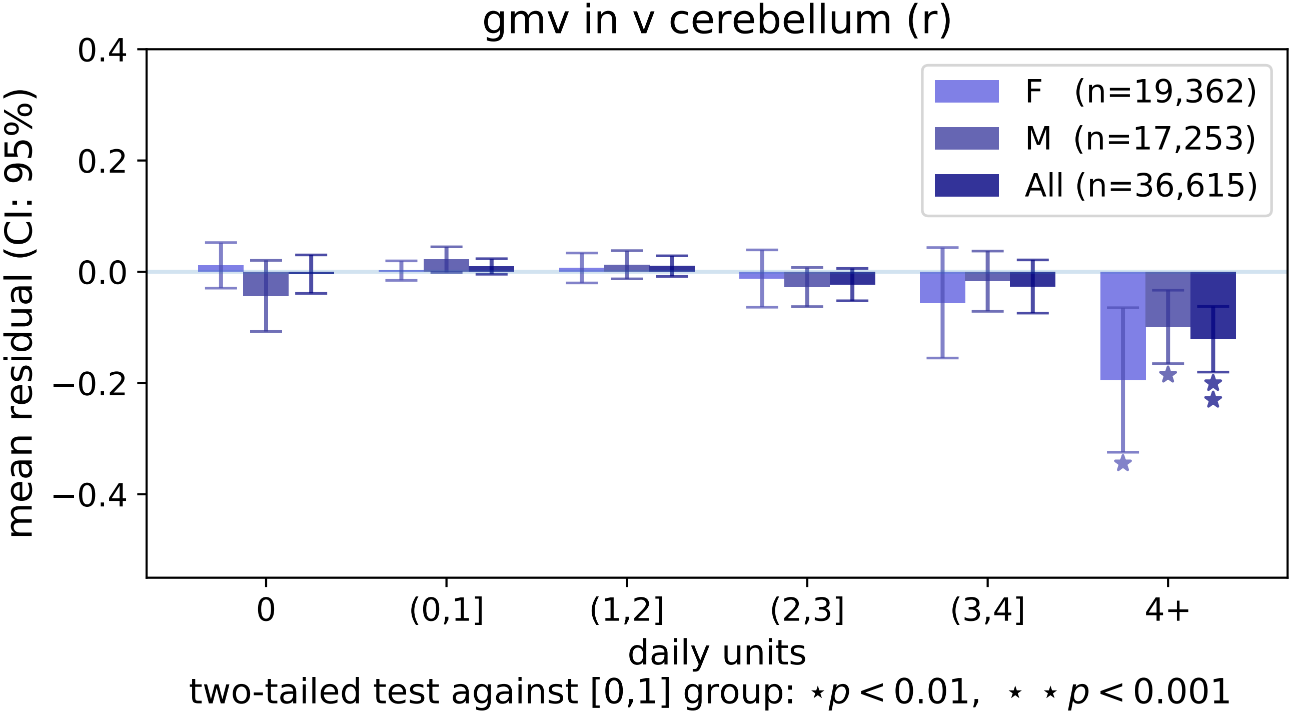

gmv in v cerebellum (r)
mean residual (CI: 95%)
0.4
0.2
0.0
−0.2
−0.4
F (n=19,362)
M (n=17,253)
All (n=36,615)
0
(0,1]
(1,2]
(2,3]
(3,4]
4+
daily units
two-tailed test against [0,1] group: ⋆ $p < 0.01$, ⋆ ⋆ $p < 0.001$

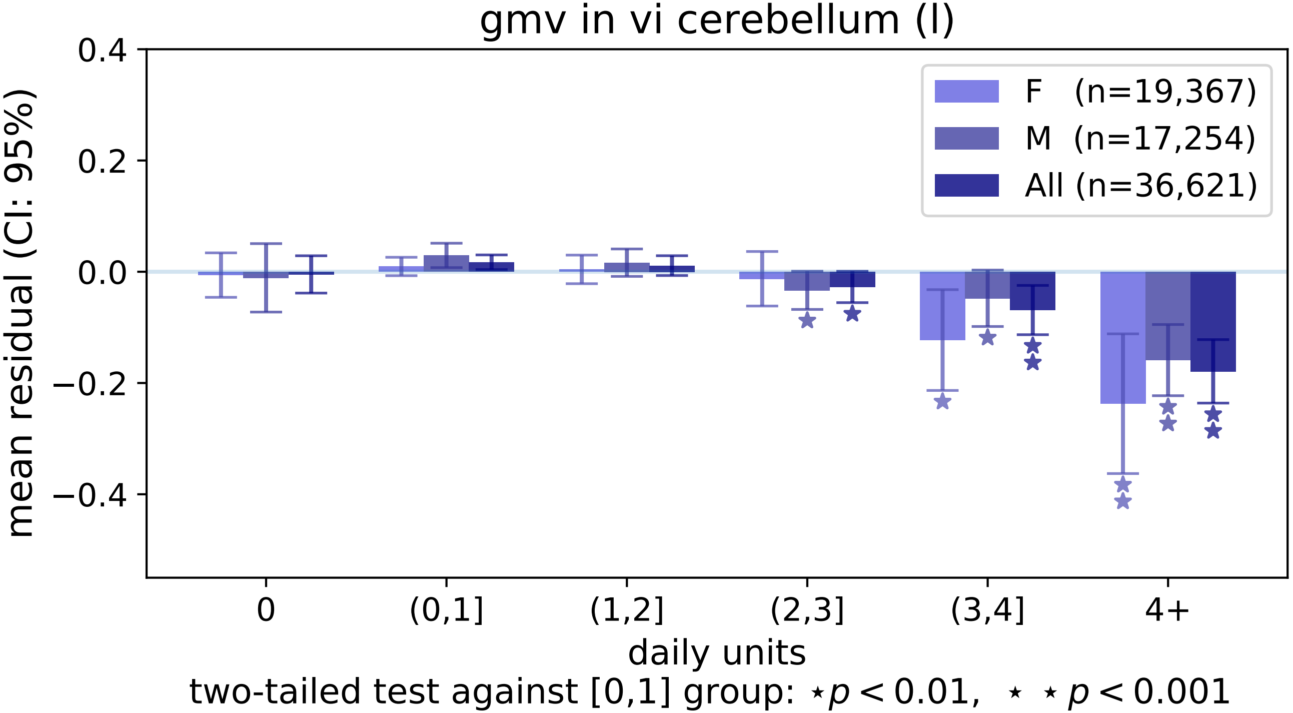
gmv in vi cerebellum (l)
mean residual (CI: 95%)
0.4
0.2
0.0
−0.2
−0.4
F (n=19,367)
M (n=17,254)
All (n=36,621)
0
(0,1]
(1,2]
(2,3]
(3,4]
4+
daily units
two-tailed test against [0,1] group: ⋆p < 0.01, ⋆ ⋆ p < 0.001

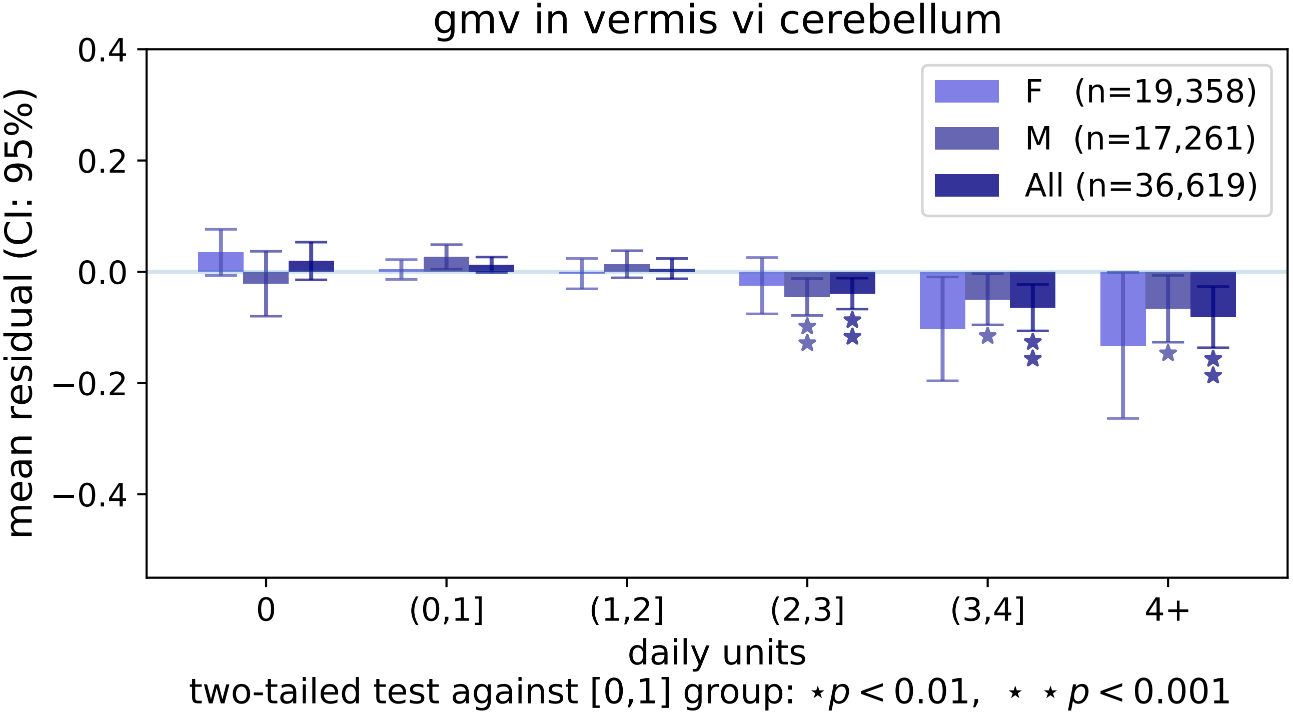

gmv in vermis vi cerebellum
mean residual (CI: 95%)
0.4
0.2
0.0
−0.2
−0.4
F (n=19,358)
M (n=17,261)
All (n=36,619)
0
(0,1]
(1,2]
(2,3]
(3,4]
4+
daily units
two-tailed test against [0,1] group: ⋆ $p < 0.01$, ⋆ ⋆ $p < 0.001$

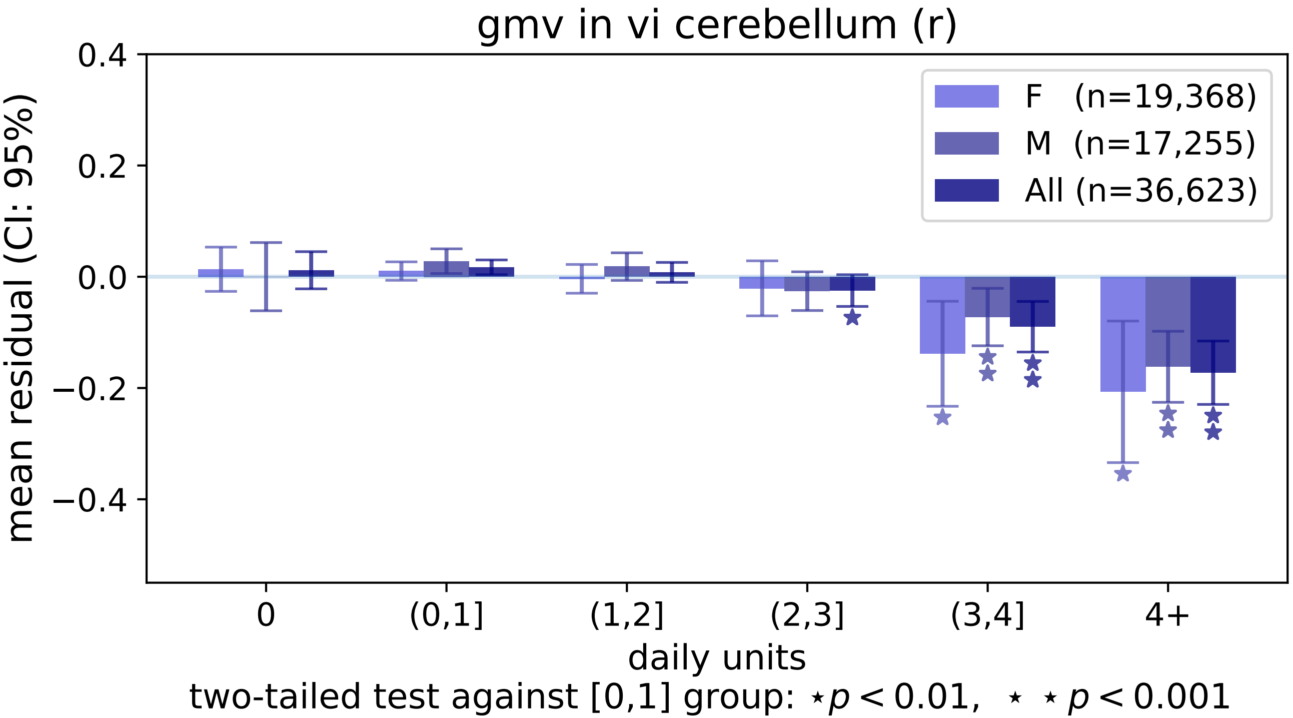
gmv in vi cerebellum (r)
mean residual (CI: 95%)
0.4
0.2
0.0
−0.2
−0.4
F (n=19,368)
M (n=17,255)
All (n=36,623)
0
(0,1]
(1,2]
(2,3]
(3,4]
4+
daily units
two-tailed test against [0,1] group: ⋆p < 0.01, ⋆ ⋆ p < 0.001

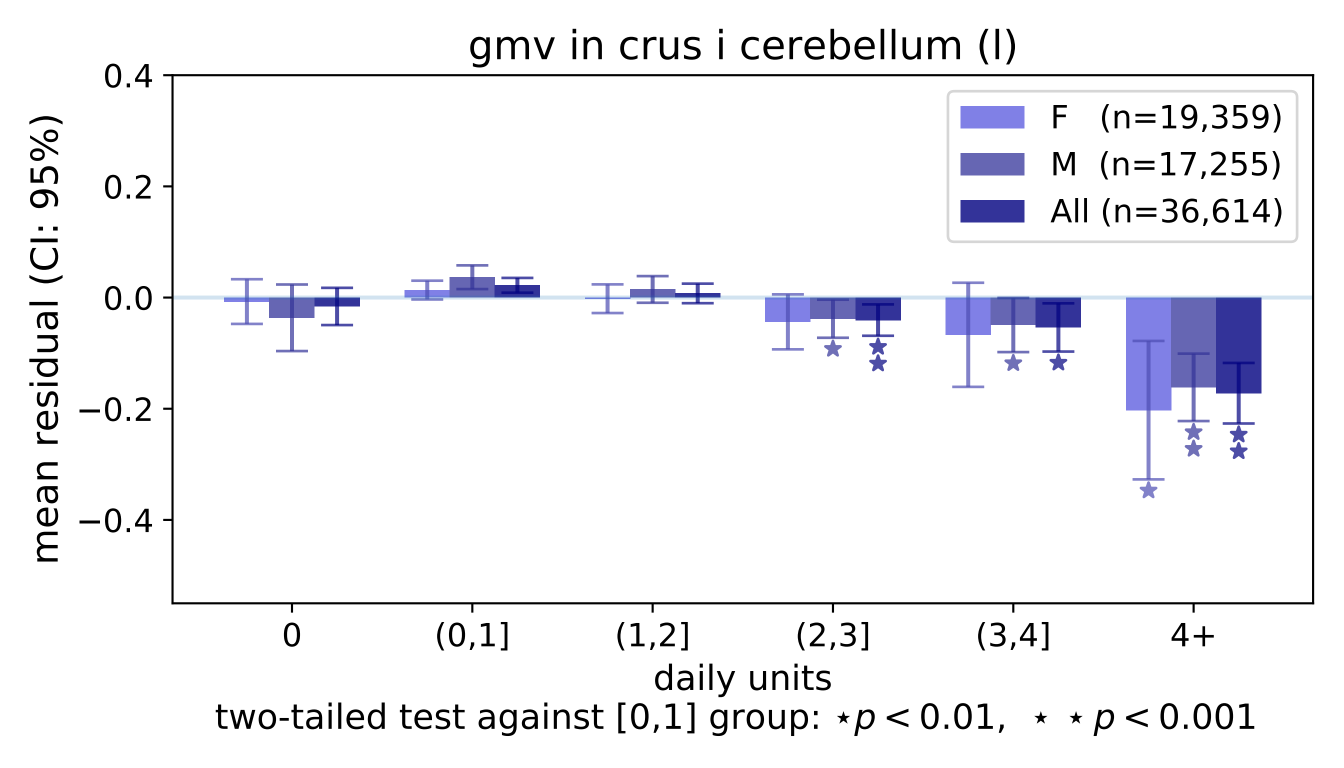
gmv in crus i cerebellum (l)
mean residual (CI: 95%)
0.4
0.2
0.0
−0.2
−0.4
F (n=19,359)
M (n=17,255)
All (n=36,614)
0
(0,1]
(1,2]
(2,3]
(3,4]
4+
daily units
two-tailed test against [0,1] group: ⋆p < 0.01, ⋆ ⋆ p < 0.001

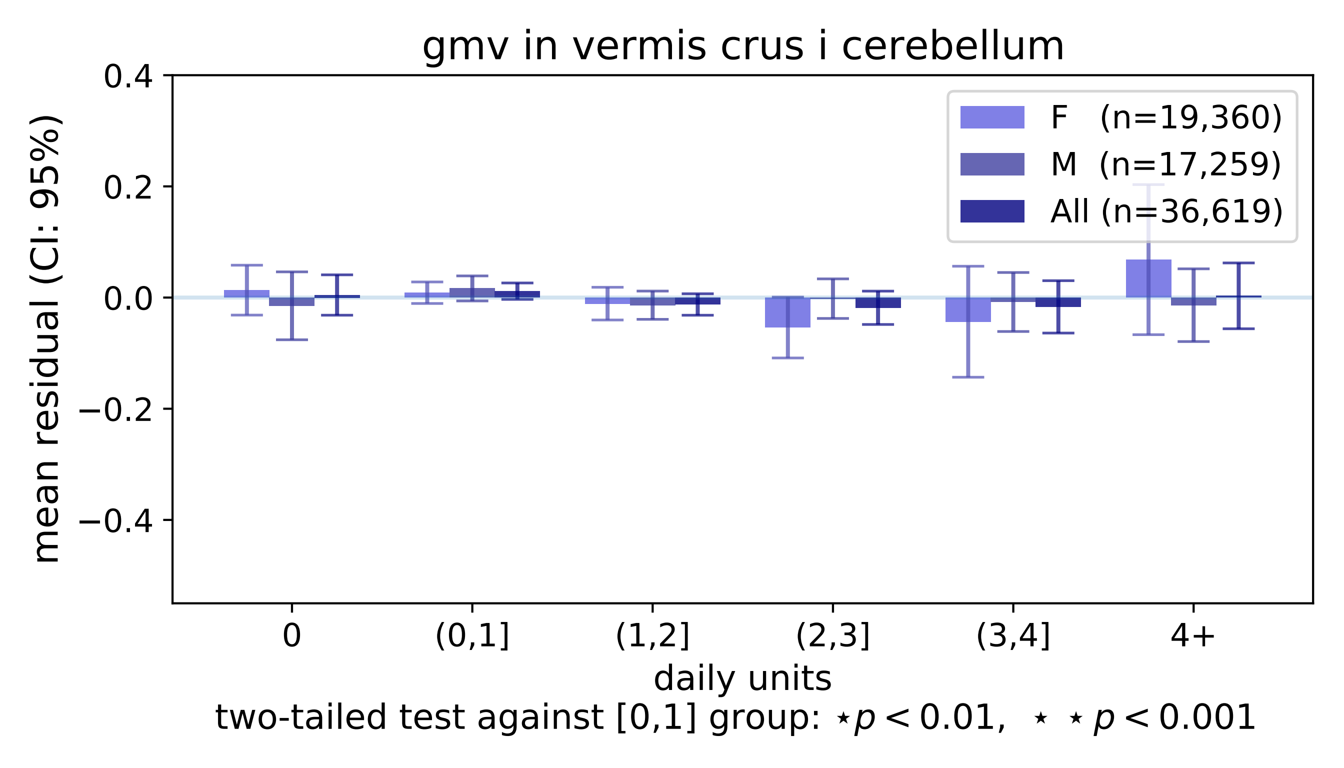
gmv in vermis crus i cerebellum
mean residual (CI: 95%)
0.4
0.2
0.0
−0.2
−0.4
F (n=19,360)
M (n=17,259)
All (n=36,619)
0
(0,1]
(1,2]
(2,3]
(3,4]
4+
daily units
two-tailed test against [0,1] group: ⋆p < 0.01, ⋆ ⋆ p < 0.001

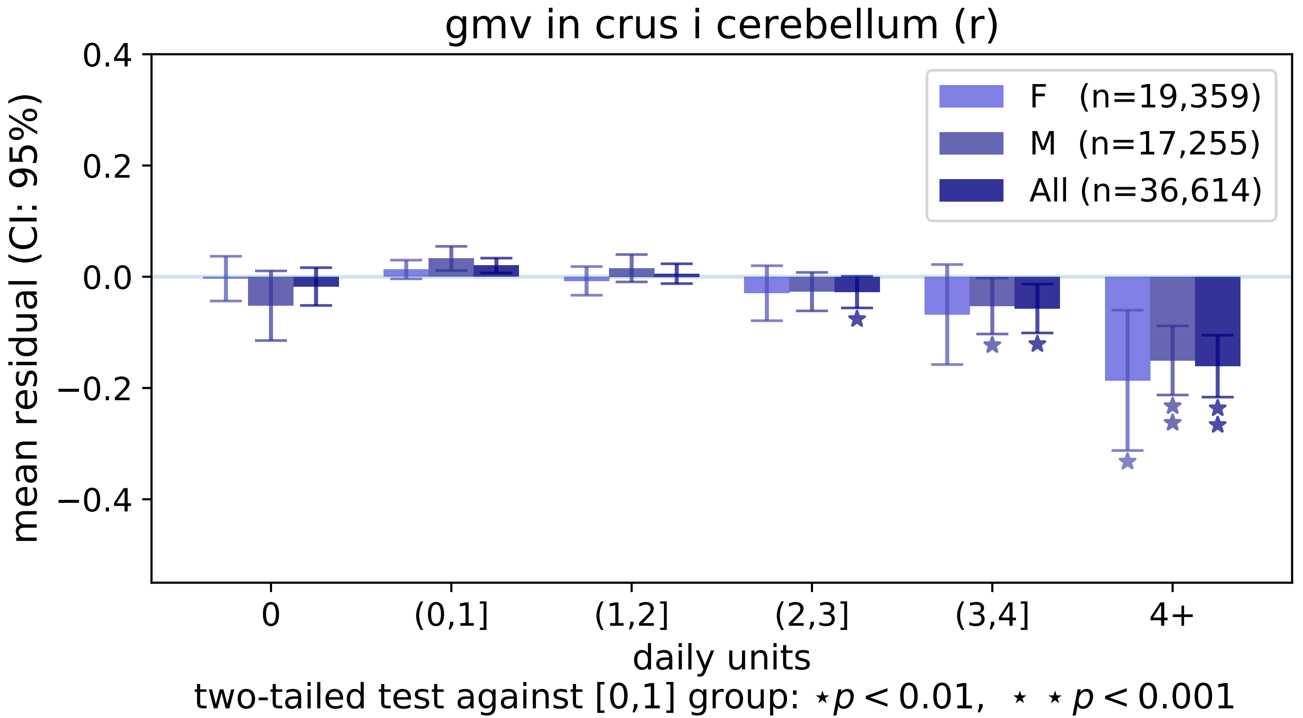

gmv in crus i cerebellum (r)
mean residual (CI: 95%)
0.4
0.2
0.0
−0.2
−0.4
F (n=19,359)
M (n=17,255)
All (n=36,614)
0
(0,1]
(1,2]
(2,3]
(3,4]
4+
daily units
two-tailed test against [0,1] group: ⋆ p < 0.01, ⋆ ⋆ p < 0.001

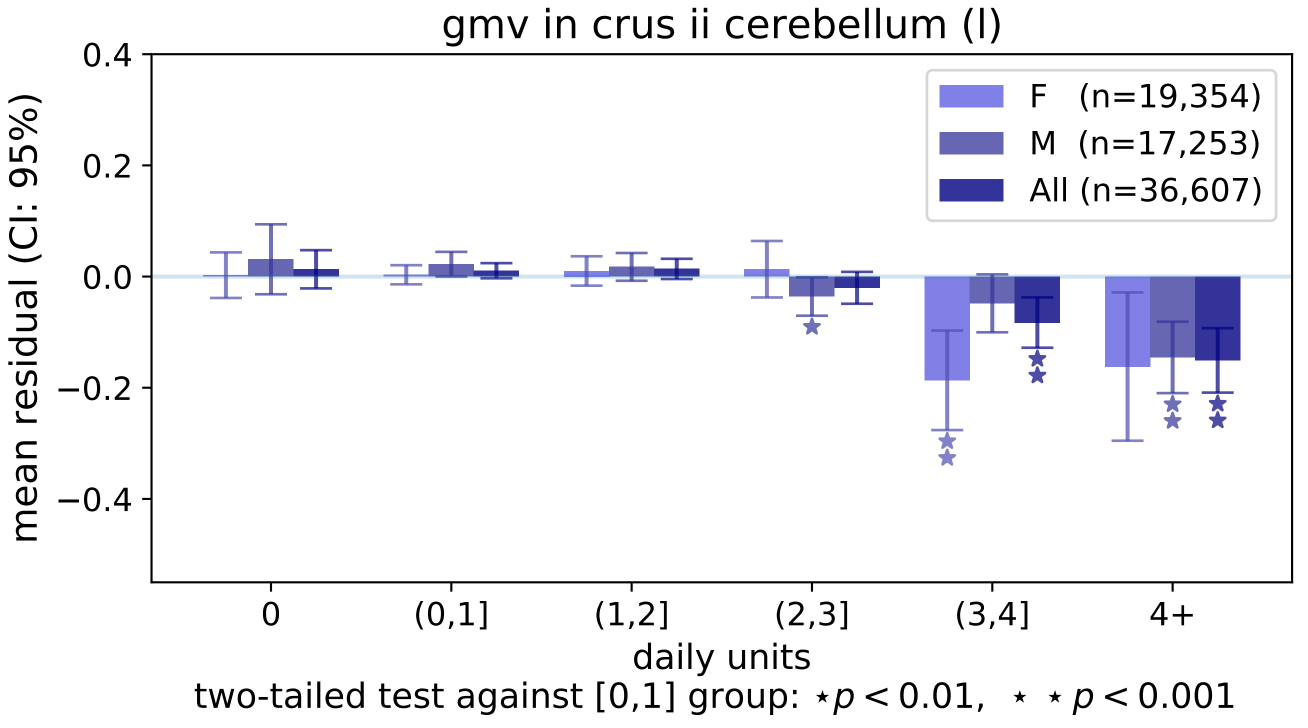

gmv in crus ii cerebellum (l)
mean residual (CI: 95%)
0.4
0.2
0.0
−0.2
−0.4
F (n=19,354)
M (n=17,253)
All (n=36,607)
0
(0,1]
(1,2]
(2,3]
(3,4]
4+
daily units
two-tailed test against [0,1] group: ⋆p < 0.01, ⋆ ⋆ p < 0.001

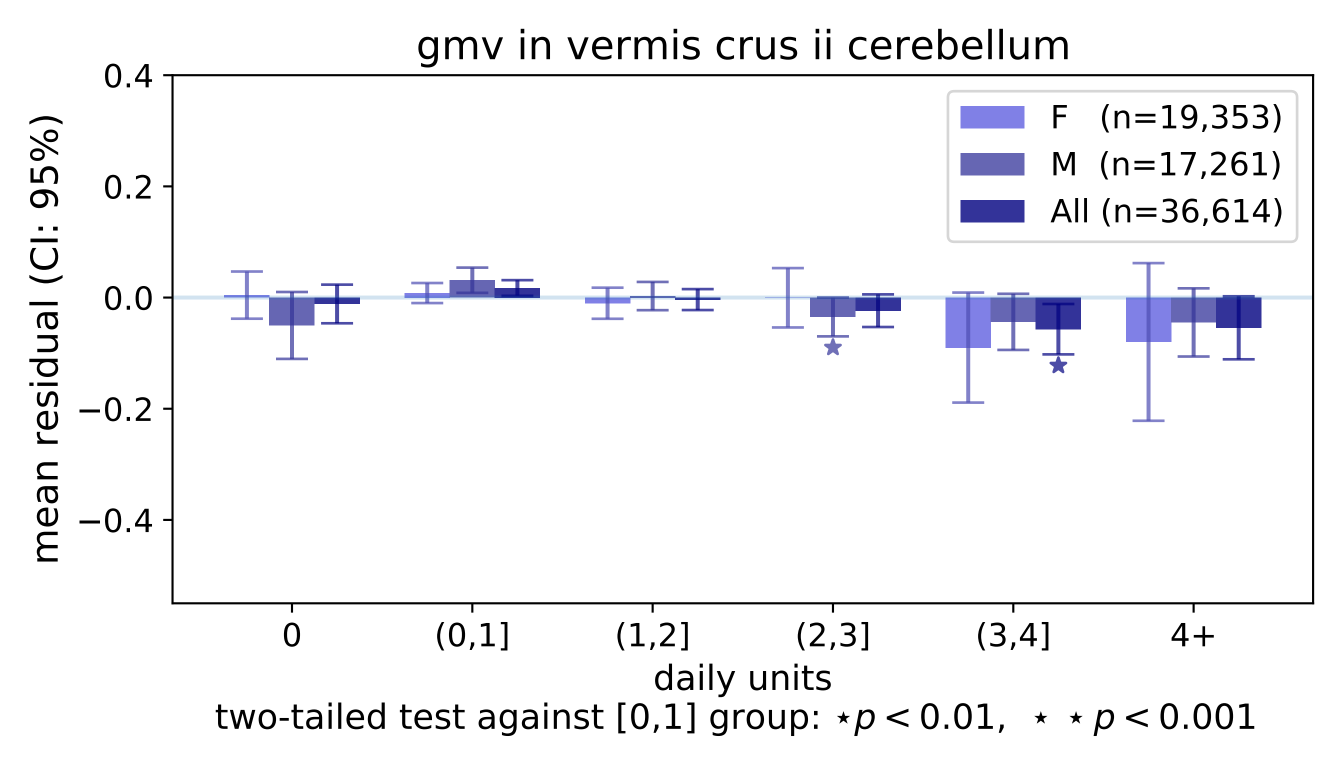
gmv in vermis crus ii cerebellum
mean residual (CI: 95%)
0.4
0.2
0.0
−0.2
−0.4
F (n=19,353)
M (n=17,261)
All (n=36,614)
0
(0,1]
(1,2]
(2,3]
(3,4]
4+
daily units
two-tailed test against [0,1] group: ⋆ $p < 0.01$, ⋆ ⋆ $p < 0.001$

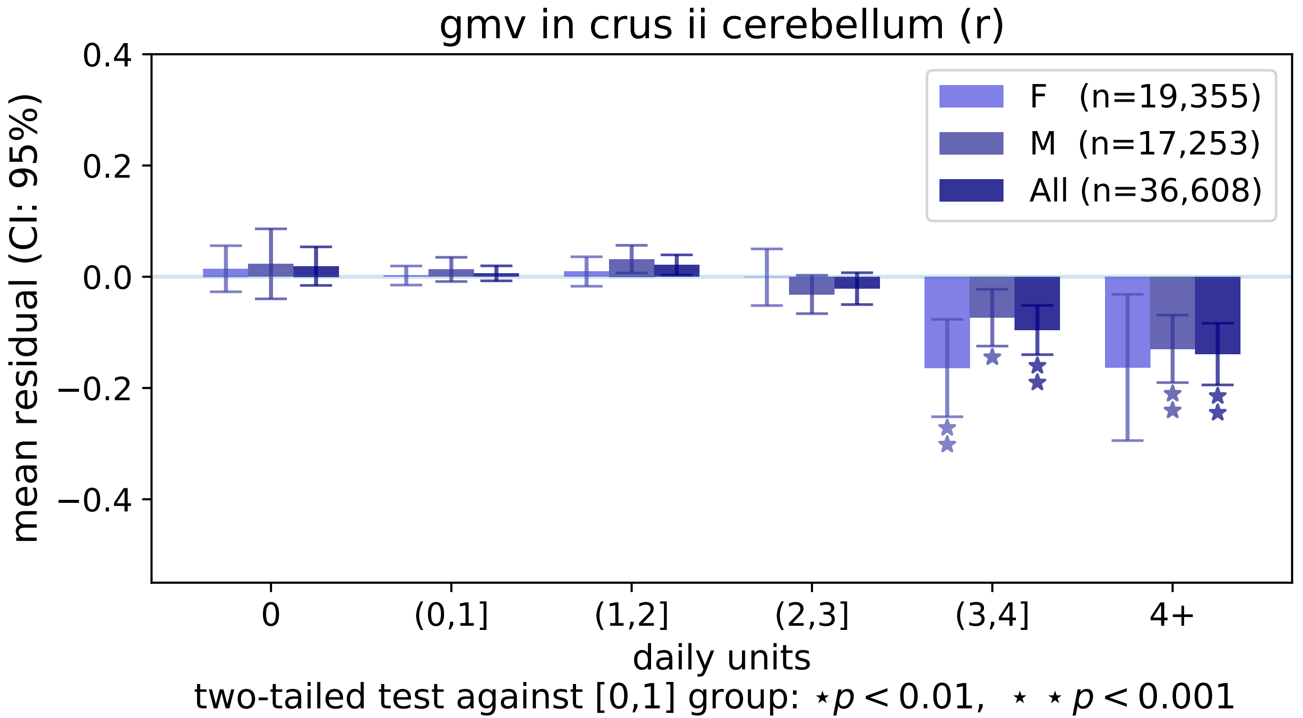
gmv in crus ii cerebellum (r)
mean residual (CI: 95%)
0.4
0.2
0.0
−0.2
−0.4
F (n=19,355)
M (n=17,253)
All (n=36,608)
0
(0,1]
(1,2]
(2,3]
(3,4]
4+
daily units
two-tailed test against [0,1] group: ⋆ $p < 0.01$, ⋆ ⋆ $p < 0.001$

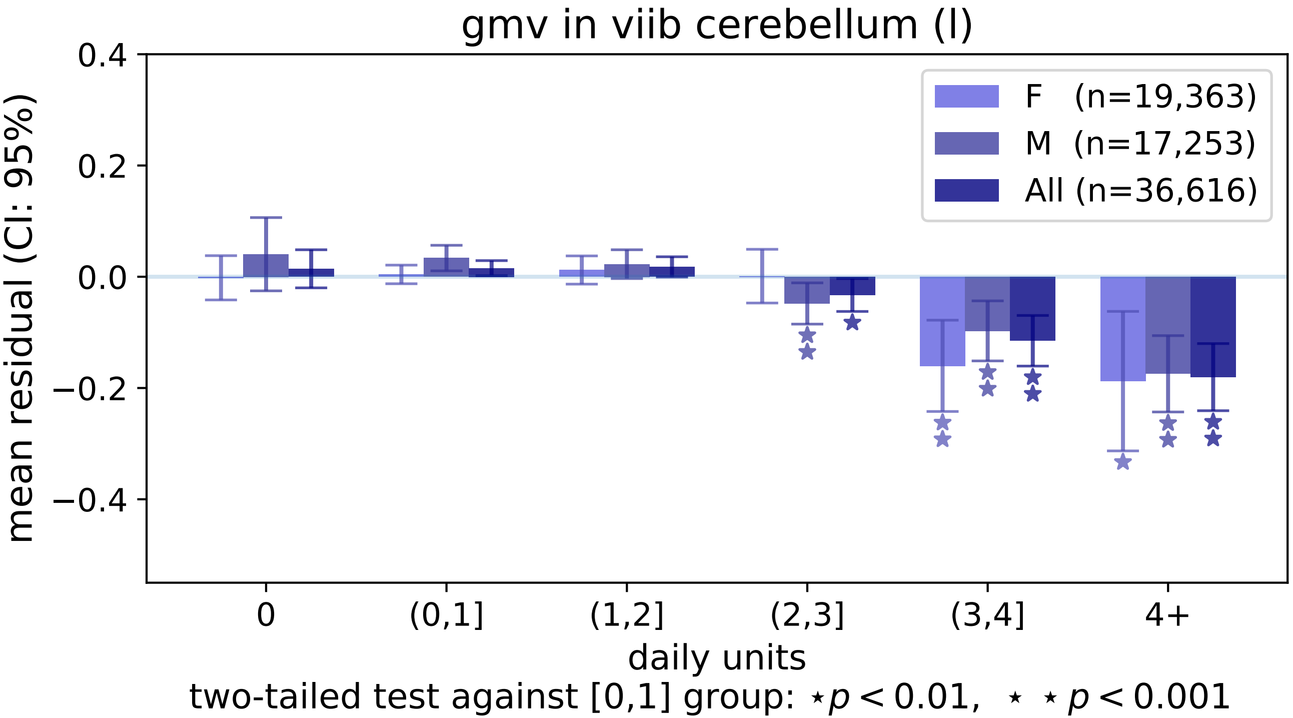
gmv in viib cerebellum (l)
0.4
0.2
0.0
−0.2
−0.4
mean residual (CI: 95%)
F (n=19,363)
M (n=17,253)
All (n=36,616)
0
(0,1]
(1,2]
(2,3]
(3,4]
4+
daily units
two-tailed test against [0,1] group: ⋆p < 0.01, ⋆ ⋆ p < 0.001

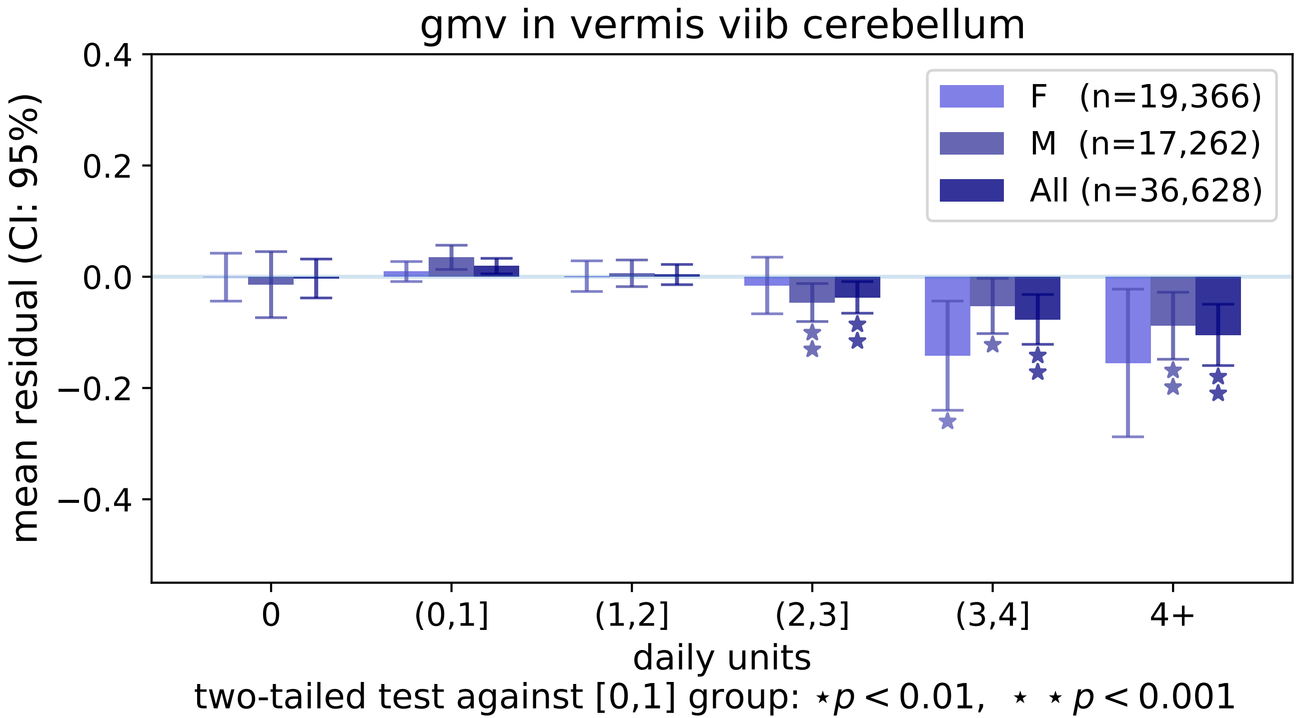
gmv in vermis viib cerebellum
mean residual (CI: 95%)
0.4
0.2
0.0
−0.2
−0.4
F (n=19,366)
M (n=17,262)
All (n=36,628)
0
(0,1]
(1,2]
(2,3]
(3,4]
4+
daily units
two-tailed test against [0,1] group: ⋆p < 0.01, ⋆ ⋆ p < 0.001

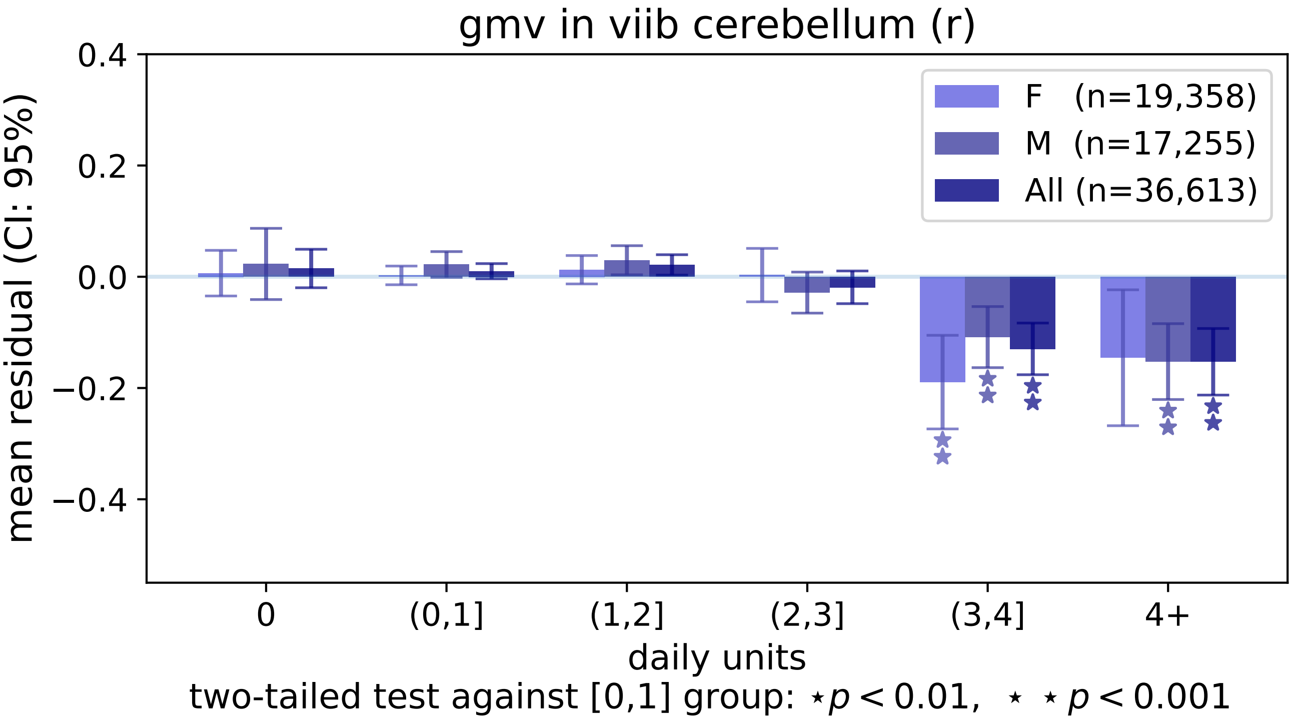
gmv in viib cerebellum (r)
mean residual (CI: 95%)
0.4
0.2
0.0
−0.2
−0.4
F (n=19,358)
M (n=17,255)
All (n=36,613)
0
(0,1]
(1,2]
(2,3]
(3,4]
4+
daily units
two-tailed test against [0,1] group: ⋆ $p < 0.01$, ⋆ ⋆ $p < 0.001$

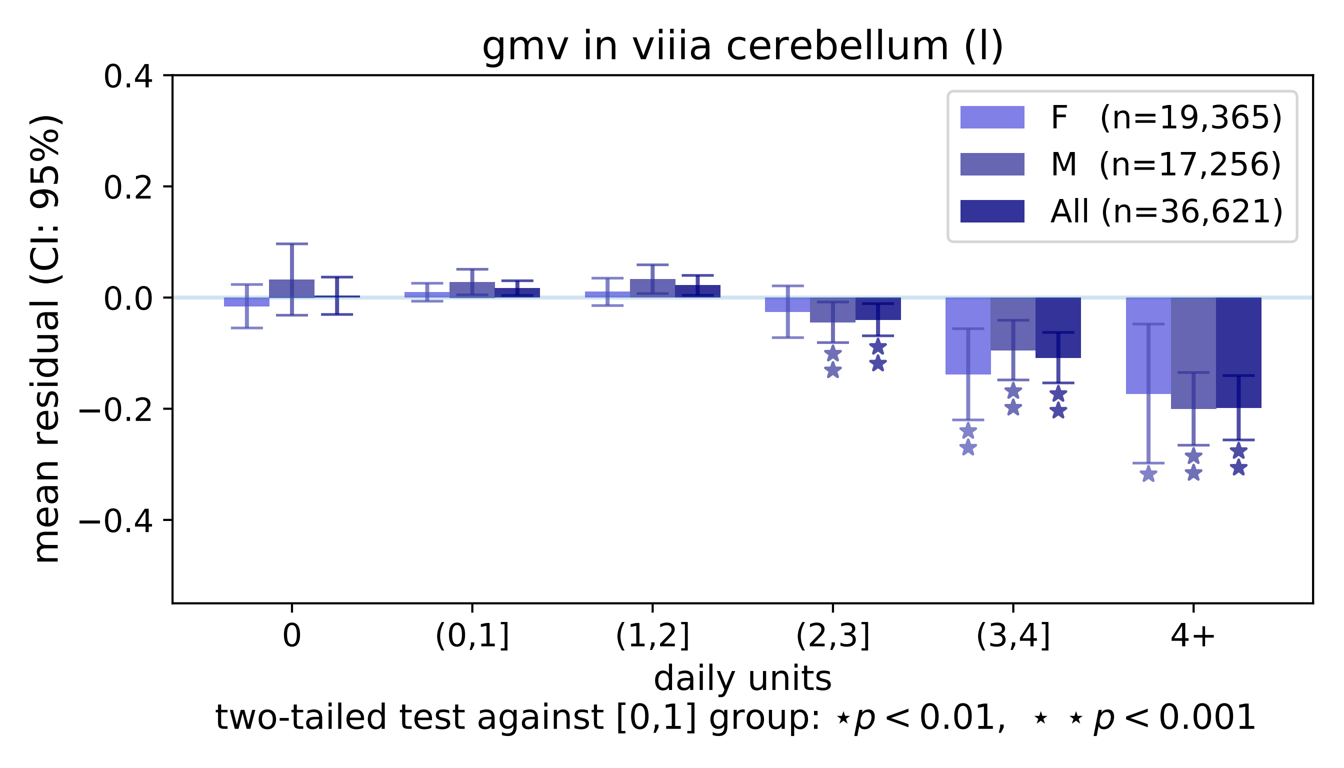

gmv in viiia cerebellum (l)
mean residual (CI: 95%)
0.4
0.2
0.0
−0.2
−0.4
F (n=19,365)
M (n=17,256)
All (n=36,621)
0
(0,1]
(1,2]
(2,3]
(3,4]
4+
daily units
two-tailed test against [0,1] group: ⋆ p < 0.01, ⋆ ⋆ p < 0.001

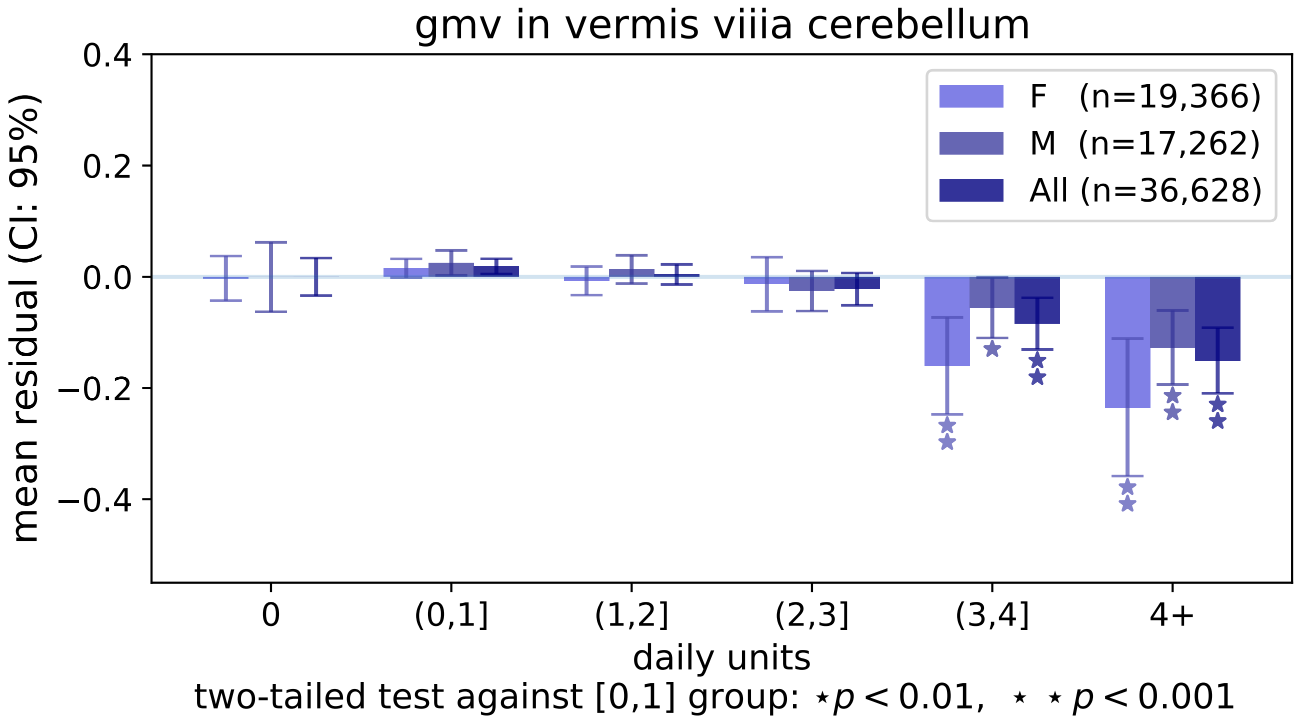
gmv in vermis viiia cerebellum
mean residual (CI: 95%)
0.4
0.2
0.0
−0.2
−0.4
F (n=19,366)
M (n=17,262)
All (n=36,628)
0
(0,1]
(1,2]
(2,3]
(3,4]
4+
daily units
two-tailed test against [0,1] group: ⋆ p < 0.01, ⋆ ⋆ p < 0.001

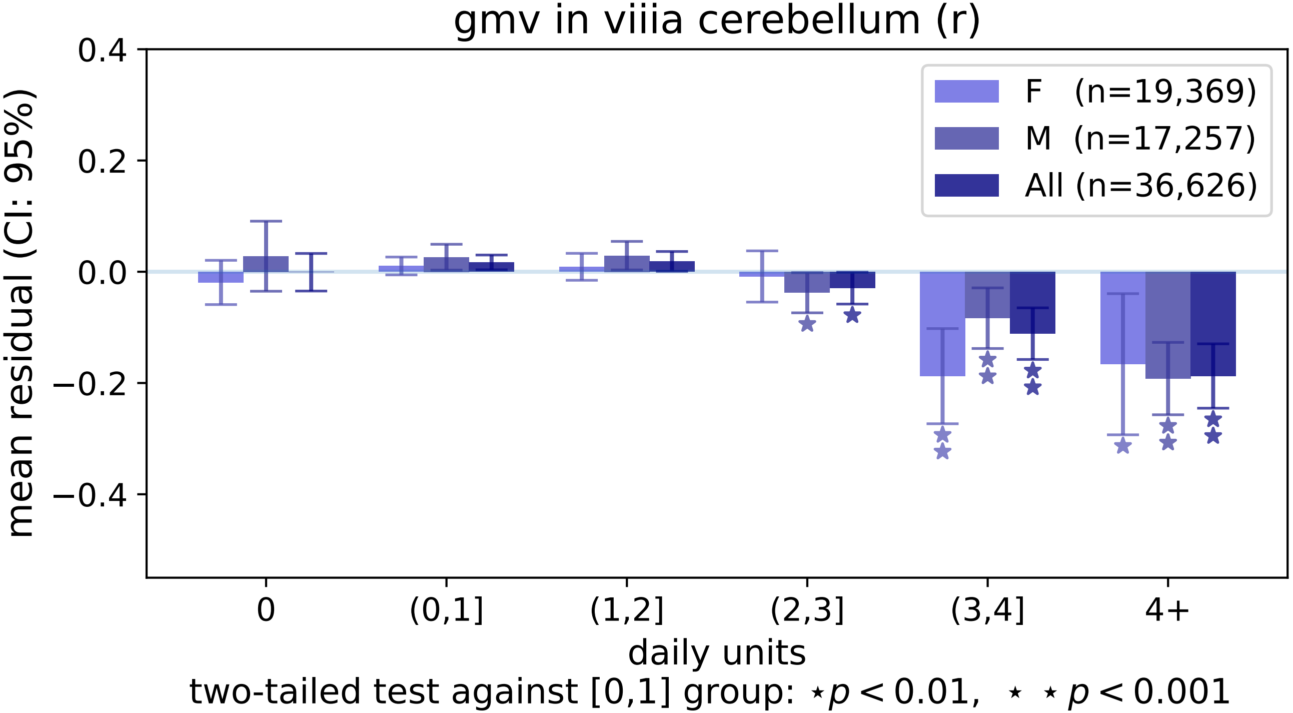

gmv in viiia cerebellum (r)
mean residual (CI: 95%)
0.4
0.2
0.0
−0.2
−0.4
F (n=19,369)
M (n=17,257)
All (n=36,626)
0
(0,1]
(1,2]
(2,3]
(3,4]
4+
daily units
two-tailed test against [0,1] group: ⋆p < 0.01, ⋆ ⋆ p < 0.001

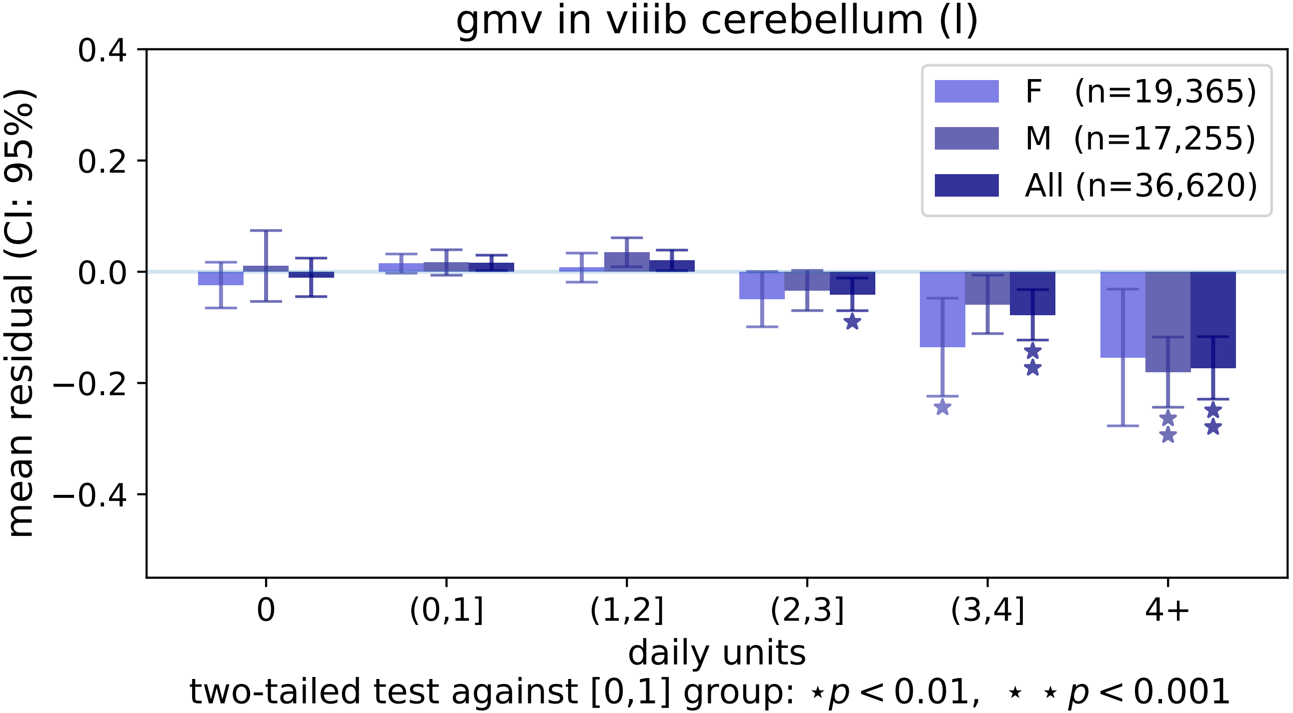

gmv in viiib cerebellum (l)
mean residual (CI: 95%)
0.4
0.2
0.0
−0.2
−0.4
F (n=19,365)
M (n=17,255)
All (n=36,620)
0
(0,1]
(1,2]
(2,3]
(3,4]
4+
daily units
two-tailed test against [0,1] group: ⋆p < 0.01, ⋆ ⋆ p < 0.001

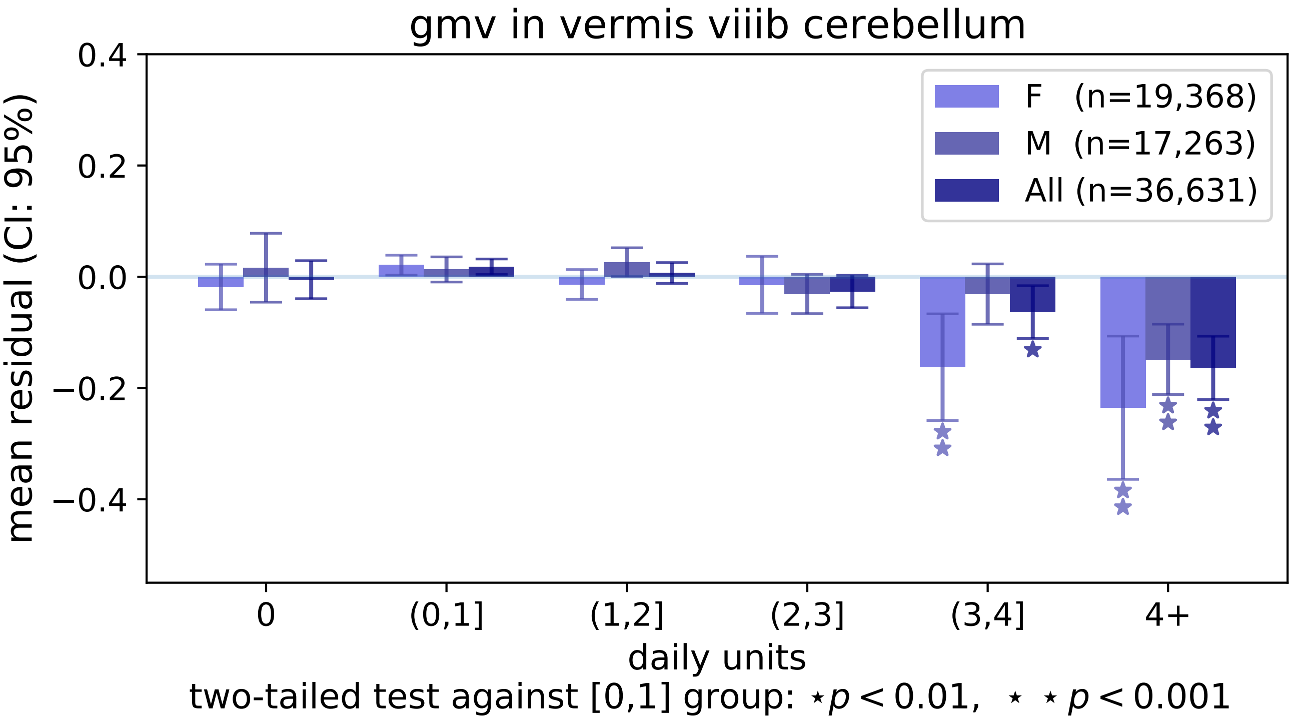

gmv in vermis viiib cerebellum
mean residual (CI: 95%)
0.4
0.2
0.0
−0.2
−0.4
F (n=19,368)
M (n=17,263)
All (n=36,631)
0
(0,1]
(1,2]
(2,3]
(3,4]
4+
daily units
two-tailed test against [0,1] group: ⋆p < 0.01, ⋆ ⋆ p < 0.001

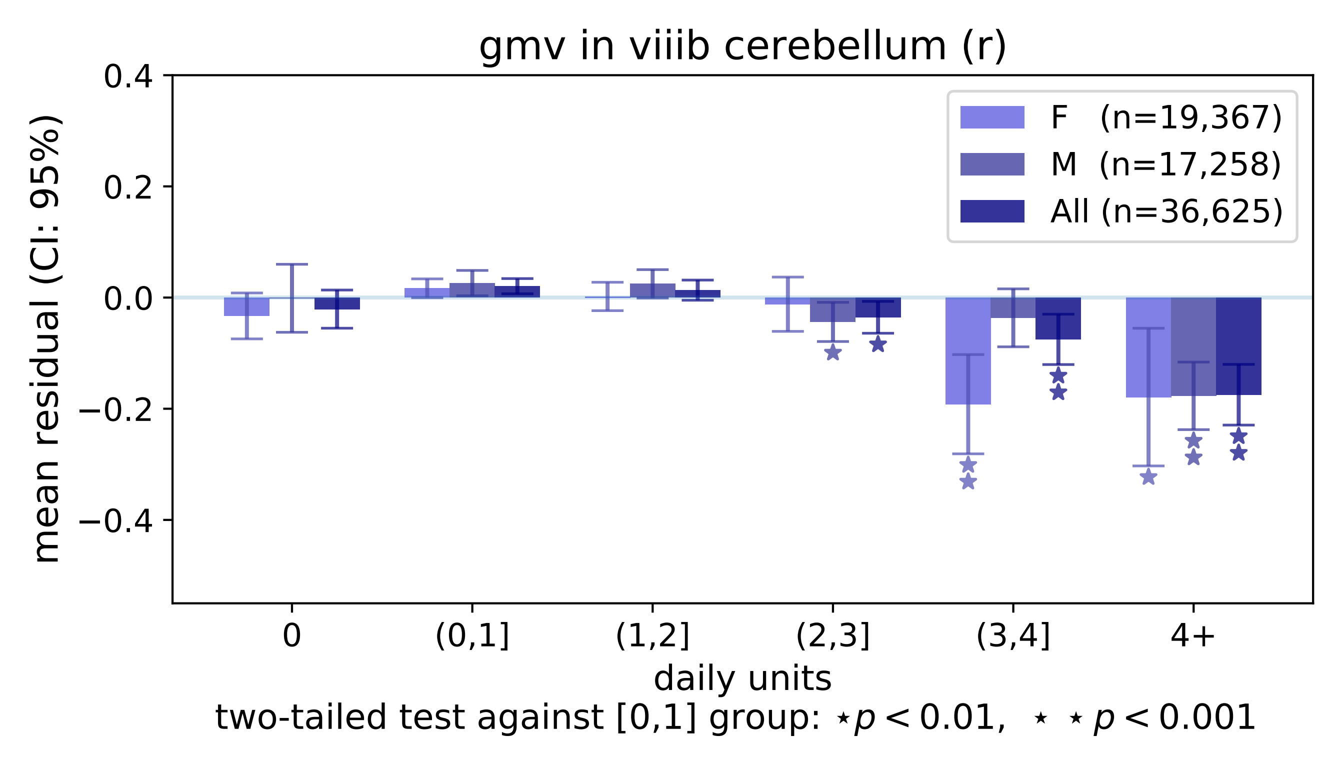
gmv in viiib cerebellum (r)
0.4
0.2
0.0
−0.2
−0.4
mean residual (CI: 95%)
F (n=19,367)
M (n=17,258)
All (n=36,625)
0
(0,1]
(1,2]
(2,3]
(3,4]
4+
daily units
two-tailed test against [0,1] group: ⋆p < 0.01, ⋆ ⋆ p < 0.001

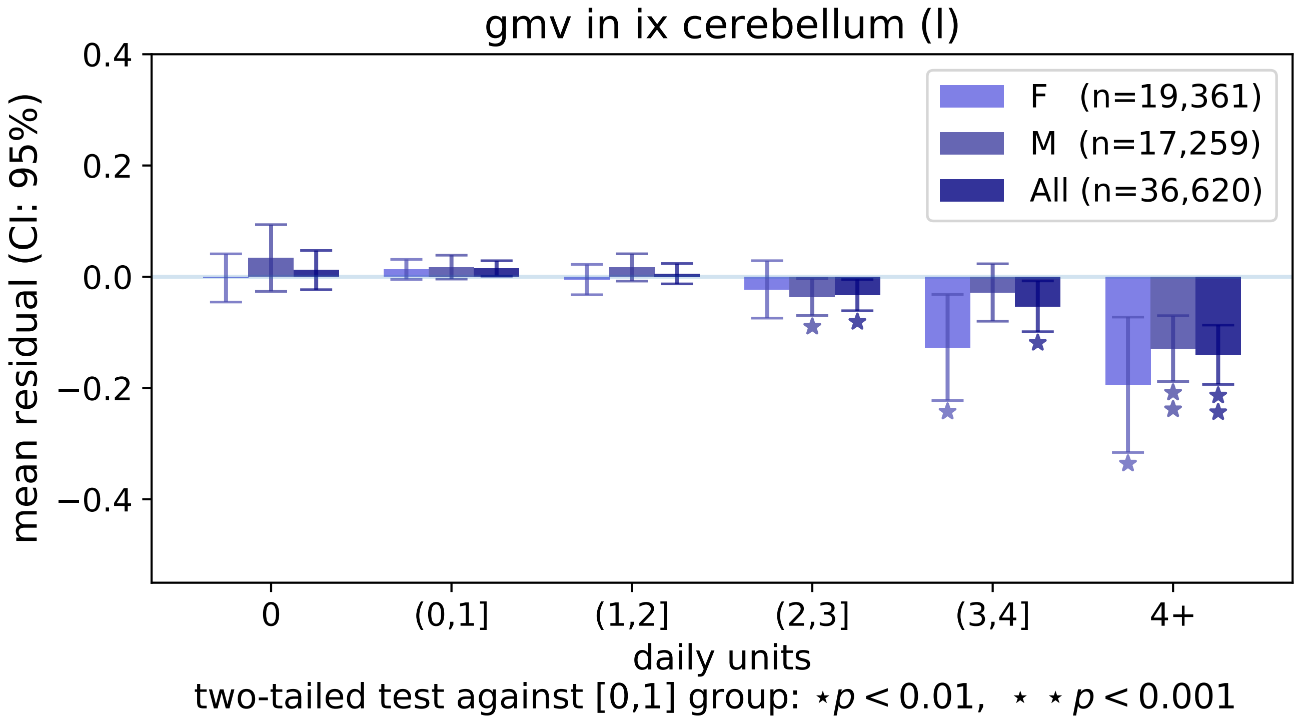

gmv in ix cerebellum (l)
0.4
0.2
0.0
−0.2
−0.4
mean residual (CI: 95%)
F (n=19,361)
M (n=17,259)
All (n=36,620)
0
(0,1]
(1,2]
(2,3]
(3,4]
4+
daily units
two-tailed test against [0,1] group: ⋆p < 0.01, ⋆ ⋆ p < 0.001

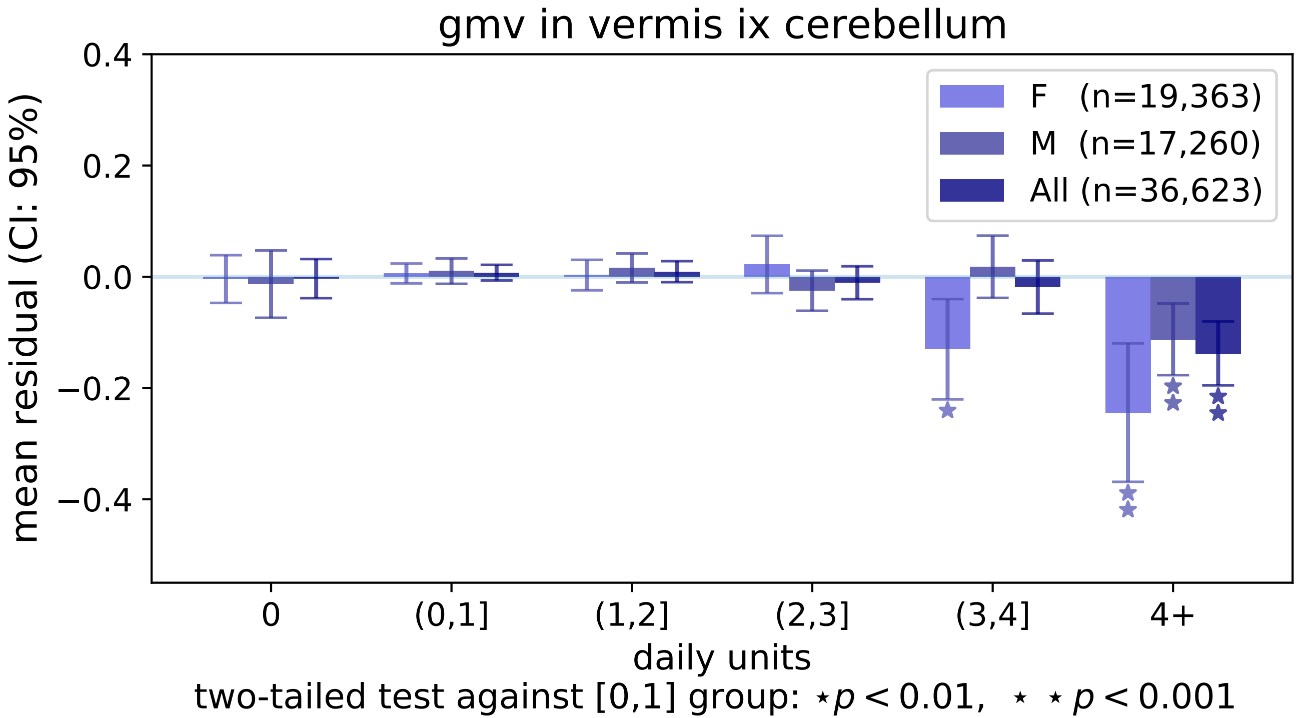
gmv in vermis ix cerebellum
mean residual (CI: 95%)
0.4
0.2
0.0
−0.2
−0.4
F (n=19,363)
M (n=17,260)
All (n=36,623)
0
(0,1]
(1,2]
(2,3]
(3,4]
4+
daily units
two-tailed test against [0,1] group: ⋆ $p < 0.01$, ⋆ ⋆ $p < 0.001$

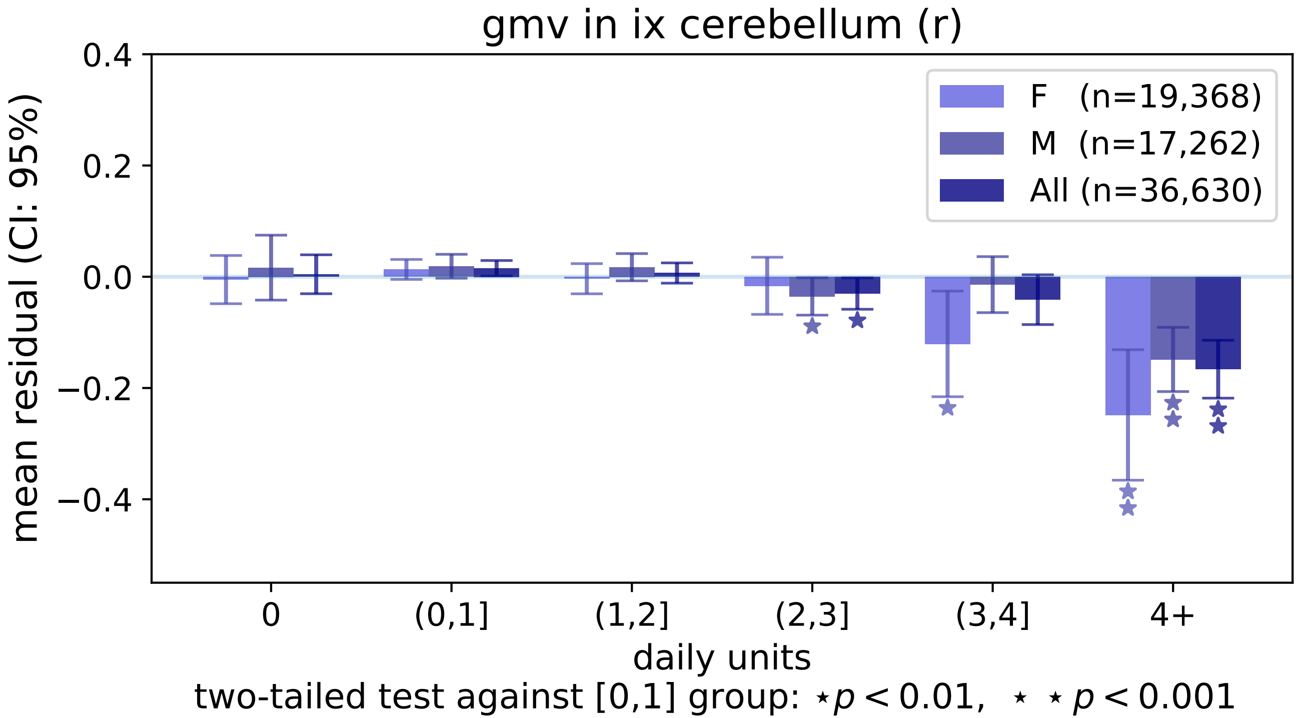
gmv in ix cerebellum (r)
mean residual (CI: 95%)
0.4
0.2
0.0
−0.2
−0.4
F (n=19,368)
M (n=17,262)
All (n=36,630)
0
(0,1]
(1,2]
(2,3]
(3,4]
4+
daily units
two-tailed test against [0,1] group: ⋆ $p < 0.01$, ⋆ ⋆ $p < 0.001$

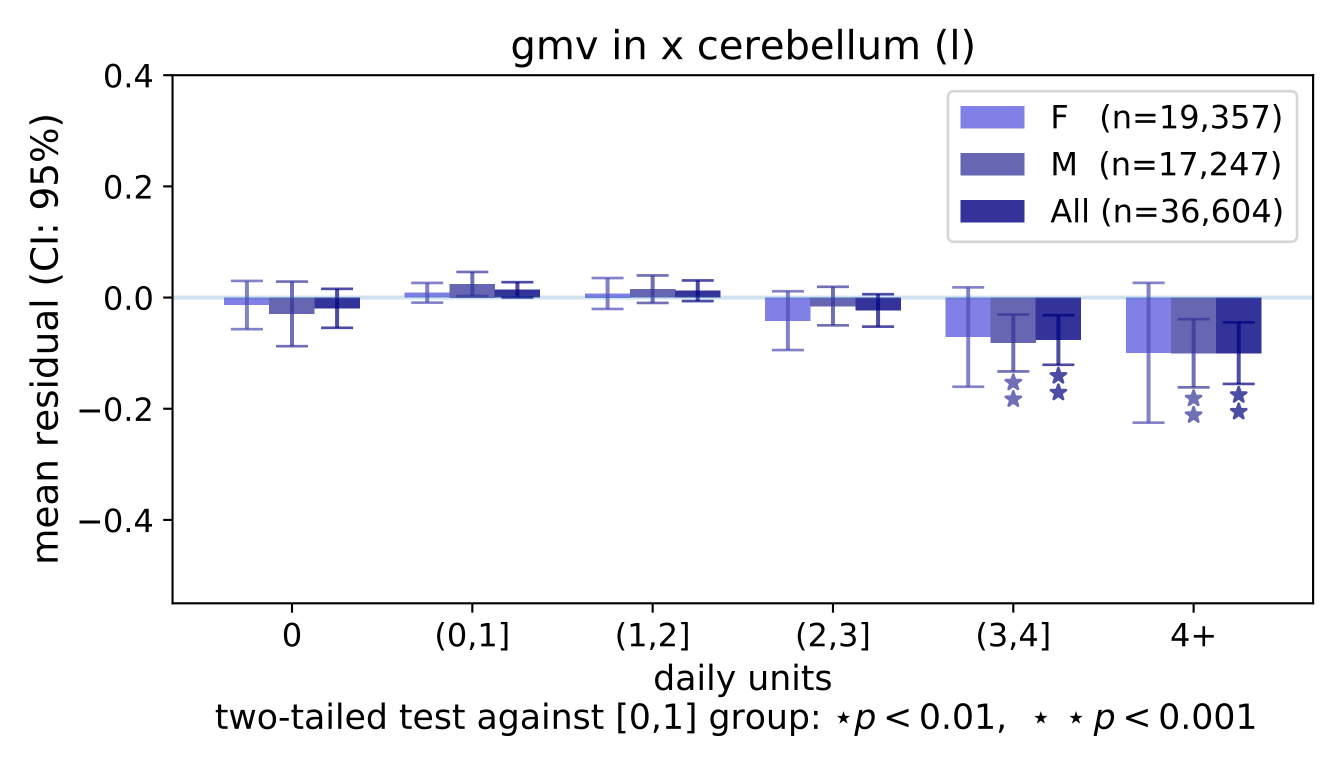

gmv in x cerebellum (l)
0.4
0.2
0.0
−0.2
−0.4
mean residual (CI: 95%)
F (n=19,357)
M (n=17,247)
All (n=36,604)
0
(0,1]
(1,2]
(2,3]
(3,4]
4+
daily units
two-tailed test against [0,1] group: ⋆$p < 0.01$, ⋆ ⋆ $p < 0.001$

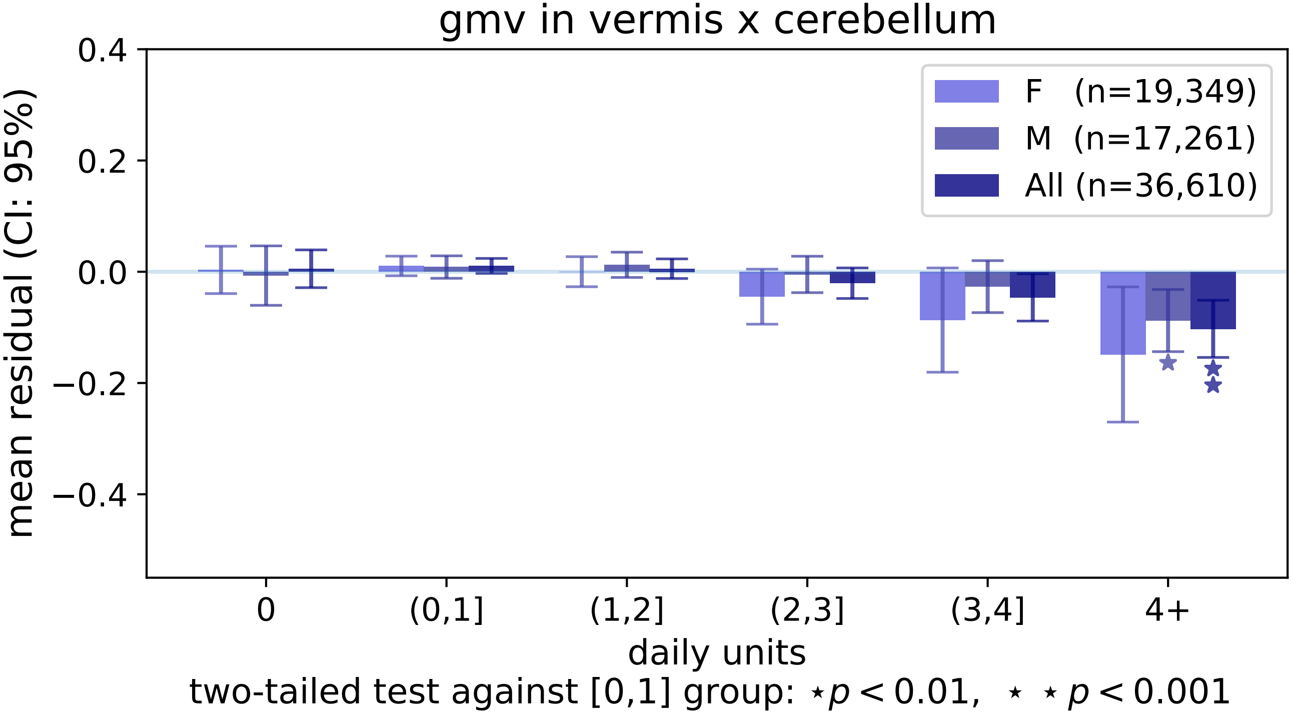

gmv in vermis x cerebellum
0.4
0.2
0.0
−0.2
−0.4
mean residual (CI: 95%)
F (n=19,349)
M (n=17,261)
All (n=36,610)
0
(0,1]
(1,2]
(2,3]
(3,4]
4+
daily units
two-tailed test against [0,1] group: ⋆ p < 0.01, ⋆ ⋆ p < 0.001

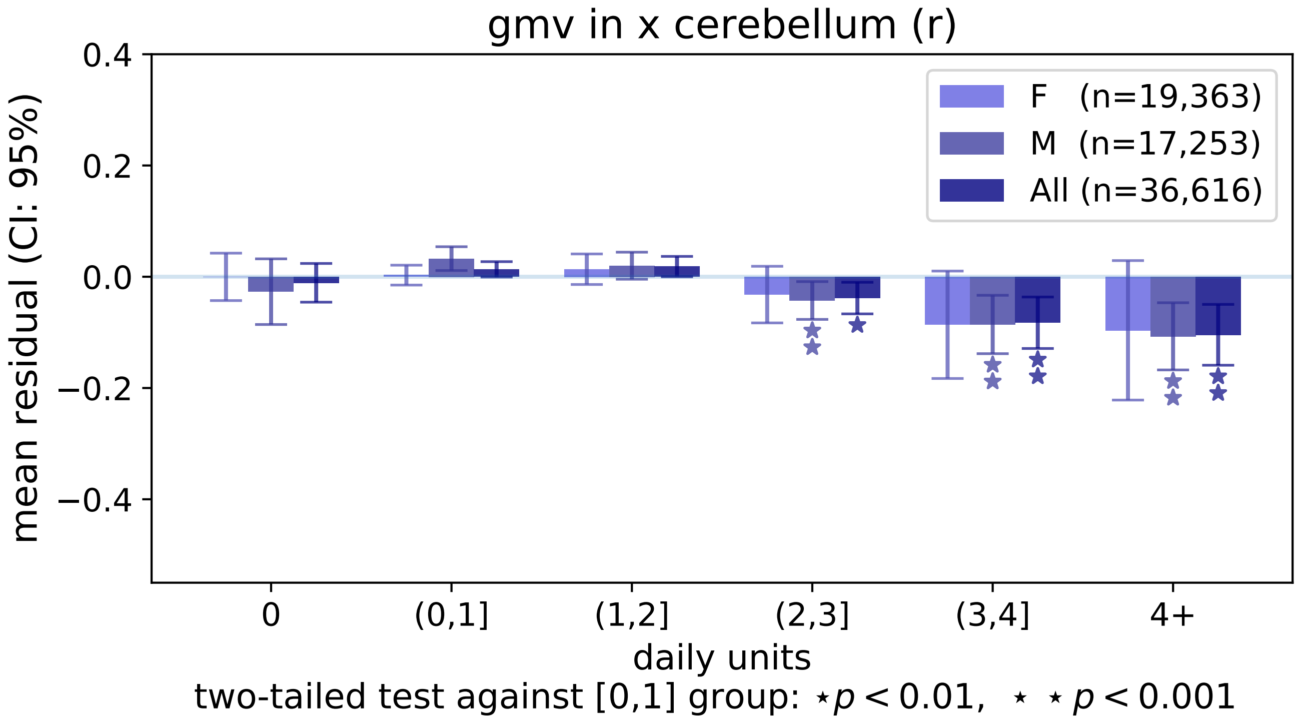
gmv in x cerebellum (r)
0.4
0.2
0.0
−0.2
−0.4
mean residual (CI: 95%)
F (n=19,363)
M (n=17,253)
All (n=36,616)
0
(0,1]
(1,2]
(2,3]
(3,4]
4+
daily units
two-tailed test against [0,1] group: ⋆ $p < 0.01$, ⋆ ⋆ $p < 0.001$
